# Macrocyclization of Broad-Spectrum Kinase Inhibitor Bosutinib leads to Potent and Selective Quinoline-based HIPK4 Inhibitor AZ137

**DOI:** 10.64898/2026.04.22.720179

**Authors:** Athina Zerva, Nicolai D. Raig, Zaile Zhuang, Andreas Krämer, Johannes Dopfer, Riley Togashi, Martin P. Schwalm, Lewis Elson, Julia M. Frischkorn, Benedict-Tilman Berger, Susanne Müller, James K. Chen, Stefan Knapp, Thomas Hanke

## Abstract

Homeodomain-interacting protein kinase 4 (HIPK4) remains an understudied member of the dark kinome. While genetic knockout studies suggest roles for HIPK4 in spermiogenesis and cutaneous squamous cell carcinoma, whether these cellular functions can be recapitulated by pharmacological inhibition remains to be determined. However, such investigations have been hampered by a lack of high-quality chemical tools. To address this, we employed a rational design strategy utilizing macrocyclization of a bosutinib-based scaffold. Systematic optimization led to the discovery of **AZ137** (**28e**), a potent and selective HIPK4 inhibitor (IC_50_ = 11 nM; cellular EC_50_ = 76 nM). **AZ137** exhibits exceptional selectivity across three comprehensive orthogonal panels, high solubility, and no detectable cytotoxicity. Its cellular activity was confirmed in cell-based assays of HIPK4-dependent F-actin remodeling. Together with a negative control compound, this probe set provides a foundational framework for the validating HIPK4 as a therapeutic target and a high-quality resource to elucidate its roles in normal physiology and disease.

## 1. INTRODUCTION

Homeodomain-interacting protein kinases (HIPKs) are a small family of serine/threonine kinases consisting of four members: HIPK1, HIPK2, HIPK3, and HIPK4.^1,2^ HIPK1–3 were originally discovered because of their interaction with homeodomain transcription factors.^3^ The fourth member, HIPK4, was later identified based on sequence homology of its kinase domain with the other HIPKs.^1^ While the HIPK family shows the highest sequence similarity to CMGC kinases known to regulate RNA-splicing, including the serine-arginine protein kinases (SRPKs), dual-specificity tyrosine-phosphorylation-regulated kinases (DYRKs) and Cdc2-like kinases (CLKs) (**Figure 1A**), HIPKs play distinct regulatory roles in cellular signaling.^1^ HIPK family members have been described regulating diverse cellular processes, such as cell proliferation, cell differentiation, cell survival and cell death. HIPK1–3 regulate these processes by phosphorylating homeodomain transcription factors, p53, and other key regulatory proteins.^3,4^ HIPK1 and HIPK2 have a in tumor suppressor role by inducing apoptosis triggered by genotoxic stress and stimulation of aberrant growth control pathways.^5–10^ HIPK2 has also been shown to promote apoptosis by phosphorylating and downregulating the transcriptional co-repressor C-terminal binding protein (CtBP1), while HIPK3 is associated with the regulation of FAS-mediated apoptotic signaling.^11,12^

**Figure 1.**
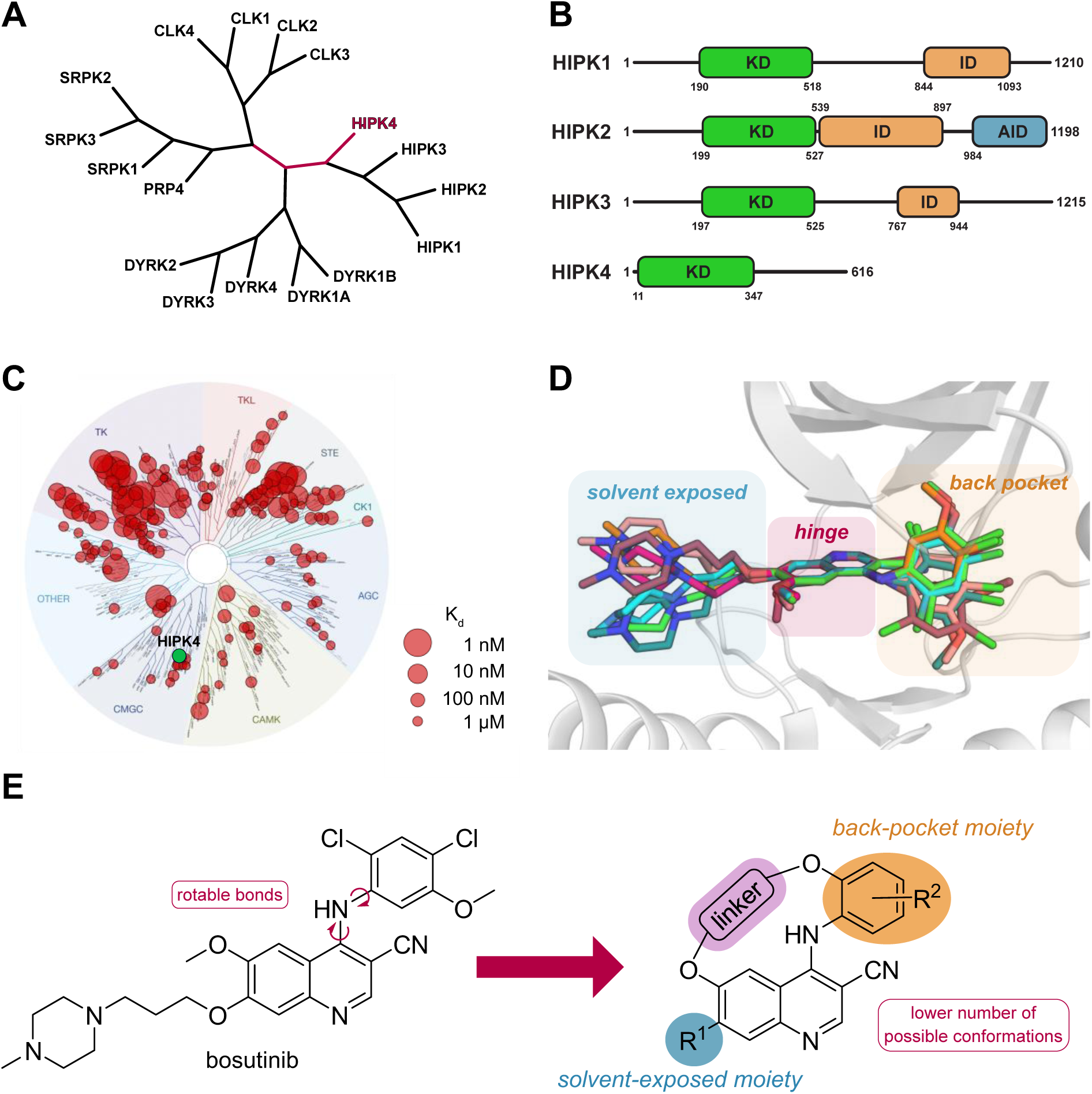
Overview of the chemical and structural foundation for the design of bosutinib-derived macrocyclic inhibitors. **A** Phylogenetic human kinome tree of the splicing kinase subfamilies, highlighting the evolutionary relationship of HIPK4 to its homologues and the closest related CMGC group of kinases. The tree was generated from an alignment of the respective kinase domains (Newick format) and visualized using iTOL^26^. **B** Schematic domain organization of human HIPK1–4 showing the kinase domains (KD), composite interaction regions (ID) and autoinhibitory domains (AID). Domain annotation and boundaries were derived from UniProt. **C** Dendrogram of bosutinib illustrating binding affinity to screened kinases. As indicated in the legend, the size of the circles reflects the K_d_ values of bosutinib binding to the respective kinase. The image was generated online via the TREEspot™ Software Tool from Eurofins. Selectivity data were taken from a study profiling kinase inhibitor selectivity.^18^ **D** Superimposition of the bioactive conformations of bosutinib in available crystal structures. PDB: 4QMN (orange, MST3), 3SOA (maroon, CaMKII), 6OP9 (hotpink, HER3), 5VC3 (salmon, WEE1), 5AJQ (cyan, STK10), 3UE4 (green, ABL1), 6FDY (petrol, ULK3). **E** Design strategy for the development of bosutinib-based macrocyclic inhibitors. To study the structure–activity relationship, the pharmacophore was modified at three positions: the linker (violet), the solvent-exposed moiety R^1^ (blue) and the back-pocket moiety R^2^ (orange).

In contrast, HIPK4 remains the least characterized member of the family and differs significantly from the other members based on its primary sequence and cellular function. Its kinase domain, exhibits only about 50% sequence identity to HIPK1–3, whereas the kinase domains of HIPK1–3 share approximately 90% sequence identity. HIPK4 also lacks the homeobox-interacting domain that defines the other HIPK family members (**Figure 1B**), and mainly resides in the cytoplasm, while HIPK1–3 are predominantly localized in the cell nucleus.^2,3,7,13^ Together with the structural divergence, this suggests that HIPK4 may have evolved to perform distinct functional roles. Unlike HIPK1–3, which are conserved in all vertebrates, HIPK4 is only expressed in mammals. Recent studies have suggested that HIPK4 is involved in a number of biological processes. In knockout mice, the loss of HIPK4 leads to male infertility, and *Hipk4* mutant sperm showed reduced oocyte binding and impaired fertilization capacity *in vitro*.^14^ Examinations using light and electron microscopy of HIPK4-null male germ cells revealed abnormal head morphologies in mature spermatozoa and defects in the acroplaxome, a filamentous actin (F-actin)- and keratin 5-scaffolded structure that is crucial for shaping the sperm nucleus, as well as alterations in the localization of actin-regulating proteins. This suggests that HIPK4 plays a role in regulating actin-based structures in sperms and supports an involvement of HIPK4 in cytoskeletal remodeling.^14^ Phosphoproteomic analyses of Hipk4^−/−^ testes further revealed reduced phosphorylation of manchette-associated proteins such as RIMS-binding protein 3.^15^ In addition to these spermatogenic functions, Larribère *et al.* linked HIPK4 to epithelial differentiation, whereby HIPK4 knockdown in keratinocyte models led to accelerated maturation, suggesting that HIPK4 serves as a negative regulator of differentiation.^16^ Further studies linked HIPK4 to cutaneous squamous cell carcinoma, where increased expression of HIPK4 correlated with higher cell proliferation and invasion, an effect that is likely through the phosphorylation of the transcription factor TAp63 and suppression of the extracellular matrix protein EFEMP1.^17^ Taken together, these findings suggest that HIPK4 is involved in reproduction, tissue differentiation, and tumor biology, but the precise molecular mechanisms mediating these effects, its substrates, and upstream regulators remain largely unknown. The lack of selective chemical tools further limits functional studies aiming to decipher the cellular roles of HIPK4, and a deeper understanding of the HIPK4 signaling network. To close this gap, the development of a potent and selective chemical tool for HIPK4 is urgently needed. While a patent by Vibliome Therapeutics LLC recently described potential HIPK4 inhibitors, no peer-reviewed data or characterized probes have been disclosed to date. However, several multi-kinase inhibitors, such as foretinib (K_d_ = 0.5 nM)^18^, sunitinib (K_d_ = 160 nM)^19^, and bosutinib (K_d_ = 130 nM)^18^, show strong binding to HIPK4 in biochemical assays, these inhibitors are highly promiscuous and unsuitable for investigating HIPK4-specific biology. Bosutinib is an ATP-competitive type I kinase inhibitor approved for the treatment of chronic myelogenous leukemia.^20^ A kinome-wide profiling reveals that bosutinib is a promiscuous kinase inhibitor that, in addition to its primary targets BCR-ABL and SRC, also binds potently to many other kinases (**Figure 1C**). Currently 15 bosutinib co-crystal structures are available in the PDB revealing its interaction with 12 different kinases. Superimposition of these co-crystal structures of bosutinib revealed that this linear molecule can adopt several distinct orientations, reflecting its notable conformational flexibility (**Figure 1D**). Given bosutinib’s broad kinome-wide activity, its conformational flexibility represents a structural hallmark that could be leveraged to enhance target selectivity.

Macrocyclization is a powerful and increasingly popular strategy for limiting small molecule flexibility.^21^ By restricting the conformation of linear ligands, macrocyclization can reduce the entropic cost of binding, stabilize target-specific bioactive conformations and enhance selectivity and affinity.^22^ Several FDA-approved macrocyclic kinase inhibitors, including lorlatinib (ALK/ROS1) and pacritinib (JAK2), have demonstrated that macrocyclization can overcome pharmacological limitations of linear parent inhibitors by for instance improving pharmacokinetics, cellular potency, selectivity or activity against resistance mutations.^23–25^ In particular, the ability of macrocycles to limit off-target interactions while maintaining or even enhancing on-target binding makes this compound class a valuable platform for the development of highly selective ligands. Guided by this rationale, we used macrocyclization as a strategy to restrict the conformational flexibility of the bosutinib scaffold and thereby improve its selectivity (**Figure 1E**).

Building on this approach, we synthesized a series of macrocyclic bosutinib derivatives and evaluated their kinome-wide profiles. This systematic screening revealed that the quinoline-based macrocycles exhibited an unexpected and exceptional affinity for HIPK4. Subsequent iterative optimization of the structure-activity relationship (SAR) led to the identification of **AZ137**, a potent and highly selective inhibitor that serves as a valuable chemical tool for studying HIPK4 function in a cellular context.

## 2. RESULTS AND DISCUSSIONS

### 2.1 Conformational flexibility drives bosutinib’s promiscuity

Analysis of published co-crystal structures of bosutinib bound to various kinases revealed that bosutinib bound with a canonical type I binding mode. The 3-cyanoquinoline moiety serves as the hinge-binding motif, a feature that remained remarkably conserved among all structures, while the aniline group pointed towards the hydrophobic back pocket and the piperazine ring was solvent exposed. The structural alignment of bosutinib crystal structures showed that acyclic bosutinib can adopt many different conformations. In particular, the aniline moiety of bosutinib was highly flexible due to the rotatable N–phenyl bond (**Figure 1D**). We hypothesized that the flexibility of the aniline group was primarily responsible for the promiscuous behavior of bosutinib, allowing it to adapt to diverse kinase backpockets. Based on our observations, we designed macrocyclic bosutinib derivatives with restricted conformational flexibility to improve selectivity. The 4-(phenylamino)quinoline-3-carbonitrile formed the core of our pharmacophore model (**Figure 1E**) in order to preserve key binding interactions. As previous macrocycle projects have demonstrated the importance of ring size in capturing the bioactive conformation, we varied the linker length as one element of our SAR to identify the optimal macrocycle size (14 to 17 atoms).^27–29^ The attachment points for the linker moieties were chosen based on the structural analysis of bosutinib and our previous experience with quinazoline-based macrocycles at position 6 of the quinoline and the *ortho* position of the aniline.^27^ As *N*-propoxy-*N*-methylpiperazine, which was exposed to the solvent, did not demonstrate any significant kinase interactions in the analyzed co-crystal structures, we initially excluded this group from the iterative SAR process and compared analogs bearing a hydrogen versus a methoxy group. After optimizing the linker length, we examined the substitution pattern of the back-pocket-targeting moiety. Based on prior observations that tailored the macrocyclic scaffold towards the back pocket can result in exceptional selectivity, we aimed to further explore these interactions by introducing ring systems with different substitution patterns.^30^

### 2.2 Synthesis of macrocyclic 3-Cyanoquinolines

Depending on the substitution pattern at the solvent-exposed moieties, distinct synthetic routes were employed to access the final macrocyclic scaffolds (R^1^, **Figure 1E**). Macrocycles **8a**–**d**, bearing a hydrogen atom at position 7 of the quinoline, were synthesized following a seven-step route starting from the commercially available 2-amino-5-methoxybenzoic acid **1** (**Scheme 1**).

Benzoic acid **1** was converted to amidine **2** using dimethylformamide dimethyl acetal. The quinoline scaffold was then assembled using a ring-closing reaction with **2** and cyanomethanide, which was prepared *in situ* by deprotonation of acetonitrile with *n*-butyllithium. The resulting hydroxyl group present in quinoline **3** was chlorinated with POCl_3_ and the methoxy group at position 6 was subsequently demethylated, yielding the quinoline building block **5**. The cumulative yield of the four-step synthesis of **5** amounted to 69%. Starting from this shared intermediate, macrocyclic derivatives **8a**–**d** were obtained in three subsequent synthetic steps, incorporating linkers of various lengths. First, the hydroxyl linkers were attached via Mitsunobu reaction conditions, followed by a nucleophilic aromatic substitution at the 4-position of the quinoline with 2-aminophenol, yielding the linear precursor compounds **7a**–**d**. The final macrocyclization involved an intramolecular nucleophilic substitution between the phenol and the halogenated linker, yielding the macrocycles **8a**–**d**.

**Scheme 1.** Seven-step synthesis of macrocycles **8a**–**d**^a,b^

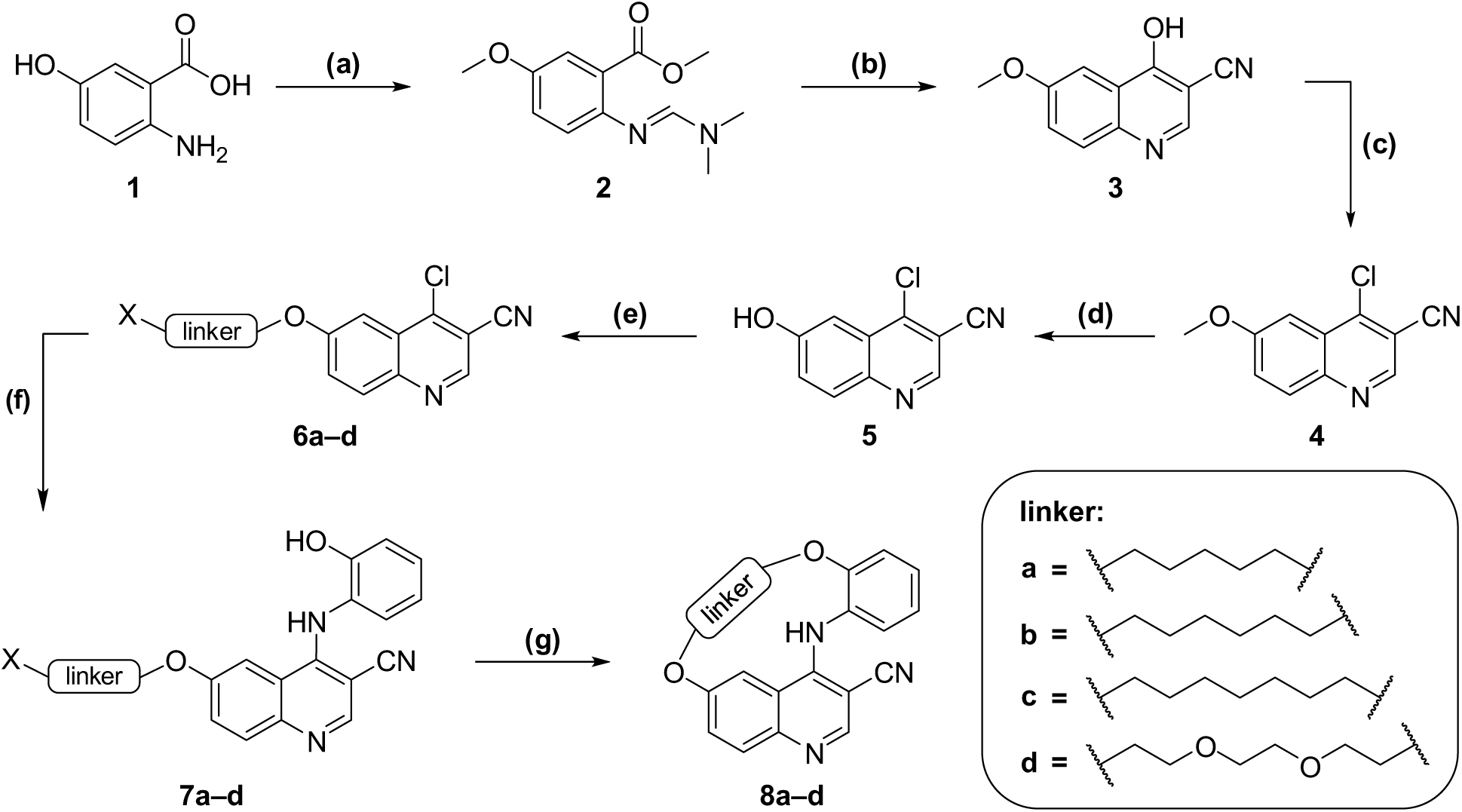

^a^Reactions and conditions: **(a)** DMF-dimethyl acetal, DMF, 150 °C, 3 h; **(b)** ACN, *n*-BuLi, THF, –78 °C to RT, 15 h; **(c)** POCl_3_, toluene, 120 °C, 4 h; **(d)** BCl_3_, TBAI, toluene, 110 °C, 16 h; **(e)** hydroxy linker, DIAD, PPh_3_, toluene/DMF, 90 °C, 18 h; **(f)** 2-aminophenol, HCl, *i*-PrOH, 100 °C, 16 h; **(g)** NaH, DMF, 60 °C, 18 h. X = Br or Cl. ^b^The synthesis of macrocycle **8a** differed slightly in the final three steps and is described in detail in the Experimental Section.

The macrocycles **23a–28e**, bearing a methoxy group at position 7 of the quinoline, were accessed through a different nine-step synthetic route (Scheme 2). To enable a late-stage derivatization of the aniline moieties and linkers the core quinoline building block **16** was synthesized on a larger scale. The seven-step synthesis was conducted according to the patented method of Heymach *et al.*^31^, starting with a Fischer esterification of commercially available 5-hydroxy-4-methoxy-2-nitrobenzoic acid **9**. The phenol was then protected with a benzyl group via a Williamson ether synthesis, followed by reduction of the nitro group under Béchamp conditions, yielding the aniline **11**. The quinoline scaffold was assembled using a ring-closing reaction utilizing cyanomethanide prepared *in situ,* before the resulting phenol was chlorinated. Finally, the quinoline building block **16** was obtained by deprotecting the 6-hydroxyl group, resulting in an overall yield of 21% over the seven steps. Starting from **16** the macrocycles **23a**–**28e** were obtained in two additional derivatizing steps. First, the acyclic precursor compounds **17**–**22** were obtained via an aromatic nucleophilic substitution using respective 2-aminophenol derivatives. Finally, double nucleophilic substitution between precursors **17**–**22** and various dibromo linkers yielded macrocycles **23a**–**28e**. To minimize oligomerization and the formation of by-products, all ring closing reactions were carried out under highly diluted conditions.

**Scheme 2.** Nine-step synthesis of macrocycles **23a–28e**^a^

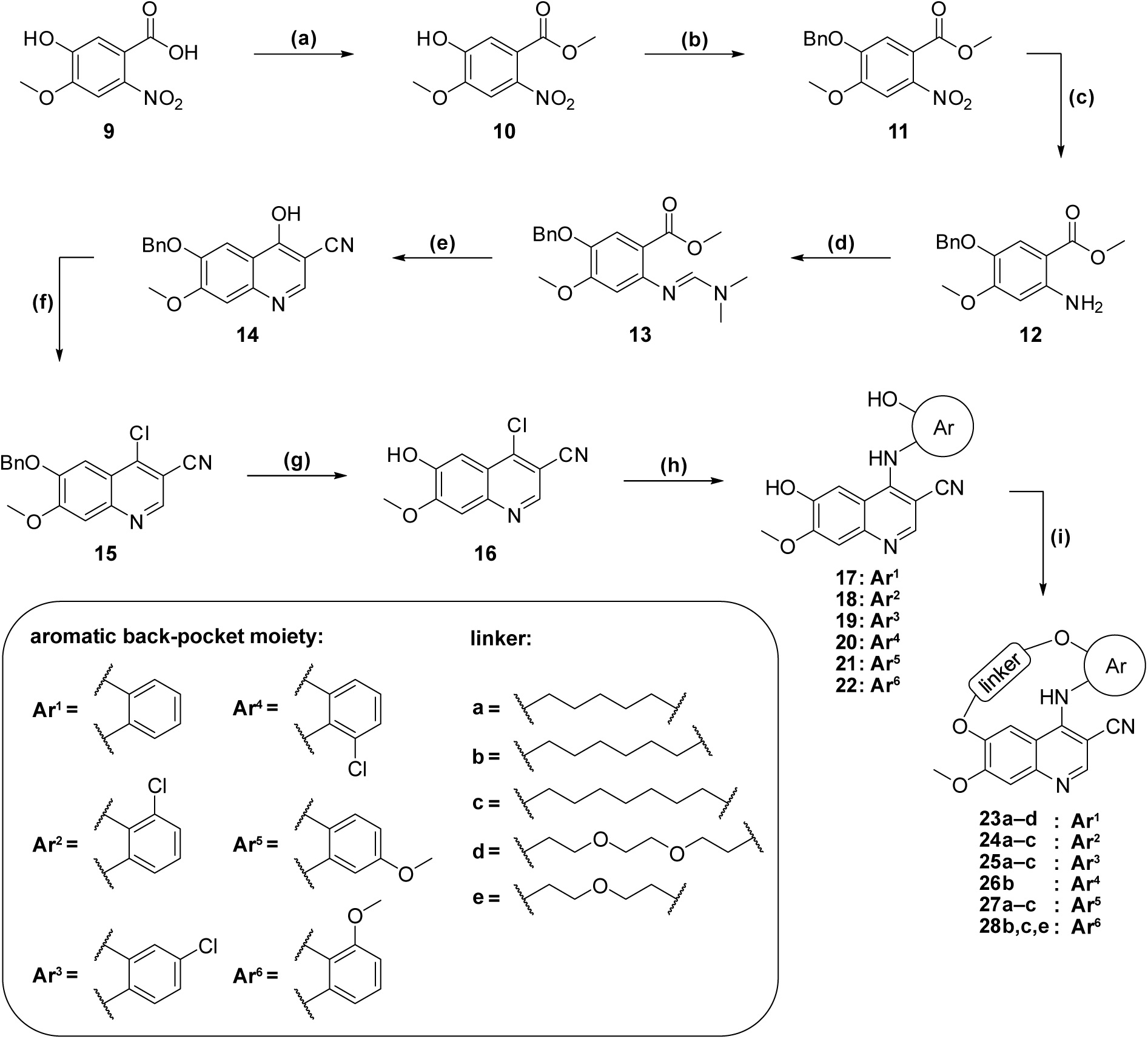

^a^Reactions and conditions: **(a)** MeOH, H_2_SO_4_, 80 °C, 16 h; **(b)** BnBr, K_2_CO_3_, DMF, 80 °C, 16 h; **(c)** Fe, NH_4_Cl, MeOH/H_2_O, 80 °C, 16 h; **(d)** DMF-dimethyl acetal, DMF, 110 °C, 5 h; **(e)** ACN, *n*-BuLi, THF, –78 °C to RT, 16 h; **(f)** POCl_3_, toluene, 120 °C, 18 h; **(g)** PhSMe, TFA, 80 °C, 3 h; **(h)** 2-aminophenol, HCl, 2-ethoxyethanol, 130 °C, 40 min (mw); **(i)** dibromo-linker, K_2_CO_3_, DMF, 80 °C, 2–4 h.

### 2.3 *In vitro* and *in cellulo* Structure–Activity Relationship study of 3-Cyanoquinolines

To assess the potential of these compounds as selective HIPK4 inhibitors, we simultaneously evaluated three key parameters. The *in vitro* potency was determined using a dose-response ADP-Glo™ assay using purified HIPK4, allowing the calculation of median inhibitory concentration (IC_50_) (**Table 1–2**). Cellular target engagement was assessed by overexpressing HIPK4-NanoLuciferase fusion constructs in HEK293T cells using NanoBRET^®^ technology to determine cellular half-maximal effective concentration (EC_50_) (**Table 1–2**). Finally, the kinome-wide selectivity of the most promising compounds (IC_50_ < 0.5 µM and EC_50_ < 1 µM) was evaluated via differential scanning fluorometry (DSF) assay against a representative panel of approximately 100 kinases.^32^ This panel also included common non-kinase off-targets of kinase inhibitors, such as bromodomains^33^ (**Table S3**). Selectivity was quantified using the S(3K) score, which reflected the fraction of kinases stabilized by Δ*T_m_* ≥ 3 K upon compound binding (**Table 1–2**).

**Table 1.** Structure–Activity Relationships of bosutinib inspired macrocycles **8a**–**d**, **23a**–**e**. The effect of quinoline-C7-substituent R (blue) and linker variation (violet) on the inhibitory effect and target engagement against HIPK4 and kinome selectivity of the macrocycles were compared with bosutinib.^a^

| comp. ID | R | linker | HIPK4 |  | S(3K) |
| --- | --- | --- | --- | --- | --- |
|  |  |  | *in vitro*<br>IC<sub>50</sub> [nM] | *in cellulo*<br>EC<sub>50</sub> [nM] |  |
| **bosutinib** |  |  | 330 | 300 | 0.35 |
| **8a** |  |  | 230 | 2360 | n.d. |
| **8b** |  |  | 110 | 400 | 0.12 |
| **8c** |  |  | 65 | 359 | 0.03 |
| **8d** |  |  | 520 | >5000 | n.d. |
| **23a** |  |  | 23 | 258 | 0.14 |
| **23e** |  |  | 8.9 | 71 | 0.23 |
| **23b** |  |  | 6.7 | 66 | 0.21 |
| **23c** |  |  | 2.2 | 46 | 0.18 |
| **23d** |  |  | 84 | 649 | 0.14 |

<sup>a</sup>*In vitro* IC<sub>50</sub> values were measured by ADP-glo assay. The EC<sub>50</sub> values were determined by a NanoBRET assay in intact cells (two biological replicates, each measured in duplicates). The selectivity (S) was calculated as the number of kinases showing a thermal shift larger than 3 K in a DSF assay divided by the number of kinases screened (approx. 100 kinases). Tabulated replicate data and associated statistical parameters are provided in the Supporting Information (**Table S1–3**).

To validate our pharmacophore model for the development of HIPK4 inhibitors, we designed an initial series of bosutinib-inspired macrocycles **8a**–**d**, **23a**–**d** (**Table 1**). This set of compounds was specifically designed to investigate the influence of the C7 substituent of the quinoline scaffold while simultaneously identifying the optimal linker length. To isolate the effects of these two variables, an unsubstituted 2-aminophenol moiety was employed as the back-pocket targeting motif across this series.

Compounds **8a**–**d**, characterized by the absence of a substituent at C7 of the quinoline scaffold (**Table 1**), showed submicromolar inhibition of HIPK4 in the ADP-Glo assay. Within these series, compounds **8b** (C6-linker) and **8c** (C7-linker) demonstrated the highest potency with IC_50_ values of 65–110 nM, which translated into a submicromolar cellular target engagement (**Table 1**). In contrast, compounds with C5-linker (**8a**) and PEG2-linker (**8d**) showed poor cellular target engagement with EC_50_ values of 2.36 µM and >5 µM, respectively. Moreover, compounds **8b** and **8c** demonstrated favorable potency and an unexpectedly high level of kinome-wide selectivity, with S(3K) values of less than 0.12.

In compounds **23a**–**e**, we introduced a methoxy group at C7 of the quinoline scaffold (**Table 1**). This modification resulted in much better *in vitro* potency with IC_50_ values ranging from 2 nM to 84 nM. This series confirmed the previous observation that C6- (**23b**) and C7-(**23c**) linkers are the most potent and cellularly active compounds, with EC_50_ values of 66 nM and 46 nM. In addition, a PEG1-linker in compound **23e** was very well tolerated with an EC_50_ value of 71 nM. Although the selectivity was slightly lower for **23a–e** (S(3K) = 0.14–0.23) than for **8b** and **8c**, the increased potency highlights the importance of the C7-methoxy modification at the quinoline and supports further optimization of this series.

In summary, the first series of compounds (**8a**–**d**, **23a**–**e**; **Table 1**) revealed that our chosen pharmacophore was a promising scaffold for the development of potent and selective HIPK4 inhibitors. In particular, **8b–c** and **23a**–2**3c** outperformed bosutinib in inhibiting HIPK4. Furthermore, the methoxy group in the C7 position of the quinoline ring had a crucial impact on the potency of the inhibitors, while the optimal linker length appeared to differ between PEG1, C6 and C7. PEG2-linkers, however, were poorly tolerated. Despite employing an unsubstituted phenyl ring, this initial series of macrocycles demonstrated promising selectivity, showing a marked reduction in off-targets activity compared to the highly promiscuous linear parent compound, bosutinib.

Based on SAR insights from the first series of macrocycles (**Table 1**), we developed a second series (compounds **24a**–**28e)** (**Table 2**). This new series retained the methoxy group at the C7-position of the quinoline and primarily utilized a linker length of 5–7-atoms. In this second set of compounds, we explored various substitution patterns on the back-pocket targeting phenyl ring to improve selectivity without compromising HIPK4 potency.

**Table 2.**
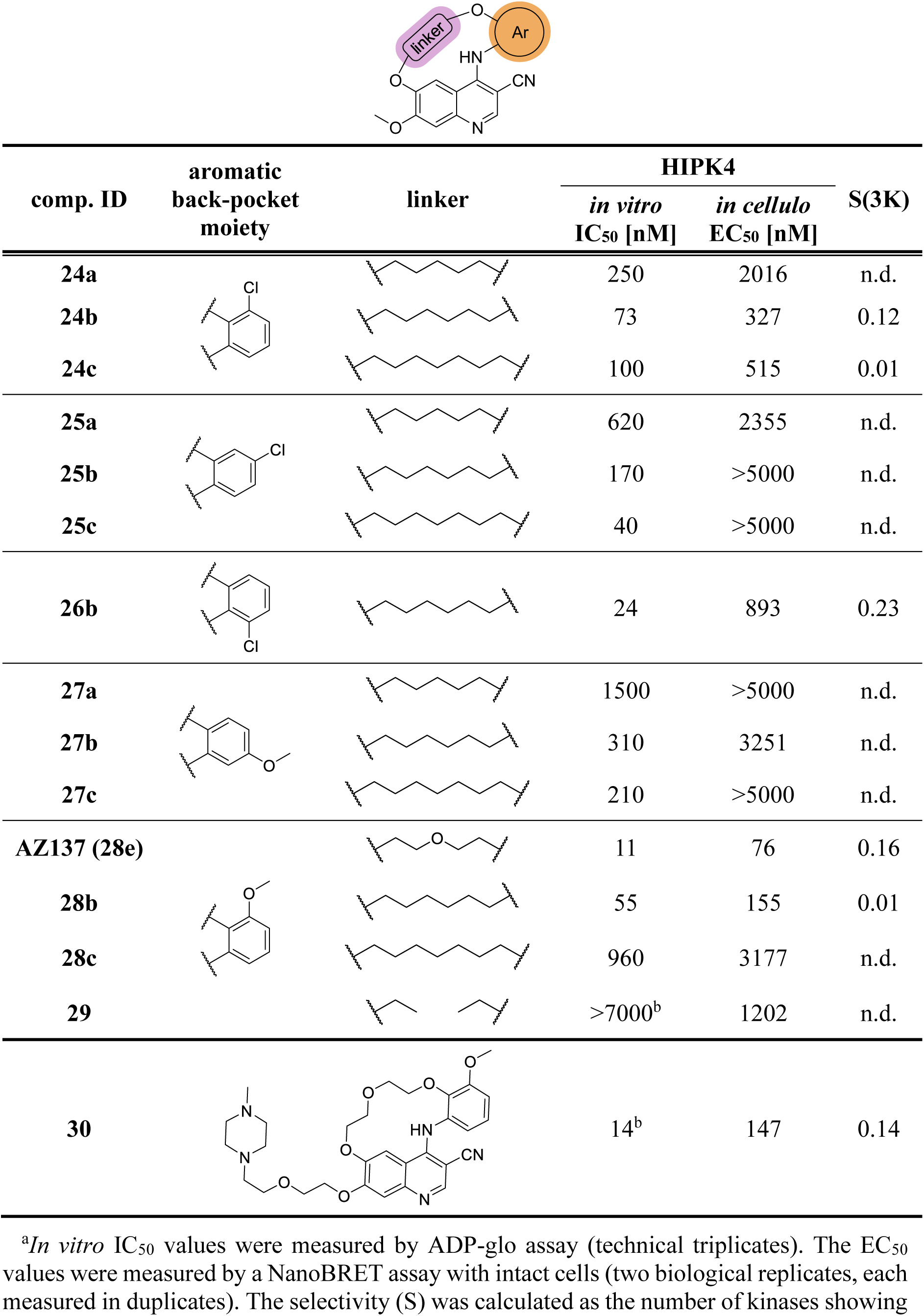
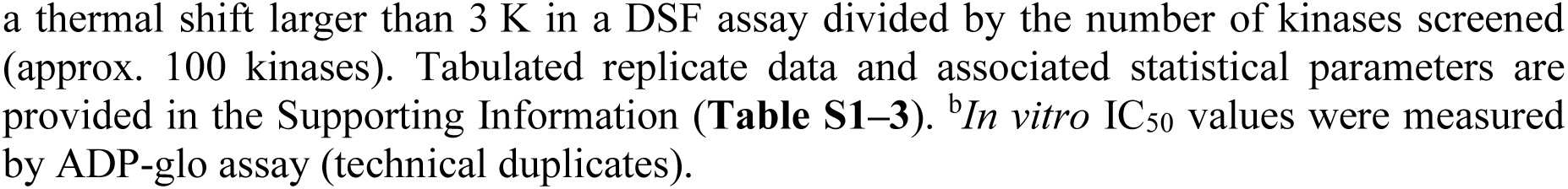
Structure–Activity Relationships of bosutinib inspired macrocycles **24a**–**28e**. The effect of aromatic back-pocket motif (orange) and linker variation (violet), on the inhibitory effect and target engagement against HIPK4 and kinome selectivity of the macrocycles were compared.^a^

| comp. ID | aromatic back-pocket moiety | linker | HIPK4 |  | S(3K) |
| --- | --- | --- | --- | --- | --- |
|  |  |  | *in vitro*<br>IC<sub>50</sub> [nM] | *in cellulo*<br>EC<sub>50</sub> [nM] |  |
| **24a** |  |  | 250 | 2016 | n.d. |
| **24b** |  |  | 73 | 327 | 0.12 |
| **24c** |  |  | 100 | 515 | 0.01 |
| **25a** |  |  | 620 | 2355 | n.d. |
| **25b** |  |  | 170 | >5000 | n.d. |
| **25c** |  |  | 40 | >5000 | n.d. |
| **26b** |  |  | 24 | 893 | 0.23 |
| **27a** |  |  | 1500 | >5000 | n.d. |
| **27b** |  |  | 310 | 3251 | n.d. |
| **27c** |  |  | 210 | >5000 | n.d. |
| **AZ137 (28e)** |  |  | 11 | 76 | 0.16 |
| **28b** |  |  | 55 | 155 | 0.01 |
| **28c** |  |  | 960 | 3177 | n.d. |
| **29** |  |  | >7000<sup>b</sup> | 1202 | n.d. |
| **30** |  |  | 14<sup>b</sup> | 147 | 0.14 |

<sup>a</sup>*In vitro* IC<sub>50</sub> values were measured by ADP-glo assay (technical triplicates). The EC<sub>50</sub> values were measured by a NanoBRET assay with intact cells (two biological replicates, each measured in duplicates). The selectivity (S) was calculated as the number of kinases showing

To systematically investigate the influence of the back-pocket substitution pattern on potency and selectivity, we synthesized compounds incorporating chloro substituents at the aniline C3, C4, or C6 positions (**24a**–**26b**) and methoxy groups at C3 or C5 position (**27a**–**28e**), thereby mimicking the parent compound, bosutinib. All chloro-substituted compounds (**24a**–**26b**) exhibited potent *in vitro* inhibition of HIPK4 (IC_50_ = 24–620 nM), but only limited cellular target engagement, with the exception of **24b** and **24c**. The C3-chloro substituted derivates **24b** and **24c** achieved pronounced cellular target engagement with EC_50_ values of 327 nM and 515 nM, respectively, while maintaining favorable selectivity scores (S(3K) < 0.12).

A similar trend was observed for the methoxy-substituted compounds (**27a**–**28e**). Consistent with the chloro-substituted series, most methoxy-substituted analogues maintained high *in vitro* potency in the sub micromolar range. However, they largely failed to translate the measured enzymatic activity into cellular potency, with the exception of **28b** and **28e** (AZ137). The C3-methoxy substituted derivatives **28b** and AZ137 (**28e**) demonstrated excellent potency in both assays (IC_50_ = 55 nM/11 nM; EC_50_ = 155 nM/76 nM) with promising selectivity profiles with a S(3K) < 0.16.

In summary, compounds featuring a hexyl or PEG1 linker in combination with a C3-substituted aniline exhibited the most robust HIPK4 inhibition and an exceptional selectivity profile compared to the parent compound bosutinib. In particular, three macrocycles (**24b**, **28b** and AZ137 (**28e**)) emerged as lead candidates, demonstrating an optimally balanced profile of high potency to HIPK4 and excellent kinome-wide selectivity.

To further contextualize these SAR trends, two additional compounds were synthesized to probe specific structural features of the scaffold. Compound **29**, the acyclic analog of **28b** and AZ137 (**28e**), was designed to isolate and directly assess the impact of macrocyclization on HIPK4 potency. Additionally, compound **30**, a bosutinib-inspired derivative featuring an extended hydrophilic front-pocket substituent, was prepared to evaluate the influence of this motif on HIPK4 binding (**Table 3**). While **30** maintained potent biochemical potency (IC_50_ = 14 nM), robust cellular activity (EC_50_ = 147 nM), and an acceptable selectivity score (S(3K) = 0.12), compound **29** exhibited no measurable effect on HIPK4 inhibition in either biochemical (IC_50_ > 7000 nM) or cellular (EC_50_ = 1202 nM) assays. This dramatic difference strongly validated our macrocyclic design approach and highlighted the benefits of the macrocyclic scaffold.

### 2.4 Selectivity Profiling revealed excellent kinome-wide selectivity of AZ137 (28e)

Our initial DSF selectivity panel contained approximately 100 kinases. We further characterized the most promising macrocycles (**24b**, **28b**, AZ137 (**28e**)) along with compound **30** in a NanoBRET panel of 192 kinases^34^. This broader profiling was conducted to evaluate the cellular target engagement and selectivity of these compounds in a relevant environment (**Figure 2A**). We observed that, across this extended selectivity panel, macrocycles bearing a C3-substituted aniline had a pronounced preference for inhibiting HIPK4, with minimal activity on the other kinases of this panel. The chloro-substituted compound **24b** had residual activity for GAK, which consistently emerged as an off-target in our DSF screening (**Table S3**). This observation was consistent with our previous studies on kinase inhibitors, in which GAK was frequently identified as a highly promiscuous kinase.^35,36^ Notably, this liability was eliminated in the methoxy-substituted compounds, for which no GAK target engagement was observed.

**Figure 2.**
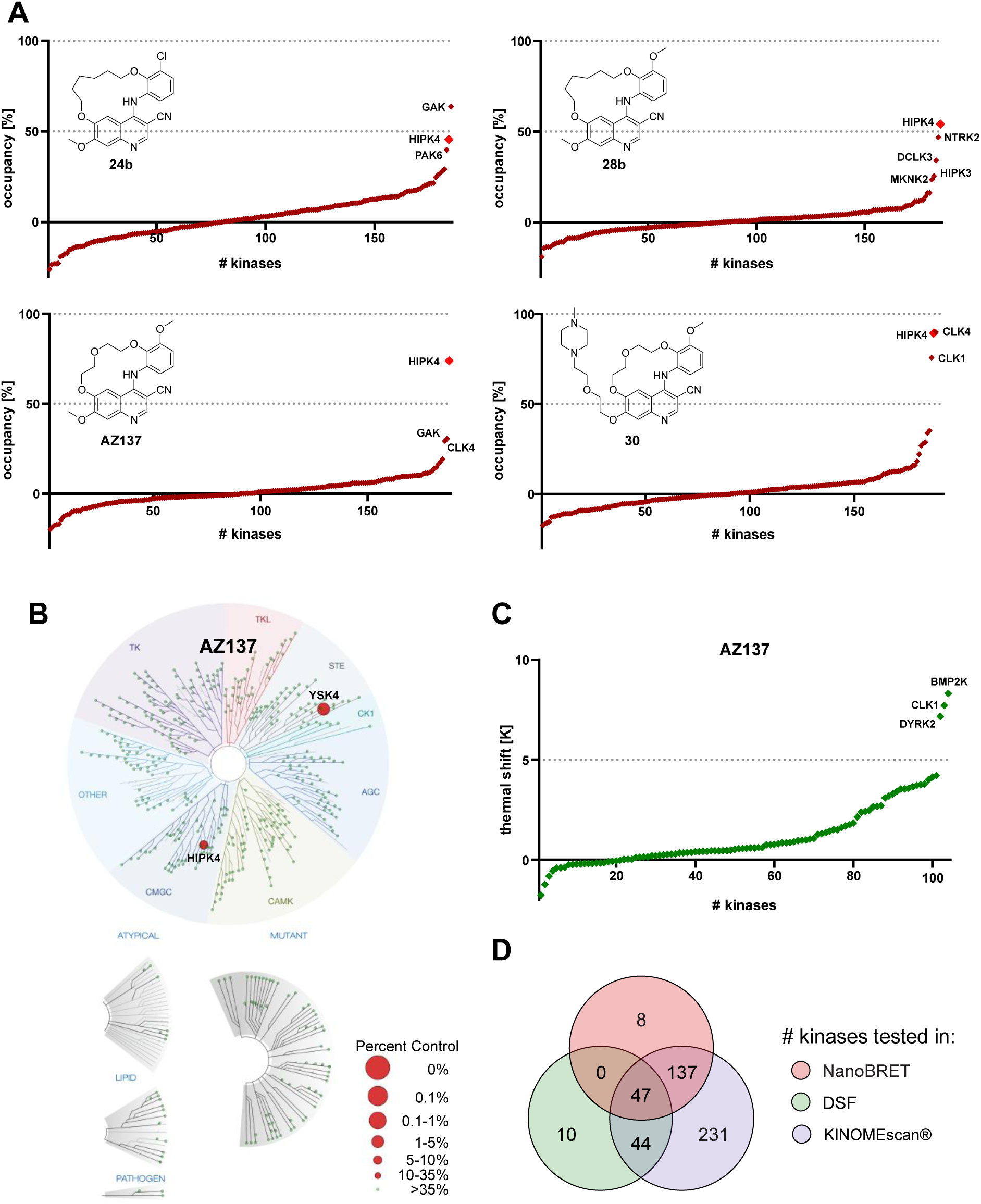
Macrocyclic HIPK4 inhibitors show excellent kinome-wide selectivity. **A** Cellular kinome selectivity profiles of compounds **24b**, **28b**, AZ137 (**28e**) and **30** assessed in a panel of 192 kinases with NanoBRET assays in HEK293T cells at 1 μM compound concentration. **B** Selectivity profile of AZ137 at 100 nM compound concentration using the Eurofins *scan*MAX Kinase Assay Panel of 468 kinases. The figure was generated using the TREEspot™ software tool. **C** Selectivity profiling of AZ137 (**28e**) in a DSF panel assay against 101 proteins at 1 μM compound concentration. Thermal shifts (Δ*T_m_*) are displayed as a waterfall plot. HIPK4 was not included in the panel. **D** Venn diagram illustrating protein overlap among the three selectivity panels. Tabulated data are provided in the Supporting Information (**Table S3-5**).

This observation suggested that the methoxy group hindered productive binding to GAK, while retaining high HIPK4 affinity. Of all the compounds tested, AZ137 (**28e**) was the most selective. HIPK4 was the only kinase with over 50% tracer displacement in the 192 kinase panel at 1 µM compound concentration (**Figure 2A**). Screening of the front-pocket extended analogue **30** also revealed good selectivity but this inhibitor also strongly bound to CLK1 and CLK4. This pattern aligns with the close structural similarity between the HIPK and CLK kinase subfamilies.

To confirm the K192 panel screening data and clarify whether the observed occupancy differences reflect genuine affinity differences within the HIPK and CLK families, dose-dependent NanoBRET assays were measured for AZ137 (**28e**) and **30** across HIPK1–4 and CLK1/2/4 (**Figure 3A/C**). Dose-response analysis for **30**, incorporating the extended front-pocket motif, corroborated the off-target profile revealed by the K192 screen. The data confirmed potent target engagement of HIPK4 (EC_50_ = 147 nM), but also activity for CLK1, and CLK4 (EC_50_ = 327 nM, and 619 nM, respectively) (**Figure 3A/C**).

**Figure 3.**
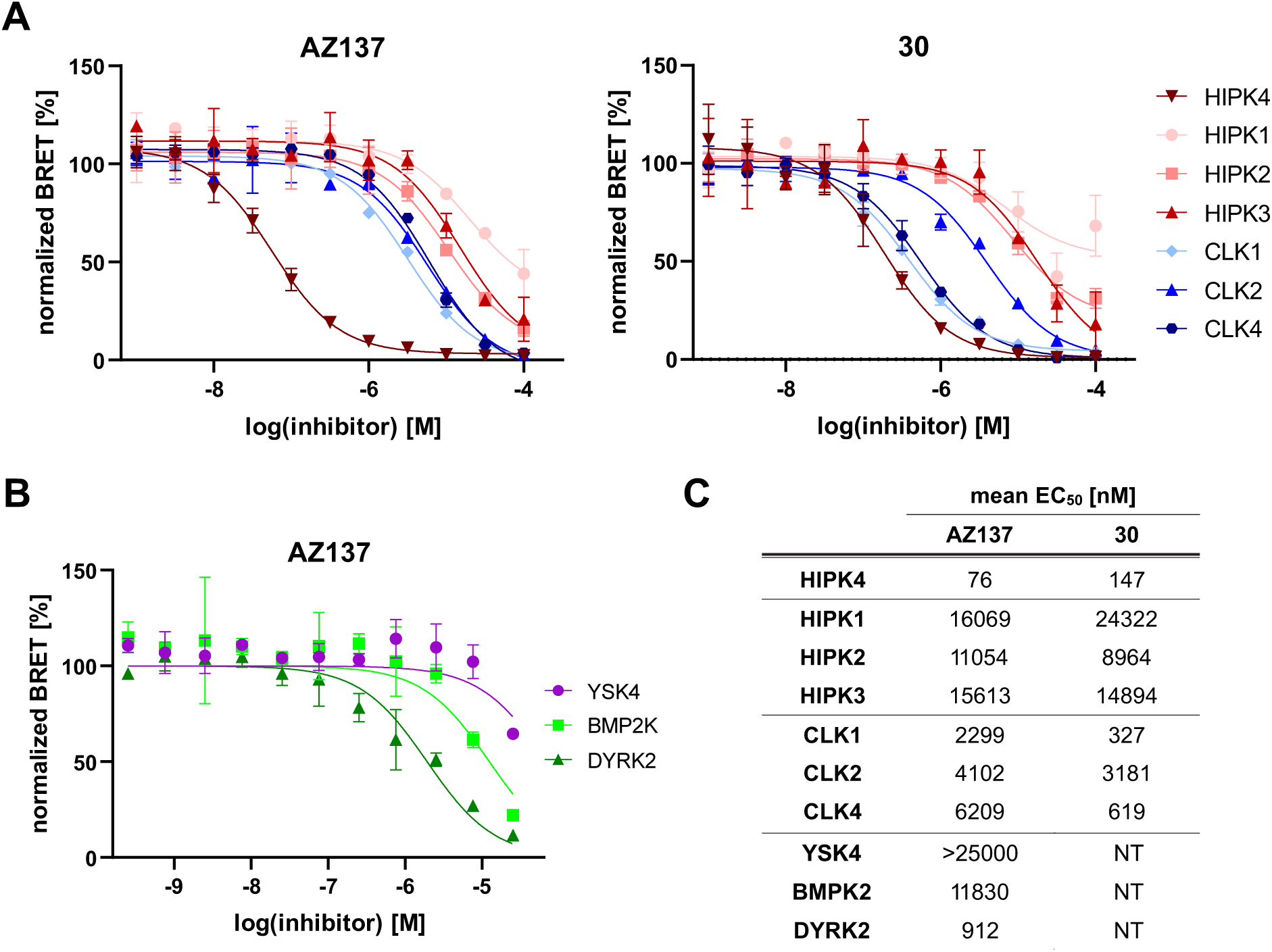
Cellular validation of kinases identified in selectivity profiling. **A** NanoBRET dose-response curves of AZ137 (**28e**) and **30** tested against HIPK1–4 (reds) and CLK1, CLK2 and CLK4 (blues). **B** NanoBRET dose-response curves of AZ137 (**28e**) determined for YSK4, DYRK2, and BMP2K, identified in the NanoBRET and DSF selectivity panels. **C** Table of mean EC_50_ values of AZ137 (**28e**) and **30** from the dose-response experiments shown in panel A and B, determined by NanoBRET in intact cells from two independent biological experiments, each performed in duplicate.

In contrast, off-target validation for AZ137 (**28e**) revealed clear isoform selectivity within the HIPK subfamily, demonstrating an EC_50_ of 76 nM for HIPK4, and over 100-fold selectivity over HIPK1–3 (EC_50_ = 11–16 µM) (**Figure 3A/C**). AZ137 (**28e**) exhibited a similarly pronounced selectivity window against the closely related CLK subfamily. With EC_50_ values ranging from 2.3 to 6.2 µM on CLK1, CLK2 and CLK4, AZ137 (**28e**) was 30- to 80-fold less potent than on HIPK4, highlighting the compound’s exceptional target specificity.

Since AZ137 (**28e**) consistently demonstrated the highest selectivity, it was selected for broader profiling against the *scan*MAX™ panel (KINOMEscan®, Eurofins DiscoverX) encompassing 468 kinases. AZ137 (**28e**) displayed an exceptionally clean profile. At a screening concentration of 100 nM, YSK4 (MAP3K19) and HIPK4 were the only substantial hits with PoC values of 2.4% and 7.3%, respectively. However, YSK4 (MAP3K19) was not identified as a potential hit in the NanoBRET K192 panel (**Figure 2B, Table S4**). To validate these findings, YSK4 (MAP3K19) was evaluated in a dose-dependent NanoBRET assay, which revealed no significant cellular target engagement with YSK4 (EC_50_ > 25 µM). This suggests that the scanMAX signal likely represents a false positive result (**Figure 3B/C**). In addition, kinases identified in the initial DSF panel as potential off-targets (Δ*T_m_* > 5 K), such as DYRK2 and BMP2K, were also measured in a dose-dependent NanoBRET assay. AZ137 (**28e**) exhibited weak cellular target engagement on BMP2K (EC_50_ = 11.8 µM), whereas more pronounced engagement was observed for DYRK2 (EC_50_ = 912 nM) (**Figure 3B/C**). Despite this activity, a substantial selectivity window relative to HIPK4 was maintained, further supporting the high target specificity of AZ137 (**28e**).

### 2.5 Crystal structure analysis revealed a preorganized conformation of quinoline-based macrocycles

In order to gain a better understanding of the binding mode and the effects of macrocyclization, we generated a crystal structure with the bosutinib-inspired macrocycles. To date no HIPK4 structure has been published and the lack of efficient expression systems precluded crystallization attempts of this kinase. The weak off-target CLK3 was therefore used as a surrogate model. As shown in **Figure 4A**, the two proteins share high sequence similarity, particularly within the ATP-binding site and major residue variations occur only in the solvent-accessible surface, specifically the exchange of HIPK4 glutamic acid (Glu93) and glutamine (Gln94) for glycine (Gly240) and lysine (Lys241) in CLK3. A minor variation was observed on the β7-strand, where Met143 in HIPK4 is replaced by Leu290 in CLK3. Additionally, Ile73 in HIPK4 corresponds to Val220 in CLK3, which is located between the αC-helix and the gatekeeper residue. All in all, the structural comparison revealed an overall high degree of similarity between these two kinases’ ATP binding sites. While no stabilization of CLK3 by AZ137 (**28e**) was observed in our thermal shift assay, the closely related derivatives **23e** and **30** were used for structure determination with the goal to evaluate how the linker and the pendant methoxy group influenced the overall binding mode of these bosutinib-inspired macrocycles.

**Figure 4.**
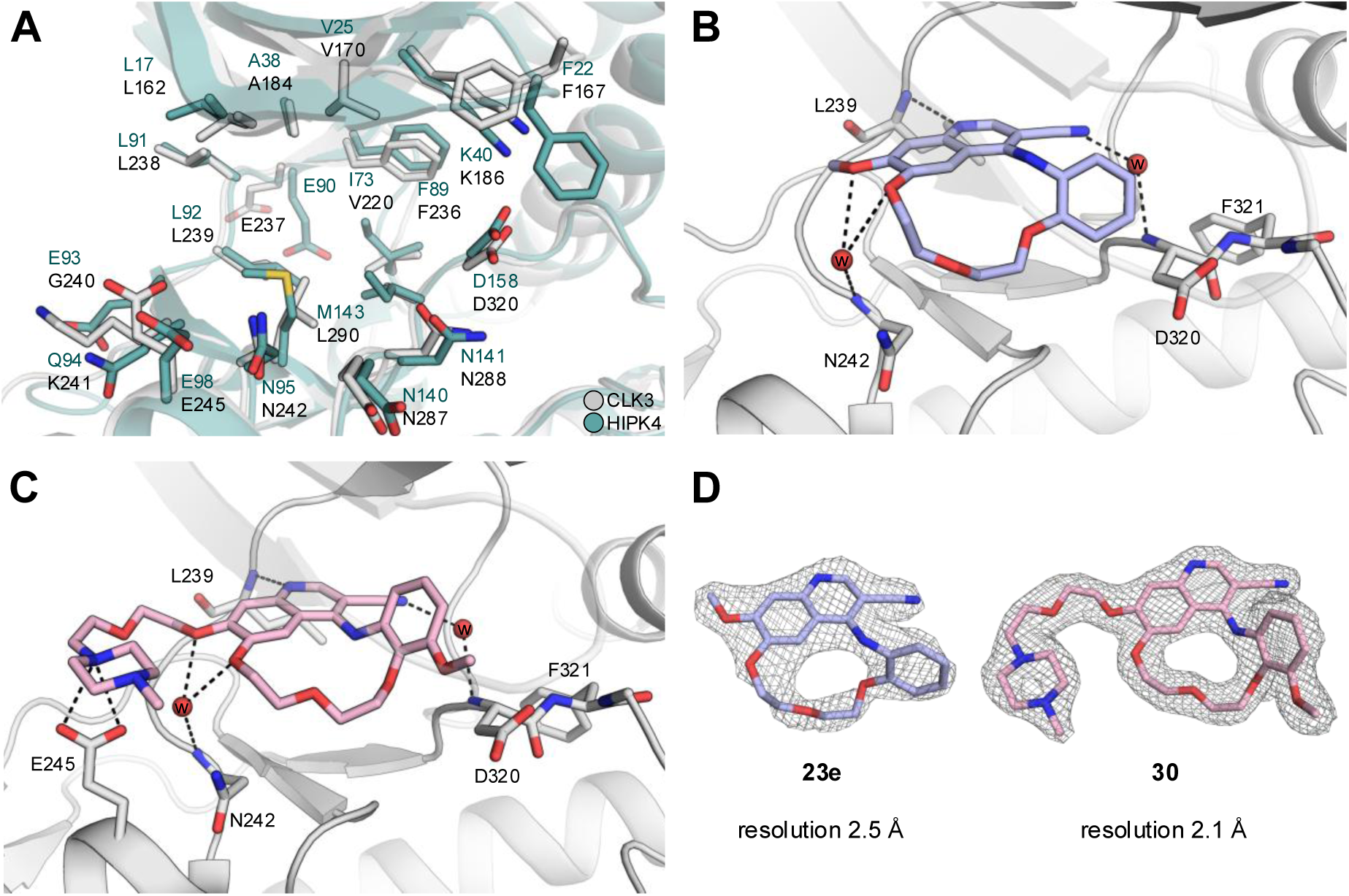
Crystal structures of CLK3. **A** Overlay of the experimental CLK3 structure (gray) with the HIPK4 (green) model predicted by AlphaFold (AF-Q8NE63-F1-v4). **B** Binding mode of **23e** (blue) in CLK3. **C** Binding mode of **30** (pink) in CLK3 (gray) shown in cartoon representation. **D** 2Fo–Fc electron density maps that defined the binding modes of the ligands contoured at 1σ.

As expected, the quinoline ring adopted a highly conserved binding mode within the ATP-binding pocket, with the quinoline nitrogen interacting with Leu239 of the hinge backbone (GK+3). Furthermore, the nitrile group formed a water-mediated interaction with the backbone of Asp320 of the DFG motif. In addition, the two oxygens of the methoxy groups pendant to the quinoline ring established an additional water-mediated hydrogen bond to the Asn242 backbone, confirming a conserved Type I binding mode for the macrocyclic compound **23e** (**Figure 4B**). In the case of the solvent-exposed methyl-piperazine derivative, **30**, a bent conformation was observed that enables an additional interaction between the piperazine moiety and the Glu245 carboxylic acid in CLK3, further validating the conserved binding mode of these series of quinoline-based macrocycles (**Figure 4C**). Notably, even the shorter PEG1 linker in **23e** and **30** retained sufficient flexibility for the pendant aniline ring to adopt a slightly different orientations, which was probably caused by the steric demand of the methoxy group. However, macrocyclization significantly impacted the positioning of the methoxy group, which was presumably one of the key drivers for the exceptional selectivity of AZ137 (**28e**). While the methoxy group on the pendant aniline ring in bosutinib exhibited high conformational variability across different kinases (**Figure 1D**), the macrocyclic constraint bosutinib analogue forces it into a preorganized conformation (**Figure 4C/D**).

### 2.6 Design of a negative control and physicochemical evaluation of Bosutinib-inspired macrocycles

In order to complete the probe set, we aimed to synthesize an inactive control. Its design was based on the insights gained from our prior SAR investigations (**Figure 5A**). Notably, potent HIPK4 inhibition was maintained by introducing a methoxy group in the solvent-exposed area, as well as substitutions at the ortho-position relative to the nitrogen and oxygen of the aniline ring. In contrast, substituents at the para-position were poorly tolerated, likely due to a steric clash within the back-pocket, as suggested by our crystal structures (**Figure 4B/4C**). Regarding the choice of the macrocyclic linker, the SAR revealed a clear trend, showing that the longer aliphatic linkers (C6 and C7) were generally better tolerated by HIPK4 than the C5 linker. Conversely, the PEG1 linker showed superior activity compared to the PEG2 linker (**Table 1 & 2**). Consequently, for the design of our inactive control, we opted to omit the methoxy group in the solvent-exposed area, utilizing a C5 linker and a para-chloro substitution relative to the oxygen on the aniline ring, which ultimately led to compound **31** (NDR44). Compound **31** displayed negligible HIPK4 activity in both the *in vitro* activity assay and binding in the cellular NanoBRET assay. Furthermore, it exhibited excellent kinome-wide selectivity within our internal DSF panel, with no kinase showing temperature shifts larger than 5 K (**Figure 5C** and **Table S3**), thereby proving to be a suitable candidate as an inactive control.

**Figure 5.**
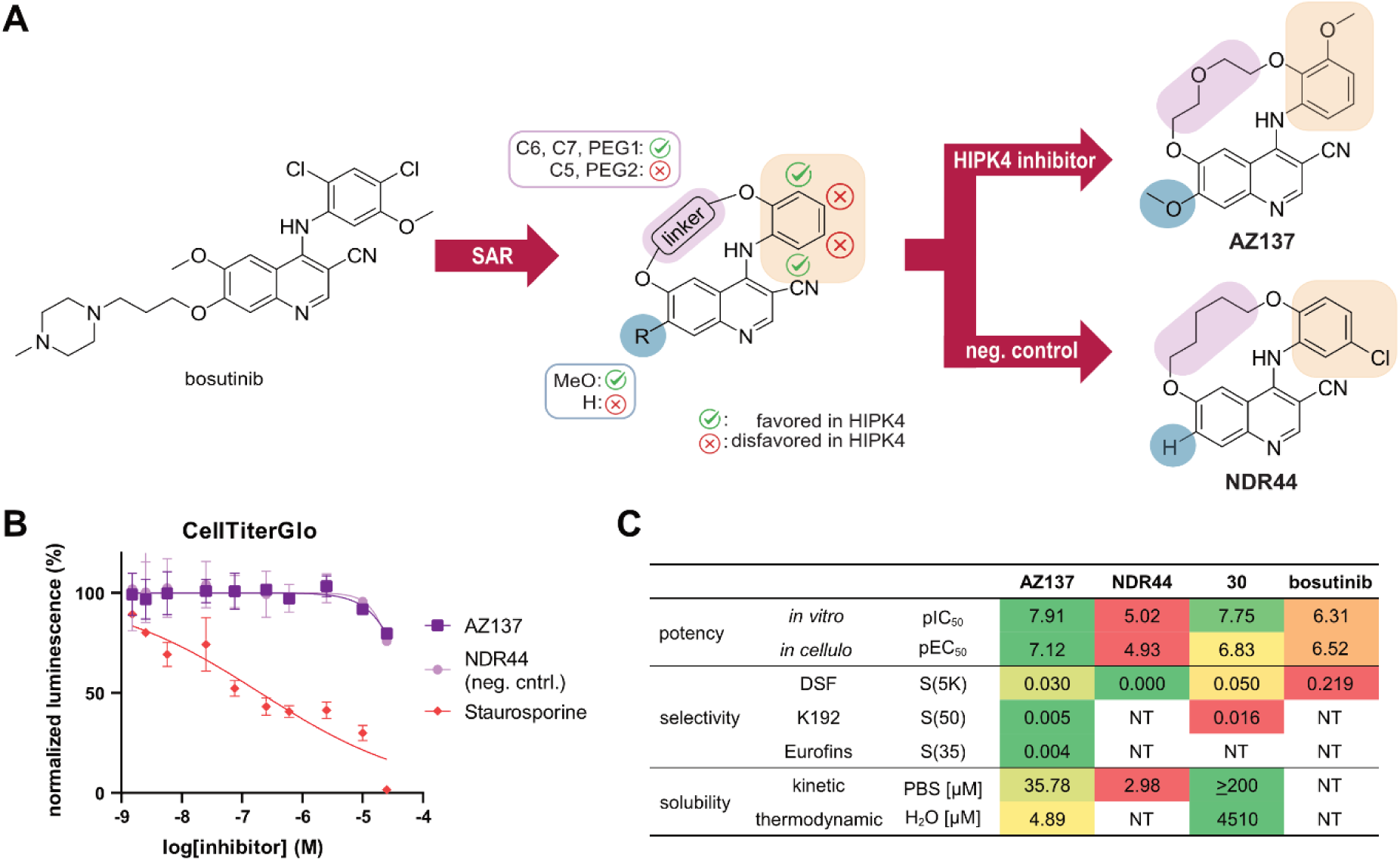
Design of inactive control compound and evaluation of cytotoxicity and physicochemical properties of bosutinib-inspired macrocycles. **A** Summary of the SAR analysis and design for an inactive control compound **31** (NDR44). **B** HEK293T cell viability after 24 h treatment with AZ137 (**28e**) and NDR44 (**31**), determined using a CellTiter-Glo assay (Promega). The curve represents one of two measured biological replicates. Error bars represent the standard deviation (SD) of two technical replicates. **C** Potency, selectivity and solubility summary of AZ137 (**28e**), NDR44 (**31**), **30** and bosutinib.

To further assess the compounds in a biological setting, their impact on cell viability was examined in HEK293T cells utilizing the CellTiter-Glo® Luminescent Cell Viability Assay (Promega). For this purpose, we employed AZ137 (**28e**) as the potential probe candidate and NDR44 (**31**) as the inactive control, while staurosporine served as a positive control for the assay. The compounds were tested in a dose-dependent manner. Notably, neither AZ137 (**28e**) nor NDR44 (**31**) exhibited significant cytotoxicity at concentrations up to a compound concentration of 25 µM (**Figure 5B**). To ensure the reliability of the biochemical data obtained, we assessed the kinetic solubility of the key compounds in PBS buffer. This allowed us to exclude solubility-related artifacts from our analysis. Consequently, AZ137 (**28e**), **30**, and NDR44 (**31**) were chosen for further solubility testing. Compound **30**, containing a piperazine moiety, demonstrated excellent solubility (> 200 µM, the assay’s upper limit). AZ137 (**28e**) also showed good solubility (35.78 µM), while the negative control NDR44 (**31**) was less soluble (2.98 µM) but given the cellular activity of the developed chemical probe this value should still be acceptable.

Our macrocycles displayed high kinetic solubility, mirroring the observations made by Walz *et al.* that macrocyclization serves as an effective strategy to improve the physicochemical properties such as kinetic solubility. ^37^ Nevertheless, as kinetic solubility measurements tend to overestimate solubility compared to thermodynamic values, we additionally determined the thermodynamic solubility of AZ137 (**28e**), and **30** in water. Our results further demonstrated favorable solubility profiles for both macrocycles. Specifically, AZ137 (**28e**), showed a solubility of 4.89 µM, whereas compound **30** displayed an exceptional solubility of 4510 µM.

### 2.7 AZ137-mediated inhibition of HIPK4-dependent F-actin remodeling in fibroblasts

Although HIPK4 has not yet been fully characterized, studies by Crapster *et al.* and Liu *et al.* showed that it actively regulated actin dynamics.^14,15^ HIPK4 ensures the proper organization and stabilization of the actin cytoskeleton network presumably by phosphorylating key target proteins. F-actin dynamics appear to play a fundamental role, particularly in the formation of the acroplaxome during spermiogenesis. The acroplaxome itself serves as a cytoskeletal anchor plate composed of keratin 5 and F-actin, linking the acrosome to the nucleus. Consequently, the acroplaxome is critical for sperm head shaping, and defects in this structure or its associated proteins frequently lead to globozoospermia.

The pivotal study by Crapster *et al.* was the first to identify the association between HIPK4 and F-actin dynamics in the acroplaxome.^14^ Specifically, they demonstrated that loss of HIPK4 in male mice results in impaired spermatogenesis due to structural dysregulation of the acrosome-acroplaxome complex. Furthermore, they showed that overexpression of HIPK4 induces significant remodeling of the F-actin cytoskeleton, although the precise downstream phosphorylation targets mediating this effect remain to be fully characterized. Consequently, we were interested to determine whether the pharmacological inhibition of HIPK4 with our potent macrocyclic inhibitors could directly modulate these F-actin dynamics.

To this end, immunofluorescence studies were conducted in NIH-3T3 cells using AZ137 (**28e**), the compound with the most favorable selectivity profile, and NDR44 (**31**), which served as a negative control. In agreement with previous observations by Crapster *et al*., our data demonstrated that DMSO-treated wild-type (WT) reference cells exhibited a spindle-like morphology with long F-actin filaments, whereas HIPK4 overexpression induced a transition toward spherical or polygonal cells with branched F-actin structures(**Figure 6A/B**). This indicates a significant depletion of F-actin stress fibers resulting from HIPK4 overexpression. Treatment with the inactive control NDR44 (**31**) (up to 5 µM) failed to alter cellular morphology, with the cells retaining their polygonal appearance, indicating that the inactive control is incapable of reversing the F-actin phenotypes induced by HIPK4 (**Figure 6C**). In contrast, treatment with varying doses of AZ137 (**28e**) revealed that the lowest concentration (0.2 µM) had only a minimal effect on F-actin dynamics (**Figure 6D**). However, at higher concentrations (1 and 5 µM), AZ137 (**28e**) successfully restored the disrupted actin architecture, returning it to a normal fibrous structure (**Figure 6E/F**). Previous studies have linked HIPK4 to the stabilization of the actin cytoskeleton, though the requirement of its catalytic activity for these functions has not been definitively established. Our findings with **AZ137** demonstrate that the pharmacological inhibition of HIPK4 blocks F-actin remodeling in NIH-3T3 cells, providing strong evidence that its kinase activity is indeed essential for these cytoskeletal rearrangements. Whether this catalytic role directly translates to the actin-rich acroplaxome during spermiogenesis, or if these processes are mechanistically divergent, remains an intriguing question for further study.

**Figure 6.**
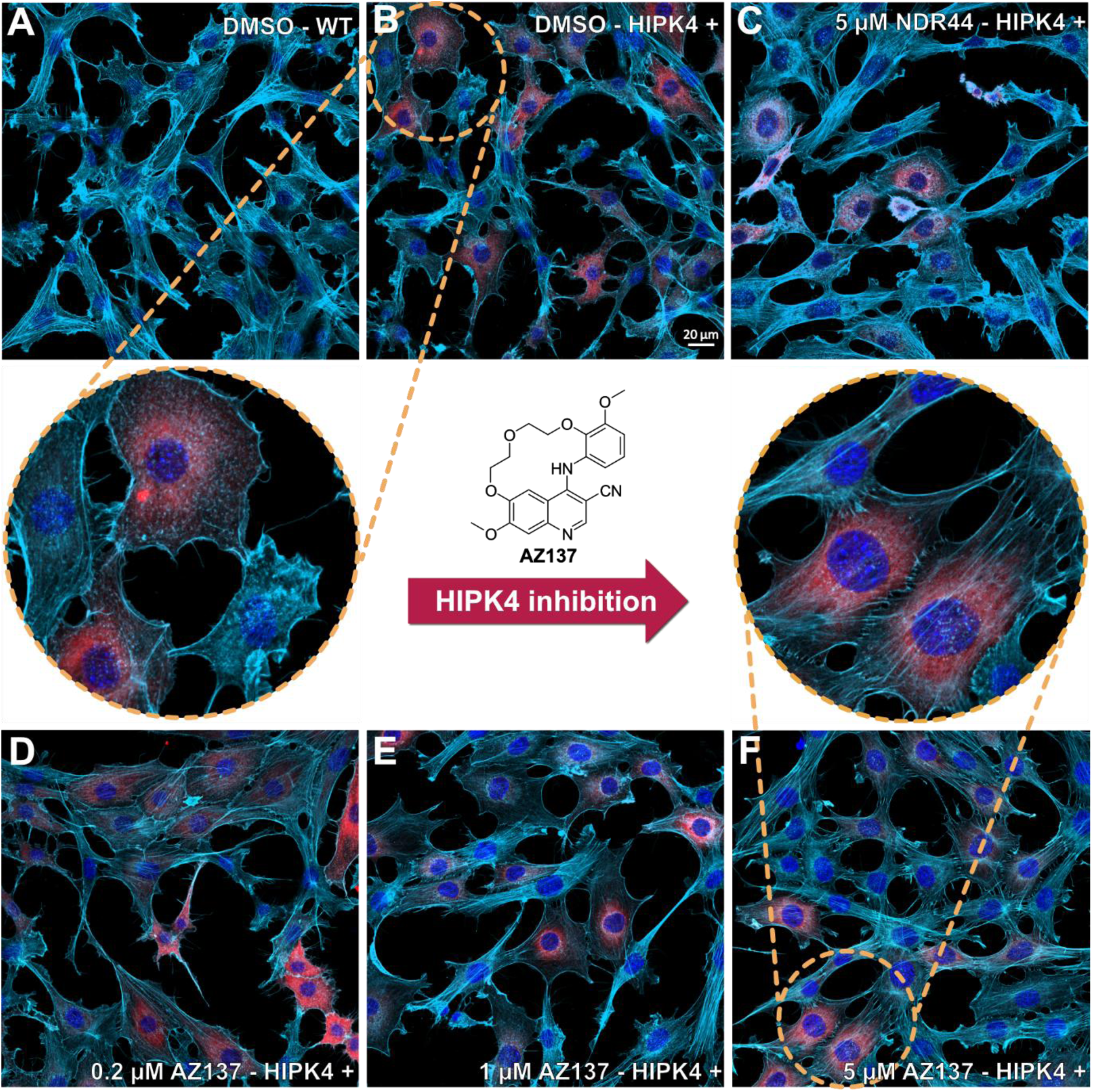
Impact of AZ137 on HIPK4-mediated actin cytoskeleton remodeling in NIH-3T3 cells. **A** Wild-type NIH-3T3 cells and **B** NIH-3T3 cells retrovirally transduced with FLAG-tagged HIPK4, stained with an anti-FLAG primary antibody followed by an Alexa Fluor 555-conjugated secondary antibody (FLAG-HIPK4, red) and Alexa Fluor 647-conjugated phalloidin (F-actin, far-red). Nuclei were counterstained with DAPI (blue). **C** Effect of the inactive control compound NDR44 (5 µM) on HIPK4-overexpressing cells. **D**–**F** Dose-dependent inhibition of HIPK4-driven F-actin remodeling by AZ137 (0.2–5 µM). Circled areas in panel B and F provide a magnified view of HIPK4-overexpressing cells.

## CONCLUSION

In this study, we utilized the non-selective acyclic compound bosutinib as a starting point to develop conformationally constrained macrocyclic inhibitors. By systematically modifying the solvent-exposed region and the back-pocket, we identified AZ137 (**28e**), which fulfills the criteria for a chemical probe for HIPK4.^38,39^ This compound exhibits a well-balanced profile of potent activity in both biochemical and cellular contexts, coupled with excellent kinome-wide selectivity and favorable physicochemical properties.

Notably, our discovery of the quinoline scaffold as a potent HIPK4 binder aligns with independent findings from the Chen Lab, which has also identified quinoline-based derivatives as effective HIPK4 inhibitors by a different approach.^53^ By evaluating our probe package in the context of HIPK4-induced F-actin remodeling, we showed that pharmacological intervention with AZ137 (**28e**), but not with the inactive analog NDR44 (**31**), effectively reverses the actin phenotype. While previous insights into HIPK4’s role in cytoskeletal dynamics were derived primarily from genetic knockout models, we here demonstrate that these effects can be effectively controlled through targeted pharmacological inhibition. This confirms that HIPK4’s catalytic activity is essential for these processes and validates the functional utility of our tools. In conclusion, the extensive characterization of AZ137 (**28e**) and its negative control provides the scientific community with a robust pharmacological toolset. While chemical matter for HIPK4 has recently begun to emerge in the patent literature, this well-characterized and accessible probe package offers a unique opportunity to further elucidate the biological roles of this dark kinase. These tools may offer additional opportunities to study HIPK4 in contexts like spermatogenesis or epithelial differentiation and could potentially help reveal new aspects of its biological signaling.

## EXPERIMENTAL SECTION

### ADP-Glo Assay

Inhibitory activity against recombinant human HIPK4 (GST-tagged) was measured using the ADP-Glo Kinase Assay kit (Promega) according to the manufacturer’s instructions with minor modifications. Reactions were performed in 96-well plates in kinase buffer containing 40 mM Tris (pH 7.5), 20 mM MgCl_2_, and 0.1 mg/mL BSA. Each 25 µL reaction mixture contained HIPK4 (3 nM), myelin basic protein (MBP, 2 µM) as substrate, and ATP (25 µM). Test compounds were prepared as serial dilutions in DMSO and added to give a final DMSO concentration of 1% (v/v). After preincubation of enzyme with compounds, reactions were initiated by addition of substrate/ATP mixture and incubated for 30 min at room temperature. Kinase reactions were terminated by addition of 25 µL ADP-Glo Reagent followed by a 40 min incubation at room temperature to deplete remaining ATP. Subsequently, 50 µL of Kinase Detection Reagent was added, and plates were incubated for an additional 30 min before luminescence measurement using a microplate reader. Percent inhibition was calculated relative to DMSO-treated controls, and IC_50_ values were determined by nonlinear regression analysis using a four-parameter logistic equation in GraphPad Prism. Reported values represent the mean of at least two independent experiments performed in duplicate.

### NanoBRET Assay

For cellular target engagement measurements, HEK293T cells were seeded at a density of 2,000 cells per well by dispensing 10 µL of cell suspension in Opti-MEM without phenol red (Thermo Fisher, 11058021) into white 384-well LDV plates (Greiner, 784075). Transfections were performed by adding 1 µL of transfection mix per well. The transfection mix was prepared by combining 0.5 µL of NLuc fusion plasmid DNA (200 ng/µL), 0.9 µL of transfection carrier DNA (Promega, E488A), and 3 µL of FuGENE® 4K Transfection Reagent (Promega, E591A) with 95 µL of Opti-MEM, yielding a total volume of 100 µL. Following transfection, cells were incubated for 16–24 h at 37 °C in a humidified atmosphere containing 5 % CO₂. Compounds were serially diluted, and NanoBRET K10 Tracer (Promega, TracerDB ID: T000008) at the Tracer K_d_ concentration taken from TracerDB (tracerdb.org)^40^ were pipetted into the 384-well plates, and incubated for 2.5 hours in the cell incubator. Before the readout, 5 µl of 1:100 Nano-Glo® Substrate (Promega, N157A) in Opti-MEM was added and incubated for 5 minutes at room temperature. BRET measurements were acquired in dual-emission mode using a PHERAstar FSX plate reader (BMG Labtech) equipped with luminescence filters centred at 450 nm (donor) and 610 nm (acceptor). For analysis, the acceptor-to-donor signal ratio was multiplied by 1000 to obtain signals in milliBRET units. Data were fitted using a normalized 3-parameter curve fit using GraphPad Prism software and following equation: Y = Bottom + (Top − Bottom)/(1 + 10^(X − LogIC50))**DSF Assay.** Recombinant protein kinase domains (2 µM) were incubated with a 10 μM compound solution in DMSO, 20 mM N-(2-hydroxyethyl)piperazine-N′-ethanesulfonicacid (HEPES), pH 7.5, and 500 mM NaCl. SYPRO Orange (5000×, Invitrogen) was added as a fluorescence probe (1 μL per mL). Thermal shift experiments were performed on a QuantStudio 5 real-time PCR system (ThermoFisher Scientific) using excitation and emission wavelengths of 465 and 590 nm, respectively. The temperature was increased at a heating rate of 3 °C per minute. Melting temperatures were determined by fitting the unfolding curves with a Boltzmann model using Thermal Shift Software v1.4 (Thermo Fisher Scientific). Thermal shifts are reported as ΔTm values in Kelvin. Measurements were carried out in technical duplicates.

### NanoBRET K192 Panel

To assess kinase inhibitor selectivity, the K192 Kinase Selectivity System (Promega, cat. no. NP4050) was performed in a miniaturized 384-well format as described in detail by Schwalm *et al*.^34^. In brief, for plate preparation, transfection mix was prepared in white 384-well small-volume plates (Greiner, cat. no. 784075) by pre plating 3 µL of 20 µL/mL FuGENE HD (Promega, cat. no. E2311), diluted in Opti-MEM medium (Gibco, cat. no. 11058-021). 1 µL DNA from both DNA vector source plates of the K192 kit was added using an Echo acoustic dispenser (Beckman Coulter). The mix was incubated for 30 min and 6 µL HEK293T cells in Opti-MEM medium were added. The proteins were allowed to express for 20 h. After expression, Tracer K-10 was added using the concentrations recommended in the K192 technical manual and 1 µM inhibitor was added to every second well. After 2 h of equilibration, detection was carried out using substrate solution comprising OptiMEM with a 1:166 dilution of NanoBRET™ Nano-Glo® Substrate and a 1:500 dilution of the Extracellular NanoLuc® Inhibitor. 5 µL of substrate solution was added to every well and filtered luminescence was measured on a PHERAstar plate reader (BMG Labtech) equipped with a luminescence filter pair (450 nm BP filter (donor) and 610 nm LP filter (acceptor)). For every kinase, occupancy was calculated and plotted using GraphPad Prism 10.

### Kinome-Wide Selectivity Profile

Kinase selectivity of the AZ137 was assessed using the scanMAX™ platform (KINOMEscan®, Eurofins DiscoverX). The compound was profiled against a panel of 468 kinases in an ATP-independent competition binding assay at concentrations of 1 µM and 100 nM. Data analysis was performed by the service provider using standard procedures.

### Protein Expression and Purification

The CLK3 kinase domain (residues 127–484), containing an N-terminal His6 tag with a TEV cleavage site, was transformed into BL21(DE3) cells. Bacteria were grown in Terrific Broth medium supplemented with 50 µg/mL kanamycin. Protein expression was induced at an OD600 of 2 with 0.5 mM IPTG at 18 °C for 12 hours. Cells were lysed by sonication in lysis buffer containing 50 mM HEPES pH 7.5, 500 mM NaCl, 25 mM imidazole, 5% glycerol, 50 mM L-glutamic acid, 50 mM L-arginine, and 0.5 mM TCEP. After centrifugation, the supernatant was loaded onto a Nickel-Sepharose column equilibrated with 30 mL of lysis buffer. The column was washed with 60 mL of lysis buffer and protein was eluted using an imidazole step gradient (50, 100, 200, and 300 mM). CLK3 was dialyzed overnight against SEC buffer (50 mM HEPES pH 7.5, 500 mM NaCl, 5% glycerol, 50 mM L-glutamic acid, 50 mM L-arginine, and 0.5 mM TCEP), and TEV protease was added to remove the tag. After reverse Ni-affinity chromatography to remove TEV protease and uncleaved protein, the sample was further purified by size-exclusion chromatography using a Superdex 75 16/60 HiLoad column. Pure protein fractions were pooled and concentrated to ∼11 mg/mL.

### Crystallization

CLK3 (∼11 mg/mL) was co-crystallized at 4 °C using the sitting-drop vapor diffusion method by mixing protein and well solutions in 2:1, 1:1, and 1:2 ratios (final drop volume 200 nL). Compounds were pre-incubated with the protein at a final concentration of ∼500 µM for approximately 30 min on ice prior to crystallization. Reservoir solutions included either 25% PEG 3350, 0.2 M NaI, and 10% ethylene glycol, or 20% PEG 3350, 0.1 M bis-tris propane, and 0.2 M ammonium sulfate. Crystals were cryoprotected with 25% ethylene glycol before flash-freezing in liquid nitrogen. Diffraction data were collected at beamline I03 (Diamond Light Source, UK) at a wavelength of 0.976 Å and a temperature of 100 K. Data were automatically integrated using XDS^41^, DIALS^42^, xia2^43^, or autoPROC^44^, and scaled with AIMLESS^45^. The PDB structure with accession code 9EZ3^46^ was used as the initial search model in MOLREP^47^. The final model was built manually in Coot^48^ and iterative refinement cycles using REFMAC5^49^. Data collection and refinement statistics are summarized in **Table S6**. Ligand dictionary files were generated using the Grade Web Server ().

### Kinetic solubility

Stock solutions of each compound were prepared in a DMSO/ACN (1:1, v/v) mixture at concentrations of 10, 25, 50, 100, and 200 µM. These standards were analysed via HPLC with UV detection at λ = 254 nm. The resulting peak areas were integrated, and a calibration curve was generated using GraphPad Prism 8. Linearity was determined via linear regression to provide a basis for quantification. To determine aqueous solubility, compounds were incubated in duplicates at a nominal concentration of 200 µM in phosphate-buffered saline (PBS, pH 7.4) containing 2% DMSO. The samples were placed in a water bath at 25°C and agitated on an orbital shaker at 100 rpm for 2 hours. Following incubation, the samples were centrifuged at 14 000 rpm for 5 min. 100 µL of dissolved compound were pipetted into 100 µL of DMSO/ACN (1:1, v/v) and quantified using the previously established HPLC-UV method. The solubility of compounds was calculated with the calibration formula determined before.

### Thermodynamic solubility

Aqueous solubility of AZ137 was determined using Whatman Uniprep filters (Whatman plc, Maidstone, UK). A precisely weighed amount of each compound and 2 mL of water were inserted into the Uniprep vessel, and the mixture was shaken at 37 °C for 24 h. The mixture was then pressed through the Uniprep filter and the concentration of dissolved compound in filtrate was quantified by HPLC (Hitachi Chromaster and Hitachi DAD 5430 Detector equipped with an ACE 5 C18, 250+4 mm column) using external calibration.

### Cell Viability Assay

Compound effects on cell viability were assessed using the CellTiter-Glo® 2.0 Cell Viability Assay (Promega), following the manufacturer’s instructions. HEK293T cells were seeded at 2,000 cells per well in 384-well plates (Greiner: 781207) and incubated overnight at 37 °C with 5% CO₂. After incubation, cells were treated for 24 hours with compounds titrated across a concentration range of 25 µM to 1.25 nM using the Echo 550 Liquid Handler (Labcyte). Following treatment, an equal volume of CellTiter-Glo® 2.0 reagent was added to each well, and plates were incubated for 10 minutes at room temperature. Luminescence was recorded using a PHERAstar plate reader (BMG Labtech). Each compound was tested as two technical replicates per biological replicate (n = 2). Technical replicates were averaged, baseline-corrected, and normalized to the negative control (DMSO). Dose–response curves were analyzed using GraphPad Prism 9 with a normalized response vs. log(inhibitor) model, applying the following equation: Y = 100 / (1 + 10^((LogIC₅₀ – X) × HillSlope)).

### Immunofluorescence studies

NIH-3T3 cells were seeded at a density of 5,000 cells per well in 24-well plates containing poly-L-lysine–coated glass coverslips and cultured in antibiotic-free Dulbecco’s modified Eagle medium (DMEM) supplemented with 10% fetal bovine serum and 1 mM sodium pyruvate. After 24 h, the culture medium was removed and cells were transduced with human HIPK4 retrovirus (multiplicity of infection = 1.5, produced as previously described [Crapster *et al*. eLife]) by addition of 10 μL of viral supernatant per well, polybrene (final concentration 4 μg/μL), and 500 μL of antibiotic-free medium. Twenty-four hours post-transduction, cells were rinsed with phosphate-buffered saline (PBS) and fixed with 4% paraformaldehyde in PBS for 15 min at room temperature. Following fixation, cells were washed with PBS, permeabilized with 0.3% Triton X-100 in PBS for 10 min, washed again with PBS, and blocked with 2% bovine serum albumin in PBS containing 0.1% Triton X-100 for 1 h at room temperature. Cells were incubated overnight at 4 °C with anti-FLAG primary antibody (1:100 dilution), washed with PBS, and subsequently incubated with Alexa Fluor–conjugated secondary antibody (AlexaFluor 555, 1:400 dilution) and phalloidin Alexa Fluor–conjugated primary antibody (AlexaFluor 647, 1:1000 dilution) for 1 h at room temperature. After final PBS washes, coverslips were mounted onto glass slides using VECTASHIELD Vibrance Antifade Mounting Medium containing DAPI. Fluorescence images were acquired using a Zeiss LSM 700 or 800 confocal microscope equipped with a 63x oil-immersion objective.

### Chemistry

The synthesis of all compounds is detailed below, and their analytical data is found in the Supporting Information (**Figure S1–49**). All commercial chemicals were purchased from common suppliers with a purity of ≥ 95% and did not undergo further purification. The solvents with an analytical grade were obtained from VWR Chemicals and the dry solvents were purchased from Acros Organics and stored under argon gas and over molecular sieves. Unless stated otherwise, all reaction flasks were purged with argon after the solid reagents had been added and without being priorly baked out. The flasks were sealed with a septum and a balloon, filled with argon and attached to a cannula, was pierced through the septum for the duration of the reaction. All microwave-assisted reactions were carried out in sealed reaction vials (2–20 mL) with a Biotage Initator+ Microwave System with Robot Eight by Biotage. To monitor the progression of the reactions and determine the purity of compounds high-performance liquid chromatography (HPLC) was performed. Analytical HPLC was performed via a 1260 Infinity II LC System consisting of the multisampler G7167A, the column compartment G7116A, the multicolumn thermostat G7116A, the flexible pump G7104C, the diode array detector HS G7117C (254, 280, 310 nm) and a single quadrupole LC/MSD system InfinityLab G6125B (ESI pos. 100−1000) by the company Agilent Technologies was used for this purpose. A Porochell 120 EC-C18 column (3.0 x 150 mm, 2.7 μm) from Agilent was used as the stationary phase and a gradient of H2O and ACN with 0.1% formic acid and with a flow rate of 0.6 mL/min served as the mobile phase. All compounds used for further biological characterization attained a purity of ≥ 95% by HPLC. The purification of the compounds was done via flash chromatography and preparative HPLC. For the purification via flash chromatography a puriFlash XS 420Plus device with a UV−vis multiwave detector (200−400nm) from Interchim was utilized with prepacked normal-phase PF-30SIHP and reversed-phase PF-30C18HP silica columns with a particle size of 30 μm (Interchim). The purification via preparative HPLC was performed using a 1260 Infinity II Preparative LC system consisting of the preparative autosampler G7157A, the preparative column organizer G9328A, the preparative binary pump G7161A, the multiple wavelength detector G7165A and the preparative-scale fraction collector G1364E. An Eclipse XDB-C18 column (21.2 x 250 mm, 7 μm) from Agilent was used as the stationary phase and a gradient of H2O and ACN with 0.1% TFA and with a flow rate of 21 mL/min served as the mobile phase. Nuclear magnetic resonance spectroscopy (NMR) was performed with DPX250 (250 MHz ^1^H), AV400HD (400 MHz ^1^H, 101 MHz ^13^C) and AV500HD (500 MHz ^1^H, 126 MHz ^13^C) spectrometers from Bruker. The measurements were performed at room temperature and in deuterated solvent (DMSO-*d*6 or CDCl3 by Eurisotop). The NMR spectra were interpreted using MestreNova. The chemical shift *δ* was stated in ppm and referenced through solvent resonance (DMSO-*d*6 ^1^H NMR: 2.50 ppm, ^13^C NMR: 39.52 ppm; CDCl3 ^1^H NMR: 7.26 ppm, ^13^C NMR: 77.16 ppm).^50^ The coupling constant *J* was stated in Hz. The multiplicity of the signals in the spectra was characterized by following abbreviations: s (singlet), br (broad singlet), d (doublet), dd (doublet of doublets), t (triplet), td (triplet of doublets), m (multiplet).

### Synthesis of Methyl (E)-2-(((dimethylamino)methylene)amino)-5-methoxybenzoate (2)

Compound **2** was synthesized following the patented conditions of Miller *et al*.^51^. 2-Amino-5-methoxy benzoic acid (**1**, 5.00 g, 29.9 mmol, 1.0 equiv) and DMF dimethyl acetal (11.5 mL, 86.2 mmol, 2.9 equiv) were solved in 22 mL dry DMF. The reaction mixture was stirred at 150 °C for 3 h. The solvent was removed in *vacuo*, the residue taken up in a saturated NaHCO3 solution and the product extracted with DCM. The combined organic phases were washed with brine and dried over MgSO4. The solvent was removed *in vacuo* to obtain the crude as a dark oil. The crude product was further converted without additional purification. ^1^H NMR (400 MHz, DMSO): *δ* 7.51 (s, 1H), 7.02 (d, *J* = 3.0 Hz, 1H), 6.96 (dd, *J* = 8.7, 3.1 Hz, 1H), 6.85 (d, *J* = 8.8 Hz, 1H), 3.73–3.72 (m, 6H), 2.93 (s, 6H) ppm. MS (ESI^+^): *m*/*z* 237.1 [M + H]^+^.

### Synthesis of 4-Hydroxy-6-methoxyquinoline-3-carbonitrile (3)

Compound **3** was synthesized following the patented conditions of Miller *et al*.^51^. To a solution of dry ACN (4.7 mL, 89.7 mmol, 3.0 equiv) diluted with 40 mL dry THF and cooled to –78 °C. *n-*Butyllithium (28.7 mL, 2.5 M in hexane, 71.8 mmol, 2.4 equiv) was added dropwise over an hour. The suspension was stirred at –78 °C for 4.5 h. A solution of the crude compound **2** solved in 40 mL dry THF was added dropwise to the reaction mixture. The reaction mixture was stirred at room temperature overnight. The reaction was quenched by adding concentrated acetic acid (13.7 mL, 239 mmol, 8.0 equiv) and stirred at room temperature for 2 h. 60 mL water were added, and the suspension was filtered. The precipitate was washed with water. The residue was dried *in vacuo* to give **3** a brown solid with a yield of 86% over two steps (5.13 g). ^1^H NMR (500 MHz, DMSO): *δ* 12.78 (s, 1H), 8.65 (s, 1H), 7.60 (d, *J* = 9.0 Hz, 1H), 7.51 (d, 4J = 2.9 Hz, 1H), 7.41 (dd, *J* = 9.0, 3.0 Hz, 1H), 3.86 (s, 3H) ppm. ^13^C NMR (126 MHz, DMSO): *δ* 174.0, 157.0, 145.1, 133.6, 126.4, 123.2, 121.1, 117.1, 104.6, 92.4, 55.6 ppm. MS (ESI^+^): *m*/*z* 223.1 [M + Na]^+^.

### Synthesis of 4-Chloro-6-methoxyquinoline-3-carbonitrile (4)

Compound **3** (9.93 g, 49.6 mmol, 1.0 equiv) was solved in 100 mL dry toluene. The solution was cooled to 0 °C and phosphoryl chloride (30 mL, 321 mmol, 6.5 equiv) was added slowly. The mixture was stirred at 120 °C for 3.5 h. The solvent was removed *in vacuo*, the residue taken up in a 5% aqueous solution of NaOH and the product extracted with DCM. The combined organic phases were dried over MgSO4, filtered and the solvent removed *in vacuo* to obtain compound **4** as a beige solid (10.67 g, 98%). ^1^H NMR (500 MHz, DMSO): *δ* 9.04 (s, 1H), 8.10 (d, *J* = 9.2 Hz, 1H), 7.70 (dd, *J* = 9.2, 2.8 Hz, 1H), 7.50 (d, *J* = 2.8 Hz, 1H), 4.00 (s, 3H) ppm. ^13^C NMR (126 MHz, DMSO): *δ* 159.6, 147.8, 144.9, 144.2, 131.6, 126.3, 125.6, 115.3, 107.1, 102.2, 56.1 ppm. MS (ESI^+^): *m*/*z* 218.9 [M + H]^+^.

### Synthesis of 4-Chloro-6-hydroxyquinoline-3-carbonitrile (5)

Compound **4** (5.0 g, 22.9 mmol, 1.0 equiv) and tertbutyl ammonium iodide (16.9 g, 45.7 mmol, 2.0 equiv) were solved in 40 mL dry toluene. The solution was cooled to 0 °C. Boron trichloride (45.7 mL, 1 M in toluene, 45.7 mmol, 2.0 equiv) was added slowly. The reaction mixture was stirred at 110 °C overnight. The reaction was quenched by adding 100 mL water and stirring for 30 min. The precipitate was filtered and washed with EtOAc. The residue was dried *in vacuo* to obtain compound **5** as a brown solid (3.83 g, 82%). ^1^H NMR (400 MHz, DMSO): *δ* 10.86 (s, 1H), 8.93 (s, 1H), 8.04 (d, *J* = 9.1 Hz, 1H), 7.57 (dd, *J* = 9.1, 2.7 Hz, 1H), 7.44 (d, *J* = 2.7 Hz, 1H) ppm. MS (ESI^+^): *m*/*z* 205.1 [M + H]^+^.

### Synthesis of 6-((5-((tert-Butyldimethylsilyl)oxy)pentyl)oxy)-4-chloroquinoline-3-carbonitrile (6a)

Triphenyl phosphine (2.56 g, 9.75 mmol, 5 equiv) was suspended in 100 mL dry toluene. Diisopropyl azodicarboxylate (1.97 g, 9.75 mmol, 5.0 equiv) solved in 10 mL dry toluene was added and the suspension was stirred for 20 min. 5-((Tert-butyldimethylsilyl)oxy)pentan-1-ol (852 mg, 3.9 mmol, 2 equiv) solved in 10 mL dry toluene and compound **5** (400 mg, 1.95 mmol, 1 equiv) solved in 20 mL dry DMF were added dropwise. The reaction mixture was stirred at 90 °C overnight. The solvent was removed *in vacuo* and the residue purified via normal-phase flash chromatography to obtain compound **6a** as a colourless oil (710 mg, 90%). ^1^H NMR (500 MHz, DMSO): *δ* 9.04 (s, 1H), 8.10 (d, *J* = 9.2 Hz, 1H), 7.68 (dd, *J* = 9.2, 2.8 Hz, 1H), 7.50 (d, *J* = 2.8 Hz, 1H), 4.22 (t, *J* = 6.4 Hz, 2H), 3.61 (t, *J* = 5.9 Hz, 2H), 1.83 (quint, *J* = 6.7 Hz, 2H), 1.59–1.46 (m, 4H), 0.85 (s, 9H), 0.02 (s, 6H) ppm. ^13^C NMR (126 MHz, DMSO): *δ* 159.0, 147.7, 144.9, 144.1, 131.6, 126.5, 125.7, 115.3, 107.1, 102.8, 68.5, 62.3, 31.9, 28.2, 25.8, 21.9, 18.0, -5.3 ppm. MS (ESI^+^): *m*/z 405.2 [M + H]^+^.

### Synthesis of 6-((5-Hydroxypentyl)oxy)-4-((2-hydroxyphenyl)amino)quinoline-3-carbonitrile (7a)

Compound **6a** (200 mg, 494 µmol, 1.0 equiv) and 2-amino phenol (108 mg, 988 µmol, 2.0 equiv) were solved in 10 mL 2-ethoxyethanol. The reaction mixture was stirred at 140 °C for 2 h and then at room temperature overnight. The reaction mixture was poured into water and the pH was adjusted to 9. The mixture was extracted with EtOAc. The combined organic phases were dried over MgSO4, filtered and the solvent was removed in vacuo. The residue was purified via normal-phase flash chromatography to obtain compound **7a** as a yellow solid (139 mg, 77%). ^1^H NMR (500 MHz, DMSO): *δ* 9.76 (s, 1H), 9.34 (s, 1H), 8.32 (s, 1H), 7.88 (d, *J* = 2.6 Hz, 1H), 7.80 (d, *J* = 9.1 Hz, 1H), 7.42 (dd, *J* = 9.1, 2.6 Hz, 1H), 7.22 (dd, *J* = 7.8, 1.7 Hz, 1H), 7.19 (td, *J* = 7.7, 1.7 Hz, 1H), 6.93 (dd, *J* = 8.1, 1.4 Hz, 1H), 6.86 (td, *J* = 7.6, 1.4 Hz, 1H), 4.38 (s, 1H), 4.12 (t, *J* = 6.6 Hz, 2H), 3.43 (t, *J* = 5.8 Hz, 2H), 1.80 (quint, *J* = 6.6 Hz, 2H), 1.55–1.44 (m, 4H) ppm. ^13^C NMR (126 MHz, DMSO): *δ* 156.8, 153.8, 150.6, 150.6, 143.7, 130.8, 128.8, 128.5, 125.2, 123.1, 119.1, 119.0, 117.2, 115.9, 102.7, 86.0, 68.3, 60.6, 32.2, 28.5, 22.2 ppm. MS (ESI^+^): *m*/*z* 364.4 [M + H]^+^.

### Synthesis of 4,10-dioxa-2-aza-1(4,6)-quinolina-3(1,2)-benzenacyclodecaphane-1^3^-carbonitrile (8a)

Triphenyl phosphine (202 mg, 770 µmol, 2.0 equiv) was solved in 50 mL dry toluene. Diisopropyl azodicarboxylate (156 mg, 770 µmol, 2.0 equiv) solved in 10 mL dry toluene was added dropwise. Compound **7a** (140 mg, 385 µmol, 1.0 equiv) solved in 10 mL dry DMF was added dropwise. The mixture was stirred at 130 °C overnight. The solvent was removed *in vacuo* and the residue was purified via normal-phase flash chromatography and subsequent reversed-phase flash chromatography. Compound **8a** was obtained as a bright yellow solid (40 mg, 30%). ^1^H NMR (500 MHz, DMSO): *δ* 8.70 (s, 1H), 8.25 (s, 1H), 7.89 (d, *J* = 9.1 Hz, 1H), 7.84 (d, *J* = 2.8 Hz, 1H), 7.43 (dd, *J* = 9.1, 2.7 Hz, 1H), 7.27 (dd, *J* = 7.8, 1.6 Hz, 1H), 7.21–7.12 (m, 1H), 7.10 (dd, *J* = 8.3, 1.5 Hz, 1H), 6.91 (td, *J* = 7.6, 1.4 Hz, 1H), 4.25 (t, *J* = 6.7 Hz, 2H), 4.18–4.14 (m, 2H), 1.81–1.75 (m, 4H), 1.68–1.61 (m, 2H) ppm. ^13^C NMR (126 MHz, DMSO): *δ* 156.7, 154.1, 152.4, 149.2, 144.9, 132.1, 130.9, 126.2, 126.1, 125.3, 122.7, 120.6, 117.5, 113.7, 105.1, 96.9, 69.1, 67.8, 26.9, 26.7, 23.2 ppm. HRMS (ESI^+^): *m/z* [M + H]^+^ calcd, 346.1550; found, 346.1555.

### Synthesis of 4-Chloro-6-((6-chlorohexyl)oxy)quinoline-3-carbonitrile (6b)

Triphenyl phosphine (2.23 g, 8.50 mmol, 3.0 equiv) was suspended in 10 mL dry toluene and 5 ml dry DMF. Diisopropyl azodicarboxylate (1.72 g, 8.50 mmol, 3.0 equiv) solved in 10 mL dry toluene was added and the suspension was stirred for 20 min. 6-Chlorohexan-1-ol (567 µL, 4.25 mmol, 1.5 equiv) solved in 10 mL dry toluene and compound **5** (580 mg, 2.83 mmol, 1.0 equiv) solved in 5 mL dry DMF were added dropwise. The reaction mixture was stirred at 100 °C overnight. The solvent was removed *in vacuo* and the residue purified via normal-phase flash chromatography to obtain compound **6b** as a colourless solid (423 mg, 46 %). ^1^H NMR (400 MHz, CDCl3): *δ* 8.83 (s, 1H), 8.14 (d, *J* = 9.2 Hz, 1H), 7.57 (dd, *J* = 9.1, 2.6 Hz, 1H), 7.46 (d, *J* = 2.5 Hz, 1H), 4.17 (t, *J* = 6.3 Hz, 2H), 3.57 (t, *J* = 6.6 Hz, 2H), 1.92 (dd, *J* = 8.9, 4.7 Hz, 2H), 1.84 (dd, *J* = 8.8, 4.8 Hz, 2H), 1.59–1.55 (m, 4H) ppm. LC–MS (ESI^+^): *m*/*z* 323.1 [M + H]^+^.

### Synthesis of 6-((7-Bromoheptyl)oxy)-4-chloroquinoline-3-carbonitrile (6c)

Compound **6c** was synthesized according to the procedure for compound **6b**. Triphenyl phosphine (2.31 g, 8.80 mmol, 3.0 equiv), diisopropyl azodicarboxylate (1.73 mL, 8.80 mmol, 3.0 equiv), 7-bromoheptan-1-ol (676 mL, 4.40 mmol, 1.5 equiv) and compound **5** (600 mg, 2.93 mmol, 1.0 equiv) gave the crude product (526 mg), which was used in the next step without further purification. LC–MS (ESI^+^): *m*/*z* 383.1 [M + H]^+^.

### Synthesis of 4-Chloro-6-(2-(2-(2-chloroethoxy)ethoxy)ethoxy)quinoline-3-carbonitrile (6d)

Compound **6d** was synthesized according to the procedure for compound **6b**. Triphenyl phosphine (2.31 g, 8.80 mmol, 3.0 equiv), diisopropyl azodicarboxylate (1.73 mL, 8.80 mmol, 3.0 equiv), 2-(2-(2-chloroethoxy)ethoxy)ethan-1-ol (639 mL, 4.40 mmol, 1.5 equiv) and compound **5** (600 mg, 2.93 mmol, 1.0 equiv) obtained **6d** as a solid (784 mg, 75%). ^1^H NMR (250 MHz, CDCl3): *δ* 8.82 (s, 1H), 8.09 (d, *J* = 9.2 Hz, 1H), 7.59 (dd, *J* = 9.2, 2.7 Hz, 1H), 7.49 (d, *J* = 2.7 Hz, 1H), 5.00–4.87 (m, 2H), 4.37–4.29 (m, 2H), 4.01–3.93 (m, 2H), 3.78–3.71 (m, 4H), 3.67–3.64 (m, 2H) ppm. LC–MS (ESI^+^): *m*/*z* 323.1 [M + H]^+^.

### Synthesis of 6-((6-Chlorohexyl)oxy)-4-((2-hydroxyphenyl)amino)quinoline-3-carbonitrile (7b)

Compound **6b** (400 mg, 1.24 mmol, 1.0 equiv) and 2-aminophenol (149 mg, 1.36 mmol, 1.1 equiv) were solved in 7 mL *n*-butanol. The reaction mixture was stirred at 130 °C overnight. The solvent was removed *in vacuo* and the crude was purified via normal- phase flash chromatography and subsequent reversed-phase flash chromatography to obtain the product as a beige solid (160 mg, 33%). LC–MS (ESI^+^): *m*/*z* 396.2 [M + H]^+^.

### Synthesis of 6-((7-Chloroheptyl)oxy)-4-((2-hydroxyphenyl)amino)quinoline-3-carbonitrile (7c)

Compound **6c** (150 mg, 393 µmol, 1.0 equiv) and 2-aminophenol (86 mg, 786 µmol, 2.0 equiv) were solved in 4 mL i-propanol. 0.2 mL of a 4 M aqueous HCl solution was added and the reaction mixture was stirred at 100 °C overnight. The solvent was removed *in vacuo* and the crude was purified via normal- phase flash chromatography to obtain **7c** as a brown solid (158 mg, 88%). ^1^H NMR (500 MHz, CDCl3): *δ* 8.31 (s, 1H), 7.90 (d, *J* = 9.0 Hz, 1H), 7.34 (d, *J* = 8.9 Hz, 1H), 7.17–7.13 (m, 1H), 6.99–6.93 (m, Hz, 1H), 6.88 (t, *J* = 7.4 Hz, 1H), 6.80–6.72 (m, 3H), 6.70–6.65 (m, 1H), 3.88–3.80 (m, 2H), 3.54 (t, *J* = 6.7 Hz, 2H), 1.82–1.69 (m, 4H), 1.49–1.41 (m, 4H), 1.39–1.35 (m, 2H) ppm. MS (ESI^+^): *m*/*z* 410.3 [M + H]^+^.

### Synthesis of 6-(2-(2-(2-Chloroethoxy)ethoxy)ethoxy)-4-((2-hydroxyphenyl)amino)quinoline-3-carbonitrile (7d)

Compound **7d** was synthesized according to the procedure for **7c.** Compound **6c** (300 mg, 845 mmol, 1.0 equiv) and 2-aminophenol (184 mg, 1.69 mmol, 2.0 equiv) obtained **7d** as a brown solid (256 mg, 71%). ^1^H NMR (500 MHz, DMSO): *δ* 11.00 (s, 1H), 10.27 (s, 1H), 8.94 (s, 1H), 8.25 (d, *J* = 2.5 Hz, 1H), 8.02 (d, *J* = 9.1 Hz, 1H), 7.72 (dd, *J* = 9.2, 2.4 Hz, 1H), 7.38–7.23 (m, 2H), 7.03 (dd, *J* = 8.2, 1.2 Hz, 1H), 6.92 (td, *J* = 7.6, 1.3 Hz, 1H), 4.33 (dd, *J* = 5.7, 3.4 Hz, 2H), 3.88–3.83 (m, 2H), 3.73–3.64 (m, 4H), 3.65–3.58 (m, 4H) ppm. MS (ESI^+^): *m*/*z* 428.3 [M + H]^+^.

### Synthesis of 4,11-dioxa-2-aza-1(4,6)-quinolina-3(1,2)-benzenacycloundecaphane-1^3^-carbonitrile (8b)

Compound **7b** (160 mg, 404 µmol, 1.0 equiv) and sodium hydride (161 mg, 60% dispersion, 4.04 mmol, 10 equiv) were suspended in 100 mL dry DMF. The reaction mixture was stirred at 60 °C overnight. The reaction mixture was cooled to room temperature and the reaction was quenched with methanol and water. The mixture was concentrated in vacuo and the product was extracted with DCM. The combined organic phases were washed with water and brine, dried over MgSO4, filtered and the solvent was removed in vacuo. The residue was purified via normal-phase flash chromatography to obtain **8b** as solid (15 mg, 10%). ^1^H NMR (500 MHz, CDCl3): *δ* 8.63 (s, 1H), 7.98 (d, *J* = 9.1 Hz, 1H), 7.78 (br, 1H), 7.39 (dd, *J* = 9.1, 2.6 Hz, 1H), 7.37–7.31 (m, 2H), 7.11 (td, *J* = 7.8, 1.5 Hz, 1H), 7.02 (td, *J* = 7.7, 1.4 Hz, 1H), 6.96 (dd, *J* = 8.1, 1.3 Hz, 1H), 4.31–4.23 (m, 2H), 4.17 (t, *J* = 4.9 Hz, 2H), 1.90–1.80 (m, 4H), 1.77–1.69 (m, 2H), 1.51–1.44 (m, 2H) ppm. ^13^C NMR (126 MHz, CDCl3): *δ* 157.7, 149.8, 149.6, 148.3, 131.5, 128.8, 125.8, 124.8, 121.1, 120.5, 120.2, 117.1, 112.3, 100.1, 91.7, 68.9, 67.6, 29.8, 29.8, 29.6, 26.2, 26.1, 23.7 ppm. HRMS (ESI^+^): *m/z* = [M + H]^+^ calcd, 360.1707; found, 360.1708.

### Synthesis of 4,12-dioxa-2-aza-1(4,6)-quinolina-3(1,2)-benzenacyclododecaphane-1^3^-carbonitrile (8c)

Compound **8c** was synthesized according to the procedure for **8b**. Compound **7c** (150 mg, 329 µmol, 1.0 equiv) and sodium hydride (132 mg, 60% dispersion, 3.29 mmol, 10 equiv) obtained **8c** as a yellow solid (63 mg, 51%).^1^H NMR (500 MHz, CDCl3): *δ* 8.66 (s, 1H), 8.02 (d, *J* = 9.1 Hz, 1H), 7.50 (s, 1H), 7.40 (dd, *J* = 9.1, 2.6 Hz, 1H), 7.24 (d, *J* = 2.7 Hz, 1H), 7.13 (dd, *J* = 7.8, 1.5 Hz, 1H), 7.11–7.06 (m, 1H), 7.00–6.95 (m, 2H), 4.28–4.24 (m, 2H), 4.23–4.18 (m, 2H), 1.93–1.84 (m, 4H), 1.70–1.62 (m, 2H), 1.61–1.55 (m, 2H), 1.50–1.44 (m, 2H) ppm. ^13^C NMR (126 MHz, CDCl3): *δ* 157.6, 149.3, 148.5, 147.9, 144.2, 131.5, 128.9, 126.2, 124.3, 121.6, 120.2, 117.9, 116.7, 112.0, 99.8, 93.4, 68.8, 68.0, 29.2, 28.2, 26.7, 25.2, 24.0 ppm. HRMS (ESI^+^): *m/z* = [M + H]^+^ calcd, 374.1863; found, 374.1879.

### Synthesis of 4,12-dioxa-2-aza-1(4,6)-quinolina-3(1,2)-benzenacyclododecaphane-1^3^-carbonitrile (8d)

Compound **8d** was synthesized according to the procedure for **8b**. Compound **7d** (170 mg, 397 µmol, 1.0 equiv) and sodium hydride (95 mg, 60% dispersion, 3.97 mmol, 10 equiv) obtained **8d** as a yellow solid (96 mg, 61%). ^1^H NMR (500 MHz, CDCl3): *δ* 8.57 (s, 1H), 7.98 (d, *J* = 9.1 Hz, 1H), 7.78 (d, *J* = 2.6 Hz, 1H), 7.52 (s, 1H), 7.43 (dd, *J* = 9.1, 2.6 Hz, 1H), 7.29 (dd, *J* = 7.9, 1.5 Hz, 1H), 7.19 (td, *J* = 7.8, 1.5 Hz, 1H), 7.05 (td, *J* = 7.7, 1.3 Hz, 1H), 6.97 (dd, *J* = 8.2, 1.3 Hz, 1H), 4.50–4.46 (m, 2H), 4.25–4.22 (m, 2H), 3.85–3.82 (m, 2H), 3.81–3.78 (m, 2H) ppm. ^13^C NMR (126 MHz, CDCl3): *δ* 159.0, 150.8, 150.2, 149.3, 130.7, 128.3, 126.4, 125.9, 125.8, 122.6, 120.9, 120.7, 116.8, 113.2, 102.7, 89.4, 72.1, 71.1, 71.0, 69.6, 69.5, 68.2 ppm. HRMS (ESI^+^): *m/z* = [M + H]^+^ calcd, 392.1605; found, 392.1612.

### Synthesis of 5-Hydroxy-4-methoxy-2-nitrobenzoic acid methyl ester (10)

Compound **10** was synthesized following the patented conditions of Heymach *et al*.^31^.5-hydroxy-4-methoxy-2-nitrobenzoic acid (**9**, 10.0 g, 46.9 mmol, 1.0 equiv) was dissolved in 35 mL dry MeOH. Concentrated sulfuric acid (4.0 mL, 75 mmol, 1.6 equiv) was added, and the solution was stirred at 85 °C overnight. The solvent was removed *in vacuo*, the residue taken up in water and the product extracted with EtOAc. The combined organic phases were washed with water and brine and dried over MgSO4. The solvent was removed *in vacuo* to obtain **10** as a yellow solid (10.2 g, 96%). ^1^H NMR (400 MHz, DMSO): *δ* 10.92 (s, 1H), 7.61 (s, 1H), 7.07 (s, 1H), 3.90 (s, 3H), 3.79 (s, 3H) ppm. MS (ESI^+^): *m/z* = 249.95 [M+Na]^+^.

### Synthesis of 5-(Benzyloxy)-4-methoxy-2-nitrobenzoic acid methyl ester (11)

Compound **11** was synthesized following the patented conditions of Heymach *et al*.^31^. Compound **10** (10.2 g, 45.1 mmol, 1.0 equiv), 1-(Bromomethyl)benzene (8.1 mL, 68.1 mmol, 1.5 equiv) and K2CO3 (12.5 g, 90.2 mmol, 2.0 equiv) were suspended in 100 mL dry ACN. The suspension was stirred at 80 °C overnight. The solvent was removed *in vacuo*, the residue taken up in water and the product extracted with EtOAc. The combined organic phases were washed with water and brine and dried over MgSO4. The solvent was removed *in vacuo* to obtain **11** as a pale-yellow solid (13.6 g, 95%). ^1^H NMR (400 MHz, DMSO): *δ* 7.66 (s, 1H), 7.48–7.36 (m, 6H), 5.26 (s, 2H), 3.91 (s, 3H), 3.83 (s, 3H) ppm. MS (ESI^+^): *m/z* = 318.05 [M+H]^+^.

### Synthesis of 2-Amino-5-(benzyloxy)-4-methoxybenzoic acid methyl ester (12)

Compound **12** was synthesized following the patented conditions of Wortmann *et al*.^52^. Compound **11** (10.0 g, 31.5 mmol, 1.0 equiv) and NH4Cl (4.36 g, 81.5 mmol, 2.6 equiv) were solved in a mixture of 15 mL water and 80 mL MeOH. Iron powder (10.56 g, 189 mmol, 6.0 equiv) was added batchwise with thorough stirring. The suspension was stirred at 80 °C overnight. The reaction mixture was filtered through Celite and washed with EtOAc. The filtrate was concentrated *in vacuo*, the residue taken up in water and the product extracted with EtOAc. The combined organic phases were washed with water and brine and dried over MgSO4. The solvent was removed *in vacuo* to obtain the crude as a pale-orange solid. The crude product was further converted without additional purification. ^1^H NMR (400 MHz, DMSO): *δ* 7.44–7.34 (m, 5H), 7.24 (s, 1H), 6.47 (s, 2H), 6.39 (s, 1H), 4.91 (s, 2H), 3.76 (s, 3H), 3.73 (s, 3H) ppm. MS (ESI^+^): *m/z* = 288.10 [M+H]^+^.

### Synthesis of Methyl (E)-5-(benzyloxy)-2-(((dimethylamino)methylene)amino)-4-methoxybenzoate (13)

Compound **13** was synthesized following the patented conditions of Heymach *et al*.^31^. Crude product **12** (11.4 g, 31.5 mmol, 1.0 equiv) and DMF dimethyl acetal (12.2 mL, 91.8 mmol, 2.9 equiv) were solved in 35 mL dry DMF. The suspension was stirred at 110 °C for 4 h. The solvent was removed *in vacuo*, the residue taken up in water and the product extracted with EtOAc. The combined organic phases were washed with water and brine and dried over MgSO4. The solvent was removed *in vacuo* to obtain the crude as a dark-purple solid. The crude product was further converted without additional purification. MS (ESI^+^): m/z = 343.15 [M+H]^+^.

### Synthesis of 6-(Benzyloxy)-3-cyano-4-hydroxy-7-methoxyquinoline (14)

Compound **14** was synthesized following the patented conditions of Heymach *et al*.^31^. To a solution of dry ACN (5.50 mL, 105 mmol, 4.2 equiv) diluted with 50 mL dry THF and cooled to –78 °C *n-*butyllithium (26.2 mL, 2.5 M in hexane, 65.5 mmol, 2.6 equiv) was added dropwise over an hour. The suspension was stirred for an additional hour at –78 °C. A solution of crude product **13** solved in 30 mL dry THF was added dropwise to the reaction mixture over 1 h. The reaction mixture was stirred at room temperature overnight. The reaction was quenched by adding concentrated acetic acid (12.0 mL, 209 mmol, 8.3 equiv) and stirred for 1 h. 70 mL water was added, and the suspension was filtered. The precipitate was washed with water and EtOAc. The residue was dried *in vacuo* to give **14** as a pale-brown solid with a yield of 42% over three steps (4.0 g). ^1^H NMR (400 MHz, DMSO): *δ* 8.57 (s, 1H), 7.54 (s, 1H), 7.48–7.32 (m, 5H), 7.05 (s, 1H), 5.18 (s, 2H), 3.88 (s, 3H) ppm. MS (ESI^+^): *m/z* = 307.05 [M+H]^+^.

### Synthesis of 6-(Benzyloxy)-4-chloro-3-cyano-7-methoxyquinoline (15)

Compound **15** was synthesized following the patented conditions of Heymach *et al*.^31^. Compound **14** (4.0 g, 13 mmol, 1.0 equiv) was solved in 40 mL dry toluene. Phosphoryl chloride (8.0 mL, 86 mmol, 6.6 equiv) was added and the reaction mixture was stirred at 120 °C overnight. The solvent was removed *in vacuo*, the residue taken up in a 5% aqueous solution of NaOH and the product extracted with DCM. The combined organic phases were washed with water and brine and dried over MgSO4. The solvent was removed *in vacuo* to obtain **15** as a beige solid (3.74 g, 88%). ^1^H NMR (400 MHz, DMSO): *δ* 8.95 (s, 1H), 7.57–7.51 (m, 4H), 7.46–7.37 (m, 3H), 5.33 (s, 2H), 4.01 (s, 3H) ppm. MS (ESI^+^): *m/z* = 325.05 [M+H]^+^.

### Synthesis of 4-Chloro-3-cyano-6-hydroxy-7-methoxyquinoline (16)

Core building block **16** was synthesized following the patented conditions of Heymach *et al*.^31^. Compound **15** (1.0 g, 3.1 mmol, 1.0 equiv) was solved in 20 mL TFA. Thioanisole (6.5 mL, 56 mmol, 18 equiv) was added and the reaction mixture was stirred at 80 °C for 4 h. The solvent was removed *in vacuo*. The residue was stirred with a 10% aqueous ammonia solution. The precipitate was filtered and washed with EtOAc. The residue was dried *in vacuo* to obtain **16** as a beige solid (440 mg, 61%). ^1^H NMR (400 MHz, DMSO): *δ* 10.74 (s, 1H), 8.90 (s, 1H), 7.51 (s, 1H), 7.45 (s, 1H), 4.02 (s, 3H) ppm. MS (ESI^+^): *m/z* = 234.95 [M+H]^+^.

### Synthesis of 3-Cyano-4-((2-hydroxyphenyl)amino)-6-hydroxy-7-methoxyquinoline (17)

Core building block **16** (200 mg, 852 μmol, 1.0 equiv) and 2-aminophenol (112 mg, 1.02 mmol, 1.2 equiv) were solved in 6 mL 2-ethoxyethanol in a microwave vial. 0.1 mL of a 4 M aqueous HCl solution was added and the vial was sealed. The microwave assisted-reaction was stirred for 40 min at 130 °C. The solvent was removed *in vacuo* and the crude was purified via reversed-phase flash chromatography to obtain **17** as a pale-yellow solid (181 mg, 69%). ^1^H NMR (400 MHz, DMSO): *δ* 9.66 (s, 1H), 9.62 (s, 1H), 9.00 (s, 1H), 8.26 (s, 1H), 7.72 (s, 1H), 7.28 (s, 1H), 7.17–7.10 (m, 2H), 6.92–6.87 (m, 1H), 6.84–6.78 (m, 1H), 3.97 (s, 3H) ppm. ^13^C NMR (101 MHz, DMSO): *δ* 153.4, 152.6, 150.4, 150.0, 146.6, 144.8, 128.2, 127.8, 125.9, 119.0, 117.5, 115.9, 113.3, 108.6, 105.7, 85.7, 55.9 ppm. MS (ESI^+^): *m/z* = 308.05 [M+H]^+^.

### Synthesis of 4-((3-Chloro-2-hydroxyphenyl)amino)-3-cyano-6-hydroxy-7-methoxy-quinoline (18)

4-Anilino-quinoline **18** was synthesized according to the procedure for **17**. Core building block **16** (160 mg, 682 μmol, 1.0 equiv) and 2-amino-6-chlorophenol (117 mg, 818 μmol, 1.2 equiv) obtained **18** as a yellow solid (135 mg, 58%). ^1^H NMR (400 MHz, DMSO): *δ* 10.62–10.16 (m, 2H), 10.09 (br, 1H), 8.80 (s, 1H), 7.90 (s, 1H), 7.45 (dd, *J* = 8.0, 1.6 Hz, 1H), 7.36 (s, 1H), 7.26 (dd, *J* = 8.0, 1.6 Hz, 1H), 6.92 (t, *J* = 8.0 Hz, 1H), 4.02 (s, 3H) ppm. ^13^C NMR (101 MHz, DMSO): *δ* 154.7, 152.3, 148.2, 147.2, 133.0, 130.0, 123.5, 119.1, 118.4, 115.7, 115.4, 114.9, 112.7, 106.5, 102.8, 85.4, 56.4 ppm. MS (ESI^+^): *m/z* = 341.95 [M+H]^+^.

### Synthesis of 4-((4-Chloro-2-hydroxyphenyl)amino)-3-cyano-6-hydroxy-7-methoxy-quinoline (19)

4-Anilino-quinoline **19** was synthesized according to the procedure for **17**. Core building block **16** (150 mg, 639 μmol, 1.0 equiv) and 2-amino-5-chlorophenol (110 mg, 767 μmol, 1.2 equiv) obtained **19** as a bright-yellow solid (166 mg, 76%). ^1^H NMR (400 MHz, DMSO): *δ* 10.69 (br, 1H), 10.57–10.10 (m, 2H), 8.83 (s, 1H), 7.91 (s, 1H), 7.36 (s, 1H), 7.31 (d, *J* = 8.4, 1H), 7.00 (s, 1H), 6.96 (d, *J* = 8.4, 1H), 4.02 (s, 3H) ppm. ^13^C NMR (101 MHz, DMSO): *δ* 154.7, 154.7, 152.3, 148.2, 147.2, 133.0, 130.0, 123.5, 119.1, 118.4, 115.7, 114.9, 112.7, 106.5, 102.8, 85.4, 56.4 ppm. MS (ESI^+^): *m/z* = 342.05 [M+H]^+^.

### Synthesis of 4-((2-Chloro-6-hydroxyphenyl)amino)-3-cyano-6-hydroxy-7-methoxy-quinoline (20)

4-Anilino-quinoline **20** was synthesized according to the procedure for **17**. The reaction time was extended to 4 h. Core building block **16** (170 mg, 725 μmol, 1.0 equiv) and 2-amino-3-chlorophenol (125 mg, 869 μmol, 1.2 equiv) obtained **20** as an orangish solid (110 mg, 44%). ^1^H NMR (400 MHz, DMSO): *δ* 10.53–10.16 (m, 2H), 10.08 (br, 1H), 8.78 (s, 1H), 7.90 (s, 1H), 7.44 (d, *J* = 8.1 Hz, 1H), 7.39 (s, 1H), 7.26 (d, *J* = 7.8 Hz, 1H), 6.92 (t, *J* = 8.0, 1.5 Hz, 1H), 4.01 (s, 3H) ppm. ^13^C NMR (101 MHz, DMSO): *δ* 154.6, 152.5, 150.1, 148.0, 147.4, 129.8, 127.7, 126.6, 121.4, 119.9, 115.4, 114.8, 113.0, 106.7, 103.0, 85.3, 56.4 ppm. MS (ESI^+^): *m/z* = 342.05 [M+H]^+^.

### Synthesis of 3-Cyano-6-hydroxy-4-((2-hydroxy-5-methoxyphenyl)amino)-7-methoxy-quinoline (21)

4-Anilino-quinoline **21** was synthesized according to the procedure for **17**. Core building block **16** (300 mg, 1.28 mmol, 1.0 equiv) and 2-amino-4-methoxyphenol (214 mg, 1.53 mmol, 1.2 equiv) obtained **21** as a yellow solid (328 mg, 76%). ^1^H NMR (400 MHz, DMSO): *δ* 10.45–10.25 (m, 2H), 9.58 (br, 1H), 8.82 (s, 1H), 7.91 (s, 1H), 7.33 (s, 1H), 6.87 (m, 3H), 4.01 (s, 3H), 3.70 (s, 3H) ppm. ^13^C NMR (101 MHz, DMSO): *δ* 154.8, 152.4, 152.0, 148.2, 147.6, 146.9, 135.6, 124.0, 116.5, 115.4, 114.6, 113.5, 112.6, 106.7, 102.3, 85.6, 56.4, 55.6 ppm. MS (ESI^+^): *m/z* = 338.05 [M+H]^+^.

### Synthesis of 3-Cyano-6-hydroxy-4-((2-hydroxy-3-methoxyphenyl)amino)-7-methoxy-quinoline (22)

4-Anilino-quinoline **22** was synthesized according to the procedure for **17**. Core building block **16** (150 mg, 639 μmol, 1.0 equiv) and 2-amino-6-methoxyphenol (107 mg, 767 μmol, 1.2 equiv) obtained **22** as a yellow solid (95 mg, 44%). ^1^H NMR (400 MHz, DMSO): *δ* 9.66 (s, 1H), 9.03 (s, 1H), 8.93 (s, 1H), 8.28 (s, 1H), 7.73 (s, 1H), 7.29 (s, 1H), 6.95–6.89 (m, 1H), 6.81–6.74 (m, 2H), 3.97 (s, 3H), 3.82 (s, 3H) ppm. ^13^C NMR (101 MHz, DMSO): *δ* 152.7, 150.3, 150.0, 148.1, 146.7, 144.7, 142.6, 126.3, 120.0, 118.3, 117.4, 113.4, 110.5, 108.5, 105.8, 86.0, 56.1, 55.9 ppm. MS (ESI^+^): *m/z* = 338.10 [M+H]^+^.

### Synthesis of 1^7^-methoxy-4,10-dioxa-2-aza-1(4,6)-quinolina-3(1,2)-benzenacyclodecaphane-1^3^-carbonitrile (23a)

4-Anilino-quinoline **17** (80 mg, 260 μmol, 1.0 equiv) and K2CO3 (360 mg, 2.60 mmol, 10 equiv) were suspended in 1.5 mL dry DMF per 1 mg 4-anilino-quionline. The reaction mixture was stirred at 80 °C. After 15 min 1,5-dibromopentane (66 mg, 286 μmol, 1.1 equiv), diluted in 5 mL dry DMF, was added dropwise to the reaction mixture. The reaction mixture was stirred for 3 h. The solvent was removed *in vacuo* and the crude was purified via reversed-phase flash chromatography and subsequent preparative HPLC to obtain compound **23a** as a yellow solid (16 mg, 16%). ^1^H NMR (500 MHz, DMSO): *δ* 8.68 (s, 1H), 8.09 (s, 1H), 7.86 (s, 1H), 7.37 (s, 1H), 7.26 (dd, *J* = 7.7, 1.6 Hz, 1H), 7.13 (td, *J* = 8.0, 1.6 Hz, 1H), 7.08 (dd, *J* = 8.1, 1.4 Hz, 1H), 6.89 (td, *J* = 7.7, 1.4 Hz, 1H), 4.24 (t, *J* = 6.6 Hz, 2H), 4.13 (t, *J* = 4.8 Hz, 2H), 3.94 (s, 3H), 1.79–1.70 (m, 4H), 1.70–1.63 (m, 2H) ppm. ^13^C NMR (126 MHz, DMSO): *δ* 154.7, 153.6, 152.3, 149.9, 148.2, 146.9, 132.3, 125.9, 125.8, 120.5, 117.8, 116.7, 113.4, 108.6, 107.0, 96.0, 68.9, 68.4, 56.0, 26.9, 26.6, 23.1 ppm. HRMS (ESI^+^): *m/z* = [M + H]^+^ calcd, 376.1656; found, 376.1671.

### Synthesis of 1^7^-methoxy-4,11-dioxa-2-aza-1(4,6)-quinolina-3(1,2)-benzenacycloundecaphane-1^3^-carbonitrile (23b)

Compound **23b** was synthesized according to the procedure for **23a**. 4-Anilino-quinoline **17** (50 mg, 163 μmol, 1.0 equiv), K2CO3 (225 mg, 1.63 mmol, 10 equiv) and 1-chloro-6-iodohexane (48 mg, 195 μmol, 1.1 equiv) obtained **23b** as a yellow solid (12 mg, 19%). ^1^H NMR (400 MHz, DMSO): *δ* 8.61 (s, 1H), 8.36 (s, 1H), 8.14 (s, 1H), 7.50 (d, *J* = 7.9 Hz, 1H), 7.32–7.28 (m, 2H), 7.02–6.96 (m, 2H), 4.51 (t, *J* = 6.2 Hz, 2H), 3.94 (s, 3H), 3.87–3.80 (m, 2H), 1.69–1.60 (m, 2H), 1.59–1.50 (m, 2H), 1.24–1.17 (m, 2H), 1.11–1.02 (m, 2H) ppm. ^13^C NMR (101 MHz, DMSO): *δ* 154.7, 154.1, 152.9, 150.8, 148.3, 145.7, 130.5, 128.8, 128.3, 119.9, 117.6, 113.6, 112.5, 108.8, 103.9, 88.0, 68.3, 65.2, 55.9, 29.0, 28.5, 24.3, 23.5 ppm. HRMS (ESI^+^): *m/z* = [M + H]^+^ calcd, 390.1812; found, 390.1812.

### Synthesis of 1^7^-methoxy-4,12-dioxa-2-aza-1(4,6)-quinolina-3(1,2)-benzenacyclododecaphane-1^3^-carbonitrile (23c)

Compound **23c** was synthesized according to the procedure for **23a**. 4-Anilino-quinoline **17** (50 mg, 163 μmol, 1.0 equiv), K2CO3 (225 mg, 1.63 mmol, 10 equiv) and 1,7-dibromoheptane (46 mg, 179 μmol, 1.1 equiv) obtained **23c** as a yellow solid (25 mg, 38%). ^1^H NMR (400 MHz, DMSO): *δ* 8.64 (s, 1H), 8.47 (s, 1H), 7.78–7.74 (m, 1H), 7.34 (s, 1H), 7.21–7.13 (m, 3H), 6.98–6.93 (m, 1H), 4.33 (t, *J* = 7.6 Hz, 2H), 4.11 (t, *J* = 5.0 Hz, 2H), 3.94 (s, 3H), 1.72–1.64 (m, 2H), 1.60–1.53 (m, 2H), 1.42–1.35 (m, 4H), 1.34–1.28 (m, 2H) ppm. ^13^C NMR (101 MHz, DMSO): *δ* 154.3, 152.0, 150.8, 149.3, 147.5, 145.9, 129.5, 126.0, 123.8, 120.6, 117.5, 114.5, 113.8, 109.0, 104.3, 88.7, 69.5, 67.0, 55.9, 28.7, 27.1, 25.8, 25.5, 23.6 ppm. HRMS (ESI^+^): *m/z* = [M + H]^+^ calcd, 404.1969; found, 404.1971.

### Synthesis of 1^7^-methoxy-4,7,10,13-tetraoxa-2-aza-1(4,6)-quinolina-3(1,2)-benzenacyclotridecaphane-1^3^-carbonitrile (23d)

Compound **23d** was synthesized according to the procedure for **23a**. Additional purification via preparative HPLC was performed. 4-Anilino-quinoline **17** (90 mg, 293 μmol, 1.0 equiv), K2CO3 (405 mg, 2.93 mmol, 10 equiv) and 1,2-bis(2-bromoethoxy)ethane (89 mg, 322 μmol, 1.1 equiv) obtained **23d** as a yellow TFA salt (19 mg, 15%). ^1^H NMR (400 MHz, DMSO): *δ* 10.55 (s, 1H), 8.88 (s, 1H), 8.33 (s, 1H), 7.49–7.43 (m, 2H), 7.39 (s, 1H), 7.18 (d, *J* = 7.3 Hz, 1H), 7.10 (td, *J* = 7.6, 1.2 Hz, 1H), 4.64 (s, 2H), 4.10 (t, *J* = 4.1 Hz, 2H), 3.99 (s, 3H), 3.74 (t, *J* = 4.2 Hz, 2H), 3.46–3.35 (m, 4H), 3.25–3.16 (m, 2H) ppm. ^13^C NMR (101 MHz, DMSO): *δ* 158.3 (q, *J* = 33.6 Hz), 155.6, 154.7, 153.9, 149.6, 147.2, 135.1, 130.4, 129.3, 125.5, 120.8, 114.1, 113.3, 111.5, 106.6, 101.2, 84.9, 71.4, 70.3, 69.8, 68.52, 68.5, 67.0, 56.3 ppm. HRMS (ESI^+^): *m/z* [M + H]^+^ calcd, 422.1711; found, 422.1729.

### Synthesis of 1^7^-methoxy-4,7,10-trioxa-2-aza-1(4,6)-quinolina-3(1,2)-benzenacyclodecaphane-1^3^-carbonitrile (23e)

Compound **23e** was synthesized according to the procedure for **23a**. Additional purification via preparative HPLC was performed. 4-Anilino-quinoline **17** (75 mg, 244 μmol, 1.0 equiv), K2CO3 (337 mg, 2.44 mmol, 10 equiv) and 2-bromoethyl ether (62 mg, 268 μmol, 1.1 equiv) obtained **23e** as a yellow TFA salt (38 mg, 41%). ^1^H NMR (400 MHz, DMSO): *δ* 8.96 (s, 1H), 8.11 (s, 1H), 7.44 (s, 1H), 7.36 (d, *J* = 7.6 Hz, 1H), 7.33–7.23 (m, 2H), 7.00 (t, *J* = 7.6 Hz, 1H), 4.38–4.27 (m, 4H), 3.97 (s, 3H), 3.67–3.60 (m, 2H), 3.56–3.49 (m, 2H) ppm. ^13^C NMR (101 MHz, DMSO): *δ* 158.5 (q, *J* = 34.8 Hz), 156.1, 154.5, 153.7, 147.7, 147.5, 139.9, 129.5, 127.8, 127.0, 120.9, 115.6, 115.2, 114.0, 109.1, 103.4, 90.7, 70.8, 69.2, 69.1, 68.8, 56.3 ppm. HRMS (ESI^+^): *m/z* = [M + H]^+^ calcd, 378.1448; found, 378.1455.

### Synthesis of 3^3^-chloro-1^7^-methoxy-4,10-dioxa-2-aza-1(4,6)-quinolina-3(1,2)-benzenacyclodecaphane-1^3^-carbonitrile (24a)

Compound **24a** was synthesized according to the procedure for **23a**. 4-Anilino-quinoline **18** (75 mg, 219 μmol, 1.0 equiv), K2CO3 (303 mg, 2.19 mmol, 10 equiv) and 1,5-dibromopentane (56 mg, 241 μmol, 1.1 equiv) obtained **24a** as a yellow solid (14 mg, 16%). ^1^H NMR (500 MHz, DMSO): *δ* 8.84 (s, 1H), 8.74 (s, 1H), 7.45 (s, 1H), 7.20 (s, 1H), 7.07 (dd, *J* = 8.1, 1.5 Hz, 1H), 6.96 (t, *J* = 8.1 Hz, 1H), 6.70 (dd, *J* = 8.1, 1.5 Hz, 1H), 4.34–4.26 (m, 2H), 4.16–4.10 (m, 2H), 3.95 (s, 3H), 2.03–1.95 (m, 2H), 1.75–1.67 (m, 2H), 1.47–1.40 (m, 2H) ppm. ^13^C NMR (126 MHz, DMSO): *δ* 154.4, 149.5, 148.9, 148.0, 147.0, 143.6, 138.2, 128.6, 125.1, 122.7, 117.7, 117.3, 117.1, 108.9, 103.4, 99.1, 70.7, 66.3, 56.1, 24.4, 22.5, 19.0 ppm. HRMS (ESI^+^): *m/z* = [M + H]^+^ calcd, 410.1266; found, 410.1271.

### Synthesis of 3^3^-chloro-1^7^-methoxy-4,11-dioxa-2-aza-1(4,6)-quinolina-3(1,2)-benzenacycloundecaphane-1^3^-carbonitrile (24b)

Compound **24b** was synthesized according to the procedure for **23a**. 4-Anilino-quinoline **18** (90 mg, 263 μmol, 1.0 equiv), K2CO3 (364 mg, 2.63 mmol, 10 equiv) and 1,6-dibromohexane (71 mg, 290 μmol, 1.1. equiv) obtained **24b** as a yellow solid (26 mg, 23%). ^1^H NMR (500 MHz, DMSO): *δ* 9.05 (s, 1H), 8.46 (s, 1H), 7.85 (s, 1H), 7.38–7.33 (m, 2H), 7.28 (d, *J* = 7.9 Hz, 1H), 7.14 (t, *J* = 8.0 Hz, 1H), 4.35 (t, *J* = 6.9 Hz, 2H), 4.02 (t, *J* = 6.6, 2H), 3.96 (s, 3H), 1.56–1.46 (m, 4H), 1.37–1.30 (m, 4H) ppm. ^13^C NMR (126 MHz, DMSO): *δ* 155.0, 150.8, 150.0, 149.9, 146.6, 135.5, 127.3, 127.0, 125.7, 124.7, 117.0, 114.0, 108.7, 107.1, 88.9, 73.8, 67.2, 56.0, 27.3, 25.0, 22.8, 22.4 ppm. HRMS (ESI^+^): *m/z* = [M + H]^+^ calcd, 424.1423; found, 424.1430.

### Synthesis of 3^3^-chloro-1^7^-methoxy-4,12-dioxa-2-aza-1(4,6)-quinolina-3(1,2)-benzenacyclododecaphane-1^3^-carbonitrile (24c)

Compound **24c** was synthesized according to the procedure for **23a**. 4-Anilino-quinoline **18** (75 mg, 219 μmol, 1.0 equiv), K2CO3 (303 mg, 2.19 mmol, 10 equiv) and 1,7-dibromoheptane (62 mg, 241 μmol, 1.1 equiv) obtained **24c** as a yellow solid (21 mg, 22%). ^1^H NMR (500 MHz, DMSO): *δ* 9.67 (s, 1H), 8.44 (s, 1H), 8.09 (s, 1H), 7.50 (dd, *J* = 8.0, 1.6 Hz, 1H), 7.41 (dd, *J* = 8.0, 1.6 Hz, 1H), 7.34 (s, 1H), 7.22 (t, *J* = 8.0 Hz, 1H), 4.53–4.43 (m, 2H), 3.96 (s, 3H), 3.92 (t, *J* = 6.6 Hz, 2H), 1.67–1.57 (m, 2H), 1.42–1.31 (m, 4H), 1.19–1.09 (m, 2H), 1.09–1.05 (m, 2H) ppm. ^13^C NMR (126 MHz, DMSO): *δ* 154.6, 151.2, 150.7, 150.6, 148.3, 144.0, 134.1, 129.5, 128.3, 127.4, 124.9, 116.4, 112.2, 108.0, 105.9, 85.8, 73.0, 66.4, 56.0, 28.7, 28.2, 26.4, 24.2, 23.7 ppm. HRMS (ESI^+^): *m/z* = [M + H]^+^ calcd, 438.1579; found, 438.1574.

### Synthesis of 3^4^-chloro-1^7^-methoxy-4,10-dioxa-2-aza-1(4,6)-quinolina-3(1,2)-benzenacyclodecaphane-1^3^-carbonitrile (25a)

Compound **25a** was synthesized according to the procedure for **23a**. Additional purification via preparative HPLC was performed. 4-Anilino-quinoline **19** (100 mg, 293 μmol, 1.0 equiv), K2CO3 (404 mg, 2.93 mmol, 10 equiv) and 1,5-dibromopentane (74 mg, 322 μmol, 1.1 equiv) obtained **25a** as a yellow solid (12 mg, 10%). ^1^H NMR (500 MHz, DMSO): *δ* 8.69 (s, 1H), 8.10 (s, 1H), 7.87 (s, 1H), 7.39 (s, 1H), 7.28 (d, *J* = 8.4 Hz, 1H), 7.16 (d, *J* = 2.3 Hz, 1H), 6.96 (dd, *J* = 8.4, 2.3 Hz, 1H), 4.26 (t, *J* = 6.6 Hz, 2H), 4.14 (t, *J* = 4.5 Hz, 2H), 3.95 (s, 3H), 1.79–1.65 (m, 6H) ppm. ^13^C NMR (126 MHz, DMSO): *δ* 154.9, 153.4, 153.1, 149.8, 148.4, 147.0, 131.4, 129.5, 126.9, 120.2, 117.7, 116.9, 113.5, 108.6, 107.2, 96.4, 69.6, 68.7, 56.0, 27.0, 26.6, 23.1 ppm. HRMS (ESI^+^): *m/z* = [M + H]^+^ calcd, 410.1266; found, 410.1266.

### Synthesis of 3^4^-chloro-1^7^-methoxy-4,11-dioxa-2-aza-1(4,6)-quinolina-3(1,2)-benzenacycloundecaphane-1^3^-carbonitrile (25b)

Compound **25b** was synthesized according to the procedure for **23a**. 4-Anilino-quinoline **19** (100 mg, 293 μmol, 1.0 equiv), K2CO3 (404 mg, 2.93 mmol, 10 equiv) and 1,6-dibromohexane (79 mg, 322 μmol, 1.1 equiv) obtained **25b** as a yellow solid (39 mg, 31%). ^1^H NMR (500 MHz, DMSO): *δ* 8.71 (s, 1H), 8.42 (s, 1H), 8.15 (s, 1H), 7.52 (d, *J* = 8.3 Hz, 1H), 7.32 (s, 1H), 7.11 (d, *J* = 2.3 Hz, 1H), 7.06 (dd, *J* = 8.3, 2.3 Hz, 1H), 4.52 (t, *J* = 6.2 Hz, 2H), 3.95 (s, 3H), 3.87 (t, *J* = 4.8 Hz, 2H), 1.68–1.63 (m, 2H), 1.56– 1.49 (m, 2H), 1.21–1.15 (m, 2H), 1.09–1.03 (m, 2H) ppm. ^13^C NMR (126 MHz, DMSO): *δ* 155.5, 154.4, 152.9, 150.4, 148.5, 145.2, 132.2, 131.6, 128.0, 119.8, 117.4, 113.7, 113.1, 108.3, 104.0, 88.3, 69.0, 65.2, 55.9, 28.9, 28.5, 24.2, 23.5 ppm. HRMS (ESI^+^): *m/z* = [M + H]^+^ calcd, 424.1423; found, 424.1425.

### Synthesis of 3^4^-chloro-1^7^-methoxy-4,12-dioxa-2-aza-1(4,6)-quinolina-3(1,2)-benzenacyclododecaphane-1^3^-carbonitrile (25c)

Compound **25c** was synthesized according to the procedure for **23a**. Additional purification via normal-phase flash chromatography was performed. 4-Anilino-quinoline **19** (100 mg, 293 μmol, 1.0 equiv), K2CO3 (404 mg, 2.93 mmol, 10 equiv) and 1,7-dibromoheptane (83 mg, 322 μmol, 1.1 equiv) obtained **25c** as a bright-yellow solid (44 mg, 34%). ^1^H NMR (500 MHz, DMSO): *δ* 8.63 (s, 1H), 8.52 (s, 1H), 7.71 (s, 1H), 7.36 (s, 1H), 7.25 (d, *J* = 2.3 Hz, 1H), 7.17 (d, *J* = 8.5 Hz, 1H), 6.99 (dd, *J* = 8.5, 2.3 Hz, 1H), 4.33 (t, *J* = 7.7 Hz, 2H), 4.17–4.12 (m, 2H), 3.95 (s, 3H), 1.72–1.65 (m, 2H), 1.61–1.54 (m, 2H) 1.42–1.30 (m, 6H) ppm. ^13^C NMR (126 MHz, DMSO): *δ* 154.5, 152.3, 150.5, 149.1, 147.7, 145.9, 129.1, 129.0, 124.0, 120.4, 117.2, 114.7, 114.2, 108.8, 104.3, 89.7, 70.1, 67.2, 56.0, 28.6, 27.0, 25.8, 25.4, 23.6 ppm. HRMS (ESI^+^): *m/z* = [M + H]^+^ calcd, 438.1579; found, 438.1579.

### Synthesis of 3^6^-chloro-1^7^-methoxy-4,11-dioxa-2-aza-1(4,6)-quinolina-3(1,2)-benzenacycloundecaphane-1^3^-carbonitrile (26b)

Compound **26b** was synthesized according to the procedure for **23a**. Additional purification via preparative HPLC was performed. 4-Anilino-quinoline **20** (110 mg, 322 μmol, 1.0 equiv), K2CO3 (445 mg, 3.22 mmol, 10 equiv) and 1,6-dibromohexane (94 mg, 386 μmol, 1.1 equiv) obtained **26b** as a yellow TFA salt (16 mg, 12%). ^1^H NMR (400 MHz, DMSO): *δ* 10.08 (s, 1H), 8.94 (s, 1H), 8.38 (s, 1H), 7.48 (t, *J* = 8.3 Hz, 1H), 7.44 (s, 2H), 7.25 (dd, *J* = 8.3, 1.1 Hz, 1H), 7.09 (dd, *J* = 8.3, 1.1 Hz, 1H), 4.89–4.80 (m, 1H), 4.38–4.30 (m, 1H), 4.20–4.14 (m, 1H), 3.63 (t, *J* = 10.3 Hz, 1H), 1.96–1.84 (m, 1H), 1.83–1.71 (m, 1H), 1.60–1.44 (m, 2H), 1.34–1.26 (m, 1H), 1.19–1.05 (m, 1H), 0.99–0.85 (m, 1H), 0.84– 0.71 (m, 1H) ppm. ^13^C NMR (101 MHz, DMSO): *δ* 158.3 (q, *J* = 36.1 Hz), 157.0, 156.4, 155.8, 149.6, 147.5, 136.5, 135.3, 131.2, 124.6, 121.1, 113.8, 112.8, 111.9, 104.8, 102.3, 86.2, 69.6, 65.4, 56.5, 29.0, 28.6, 24.2, 23.5 ppm. HRMS (ESI^+^): *m/z* = [M + H]^+^ calcd, 424.1423; found, 424.1425.

### Synthesis of 1^7^,3^5^-dimethoxy-4,10-dioxa-2-aza-1(4,6)-quinolina-3(1,2)-benzenacyclo-decaphane-1^3^-carbonitrile (27a)

Compound **27a** was synthesized according to the procedure for **23a**. 4-Anilino-quinoline **21** (150 mg, 445 μmol, 1.0 equiv), K2CO3 (615 mg, 4.45 mmol, 10 equiv) and 1,5-dibromopentane (112 mg, 489 μmol, 1.1 equiv) obtained **27a** as a colourless solid (23 mg, 13%). ^1^H NMR (500 MHz, DMSO): *δ* 8.75 (s, 1H), 8.11 (s, 1H), 7.76 (s, 1H), 7.40 (s, 1H), 7.00 (d, *J* = 8.9 Hz, 1H), 6.81–6.73 (m, 1H), 6.64 (dd, *J* = 8.9, 3.0 Hz, 1H), 4.25 (t, *J* = 7.1 Hz, 2H), 4.10 (t, *J* = 5.3 Hz, 2H), 3.95 (s, 3H), 3.66 (s, 3H), 1.84–1.75 (m, 4H), 1.66–1.59 (m, 2H) ppm. ^13^C NMR (126 MHz, DMSO): *δ* 154.7, 153.4, 152.6, 149.7, 148.5, 146.9, 144.9, 133.7, 117.9, 117.2, 115.0, 109.8, 108.8, 108.5, 105.9, 97.4, 69.7, 68.2, 56.0, 55.3, 26.3, 26.1, 22.7 ppm. HRMS (ESI^+^): *m/z* = [M + H]^+^ calcd, 406.1761; found, 406.1760.

### Synthesis of 1^7^,3^5^-dimethoxy-4,11-dioxa-2-aza-1(4,6)-quinolina-3(1,2)-benzenacyclo-undecaphane-1^3^-carbonitrile (27b)

Compound **27b** was synthesized according to the procedure for **23a**. 4-Anilino-quinoline **21** (100 mg, 296 μmol, 1.0 equiv), K2CO3 (410 mg, 2.96 mmol, 10 equiv) and 1,6-dibromohexane (80 mg, 326 μmol, 1.1 equiv) obtained **27b** as a yellow solid (35 mg, 28%). ^1^H NMR (500 MHz, DMSO): δ 9.37 (s, 1H), 8.65 (s, 1H), 8.19 (s, 1H), 7.37 (s, 1H), 7.14 (d, *J* = 3.0 Hz, 1H), 6.98 (d, *J* = 8.9 Hz, 1H), 6.91 (dd, *J* = 8.9, 3.0 Hz, 1H), 4.52 (t, *J* = 6.4 Hz, 2H), 3.97 (s, 3H), 3.84 (t, *J* = 5.0 Hz, 2H), 3.75 (s, 3H), 1.68–1.61 (m, 2H), 1.56–1.49 (m, 2H), 1.28–1.22 (m, 2H), 1.13–1.06 (m, 2H). ^13^C NMR (126 MHz, DMSO): δ 155.2, 153.5, 152.8, 149.1, 148.7, 128.4, 116.2, 115.8, 113.9, 113.7, 113.3, 105.6, 104.7, 87.7, 69.4, 65.7, 56.2, 55.7, 29.0, 28.1, 24.5, 23.5 ppm. HRMS (ESI^+^): *m/z* = [M + H]^+^ calcd, 420.1918; found, 420.1926.

### Synthesis of 1^7^,3^5^-dimethoxy-4,12-dioxa-2-aza-1(4,6)-quinolina-3(1,2)-benzenacyclo-dodecaphane-1^3^-carbonitrile (27c)

Compound **27c** was synthesized according to the procedure for **23a**. Additional purification via preparative HPLC was performed. 4-Anilino-quinoline **21** (100 mg, 296 μmol, 1.0 equiv), K2CO3 (410 mg, 2.96 mmol, 10 equiv) and 1,7-dibromoheptane (84 mg, 326 μmol, 1.1 equiv) obtained **27c** as a yellow solid (22 mg, 17%). ^1^H NMR (500 MHz, DMSO): δ 9.00 (s, 1H), 8.51 (s, 1H), 7.84 (s, 1H), 7.36 (s, 1H), 7.08 (d, *J* = 8.8 Hz, 1H), 6.82–6.73 (m, 2H), 4.38 (t, *J* = 7.3 Hz, 2H), 4.04–3.99 (m, 2H), 3.95 (s, 3H), 3.71 (s, 3H), 1.70–1.63 (m, 2H), 1.53–1.47 (m, 2H), 1.40–1.34 (m, 2H), 1.31–1.25 (m, 4H) ppm.^13^C NMR (126 MHz, DMSO): *δ* 154.5, 153.5, 150.4, 149.7, 147.8, 146.4, 130.5, 117.0, 116.6, 113.6, 111.3, 110.3, 108.1, 104.5, 88.6, 70.7, 66.8, 59.8, 56.0, 55.5, 28.6, 26.9, 26.0, 25.8, 23.7 ppm. HRMS: *m/z* = [M + H]^+^ calcd, 434.2074; found, 434.2078.

### Synthesis of 1^7^,3^3^-dimethoxy-4,11-dioxa-2-aza-1(4,6)-quinolina-3(1,2)-benzenacyclo-undecaphane-1^3^-carbonitrile (28b)

Compound **28b** was synthesized according to the procedure for **23a**. Additional purification via preparative HPLC was performed. 4-Anilino-quinoline **22** (80 mg, 237 μmol, 1.0 equiv), K2CO3 (328 mg, 2.37 mmol, 10 equiv) and 1,6-dibromohexane (69 mg, 285 μmol, 1.1 equiv) obtained **28b** as a colourless TFA salt (24 mg, 23%). ^1^H NMR (400 MHz, DMSO): *δ* 10.10 (s, 1H), 8.82 (s, 1H), 8.12 (s, 1H), 7.42 (s, 1H), 7.18–7.06 (m, 3H), 4.48 (t, *J* = 6.6 Hz, 2H), 4.05–3.92 (m, 5H), 3.81 (s, 3H), 1.63– 1.51 (m, 2H), 1.44–1.28 (m, 4H), 1.25–1.16 (m, 2H) ppm. ^13^C NMR (101 MHz, DMSO): *δ* 158.2 (q, *J* = 33.3 Hz), 156.1, 153.2, 152.5, 148.4, 148.0, 144.3, 138.3, 131.0, 123.3, 120.7, 114.6, 113.1, 112.5, 106.6, 103.7, 86.6, 73.8, 66.7, 56.3, 56.0, 28.8, 26.6, 24.2, 23.1 ppm. HRMS (ESI^+^): *m/z* = [M + H]^+^ calcd, 420.1918; found, 420.1928.

### Synthesis of 1^7^,3^3^-Dimethoxy-4,12-dioxa-2-aza-1(4,6)-quinolina-3(1,2)-benzenacyclododecaphane-1^3^-carbonitrile (28c)

Compound **28c** was synthesized according to the procedure for **23a**. 4-Anilino-quinoline **22** (100 mg, 296 μmol, 1.0 equiv), K2CO3 (410 mg, 2.96 mmol, 10 equiv) and 1,7-dibromoheptane (61 µL, 356 μmol, 1.2 equiv) obtained **28c** as a pale-yellow solid (22 mg, 17%). ^1^H NMR (400 MHz, DMSO): *δ* 9.17 (s, 1H), 8.30 (s, 1H), 7.97 (s, 1H), 7.30 (s, 1H), 7.12–7.03 (m, 2H), 6.98–6.91 (m, 1H), 4.51–4.38 (m, 2H), 3.95–3.88 (m, 5H), 3.82 (s, 3H), 1.65–1.57 (m, 2H), 1.39–1.30 (m, 4H), 1.19–1.13 (m, 2H), 1.10–1.03 (m, 2H) ppm. ^13^C NMR (101 MHz, DMSO): *δ* 154.0, 153.0, 151.3, 150.5, 147.8, 145.5, 143.7, 132.8, 123.5, 120.6, 117.0, 112.3, 112.3, 109.1, 105.2, 85.8, 71.9, 66.0, 56.0, 55.8, 28.9, 28.0, 26.5, 25.1, 23.7 ppm. HRMS: *m/z* = [M + H]^+^ calcd, 434.2074; found, 434.2068.

### Synthesis of 1^7^,3^3^-Dimethoxy-4,7,10-trioxa-2-aza-1(4,6)-quinolina-3(1,2)-benzenacyclodecaphane-1^3^-carbonitrile (AZ137, 28e)

AZ137 (**28e**) was synthesized according to the procedure for **23a**. 4-Anilino-quinoline **22** (110 mg, 326 μmol, 1.0 equiv), K2CO3 (451 mg, 3.26 mmol, 10 equiv) and 1-bromo-2-(2-bromoethoxy)ethane (49 µL, 391 μmol, 1.2 equiv) obtained AZ137 (**28e**) as a colorless solid (24 mg, 18%). ^1^H NMR (400 MHz, DMSO): *δ* 8.72 (s, 1H), 8.27–8.18 (m, 2H), 7.39 (s, 1H), 6.94 (t, *J* = 8.2 Hz, 1H), 6.74 (d, *J* = 8.2 Hz, 1H), 6.65 (d, *J* = 8.0 Hz, 1H), 4.42–4.33 (m, 4H), 3.85 (s, 3H), 3.82–3.77 (m, 2H), 3.77–3.71 (m, 2H) ppm. ^13^C NMR (101 MHz, DMSO): *δ* 154.2, 151.6, 150.0, 148.1, 148.0, 146.6, 137.4, 134.2, 122.6, 117.3, 116.1, 109.8, 108.4, 106.9, 105.3, 94.0, 73.7, 70.6, 70.4, 69.9, 56.0, 56.0 ppm. HRMS (ESI^+^): *m/z* = [M + H]^+^ calcd, 408.1554; found, 408.1567.

### Synthesis of 6-Ethoxy-4-((2-ethoxy-3-methoxyphenyl)amino)-7-methoxyquinoline-3-carbonitrile (29)

4-Anilino-quinoline **22** (30 mg, 89 μmol, 1.0 equiv) and K2CO3 (61 mg, 445 mmol, 5.0 equiv) were suspended in 4 mL dry DMF. The reaction mixture was stirred at 80 °C. After 15 min iodoethane (16 µL, 196 μmol, 2.2 equiv), diluted in 1 mL dry DMF, was added dropwise to the reaction mixture. The reaction mixture was stirred at room temperature for 3 h. The solvent was removed *in vacuo* and the crude was purified via reversed-phase flash chromatography to obtain compound **29** as a colorless solid (10 mg, 29%). ^1^H NMR (500 MHz, DMSO): *δ* 9.23 (s, 1H), 8.36 (s, 1H), 7.81 (s, 1H), 7.30 (s, 1H), 7.08 (t, *J* = 8.1 Hz, 1H), 6.99 (dd, *J* = 8.4, 1.4 Hz, 1H), 6.88 (dd, *J* = 7.9, 1.3 Hz, 1H), 4.16 (q, *J* = 7.0 Hz, 2H), 3.94 (s, 3H), 3.87–3.81 (m, 5H), 1.40 (t, *J* = 7.0 Hz, 3H), 0.87 (t, *J* = 7.0 Hz, 3H) ppm. ^13^C NMR (126 MHz, DMSO): *δ* 153.3, 153.1, 150.8, 150.3, 148.3, 145.6, 143.3, 133.4, 123.6, 119.3, 117.2, 113.0, 111.2, 108.7, 102.5, 87.2, 67.8, 64.3, 56.0, 55.8, 39.5, 15.2, 14.5 ppm. HRMS (ESI^+^): *m/z* = [M + H]^+^ calcd, 394.1761; found, 394.1765.

### Synthesis of 4-Chloro-6,7-dihydroxyquinoline-3-carbonitrile (33)

4-Chloro-7-(3-chloropropoxy)-6-methoxyquinoline-3-carbonitrile (**32**, 1.00 g, 3.21 mmol, 1.0 equiv) and tertbutyl ammonium iodide (1.42 g, 3.86 mmol, 1.2 equiv) were solved in 10 mL dry toluene. The solution was cooled to 0 °C. Boron trichloride (11 mL, 1 M in toluene, 11.0 mmol, 3.4 equiv) was added slowly. The reaction mixture was stirred at 100 °C for 5 h. The reaction was quenched by adding 100 mL water and stirring for 30 min. The precipitate was filtered and washed with EtOAc. The residue was dried *in vacuo* to obtain compound **33** as a yellow solid (640 mg, 90%). ^1^H NMR (300 MHz, DMSO): *δ* 11.08 (br, 1H), 10.81 (br, 1H), 8.81 (s, 1H), 7.43 (s, 1H), 7.36 (s, 1H) ppm. LC–MS (ESI^+^): *m*/*z* = 221.00 [M + H]^+^.

### Synthesis of 6,7-Dihydroxy-4-((2-hydroxy-3-methoxyphenyl)amino)quinoline-3-carbonitrile (34)

The title compound **34** was synthesized according to the procedure for 4-anilino-quinoline **17**. Additionally, after all reagents were added, the reaction mixture was degassed under argon for 15 min. Compound **33** (210 mg, 952 μmol, 1.0 equiv) and 2-amino-6-methoxyphenol (159 mg, 1.14 mmol, 1.2 equiv) obtained the product as a bright-yellow solid (166 mg, 54%). ^1^H NMR (400 MHz, DMSO): *δ* 11.88 (br, 1H), 10.49 (s, 1H), 10.39 (br, 1H) 9.39 (s, 1H), 8.82 (s, 1H), 7.94 (s, 1H), 7.30 (s, 1H), 7.05 (dd, *J* = 7.9, 1.8 Hz, 1H), 6.90 (dd, *J* = 8.1, 1.8 Hz, 1H), 6.85 (t, *J* = 7.9 Hz, 1H), 3.84 (s, 3H) ppm. LC–MS (ESI^+^): *m*/*z* = 324.10 [M + H]^+^.

### Synthesis of 3^3^-Methoxy-1^7^-(2-(2-(4-methylpiperazin-1-yl)ethoxy)ethoxy)-4,7,10-trioxa-2-aza-1(4,6)-quinolina-3(1,2)-benzenacyclodecaphane-1^3^-carbonitrile (30)

Compound **34** (20.0 mg, 61.9 µmol, 1.0 equiv), K2CO3 (59.9 mg, 433 µmol, 7.0 equiv) and 1-bromo-2-(2-bromoethoxy)ethane (8 µL, 63.6 μmol, 1.0 equiv) were suspended in 20 mL dry DMF. After the reaction mixture was degassed under argon for 15 min the reaction mixture was stirred at 80 °C for 2 h. 1-Bromo-2-(2-bromoethoxy)ethane (8 µL, 63.6 μmol, 1.0 equiv) added and the reaction mixture was stirred for an additional hour at 80 °C. Afterwards the reaction mixture was cooled to 60 °C and 1-methylpiperazine (137 µL, 1.24 mmol, 20 equiv) was added and the reaction mixture stirred for 4 h. The solvent was removed *in vacuo* and the crude was purified via reversed-phase flash chromatography and subsequent preparative HPLC to obtain the product as a yellow solid (7.2 mg, 21%). ^1^H NMR (500 MHz, DMSO): *δ* 8.71 (s, 1H), 8.22 (d, *J* = 20.7 Hz, 2H), 7.42 (s, 1H), 6.94 (t, *J* = 8.2 Hz, 1H), 6.74 (dd, *J* = 8.4, 1.4 Hz, 1H), 6.65 (dd, *J* = 8.2, 1.3 Hz, 1H), 4.41–4.36 (m, 4H), 4.32–4.28 (m, 2H), 3.85 (s, 3H), 3.80 (t, *J* = 4.2 Hz, 4H), 3.76–3.73 (m, 2H), 3.60 (t, *J* = 5.8 Hz, 2H), 2.54 (s, 2H), 2.48–2.30 (m, 8H), 2.20 (s, 3H) ppm. ^13^C NMR (126 MHz, DMSO): *δ* 153.4, 151.60, 150.0, 148.0, 147.9, 146.5, 137.4, 134.2, 122.6, 117.3, 116.1, 109.7, 109.1, 106.8, 105.3, 94.0, 73.7, 70.5, 70.4, 69.9, 68.4, 68.3, 68.2, 57.0, 56.0, 54.3, 52.6, 45.2 ppm. HRMS: *m/z* = [M + H]^+^ calcd, 564.2817; found, 564.2826.

### Synthesis of 4-((5-chloro-2-hydroxyphenyl)amino)-6-((5-hydroxypentyl)oxy)quinoline-3-carbonitrile (35)

Compound **35** was synthesized according to the procedure for **7a.** Compound **6a** (200 mg, 494 mmol, 1.0 equiv) and 2-amino-4-chlorophenol (142 mg, 988 µmol, 2.0 equiv) obtained **35** as a yellow solid (196 mg, 60%). ^1^H NMR (500 MHz, DMSO): *δ* 10.11 (s, 1H), 9.39 (s, 1H), 8.38 (s, 1H), 7.84 (d, *J* = 2.7 Hz, 1H), 7.82 (d, *J* = 9.1 Hz, 1H), 7.44 (dd, *J* = 9.1, 2.6 Hz, 1H), 7.31 (d, *J* = 2.6 Hz, 1H), 7.22 (dd, *J* = 8.7, 2.7 Hz, 1H), 6.94 (d, *J* = 8.7 Hz, 1H), 4.39 (t, *J* = 5.1 Hz, 1H), 4.12 (t, *J* = 6.5 Hz, 2H), 3.47– 3.38 (m, 2H), 1.81 (quint, *J* = 6.7 Hz, 2H), 1.57–1.40 (m, 4H) ppm. ^13^C NMR (126 MHz, DMSO): *δ* 156.9, 152.6, 150.3, 150.0, 143.7, 130.9, 127.7, 127.7, 126.7, 123.2, 122.0, 119.2, 117.2, 102.6, 86.8, 68.3, 60.6, 32.2, 30.4, 28.5, 22.2 ppm. MS (ESI^+^): *m*/*z* 398.4 [M + H]^+^.

### Synthesis of 3^5^-chloro-4,10-dioxa-2-aza-1(4,6)-quinolina-3(1,2)-benzenacyclodecaphane-1^3^-carbonitrile (31)

Compound **31** was synthesized according to the procedure for **8a.** Compound **35** (90 mg, 226 µmol, 1.0 equiv), diisopropyl azodicarboxylate (229 mg, 1.13 µmol, 5.0 equiv) and triphenyl phosphine (297 mg, 1.13 µmol, 5.0 equiv) obtained **31** as a yellow solid (10 mg, 12%). ^1^H NMR (500 MHz, DMSO): *δ* 8.80 (s, 1H), 8.30 (s, 1H), 7.94 (d, *J* = 9.1 Hz, 1H), 7.75 (d, *J* = 2.8 Hz, 1H), 7.46 (dd, *J* = 9.1, 2.7 Hz, 1H), 7.22–7.18 (m, 1H), 7.14–7.13 (m, 2H), 4.26 (t, *J* = 7.0 Hz, 2H), 4.22–4.19 (m, 2H), 1.85–1.79 (m, 4H), 1.67–1.61 (m, 2H) ppm. ^13^C NMR (126 MHz, DMSO): *δ* 157.1, 153.2, 150.1, 149.0, 145.1, 134.2, 131.0, 125.6, 124.4, 124.2, 123.4, 123.2, 117.4, 115.1, 104.5, 99.5, 69.8, 68.1, 26.5, 26.3, 23.2 ppm. HRMS: *m/z* = [M + H]^+^ calcd, 380.1160; found, 380.1169.

## Associated content Supporting Information

Supporting Information Available, including: Synthetic procedures, DSF kinase selectivity panel, NanoBRET data, X ray data collection and refinement statistics, and HPLC and NMR spectra for all compounds. Molecular Formular Strings are available.

## Data availability

The coordinates and structure factors of the CLK3 complex with macrocyclic inhibitor **23e** and **30** have been deposited in the Protein Data Bank (PDB) under accession code 29MP & 29MO.

## Author contributions

A.Z., N.D.R., and T.H. designed the project; A.Z. and N.D.R. synthesized the compounds; Z.Z. performed ADP-Glo assays, A.K. expressed, purified and co-crystallized CLK3; A.K., refined and analyzed the structures of CLK3 inhibitor complexes; A.K., L.E. performed DSF assays; J.D.,L.E. and B.-T.B performed NanoBRET assays; R.T. performed actin staining assays M.P.S. and J.F. performed K192-screens; N.D.R performed the kinetic solubility assay; J.K.C, S.K. and T.H. supervised the research. The manuscript was written by A.Z., N.D.R., and T.H. with contributions from all authors.

## Acknowledgments

The authors acknowledge support by the Structural Genomics Consortium (SGC), a registered charity (no: 1097737) that receives funds from Bayer AG, Boehringer Ingelheim, Bristol Meyer Squibb, Genentech, Genome Canada through Ontario Genomics Institute [OGI-196], EU/EFPIA/OICR/McGill/KTH/Diamond Innovative Medicines Initiative 2 Joint Undertaking [EUbOPEN grant 875510], Janssen, Pfizer and Takeda. S.K. is funded by the German Cancer Research Center (DKTK) and the Frankfurt Cancer Institute (FCI) as well as the DKH (Krebshilfe) translational research network TACTIC and the BMFTR grant PREVENT (01GR2501A). This work was further supported by the National Institutes of Health (R35 GM127030 to J.K.C.), a Male Contraceptive Initiative Agility Grant (to J.K.C.), a Male Contraceptive Initiative Postdoctoral Fellowship (to Z.Z.), and the Stanford Molecular Pharmacology Training Program (T32 GM136631 to R.K.T.). In addition, we would like to thank Hanna Holzmann for her support with the CellTiter-Glo assay, Nikos To for his support with DSF selectivity panels and Astrid Kaiser for her support with the thermodynamic solubility assay.

## Notes

B.-T.B. is a cofounder and the CEO of the Contract Research Organization CELLinib GmbH, Frankfurt, Germany. The other authors declare no conflict of interest.

## Abbreviations

CLK: CDC-like kinase
DSF: differential scanning fluorimetry
ESI: electrospray ionization
GAK: cyclin G-associated kinase
HIPK: homeodomain-interacting protein kinase
LC–MS: liquid chromatography–mass spectrometry
NanoBRET: nanoluciferase-based bioluminescence resonance energy transfer
DYRK2: dual-specificity tyrosine-phosphorylation-regulated kinase 2
BMP2K: BMP-2-inducible kinase
YSK4: Mitogen-activated protein kinase kinase kinase 19

---

For Table of Contents Only

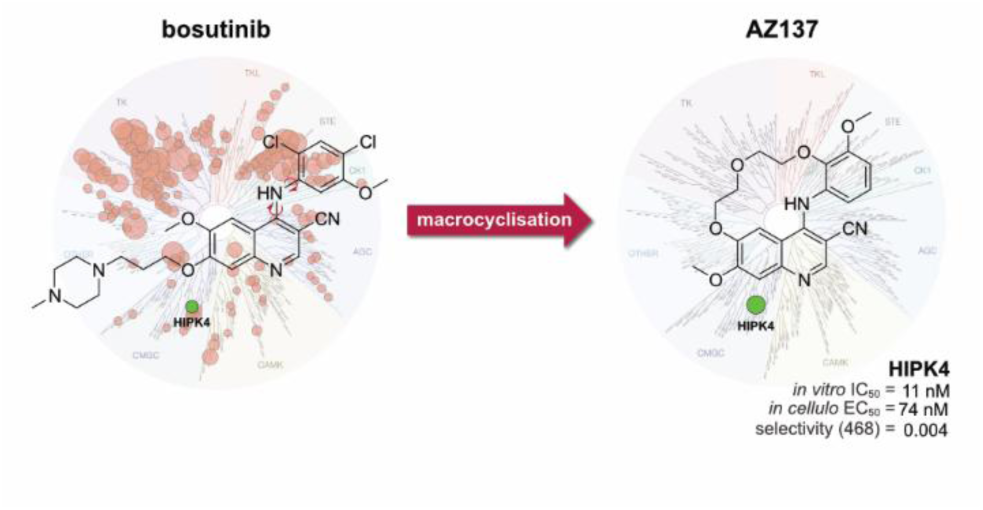

## REFERENCES

(1) Manning, G.; Whyte, D. B.; Martinez, R.; Hunter, T.; Sudarsanam, S. The Protein Kinase Complement of the Human Genome. Science 2002, 298 (5600), 1912–1934. 10.1126/science.1075762.

(2) Arai, S.; Matsushita, A.; Du, K.; Yagi, K.; Okazaki, Y.; Kurokawa, R. Novel Homeodomain-interacting Protein Kinase Family Member, HIPK4, Phosphorylates Human P53 at Serine 9. FEBS Lett. 2007, 581 (29), 5649–5657. 10.1016/j.febslet.2007.11.022.

(3) Kim, Y. H.; Choi, C. Y.; Lee, S.-J.; Conti, M. A.; Kim, Y. Homeodomain-Interacting Protein Kinases, a Novel Family of Co-Repressors for Homeodomain Transcription Factors*. J. Biol. Chem. 1998, 273 (40), 25875–25879. 10.1074/jbc.273.40.25875.

(4) Choi, C. Y.; Kim, Y. H.; Kwon, H. J.; Kim, Y. The Homeodomain Protein NK-3 Recruits Groucho and a Histone Deacetylase Complex to Repress Transcription. J. Biol. Chem. 1999, 274 (47), 33194–33197. 10.1074/jbc.274.47.33194.

(5) Kondo, S.; Lu, Y.; Debbas, M.; Lin, A. W.; Sarosi, I.; Itie, A.; Wakeham, A.; Tuan, J.; Saris, C.; Elliott, G.; Ma, W.; Benchimol, S.; Lowe, S. W.; Mak, T. W.; Thukral, S. K. Characterization of Cells and Gene-Targeted Mice Deficient for the P53-Binding Kinase Homeodomain-Interacting Protein Kinase 1 (HIPK1). Proc. Natl. Acad. Sci. U. S. A. 2003, 100 (9), 5431–5436. 10.1073/pnas.0530308100.

(6) D’Orazi, G.; Cecchinelli, B.; Bruno, T.; Manni, I.; Higashimoto, Y.; Saito, S.; Gostissa, M.; Coen, S.; Marchetti, A.; Del Sal, G.; Piaggio, G.; Fanciulli, M.; Appella, E.; Soddu, S. Homeodomain-Interacting Protein Kinase-2 Phosphorylates P53 at Ser 46 and Mediates Apoptosis. Nat. Cell Biol. 2002, 4 (1), 11–19. 10.1038/ncb714.

(7) Rinaldo, C.; Siepi, F.; Prodosmo, A.; Soddu, S. HIPKs: Jack of All Trades in Basic Nuclear Activities. Biochim. Biophys. Acta BBA - Mol. Cell Res. 2008, 1783 (11), 2124–2129. 10.1016/j.bbamcr.2008.06.006.

(8) Isono, K.; Nemoto, K.; Li, Y.; Takada, Y.; Suzuki, R.; Katsuki, M.; Nakagawara, A.; Koseki, H. Overlapping Roles for Homeodomain-Interacting Protein Kinases Hipk1 and Hipk2 in the Mediation of Cell Growth in Response to Morphogenetic and Genotoxic Signals. Mol. Cell. Biol. 2006, 26 (7), 2758–2771. 10.1128/MCB.26.7.2758-2771.2006.

(9) Calzado, M. A.; Renner, F.; Roscic, A.; Schmitz, M. L. HIPK2, a Versatile Switchboard Regulating the Transcription Machinery and Cell Death. Cell Cycle 2007, 6 (2), 139–143. 10.4161/cc.6.2.3788.

(10) Hofmann, T. G.; Möller, A.; Sirma, H.; Zentgraf, H.; Taya, Y.; Dröge, W.; Will, H.; Schmitz, M. L. Regulation of P53 Activity by Its Interaction with Homeodomain-Interacting Protein Kinase-2. Nat. Cell Biol. 2002, 4 (1), 1–10. 10.1038/ncb715.

(11) Zhang, Q.; Yoshimatsu, Y.; Hildebrand, J.; Frisch, S. M.; Goodman, R. H. Homeodomain Interacting Protein Kinase 2 Promotes Apoptosis by Downregulating the Transcriptional Corepressor CtBP. Cell 2003, 115 (2), 177–186. 10.1016/s0092-8674(03)00802-x.

(12) Rochat-Steiner, V.; Becker, K.; Micheau, O.; Schneider, P.; Burns, K.; Tschopp, J. Fist/Hipk3. J. Exp. Med. 2000, 192 (8), 1165–1174. 10.1084/jem.192.8.1165.

(13) Van Der Laden, J.; Soppa, U.; Becker, W. Effect of Tyrosine Autophosphorylation on Catalytic Activity and Subcellular Localisation of Homeodomain-Interacting Protein Kinases (HIPK). Cell Commun. Signal. 2015, 13 (1), 3. 10.1186/s12964-014-0082-6.

(14) Crapster, J. A.; Rack, P. G.; Hellmann, Z. J.; Le, A. D.; Adams, C. M.; Leib, R. D.; Elias, J. E.; Perrino, J.; Behr, B.; Li, Y.; Lin, J.; Zeng, H.; Chen, J. K. HIPK4 Is Essential for Murine Spermiogenesis. eLife 2020, 9, e50209. 10.7554/eLife.50209.

(15) Liu, X.; Zang, C.; Wu, Y.; Meng, R.; Chen, Y.; Jiang, T.; Wang, C.; Yang, X.; Guo, Y.; Situ, C.; Hu, Z.; Zhang, J.; Guo, X. Homeodomain-Interacting Protein Kinase HIPK4 Regulates Phosphorylation of Manchette Protein RIMBP3 during Spermiogenesis. J. Biol. Chem. 2022, 298 (9), 102327. 10.1016/j.jbc.2022.102327.

(16) Larribère, L.; Galach, M.; Novak, D.; Arévalo, K.; Volz, H. C.; Stark, H.-J.; Boukamp, P.; Boutros, M.; Utikal, J. An RNAi Screen Reveals an Essential Role for HIPK4 in Human Skin Epithelial Differentiation from iPSCs. Stem Cell Rep. 2017, 9 (4), 1234–1245. 10.1016/j.stemcr.2017.08.023.

(17) Guo, Z.; Chen, B.; Zhang, M.; Gao, M.; Wang, Z.; Tang, H.; Tang, X.; Zhang, Q.; Utikal, J. HIPK4 Accelerates Cutaneous Squamous Cell Carcinoma Progression by Phosphorylating TAp63 and Inhibiting EFEMP1 Expression. J. Biol. Chem. 2025, 301 (7), 108564. 10.1016/j.jbc.2025.108564.

(18) Davis, M. I.; Hunt, J. P.; Herrgard, S.; Ciceri, P.; Wodicka, L. M.; Pallares, G.; Hocker, M.; Treiber, D. K.; Zarrinkar, P. P. Comprehensive Analysis of Kinase Inhibitor Selectivity. Nat. Biotechnol. 2011, 29 (11), 1046–1051. 10.1038/nbt.1990.

(19) Zarrinkar, P. P.; Gunawardane, R. N.; Cramer, M. D.; Gardner, M. F.; Brigham, D.; Belli, B.; Karaman, M. W.; Pratz, K. W.; Pallares, G.; Chao, Q.; Sprankle, K. G.; Patel, H. K.; Levis, M.; Armstrong, R. C.; James, J.; Bhagwat, S. S. AC220 Is a Uniquely Potent and Selective Inhibitor of FLT3 for the Treatment of Acute Myeloid Leukemia (AML). Blood 2009, 114 (14), 2984–2992. 10.1182/blood-2009-05-222034.

(20) Steinbach, PharmD, A.; Clark, PharmD, S. M.; Clemmons, PharmD, Bcop, A. B. Bosutinib: A Novel Src/Abl Kinase Inhibitor for Chronic Myelogenous Leukemia. J. Adv. Pract. Oncol. 2013, 4 (6). 10.6004/jadpro.2013.4.6.8.

(21) Amrhein, J. A.; Knapp, S.; Hanke, T. Synthetic Opportunities and Challenges for Macrocyclic Kinase Inhibitors. J. Med. Chem. 2021, 64 (12), 7991–8009. 10.1021/acs.jmedchem.1c00217.

(22) Mallinson, J.; Collins, I. Macrocycles In New Drug Discovery. Future Med. Chem. 2012, 4 (11), 1409–1438. 10.4155/fmc.12.93.

(23) Zou, H. Y.; Friboulet, L.; Kodack, D. P.; Engstrom, L. D.; Li, Q.; West, M.; Tang, R. W.; Wang, H.; Tsaparikos, K.; Wang, J.; Timofeevski, S.; Katayama, R.; Dinh, D. M.; Lam, H.; Lam, J. L.; Yamazaki, S.; Hu, W.; Patel, B.; Bezwada, D.; Frias, R. L.; Lifshits, E.; Mahmood, S.; Gainor, J. F.; Affolter, T.; Lappin, P. B.; Gukasyan, H.; Lee, N.; Deng, S.; Jain, R. K.; Johnson, T. W.; Shaw, A. T.; Fantin, V. R.; Smeal, T. PF-06463922, an ALK/ROS1 Inhibitor, Overcomes Resistance to First and Second Generation ALK Inhibitors in Preclinical Models. Cancer Cell 2015, 28 (1), 70–81. 10.1016/j.ccell.2015.05.010.

(24) Al-Fayoumi, S.; Campbell, M. S.; Amberg, S.; Zhou, H.; Millard, L.; Dean, J. P. Clinical Pharmacology Profile of Pacritinib (PAC), a Novel JAK2/FLT3 Inhibitor. Blood 2015, 126 (23), 5178–5178. 10.1182/blood.V126.23.5178.5178.

(25) De, S. K. First Approval of Pacritinib as a Selective Janus Associated Kinase-2 Inhibitor Forthe Treatment of Patients with Myelofibrosis. Anticancer Agents Med. Chem. 2023, 23 (12), 1355–1360. 10.2174/1871520623666230320120915.

(26) Letunic, I.; Bork, P. Interactive Tree of Life (iTOL) v6: Recent Updates to the Phylogenetic Tree Display and Annotation Tool. Nucleic Acids Res. 2024, 52 (W1), W78–W82. 10.1093/nar/gkae268.

(27) Amrhein, J. A.; Beyett, T. S.; Feng, W. W.; Krämer, A.; Weckesser, J.; Schaeffner, I. K.; Rana, J. K.; Jänne, P. A.; Eck, M. J.; Knapp, S.; Hanke, T. Macrocyclization of Quinazoline-Based EGFR Inhibitors Leads to Exclusive Mutant Selectivity for EGFR L858R and Del19. J. Med. Chem. 2022, 65 (23), 15679–15697. 10.1021/acs.jmedchem.2c01041.

(28) Amrhein, J. A.; Berger, L. M.; Balourdas, D.-I.; Joerger, A. C.; Menge, A.; Krämer, A.; Frischkorn, J. M.; Berger, B.-T.; Elson, L.; Kaiser, A.; Schubert-Zsilavecz, M.; Müller, S.; Knapp, S.; Hanke, T. Synthesis of Pyrazole-Based Macrocycles Leads to a Highly Selective Inhibitor for MST3. J. Med. Chem. 2024, 67 (1), 674–690. 10.1021/acs.jmedchem.3c01980.

(29) Amrhein, J. A.; Wang, G.; Berger, B.-T.; Berger, L. M.; Kalampaliki, A. D.; Krämer, A.; Knapp, S.; Hanke, T. Design and Synthesis of Pyrazole-Based Macrocyclic Kinase Inhibitors Targeting BMPR2. ACS Med. Chem. Lett. 2023, 14 (6), 833–840. 10.1021/acsmedchemlett.3c00127.

(30) Dettenhöfer, M.; Tandara, L. N.; Amrhein, J. A.; Kurz, C. G.; Schwalm, M. P.; Mensing, T. E.; Wahl, L. M.; Krämer, A.; Gerninghaus, J.; Lenz, C.; Elson, L.; Berger, B.-T.; Schröder, M.; Saxena, K.; Müller, S.; Knapp, S.; Greco, F. A.; Hanke, T. Un-LOK-Ing a New Approach for Conformational Selective Targeting of STK10 (LOK). ACS Med. Chem. Lett. 2025, 16 (11), 2086–2096. 10.1021/acsmedchemlett.5c00479.

(31) Heymach, J.; Robichaux, J.; Nilsson, M.; Jones, P.; Cross, J.; Theroff, J. Heterocyclic Inhibitors of Tyrosine Kinase. WO2020219904A1.

(32) Amrhein, J. A.; Berger, L. M.; Tjaden, A.; Krämer, A.; Elson, L.; Tolvanen, T.; Martinez-Molina, D.; Kaiser, A.; Schubert-Zsilavecz, M.; Müller, S.; Knapp, S.; Hanke, T. Discovery of 3-Amino-1H-Pyrazole-Based Kinase Inhibitors to Illuminate the Understudied PCTAIRE Family. Int. J. Mol. Sci. 2022, 23 (23), 14834. 10.3390/ijms232314834.

(33) Klaeger, S.; Heinzlmeir, S.; Wilhelm, M.; Polzer, H.; Vick, B.; Koenig, P.-A.; Reinecke, M.; Ruprecht, B.; Petzoldt, S.; Meng, C.; Zecha, J.; Reiter, K.; Qiao, H.; Helm, D.; Koch, H.; Schoof, M.; Canevari, G.; Casale, E.; Depaolini, S. R.; Feuchtinger, A.; Wu, Z.; Schmidt, T.; Rueckert, L.; Becker, W.; Huenges, J.; Garz, A.-K.; Gohlke, B.-O.; Zolg, D. P.; Kayser, G.; Vooder, T.; Preissner, R.; Hahne, H.; Tõnisson, N.; Kramer, K.; Götze, K.; Bassermann, F.; Schlegl, J.; Ehrlich, H.-C.; Aiche, S.; Walch, A.; Greif, P. A.; Schneider, S.; Felder, E. R.; Ruland, J.; Médard, G.; Jeremias, I.; Spiekermann, K.; Kuster, B. The Target Landscape of Clinical Kinase Drugs. Science 2017, 358 (6367), eaan4368. 10.1126/science.aan4368.

(34) Schwalm, M. P.; Knapp, S. Single-Plate Kinome Screening in Live-Cells to Enable Highly Cost-Efficient Kinase Inhibitor Profiling. SLAS Discov. Adv. Life Sci. R D 2025, 31, 100214. 10.1016/j.slasd.2025.100214.

(35) Gerninghaus, J.; Zhubi, R.; Krämer, A.; Karim, M.; Tran, D. H. N.; Joerger, A. C.; Schreiber, C.; Berger, L. M.; Berger, B.-T.; Ehret, T. A. L.; Elson, L.; Lenz, C.; Saxena, K.; Müller, S.; Einav, S.; Knapp, S.; Hanke, T. Back-Pocket Optimization of 2-Aminopyrimidine-Based Macrocycles Leads to Potent EPHA2/GAK Kinase Inhibitors. J. Med. Chem. 2024, 67 (15), 12534–12552. 10.1021/acs.jmedchem.4c00411.

(36) Mensing, T. E.; Kurz, C. G.; Amrhein, J. A.; Ehret, T. A. L.; Preuss, F.; Mathea, S.; Karim, M.; Tran, D. H. N.; Kadlecova, Z.; Tolvanen, T. A.; Martinez-Molina, D.; Müller, S.; Einav, S.; Knapp, S.; Hanke, T. Development of Pyrazolo[1,5-a]Pyrimidine Based Macrocyclic Kinase Inhibitors Targeting AAK1. Eur. J. Med. Chem. 2025, 299, 118076. 10.1016/j.ejmech.2025.118076.

(37) Walz, C.; Spiske, M.; Walter, M.; Keller, B.-L.; Mezler, M.; Hoft, C.; Pohlki, F.; Vukelić, S.; Hausch, F. Macrocyclization as a Strategy for Kinetic Solubility Improvement: A Comparative Analysis of Matched Molecular Pairs. J. Med. Chem. 2025, 68 (3), 2639–2656. 10.1021/acs.jmedchem.4c01822.

(38) Müller, S.; Ackloo, S.; Arrowsmith, C. H.; Bauser, M.; Baryza, J. L.; Blagg, J.; Böttcher, J.; Bountra, C.; Brown, P. J.; Bunnage, M. E.; Carter, A. J.; Damerell, D.; Dötsch, V.; Drewry, D. H.; Edwards, A. M.; Edwards, J.; Elkins, J. M.; Fischer, C.; Frye, S. V.; Gollner, A.; Grimshaw, C. E.; IJzerman, A.; Hanke, T.; Hartung, I. V.; Hitchcock, S.; Howe, T.; Hughes, T. V.; Laufer, S.; Li, V. M.; Liras, S.; Marsden, B. D.; Matsui, H.; Mathias, J.; O’Hagan, R. C.; Owen, D. R.; Pande, V.; Rauh, D.; Rosenberg, S. H.; Roth, B. L.; Schneider, N. S.; Scholten, C.; Singh Saikatendu, K.; Simeonov, A.; Takizawa, M.; Tse, C.; Thompson, P. R.; Treiber, D. K.; Viana, A. Y.; Wells, C. I.; Willson, T. M.; Zuercher, W. J.; Knapp, S.; Mueller-Fahrnow, A. Donated Chemical Probes for Open Science. eLife 2018, 7, e34311. 10.7554/eLife.34311.

(39) Workman, P.; Collins, I. Probing the Probes: Fitness Factors For Small Molecule Tools. Chem. Biol. 2010, 17 (6), 561–577. 10.1016/j.chembiol.2010.05.013.

(40) Dopfer, J.; Vasta, J. D.; Müller, S.; Knapp, S.; Robers, M. B.; Schwalm, M. P. tracerDB: A Crowdsourced Fluorescent Tracer Database for Target Engagement Analysis. Nat. Commun. 2024, 15 (1), 5646. 10.1038/s41467-024-49896-5.

(41) Kabsch, W. *XDS*. *Acta Crystallogr*. D Biol. Crystallogr. 2010, 66 (2), 125–132. 10.1107/S0907444909047337.

(42) Winter, G.; Waterman, D. G.; Parkhurst, J. M.; Brewster, A. S.; Gildea, R. J.; Gerstel, M.; Fuentes-Montero, L.; Vollmar, M.; Michels-Clark, T.; Young, I. D.; Sauter, N. K.; Evans, G. *DIALS* : Implementation and Evaluation of a New Integration Package. Acta Crystallogr. Sect. Struct. Biol. 2018, 74 (2), 85–97. 10.1107/S2059798317017235.

(43) Winter, G.; Lobley, C. M. C.; Prince, S. M. Decision Making in *Xia* 2. Acta Crystallogr. D Biol. Crystallogr. 2013, 69 (7), 1260–1273. 10.1107/S0907444913015308.

(44) Vonrhein, C.; Flensburg, C.; Keller, P.; Sharff, A.; Smart, O.; Paciorek, W.; Womack, T.; Bricogne, G. Data Processing and Analysis with the *autoPROC* Toolbox. Acta Crystallogr. D Biol. Crystallogr. 2011, 67 (4), 293–302. 10.1107/S0907444911007773.

(45) Evans, P. R.; Murshudov, G. N. How Good Are My Data and What Is the Resolution? Acta Crystallogr. D Biol. Crystallogr. 2013, 69 (7), 1204–1214. 10.1107/S0907444913000061.

(46) Raig, N. D.; Surridge, K. J.; Sanz-Murillo, M.; Dederer, V.; Krämer, A.; Schwalm, M. P.; Lattal, N. M.; Elson, L.; Chatterjee, D.; Mathea, S.; Hanke, T.; Leschziner, A. E.; Reck-Peterson, S. L.; Knapp, S. Type II Kinase Inhibitors That Target Parkinson’s Disease–Associated LRRK2. Sci. Adv. 2025, 11 (23), eadt2050. 10.1126/sciadv.adt2050.

(47) Lebedev, A. A.; Vagin, A. A.; Murshudov, G. N. Model Preparation in *MOLREP* and Examples of Model Improvement Using X-Ray Data. Acta Crystallogr. D Biol. Crystallogr. 2008, 64 (1), 33–39. 10.1107/S0907444907049839.

(48) Emsley, P.; Cowtan, K. *Coot* : Model-Building Tools for Molecular Graphics. Acta Crystallogr. D Biol. Crystallogr. 2004, 60 (12), 2126–2132. 10.1107/S0907444904019158.

(49) Vagin, A. A.; Steiner, R. A.; Lebedev, A. A.; Potterton, L.; McNicholas, S.; Long, F.; Murshudov, G. N. *REFMAC* 5 Dictionary: Organization of Prior Chemical Knowledge and Guidelines for Its Use. Acta Crystallogr. D Biol. Crystallogr. 2004, 60 (12), 2184–2195. 10.1107/S0907444904023510.

(50) Fulmer, G. R.; Miller, A. J. M.; Sherden, N. H.; Gottlieb, H. E.; Nudelman, A.; Stoltz, B. M.; Bercaw, J. E.; Goldberg, K. I. NMR Chemical Shifts of Trace Impurities: Common Laboratory Solvents, Organics, and Gases in Deuterated Solvents Relevant to the Organometallic Chemist. Organometallics 2010, 29 (9), 2176–2179. 10.1021/om100106e.

(51) Miller, W. H.; Pendrak, I.; Seefeld, M. A. ANTIBACTERIAL AGENTS. WO 2006/002047A2, January 5, 2006.

(52) Wortmann, L.; Sautier, B.; Eis, K.; Briem, H.; Böhnke, N.; Von Nussbaum, F.; Hillig, R.; Bader, B.; Schröder, J.; Petersen, K.; Lienau, P.; Wengner, A. M.; Moosmayer, D.; Wang, Q.; Schick, H. 2-METHYL-QUINAZOLINES. WO2018172250A1, September 27, 2018.

(53) Zaile Zhuang, Riley K. Togashi, Patrick Kearney, Ian Pass, Steven M. Swick, Fu-Yue Zeng, Andrey A. Bobkov, Lynn M. Fujimoto, Shubhankar Dutta, Athina Zerva, Nicolai D. Raig, Debasmita Saha, Atoosa Emami, Martin P. Schwalm, Bradley K. Moon, Samuel T. Howard, Stefan Knapp, Thomas Hanke, Thomas D.Y. Chung, James K. Chen. Small-molecule modulators of HIPK4 activity and proteostasis. bioRxiv 2026.05.12.724395; doi: 10.64898/2026.05.12.724395

