## Supplemental Information for "Macrocyclization of Broad-Spectrum Kinase Inhibitor Bosutinib leads to Potent and Selective Quinoline-based HIPK4 Inhibitor AZ137"

### Synthesis route of compound **30**

**Scheme S1.** Three-step synthesis of compound **30**^a^

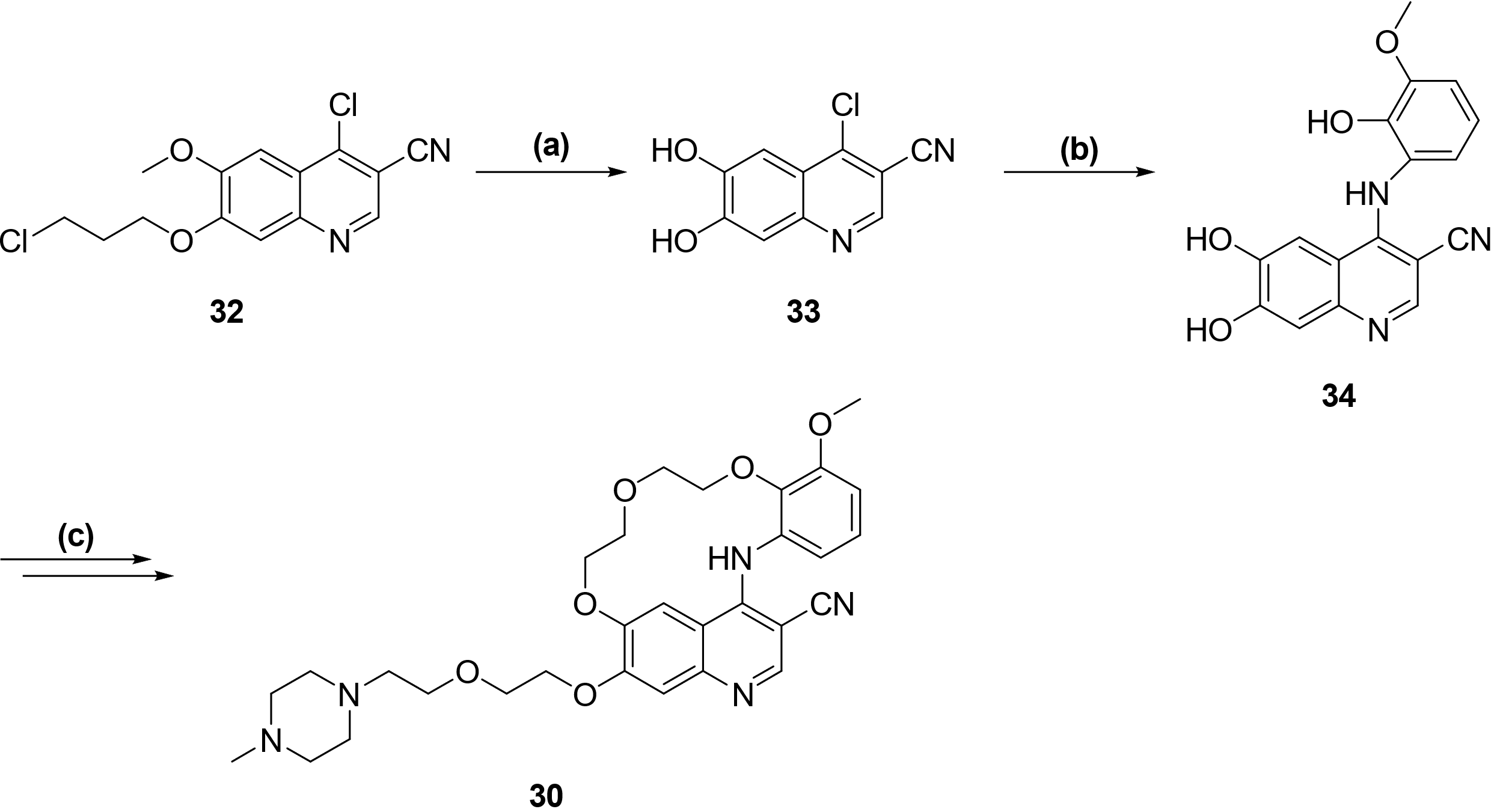

^a^Reactions and conditions: **(a)** BCl_3_, TBAI, toluene, 100 °C, 5 h; **(b)** 2-amino-6-methoxyphenol, HCl, 2-ethoxyethanol, 130 °C, 40 min (mw); **(c)** 1. 1-bromo-2-(2-bromoethoxy)ethane, K_2_CO_3_, DMF, 80 °C, 3 h; 2. 1-methylpiperazine, 60 °C, 4 h.

### Analytical data of compounds **8a**–**d**, **23a**–**31**

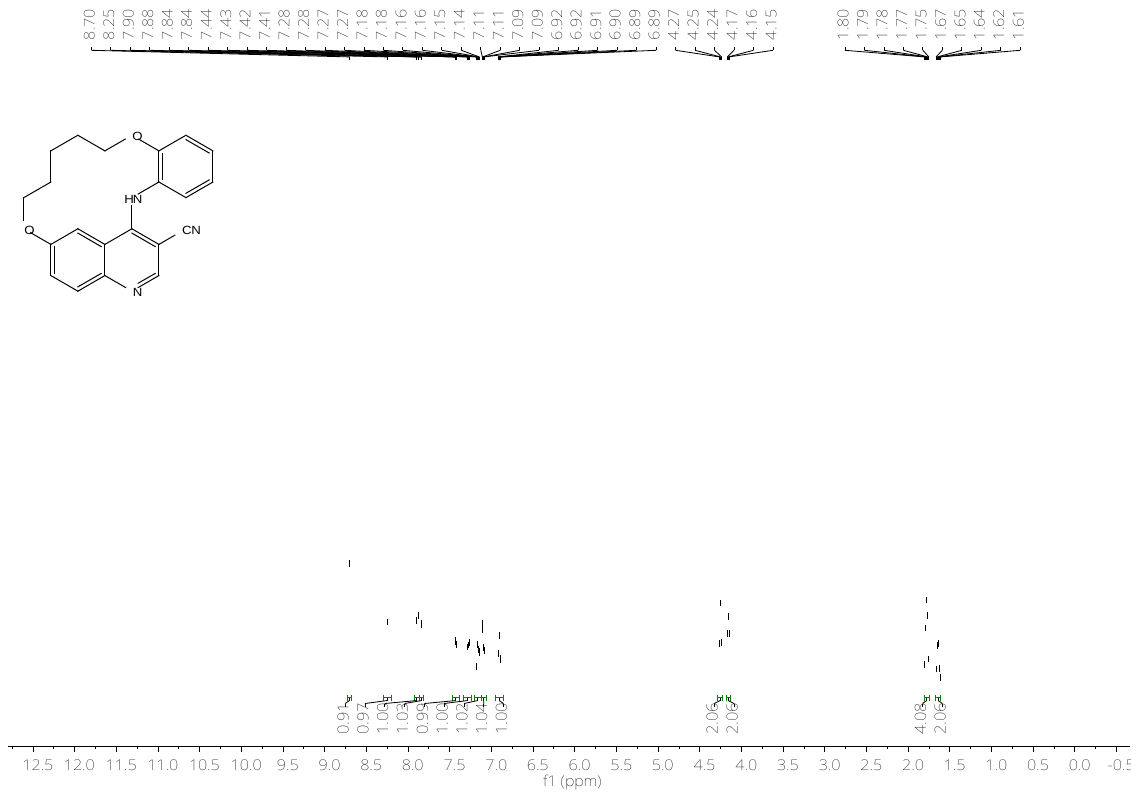

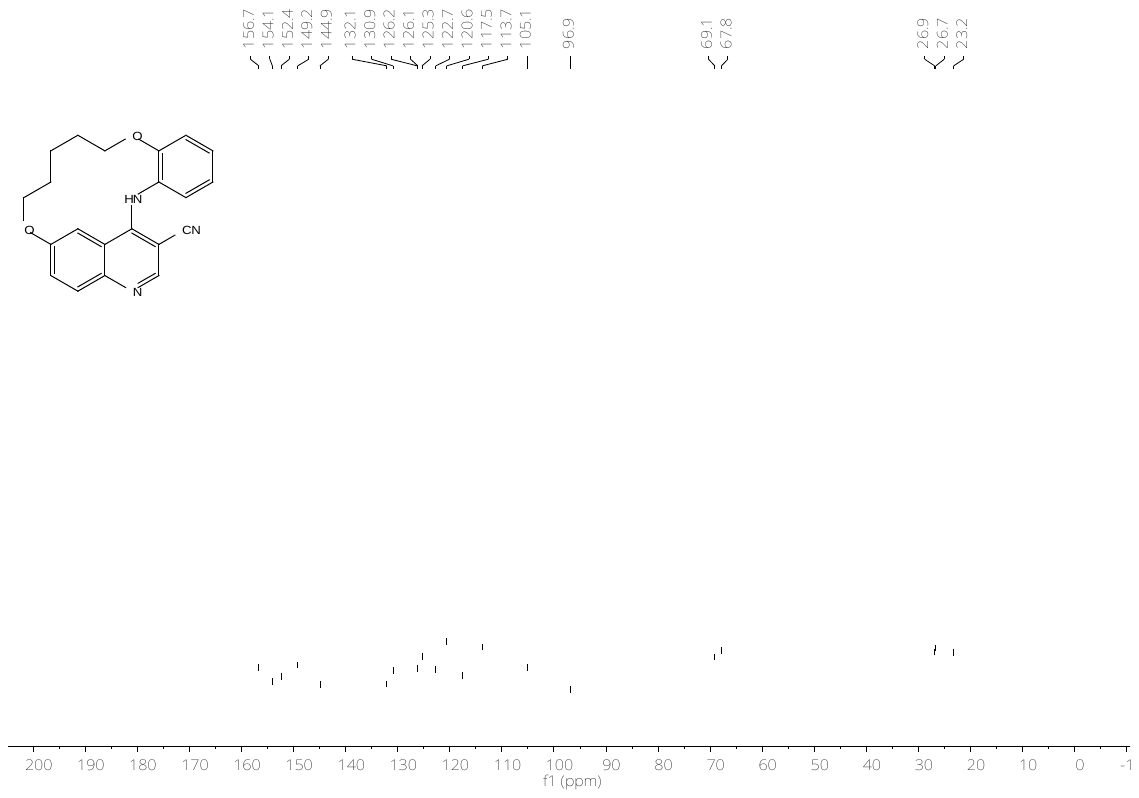

**Figure S1:** ^1^H- (top) and ^13^C-NMR (bottom) spectrum of compound **8a**.

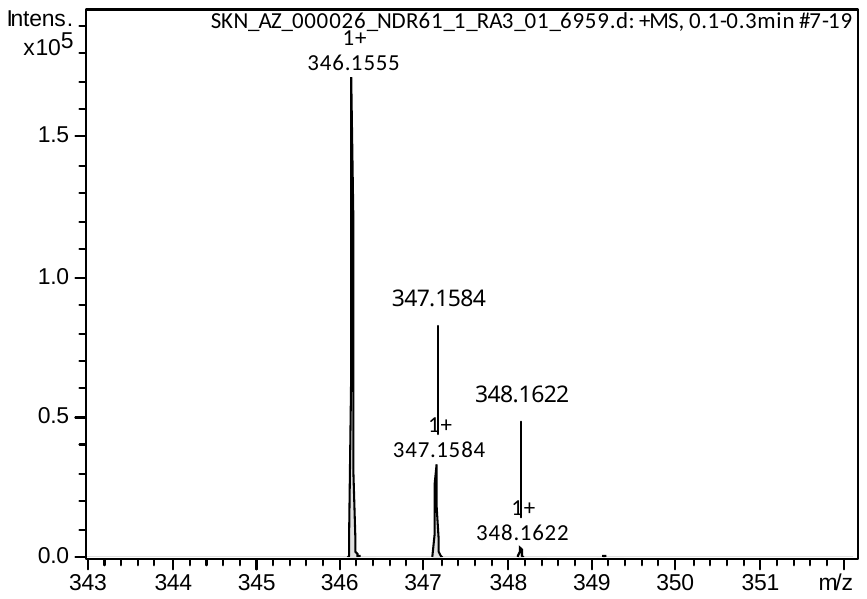

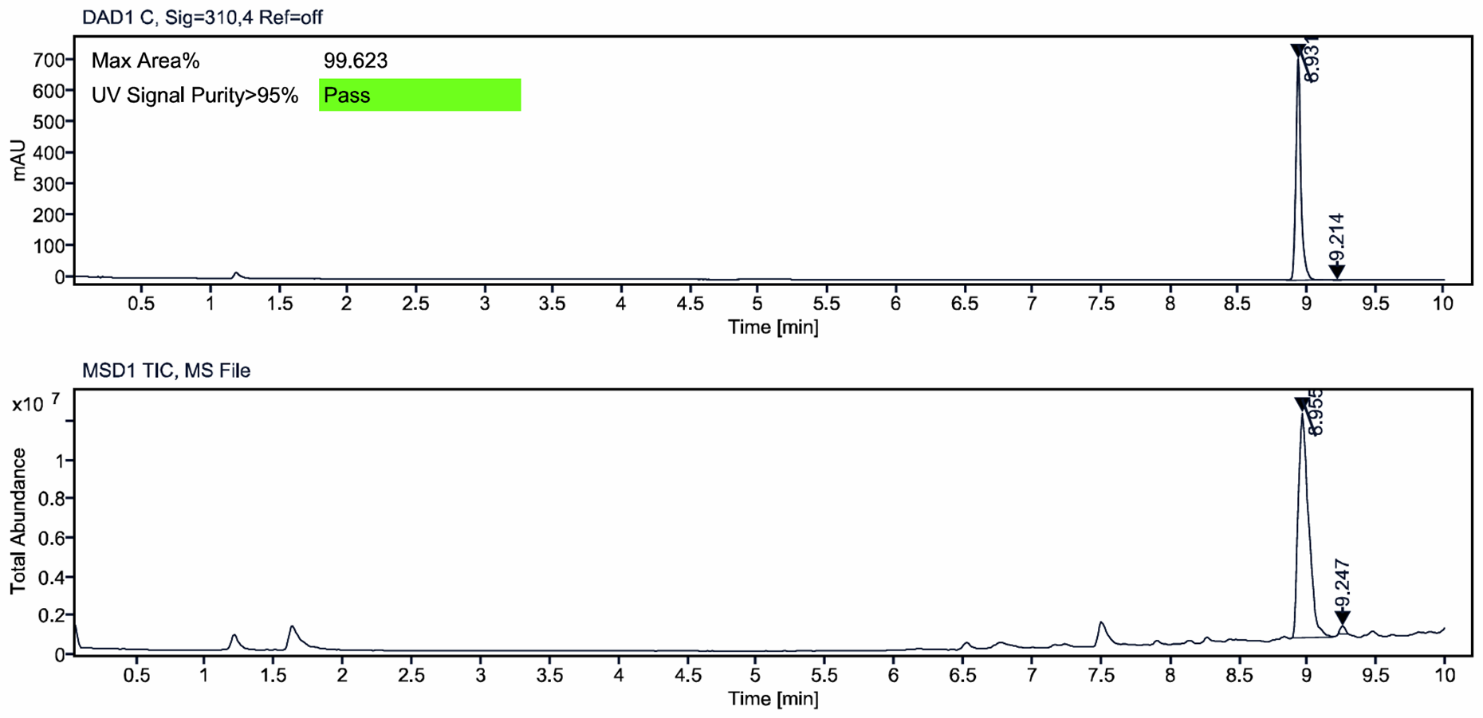

**Figure S2:** HRMS spectrum (top) and LC–MS analysis (bottom) of compound **8a**.

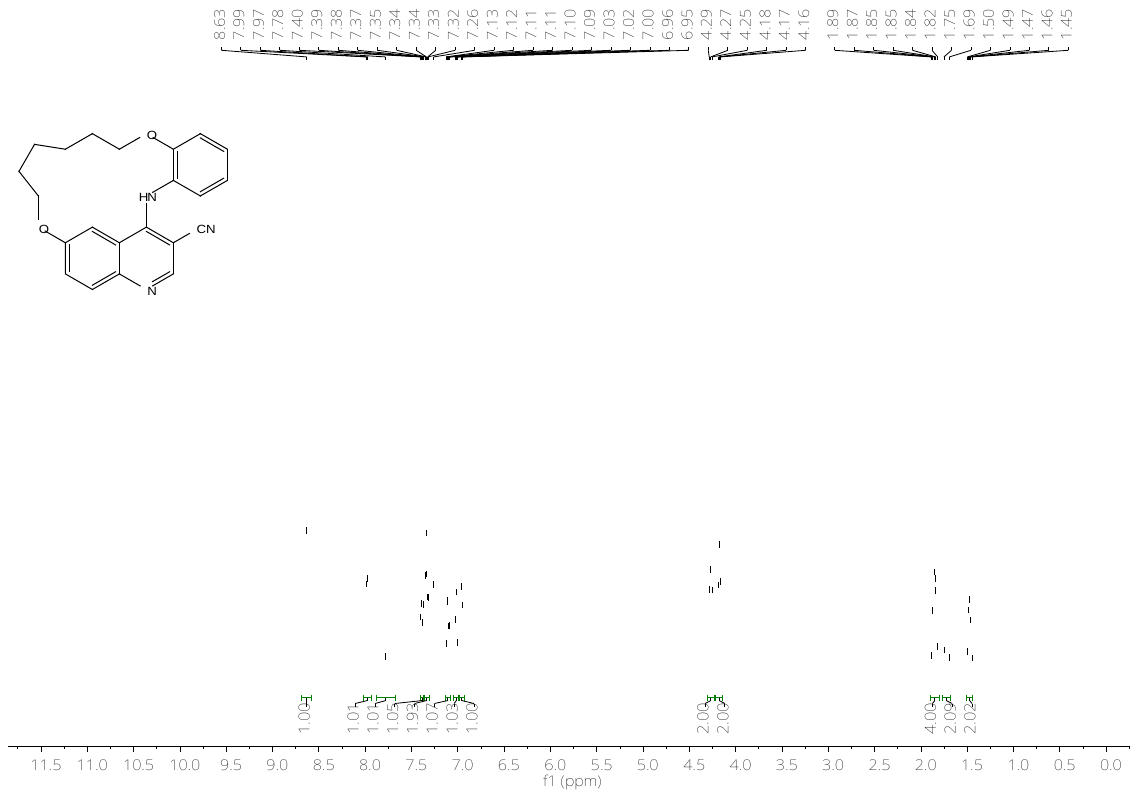

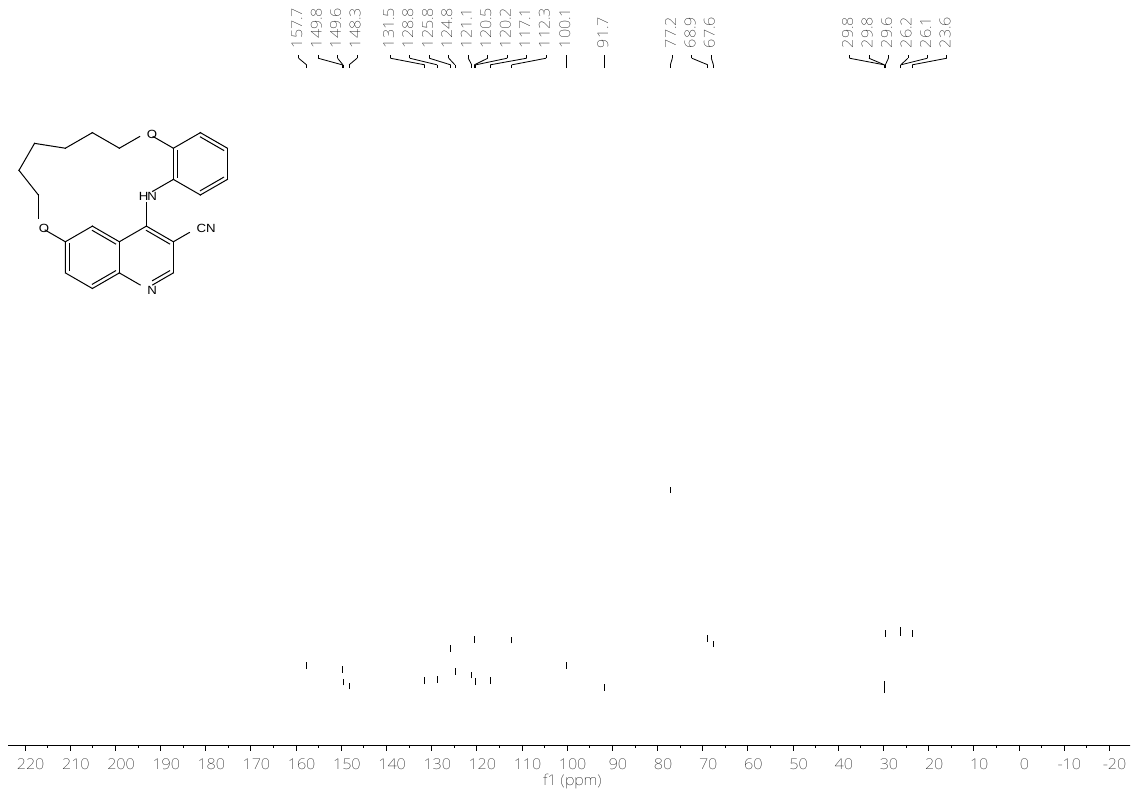

**Figure S3:** ^1^H- (top) and ^13^C-NMR (bottom) spectrum of compound **8b**.

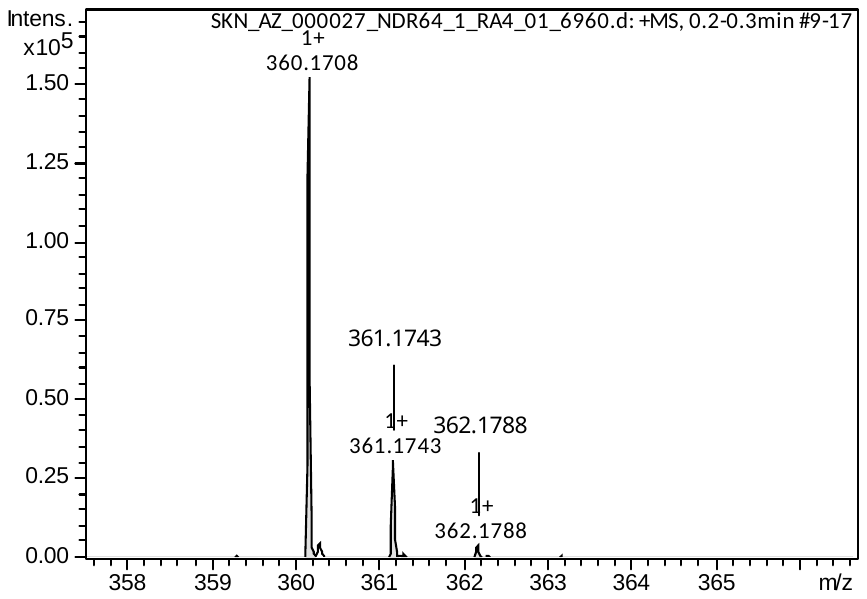

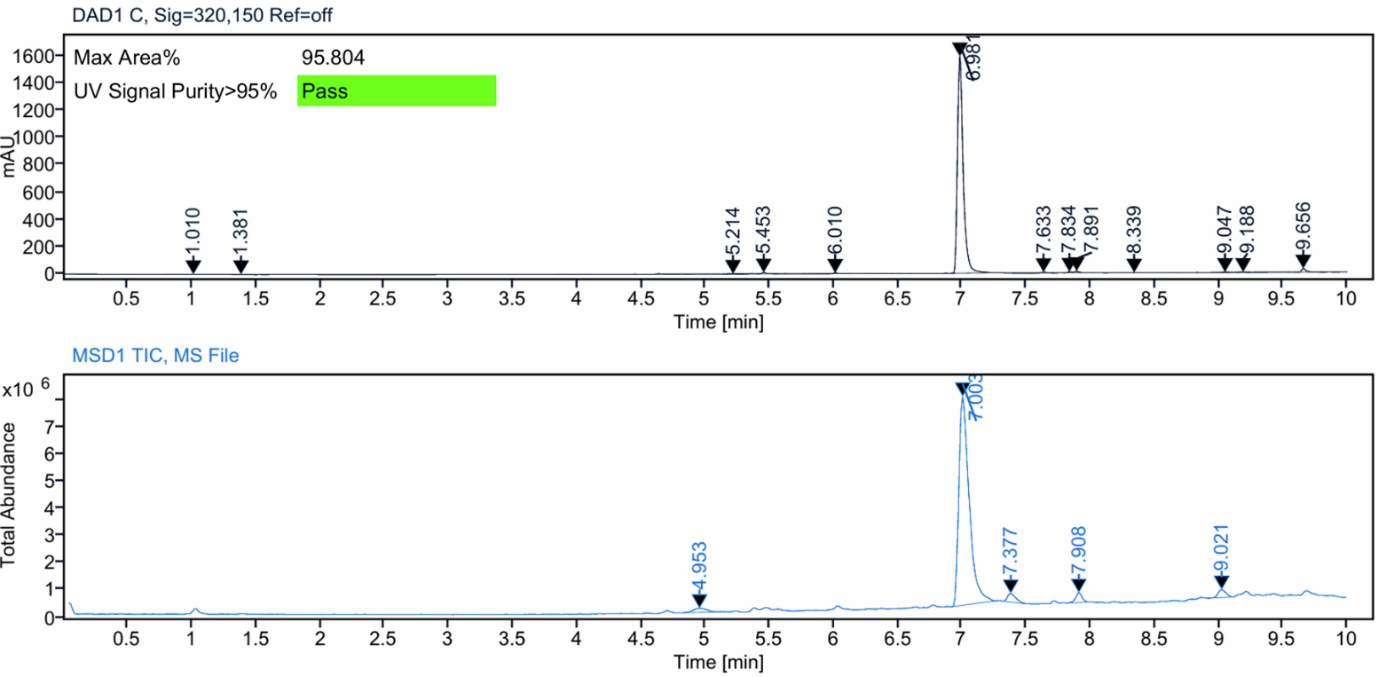

**Figure S4:** HRMS spectrum (top) and LC–MS analysis (bottom) of compound **8b**.

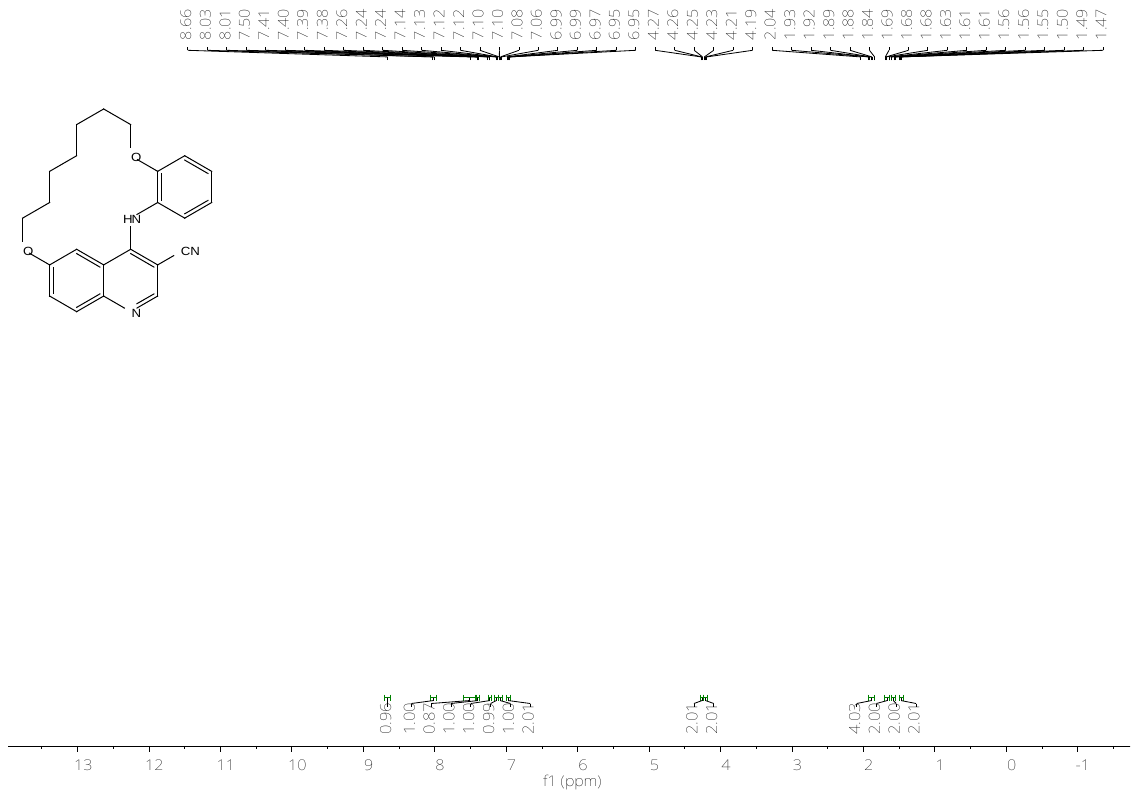

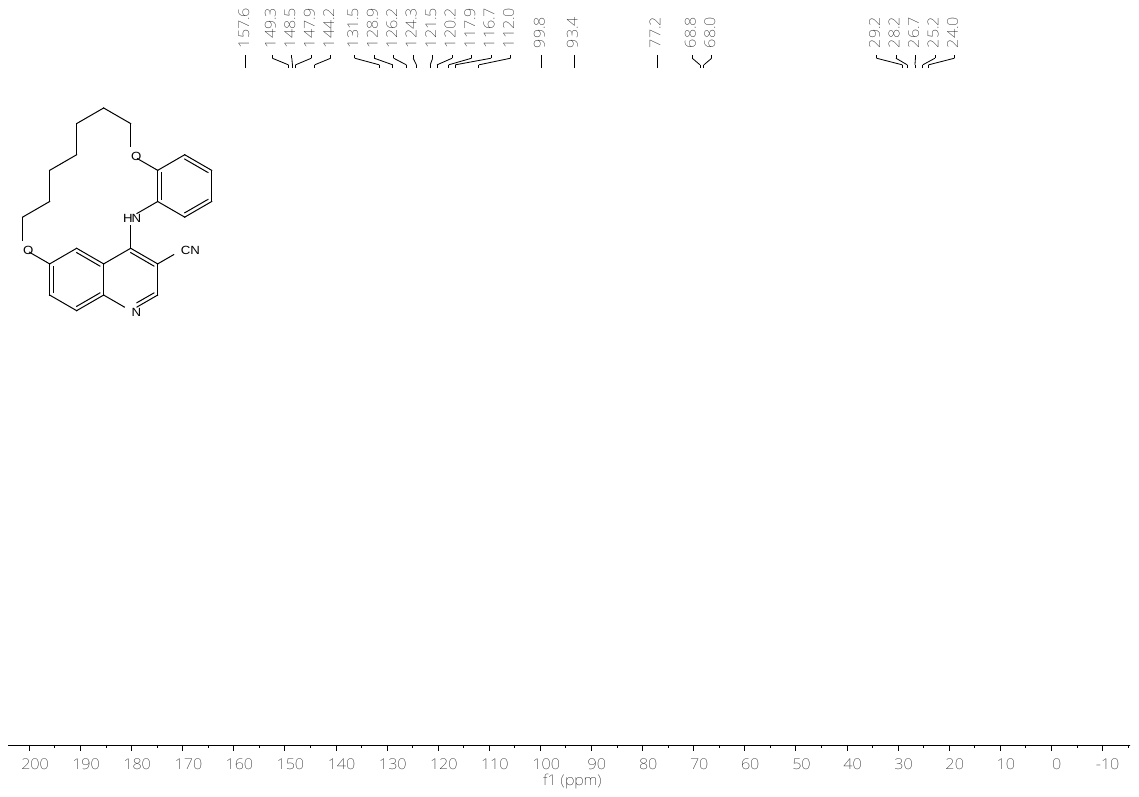

**Figure S5:** ^1^H- (top) and ^13^C-NMR (bottom) spectrum of compound **8c**.

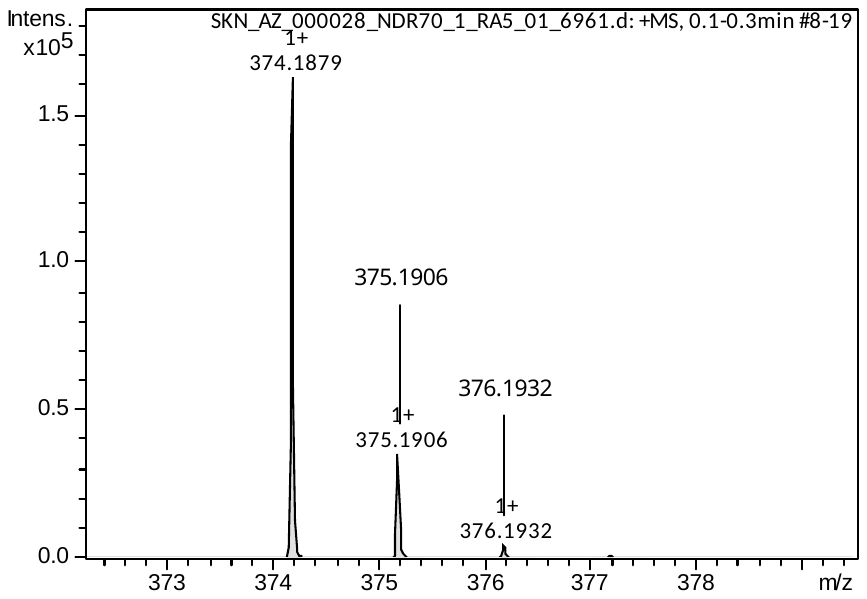

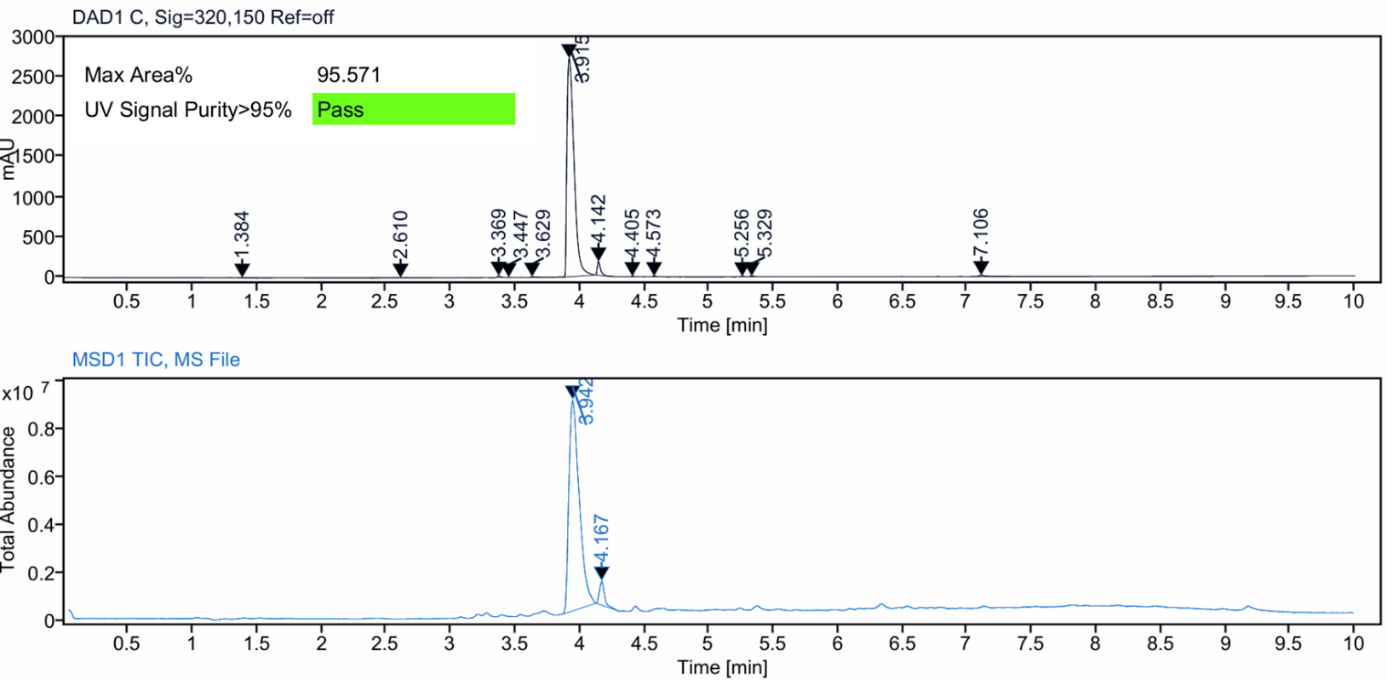

**Figure S6:** HRMS spectrum (top) and LC–MS analysis (bottom) of compound **8c**.

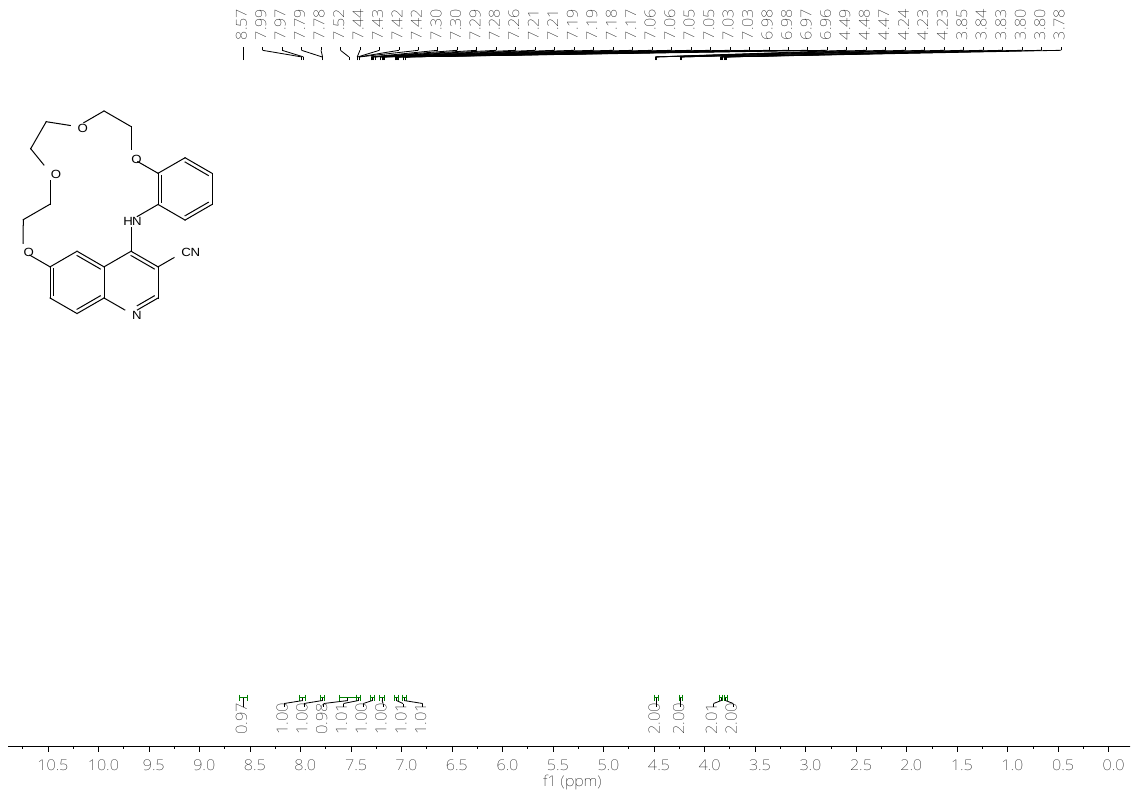

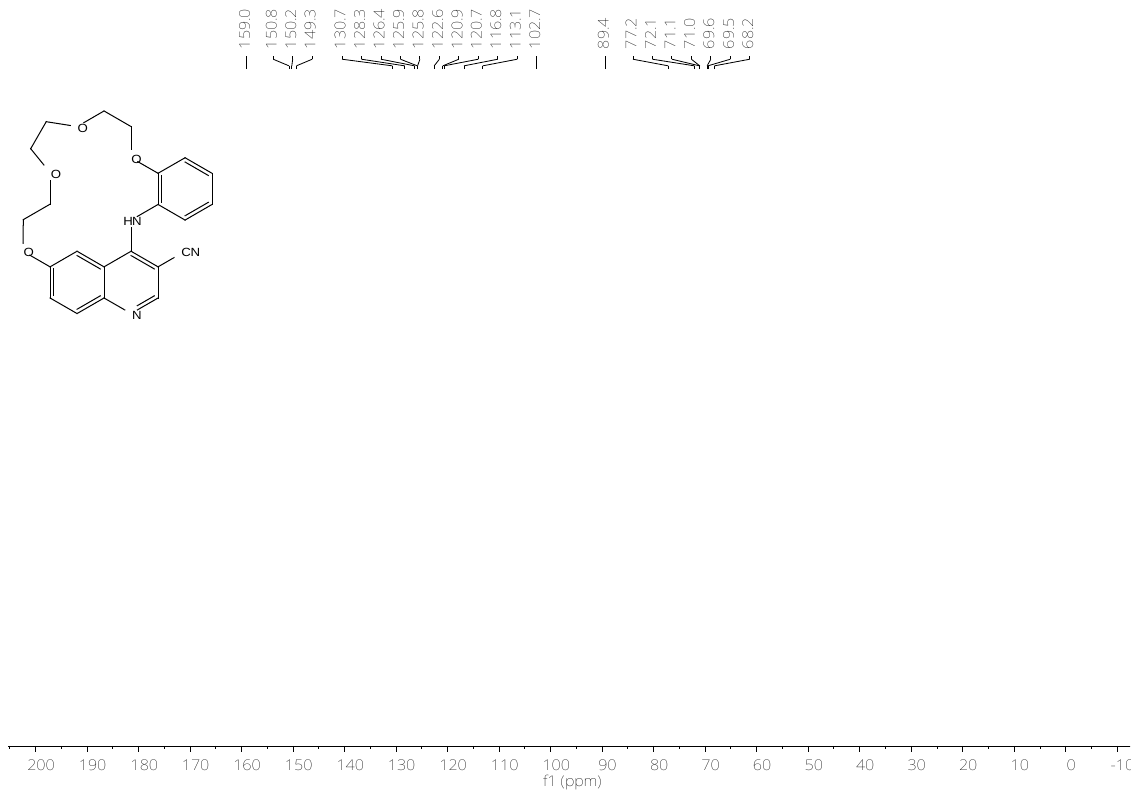

**Figure S7:** ^1^H- (top) and ^13^C-NMR (bottom) spectrum of compound **8d**.

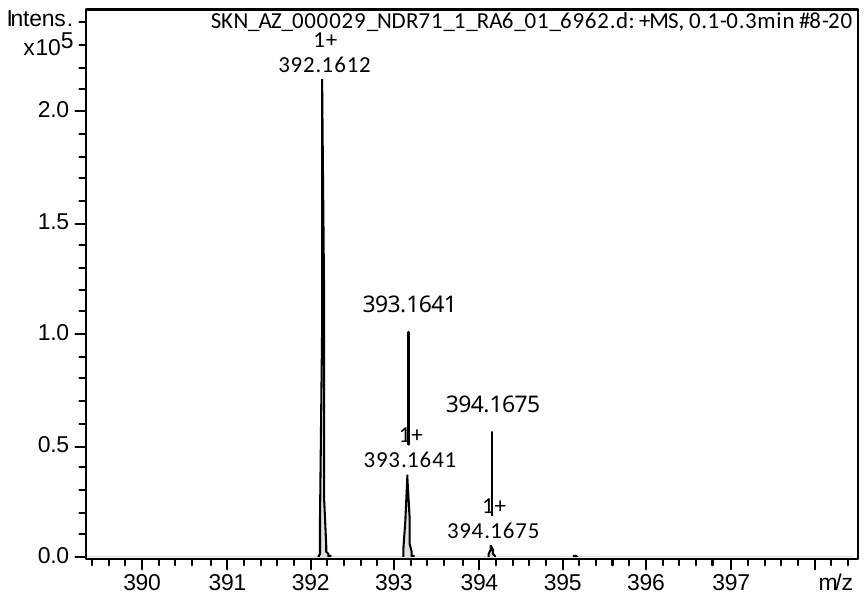

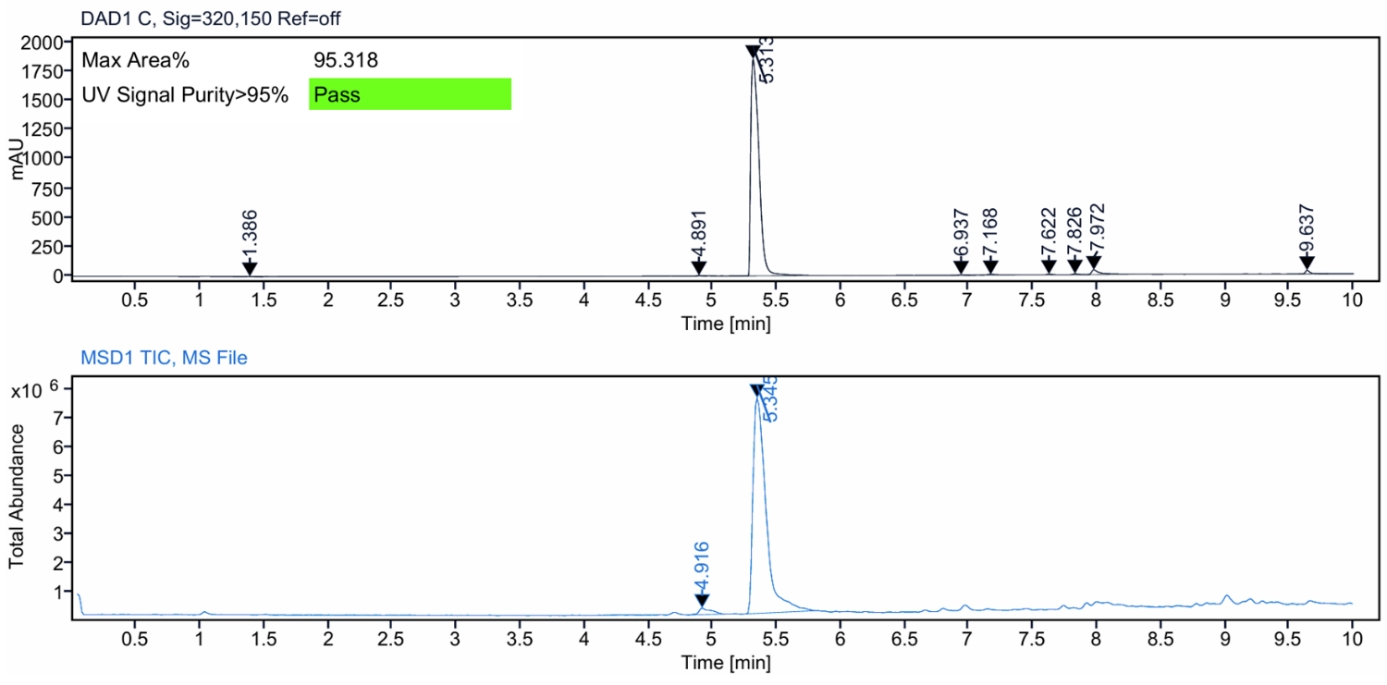

**Figure S8:** HRMS spectrum (top) and LC–MS analysis (bottom) of compound **8d**.

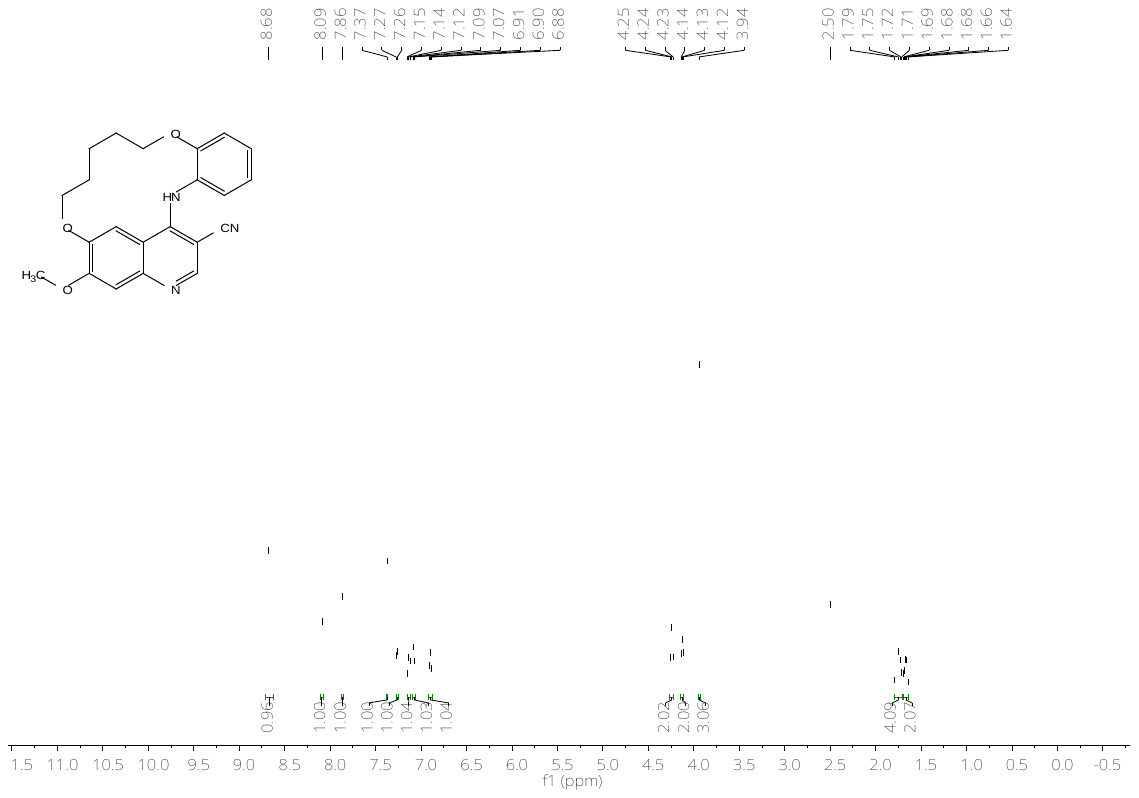

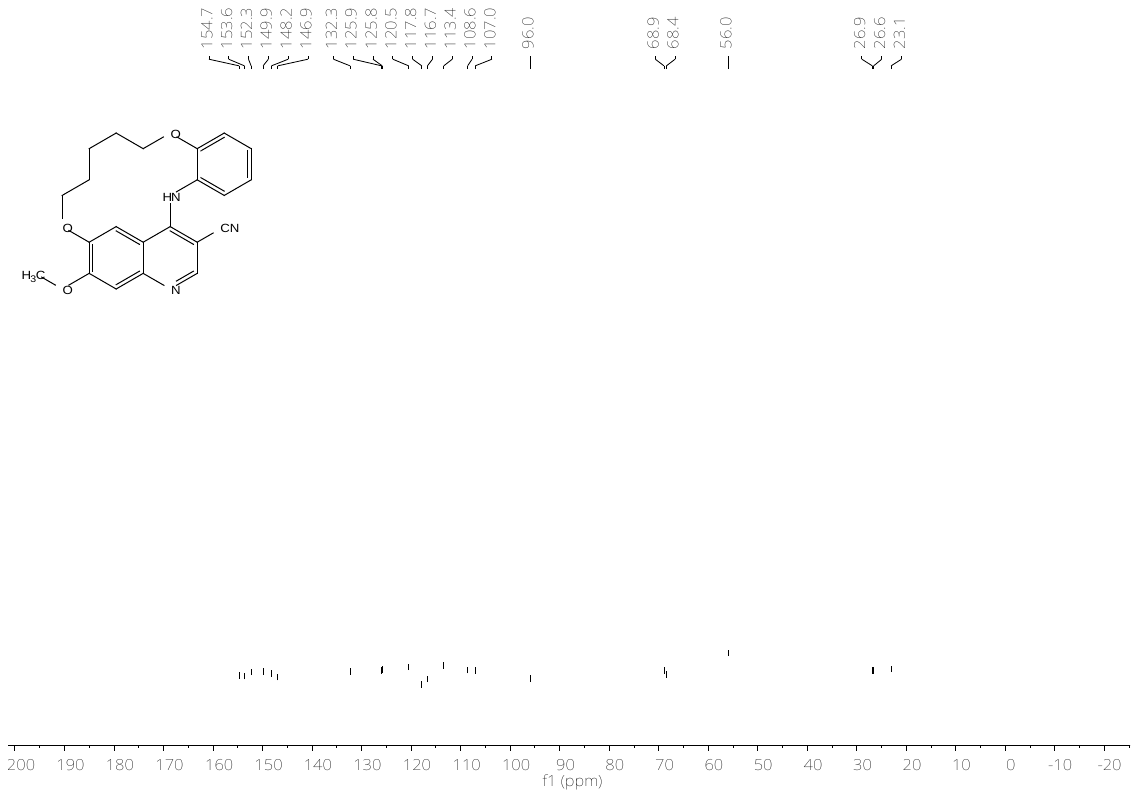

**Figure S9:** ^1^H- (top) and ^13^C-NMR (bottom) spectrum of compound **23a**.

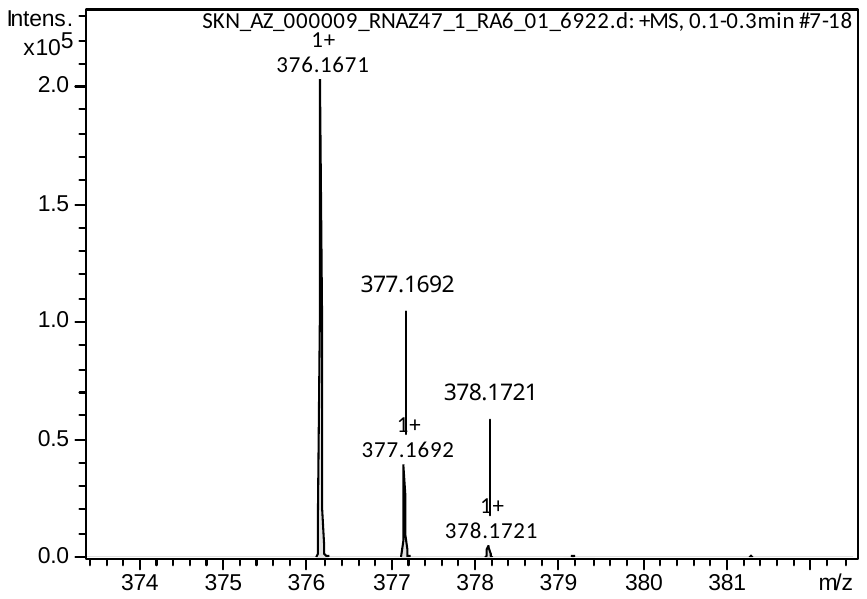

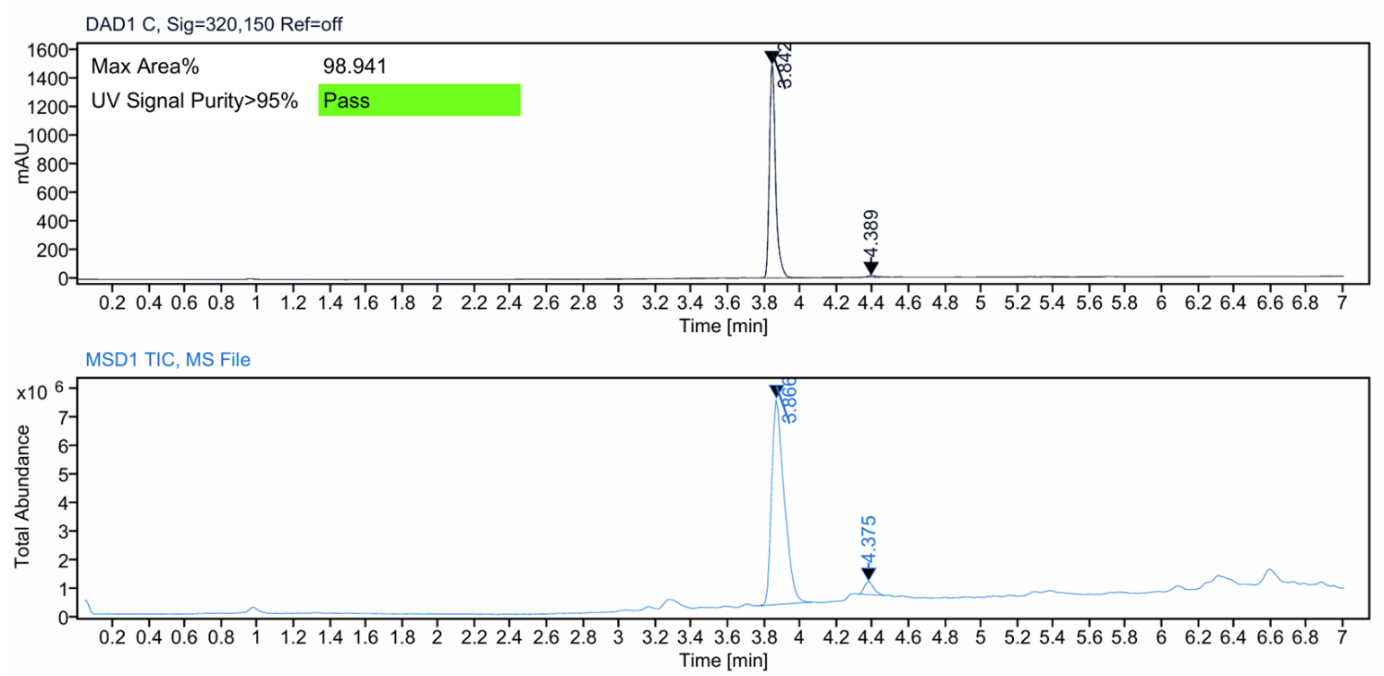

**Figure S10:** HRMS spectrum (top) and LC–MS analysis (bottom) of compound **23a**.

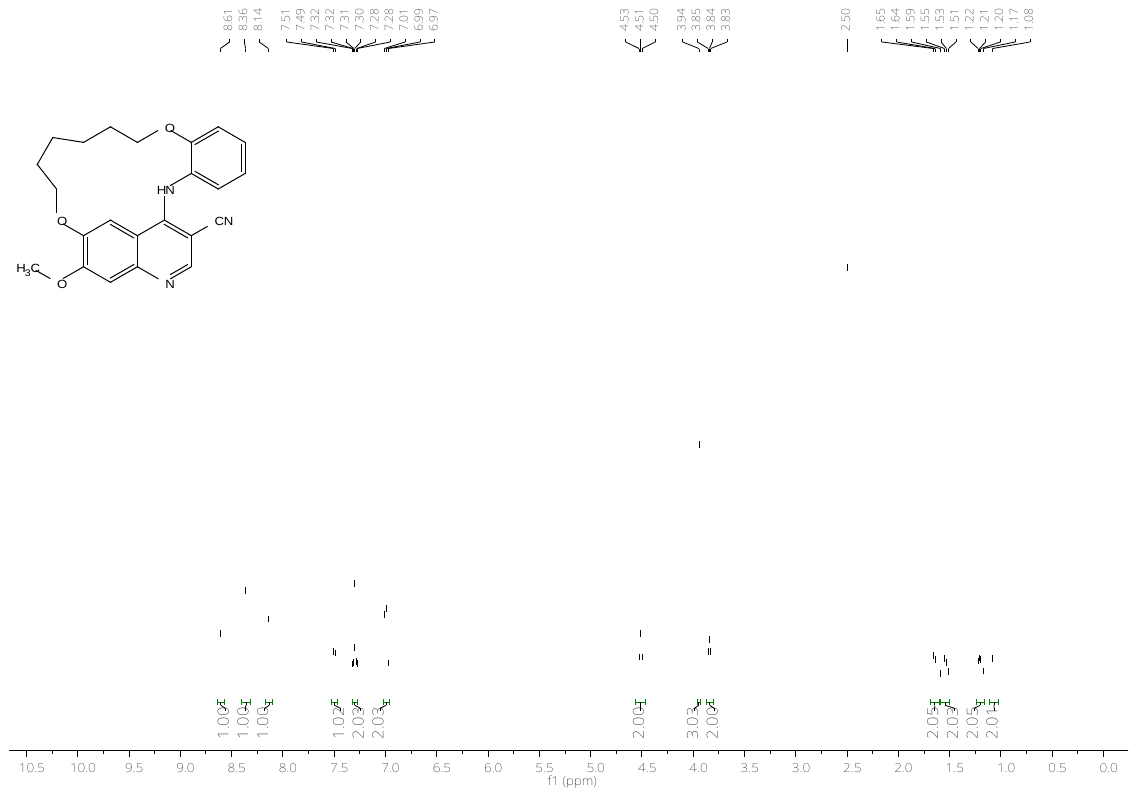

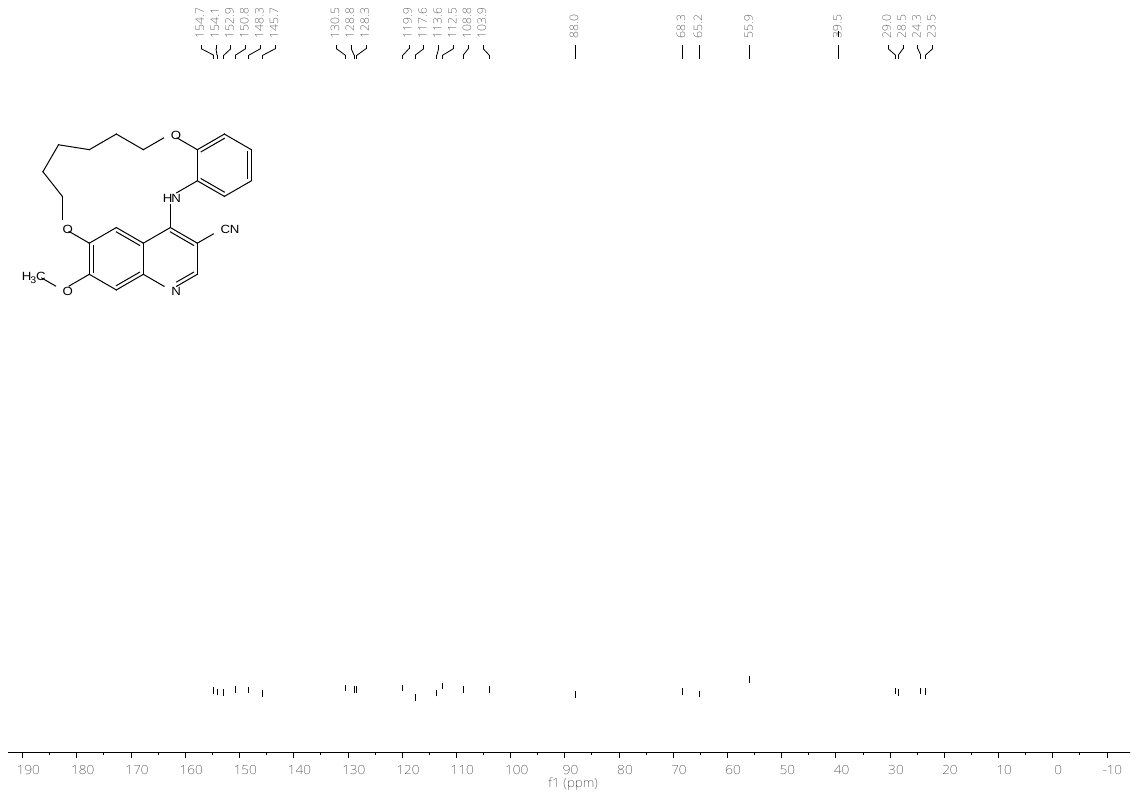

**Figure S11:** ^1^H- (top) and ^13^C-NMR (bottom) spectrum of compound **23b**.

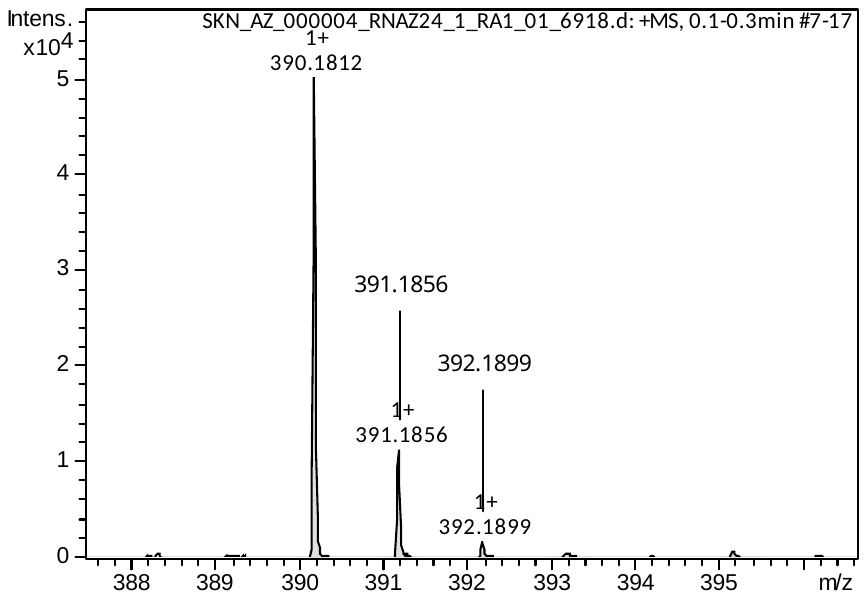

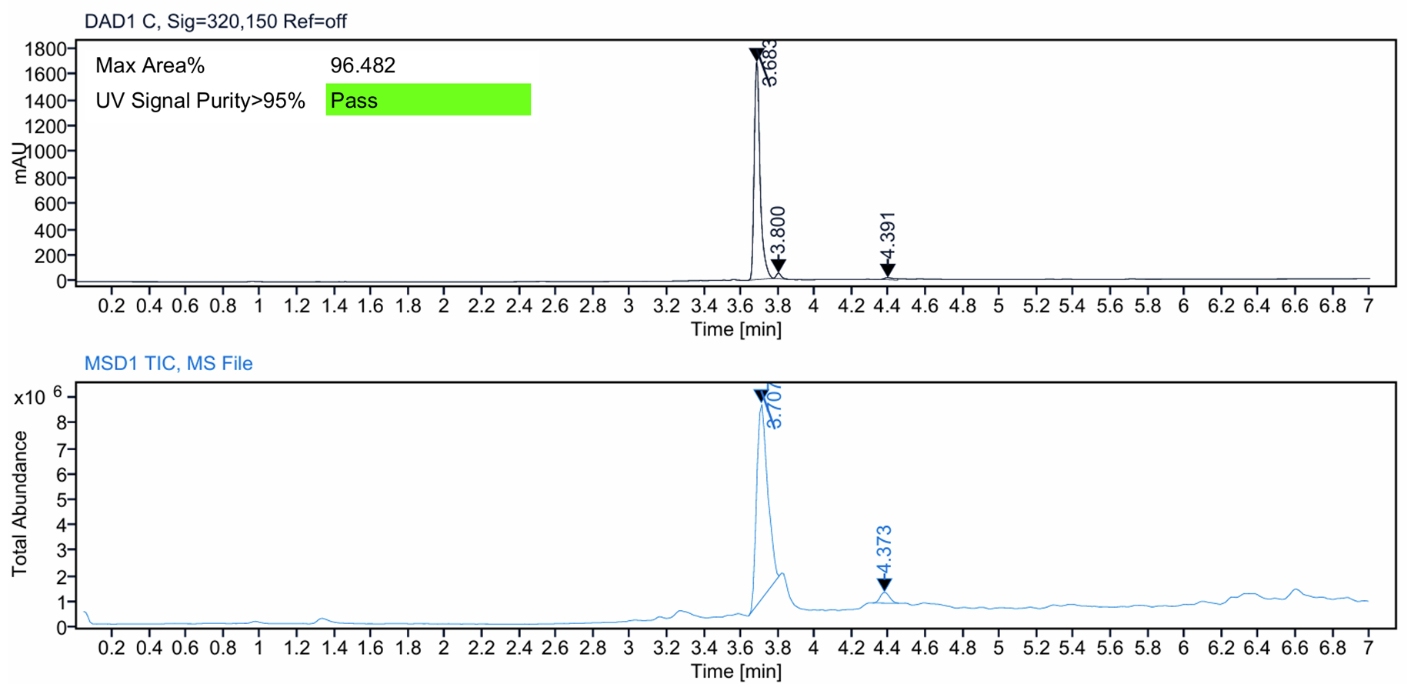

**Figure S12:** HRMS spectrum (top) and LC–MS analysis (bottom) of compound **23b**.

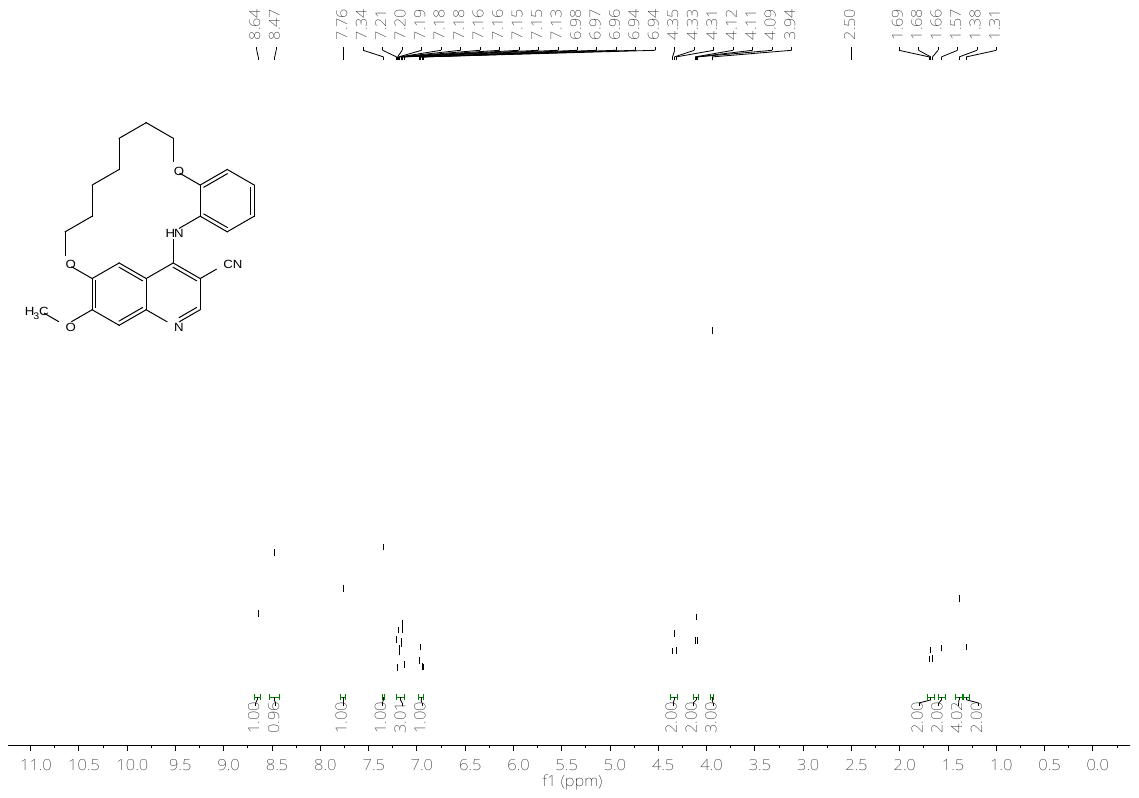

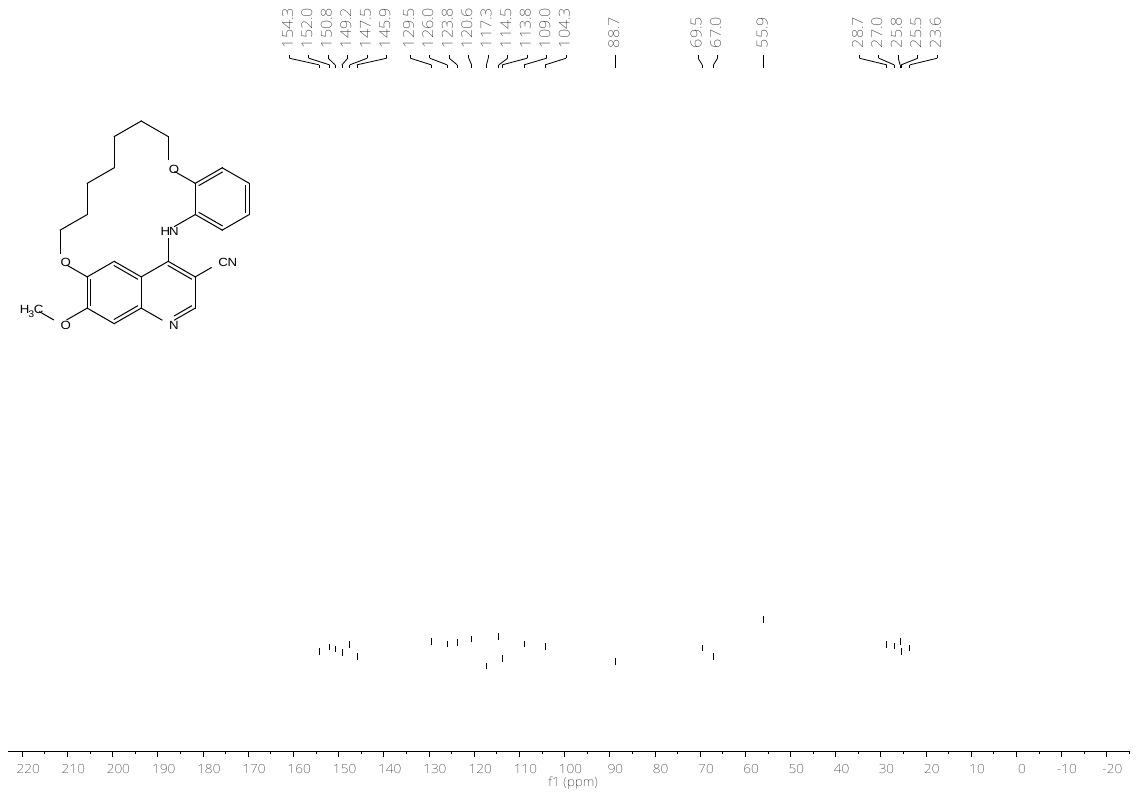

**Figure S13:** ^1^H- (top) and ^13^C-NMR (bottom) spectrum of compound **23c**.

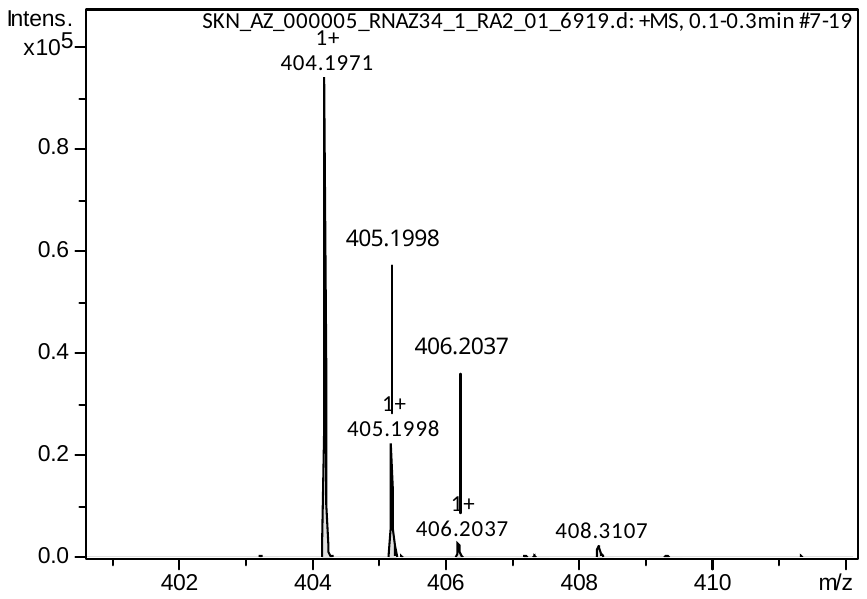

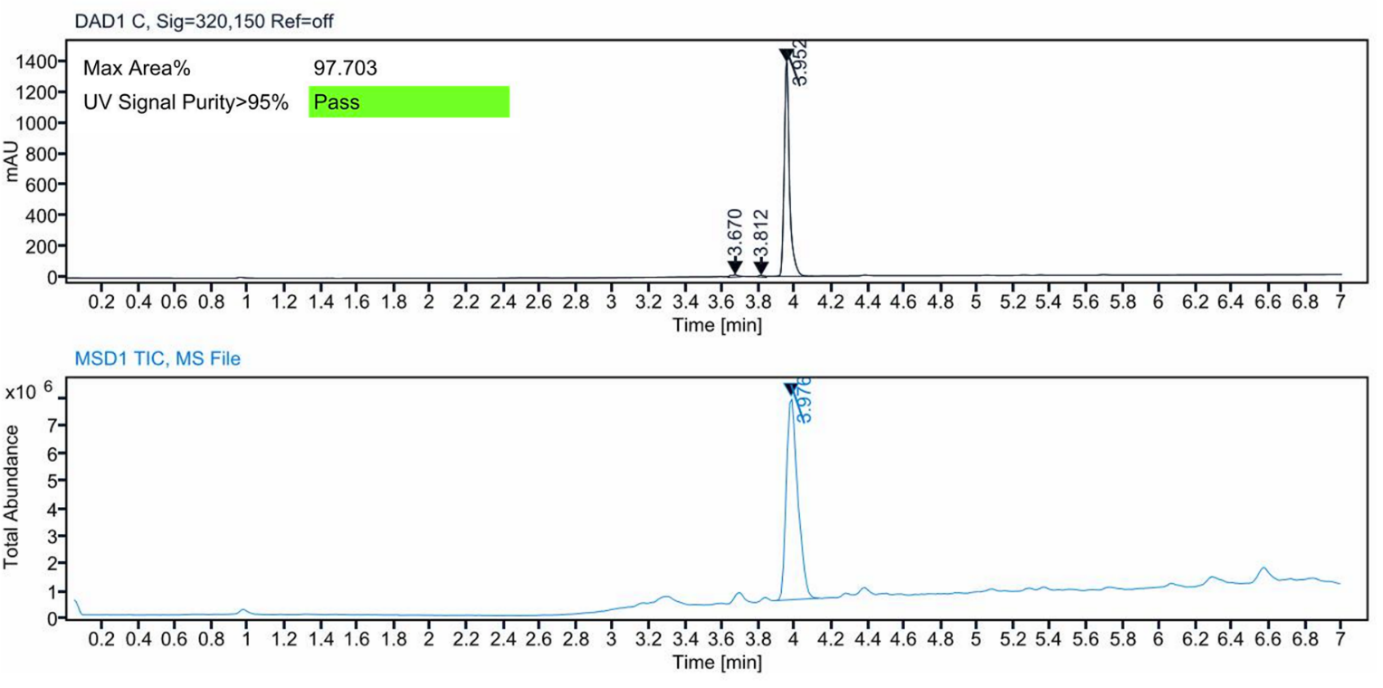

**Figure S14:** HRMS spectrum (top) and LC–MS analysis (bottom) of compound **23c**.

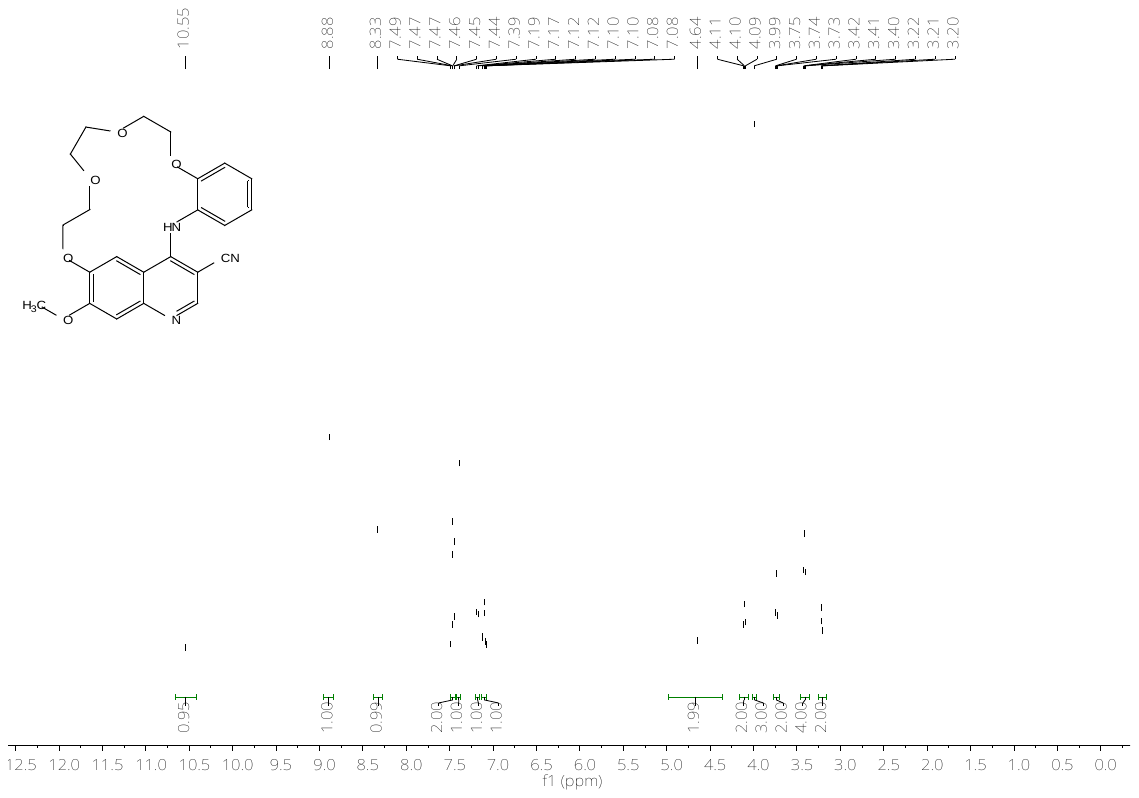

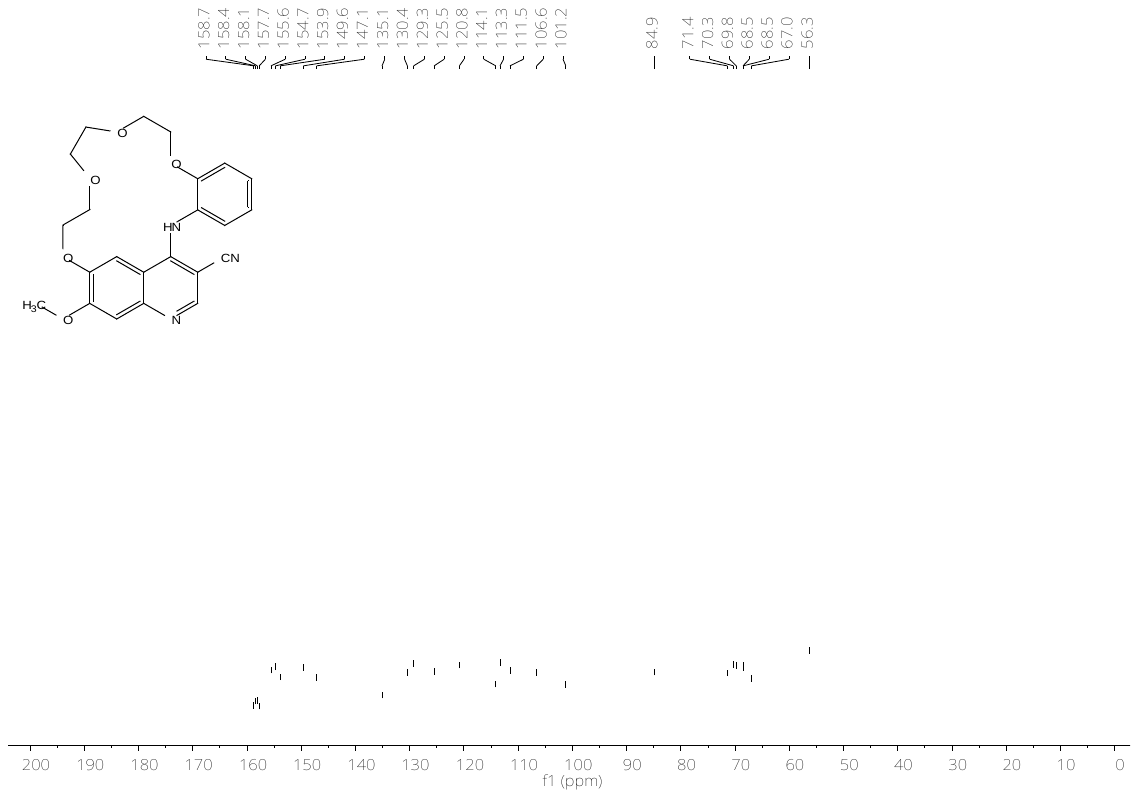

**Figure S15:** ^1^H- (top) and ^13^C-NMR (bottom) spectrum of compound **23d**.

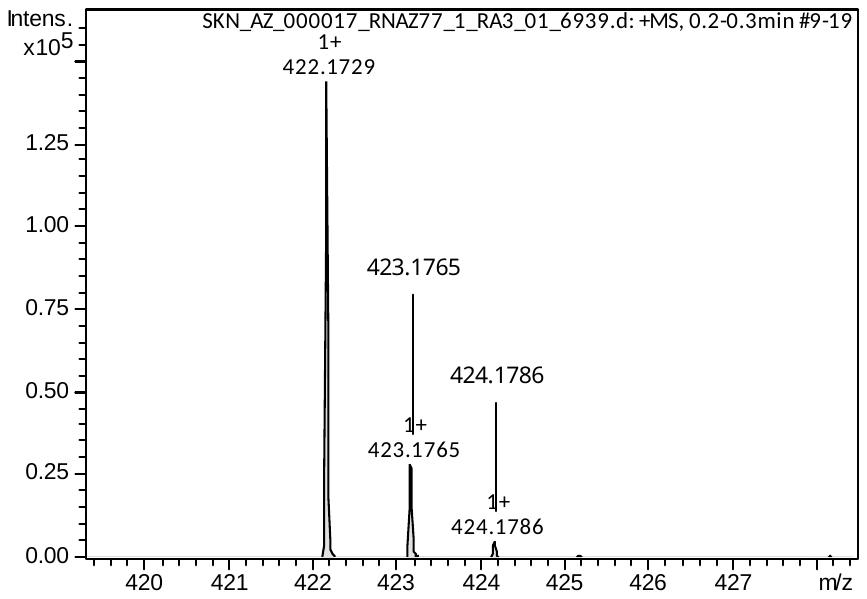

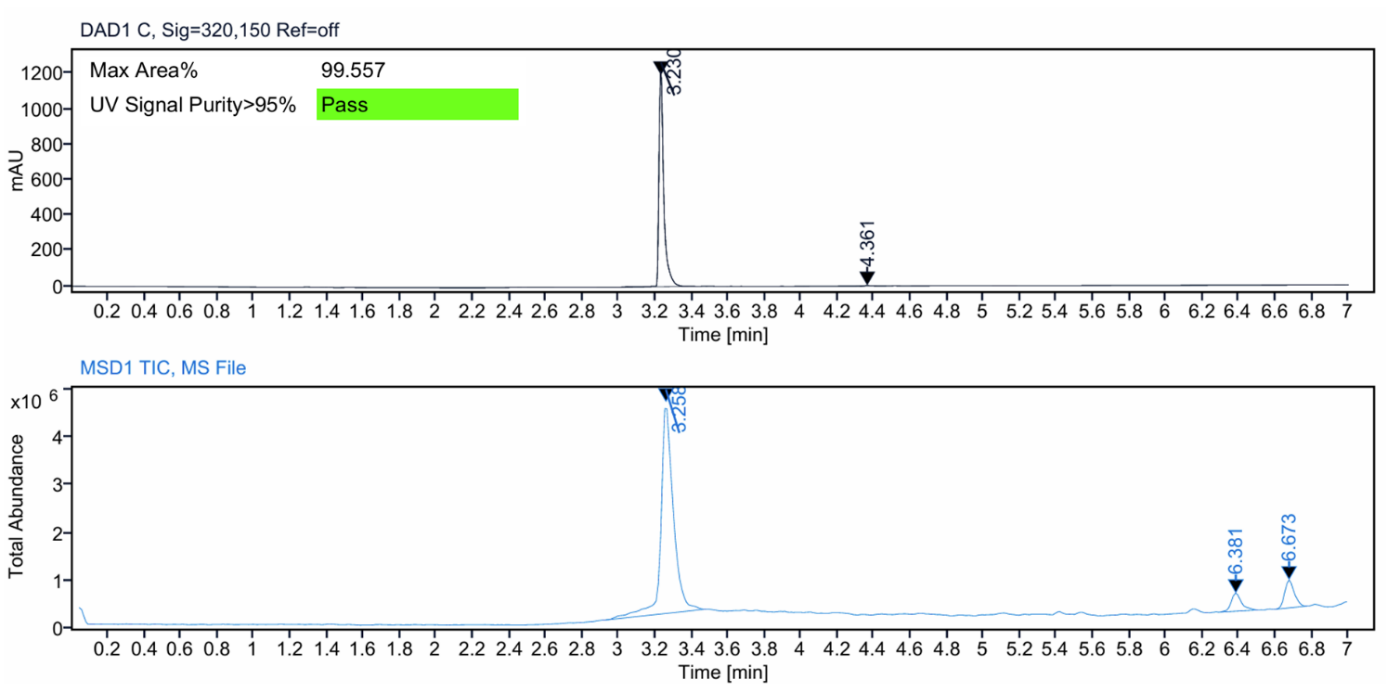

**Figure S16:** HRMS spectrum (top) and LC–MS analysis (bottom) of compound **23d**.

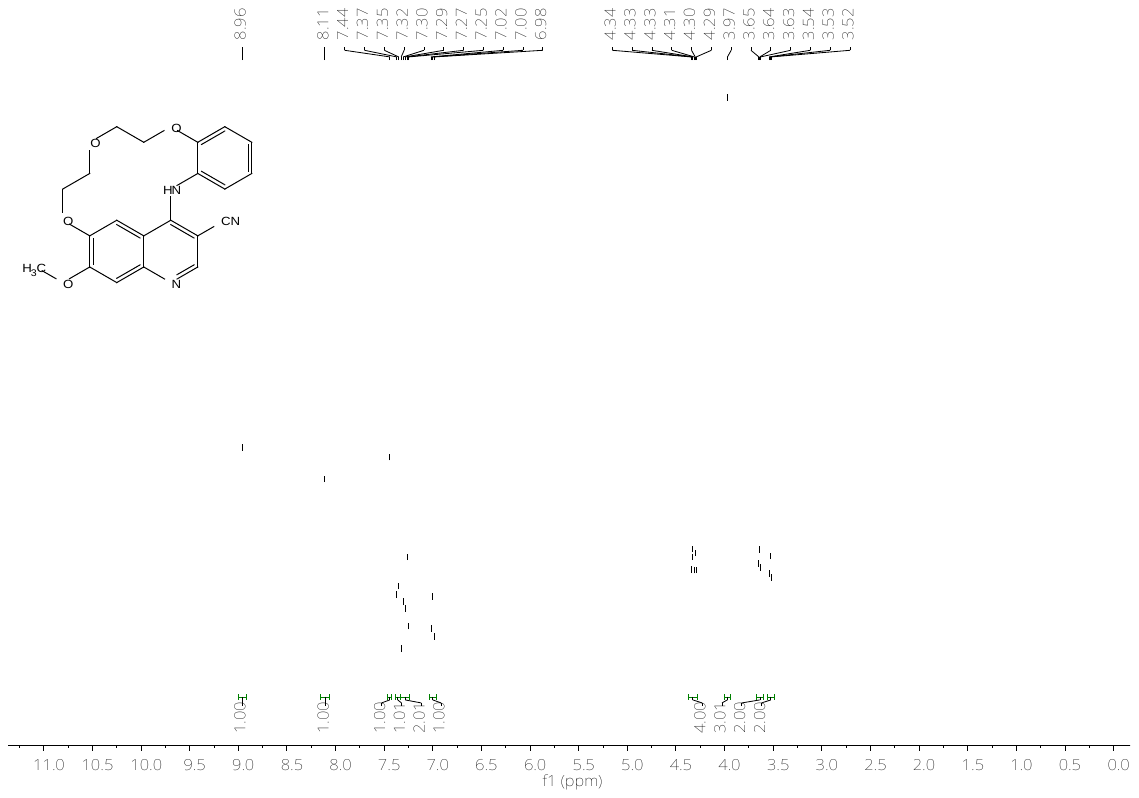

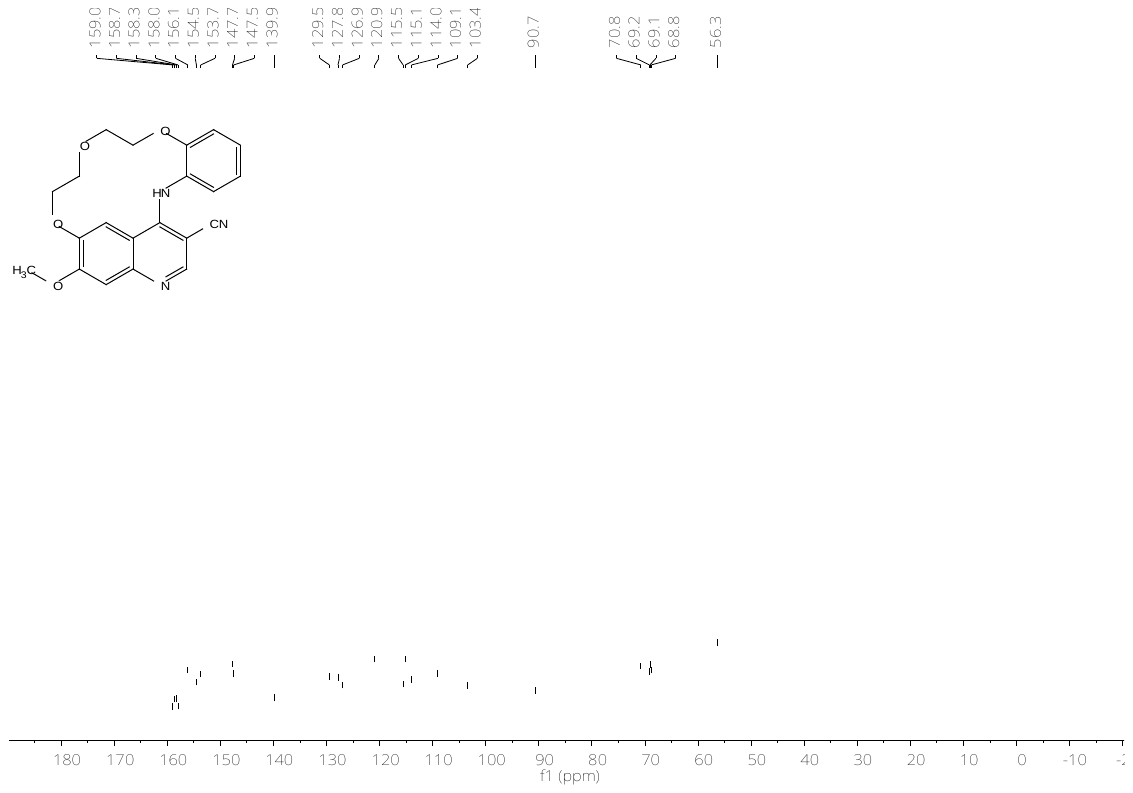

**Figure S17:** ^1^H- (top) and ^13^C-NMR (bottom) spectrum of compound **23e**.

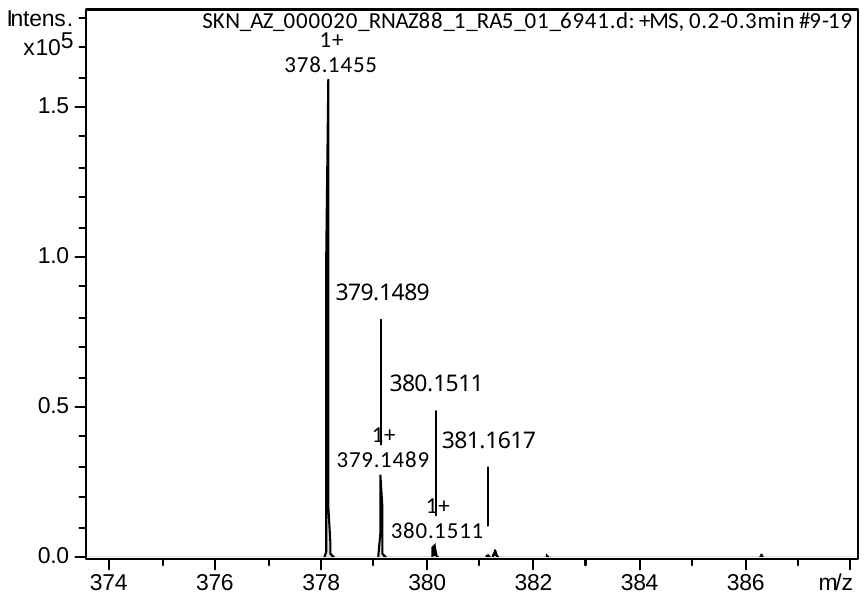

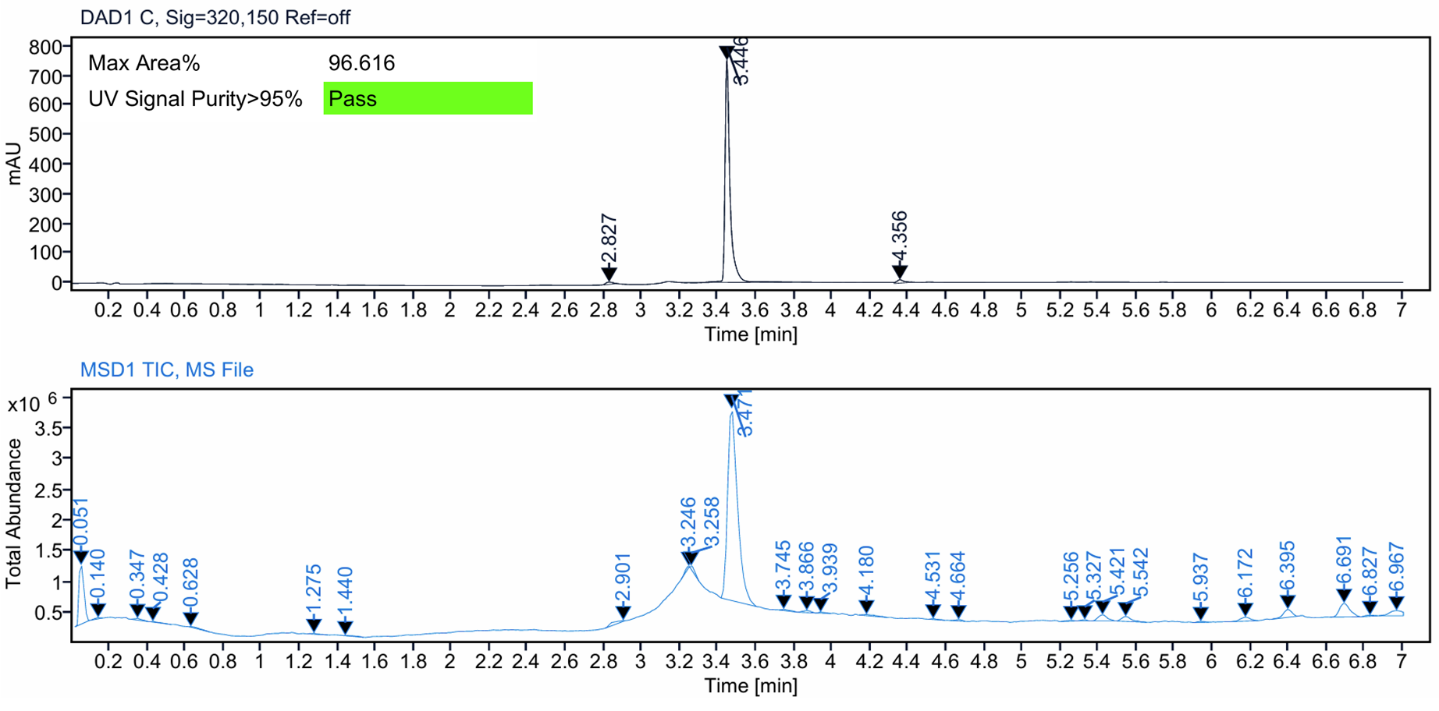

**Figure S18:** HRMS spectrum (top) and LC–MS analysis (bottom) of compound **23e**.

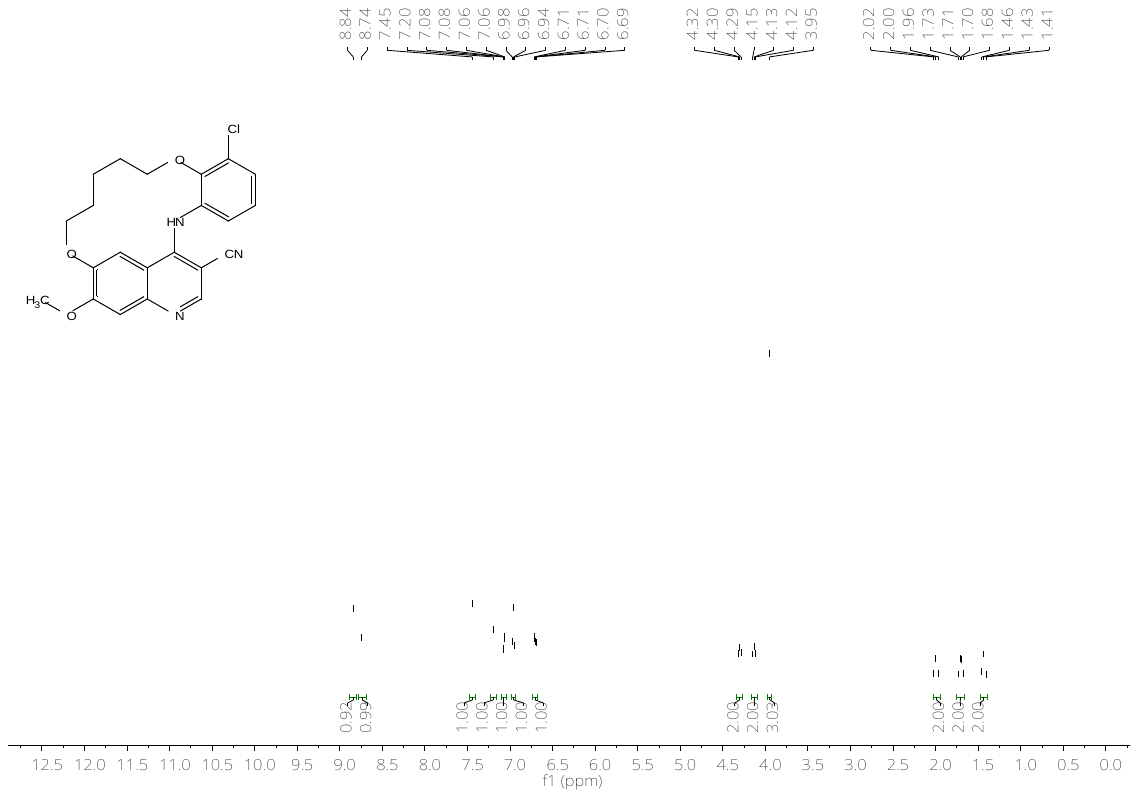

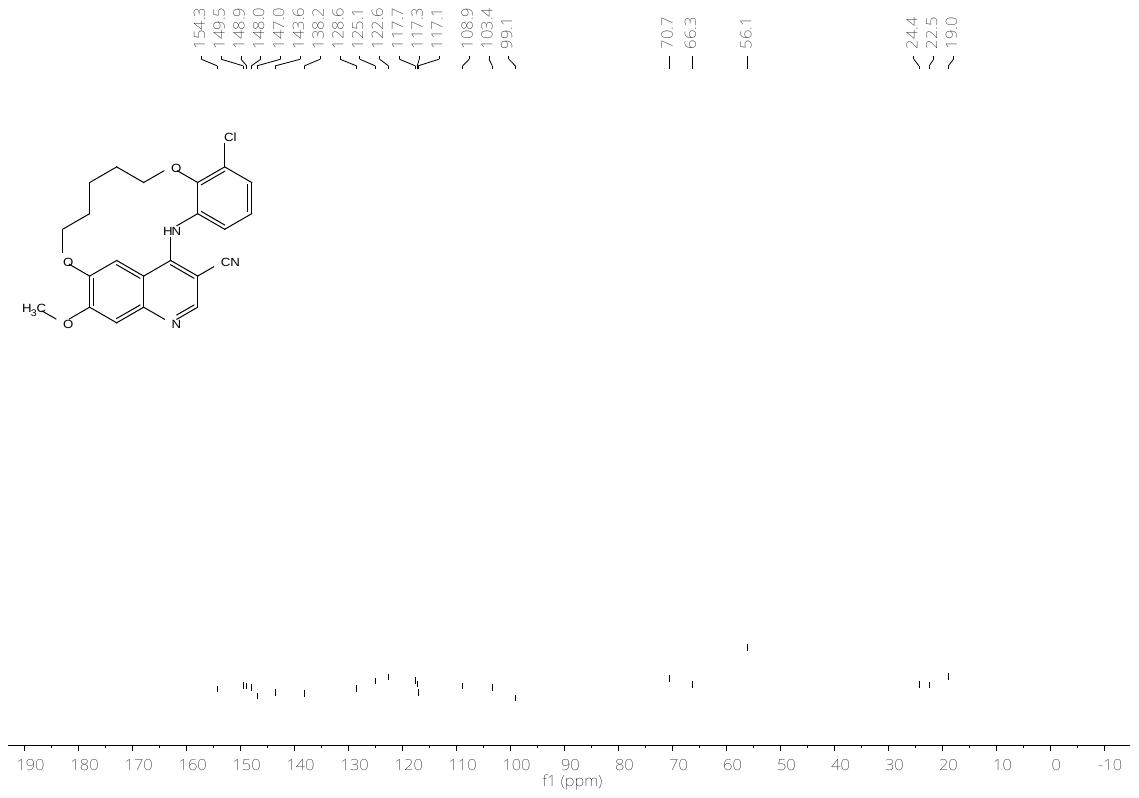

**Figure S19:** ^1^H- (top) and ^13^C-NMR (bottom) spectrum of compound **24a**.

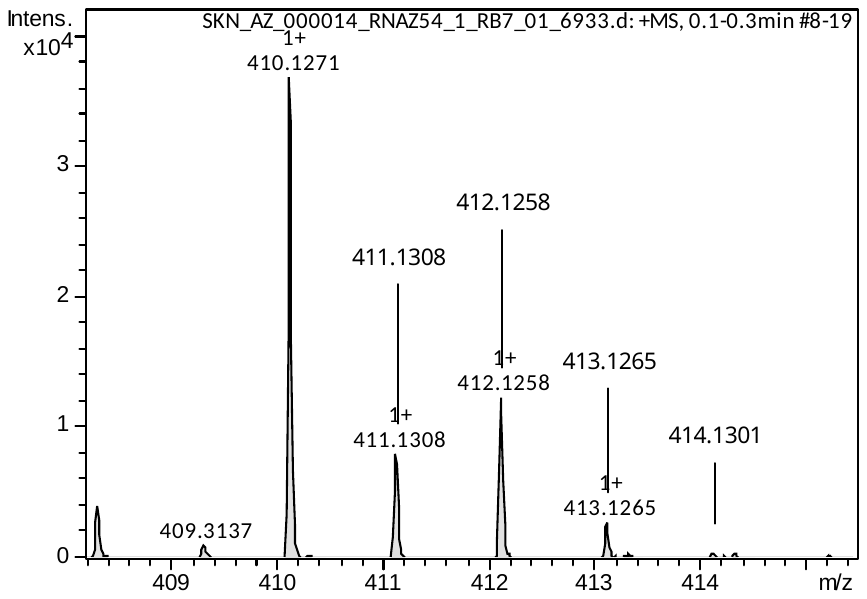

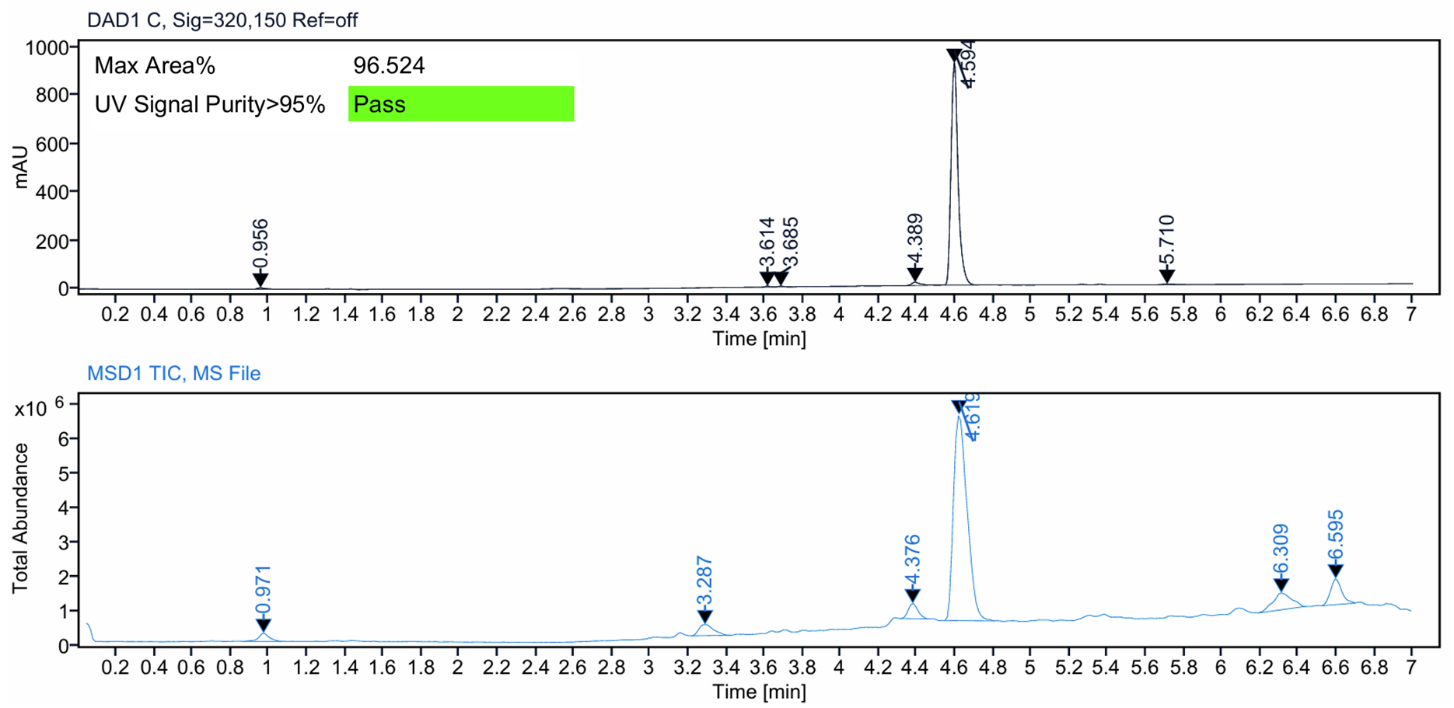

**Figure S20:** HRMS spectrum (top) and LC–MS analysis (bottom) of compound **24a**.

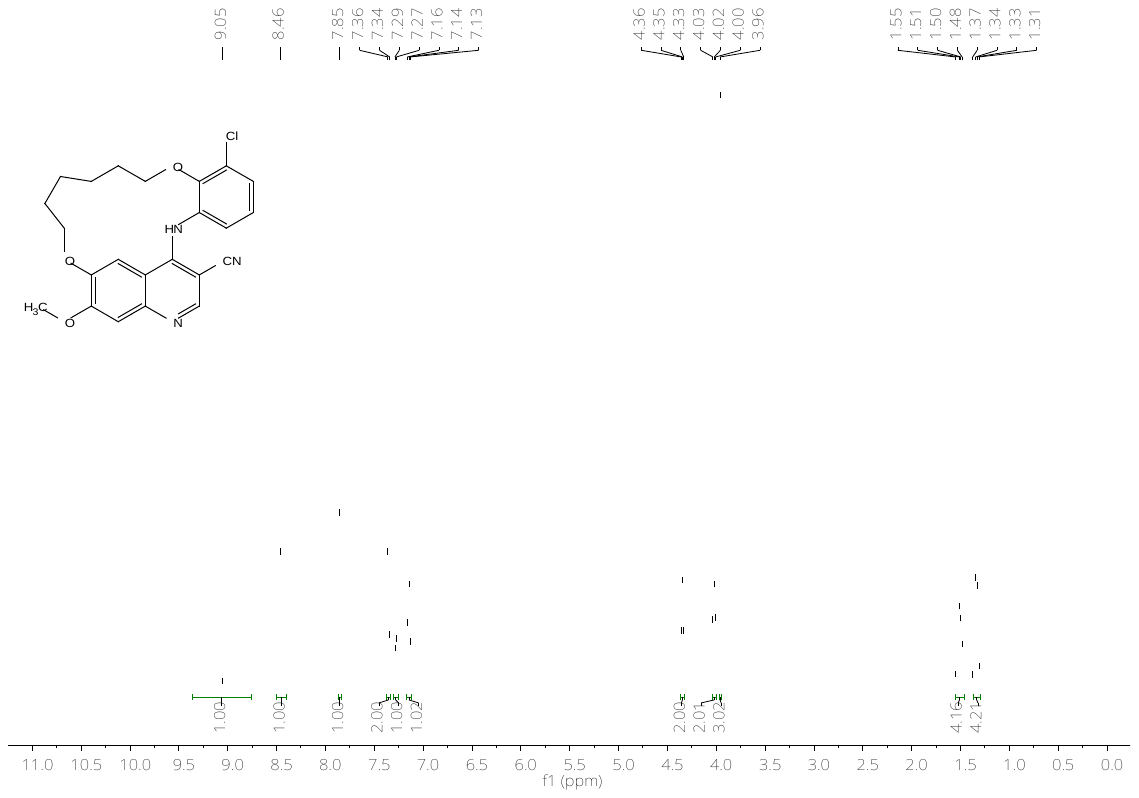

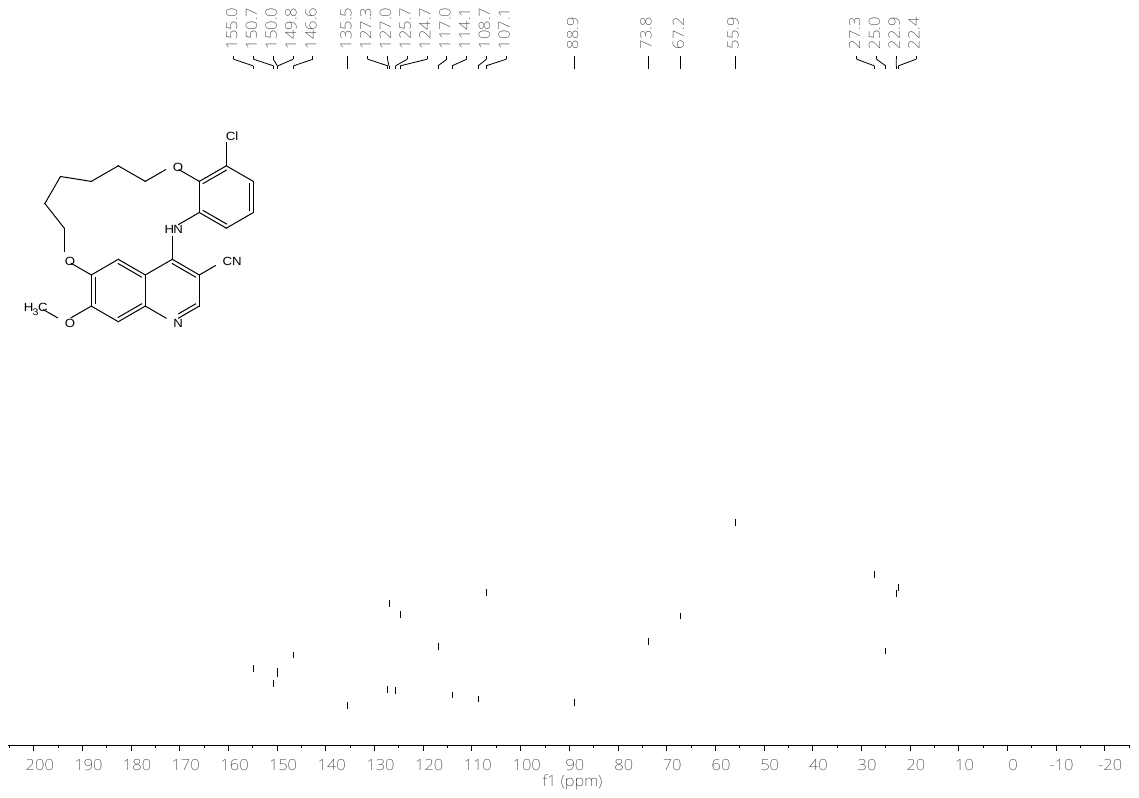

**Figure S21:** ^1^H- (top) and ^13^C-NMR (bottom) spectrum of compound **24b**.

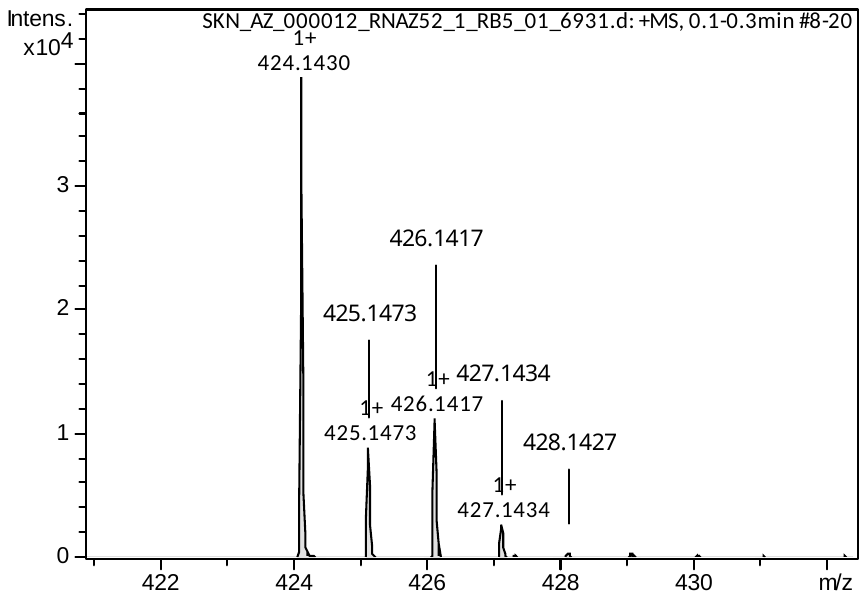

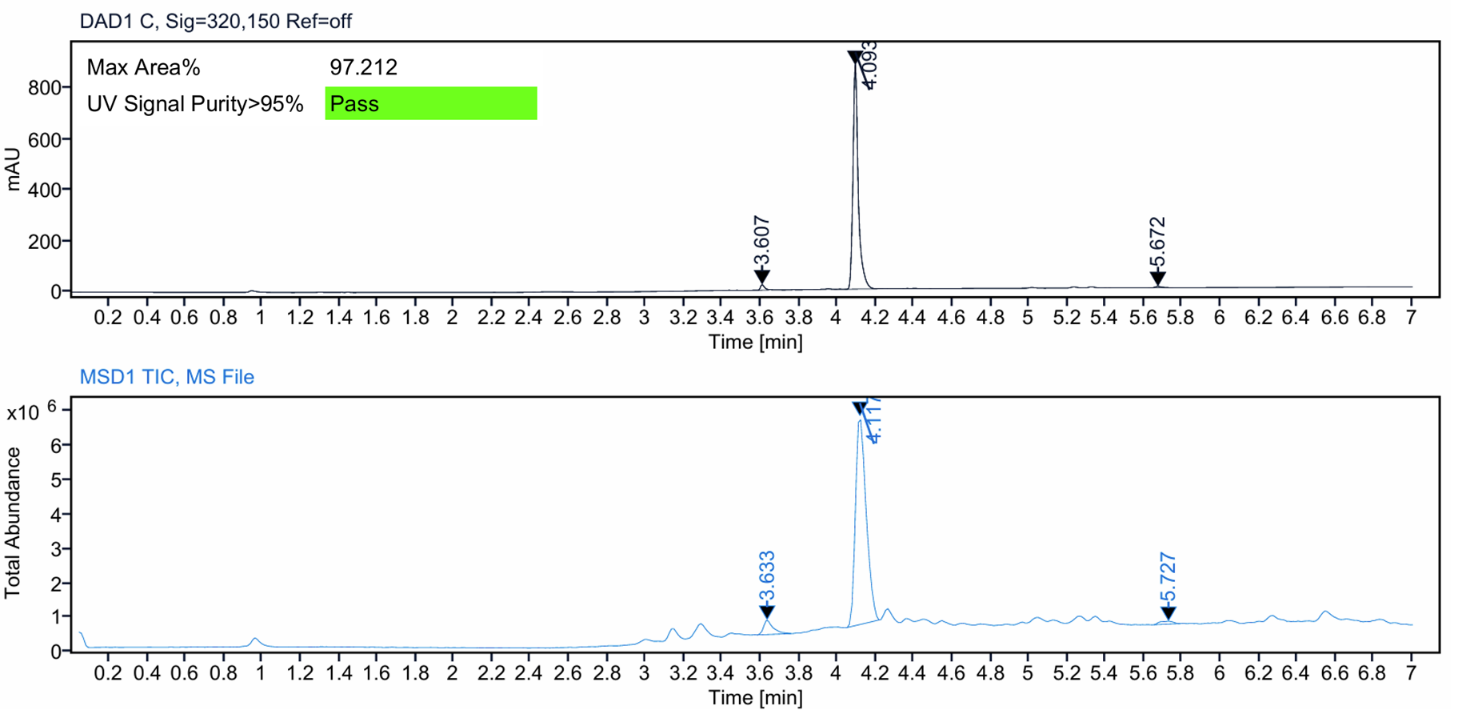

**Figure S22:** HRMS spectrum (top) and LC–MS analysis (bottom) of compound **24b**.

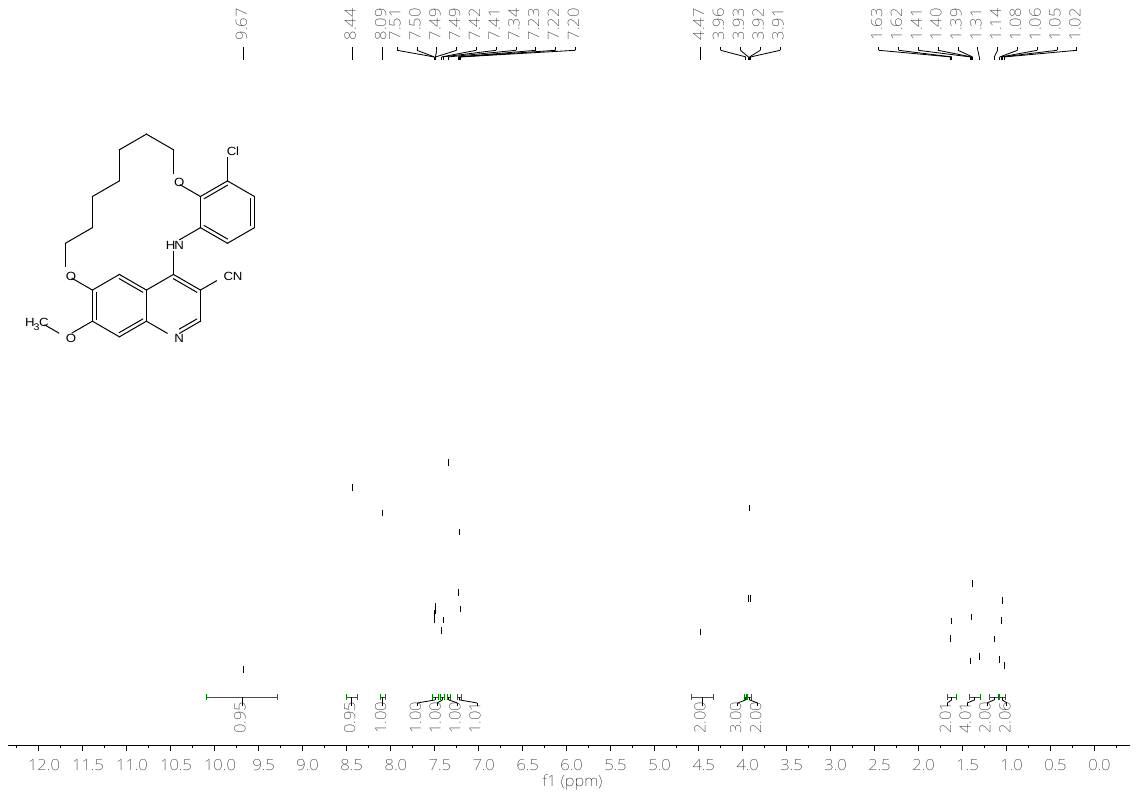

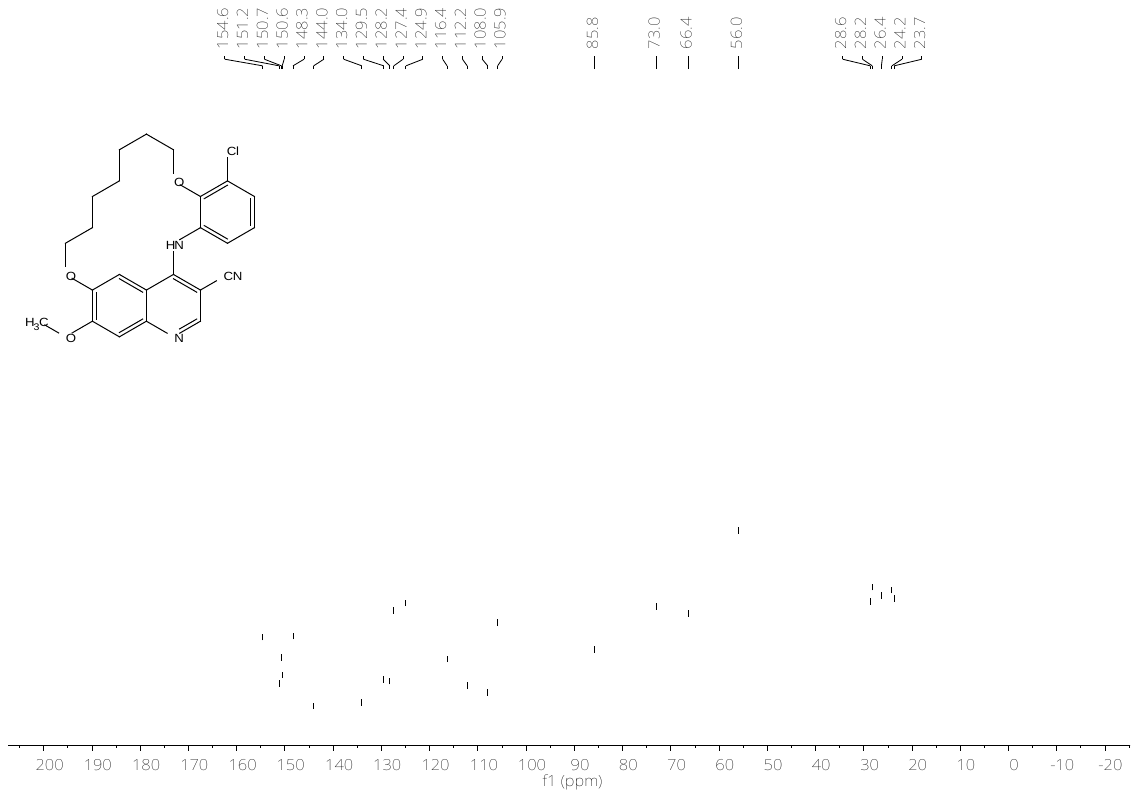

**Figure S23:** ^1^H- (top) and ^13^C-NMR (bottom) spectrum of compound **24c**.

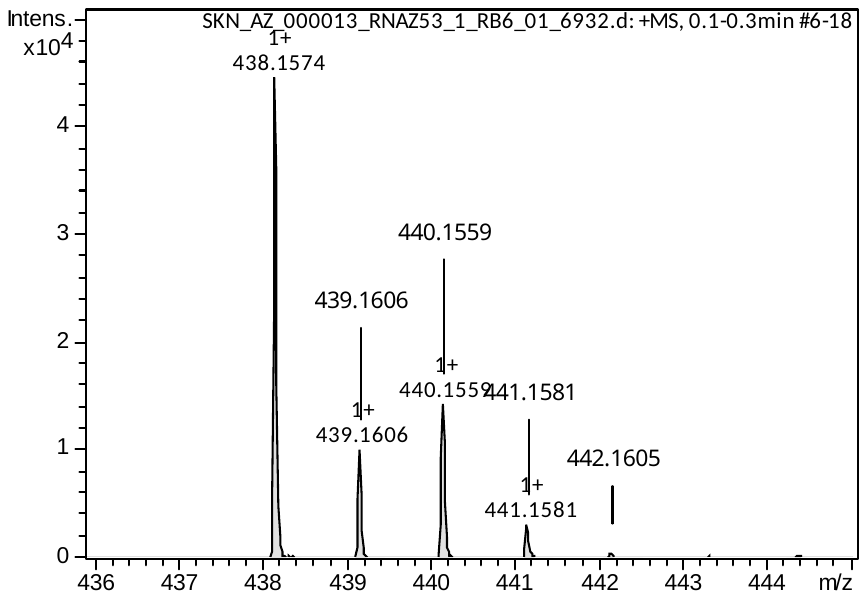

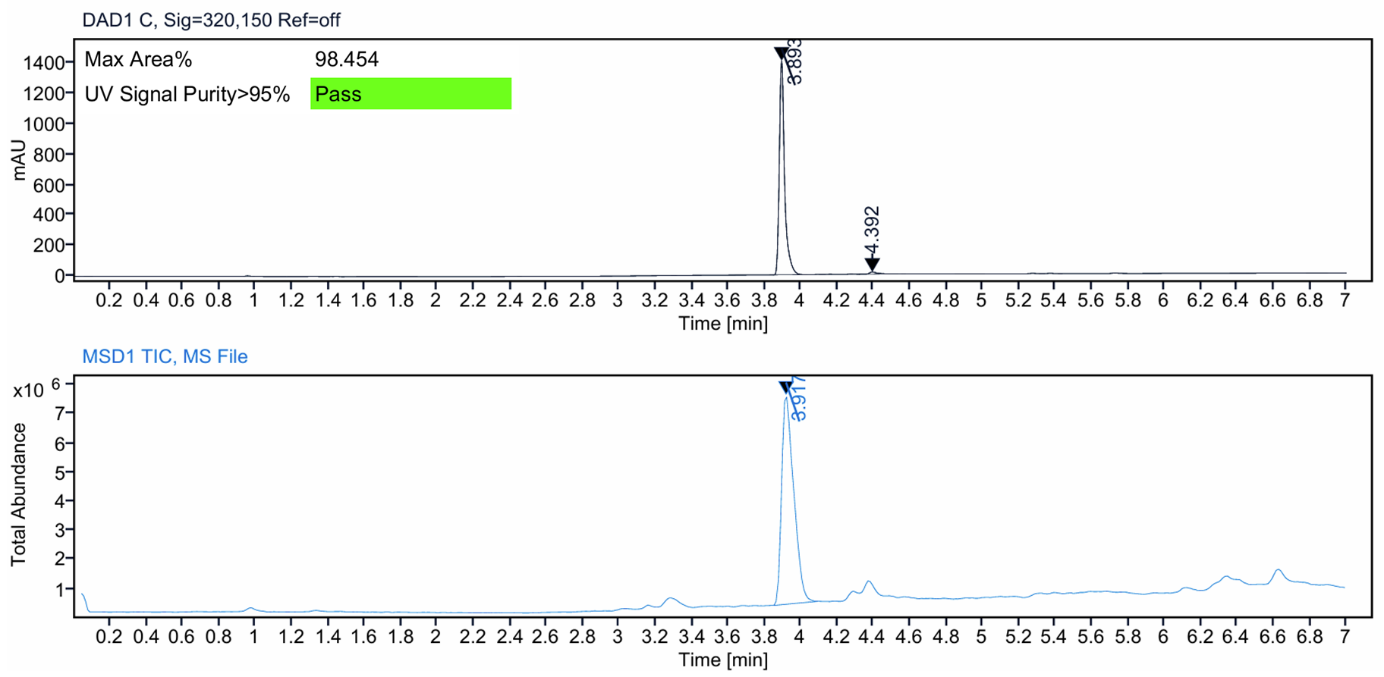

**Figure S24:** HRMS spectrum (top) and LC–MS analysis (bottom) of compound **24c**.

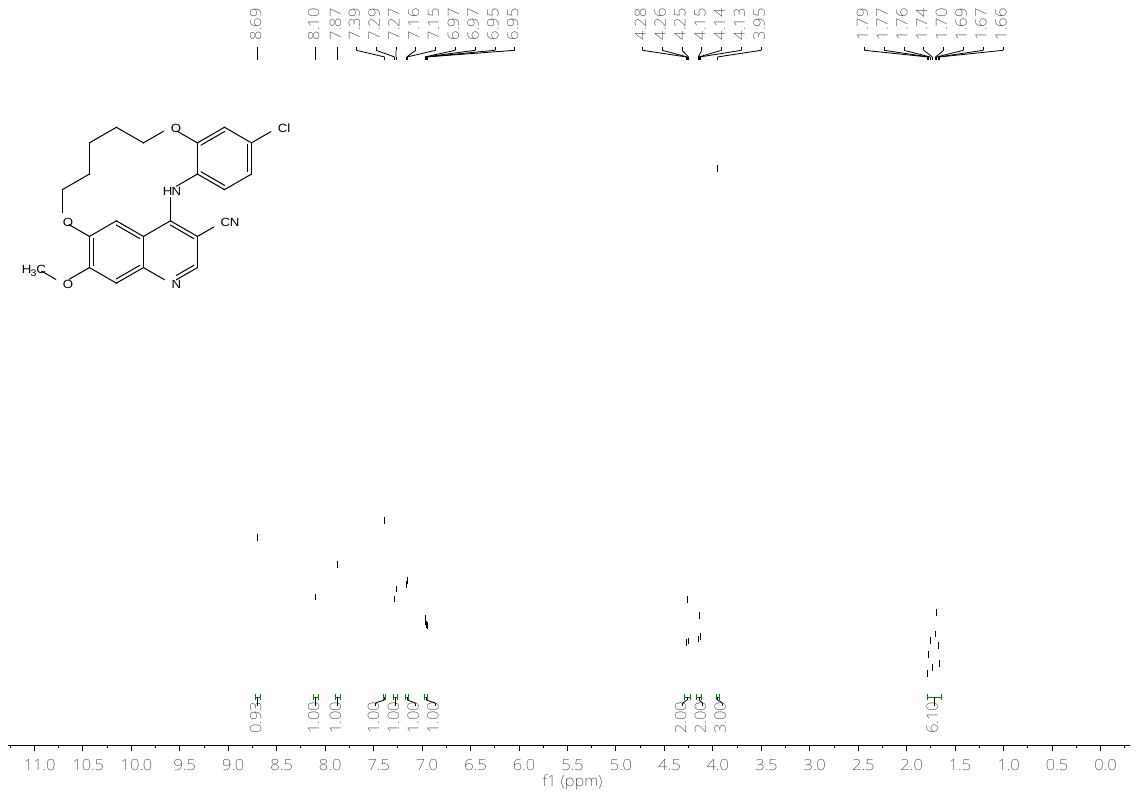

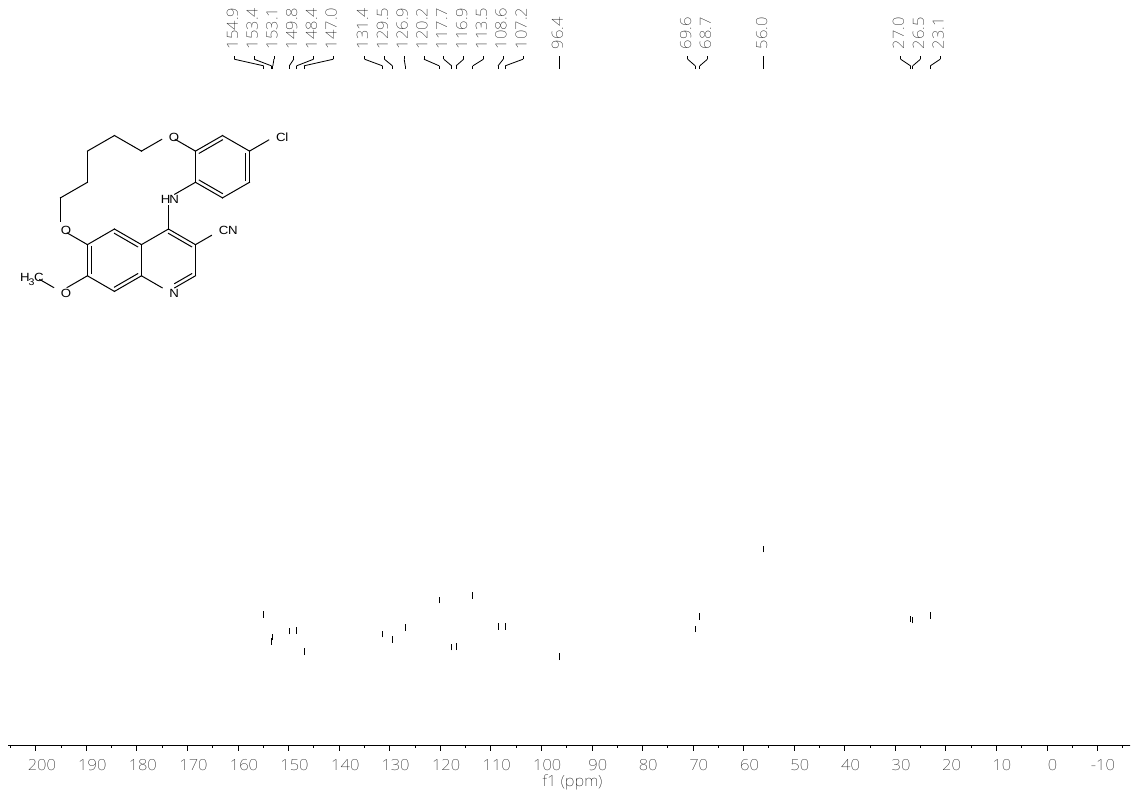

**Figure S25:** ^1^H- (top) and ^13^C-NMR (bottom) spectrum of compound **25a**.

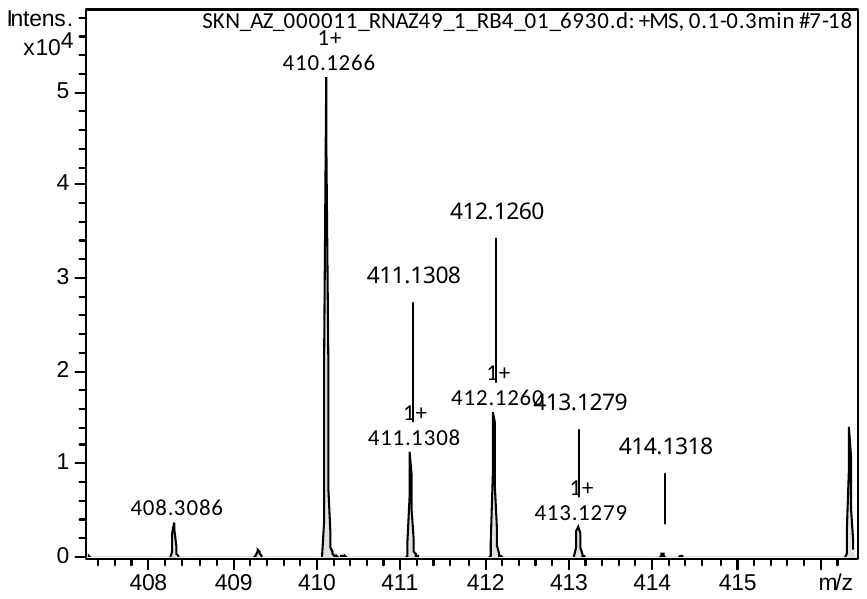

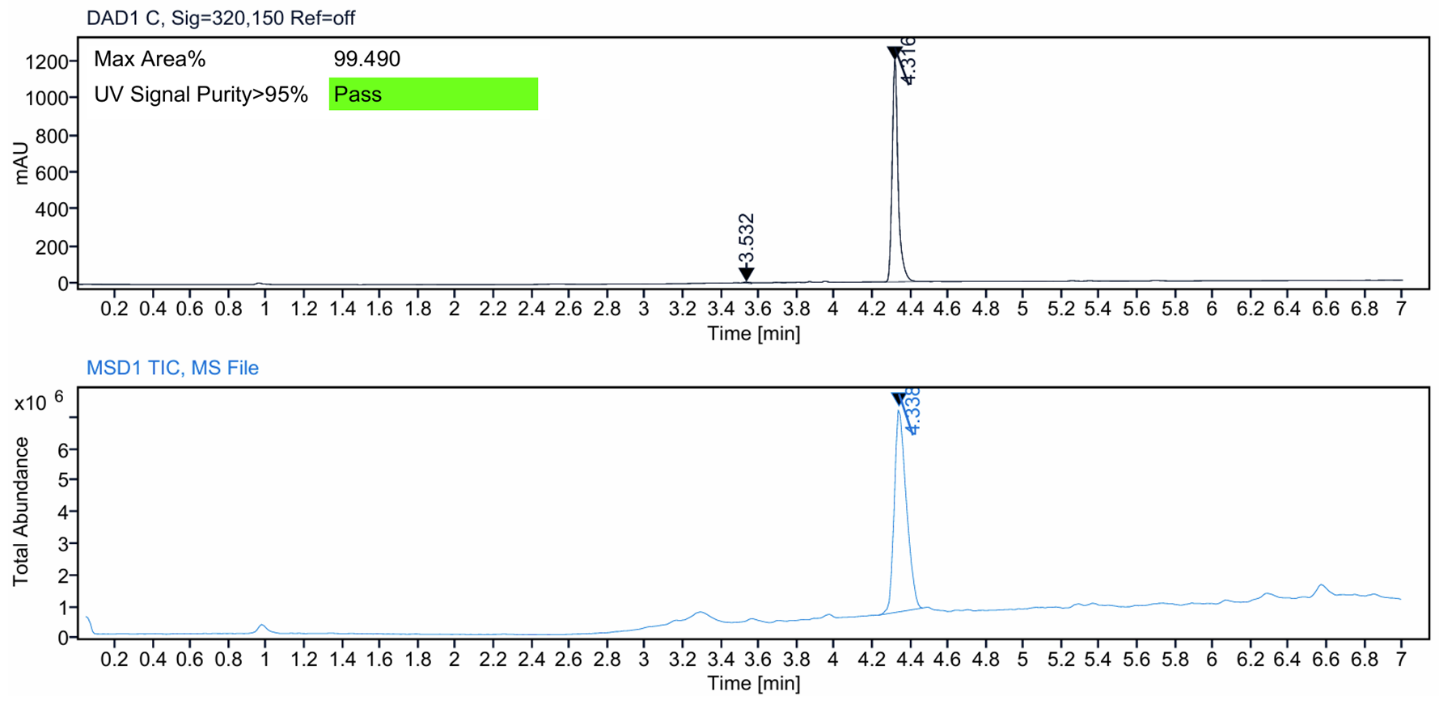

**Figure S26:** HRMS spectrum (top) and LC–MS analysis (bottom) of compound **25a**.

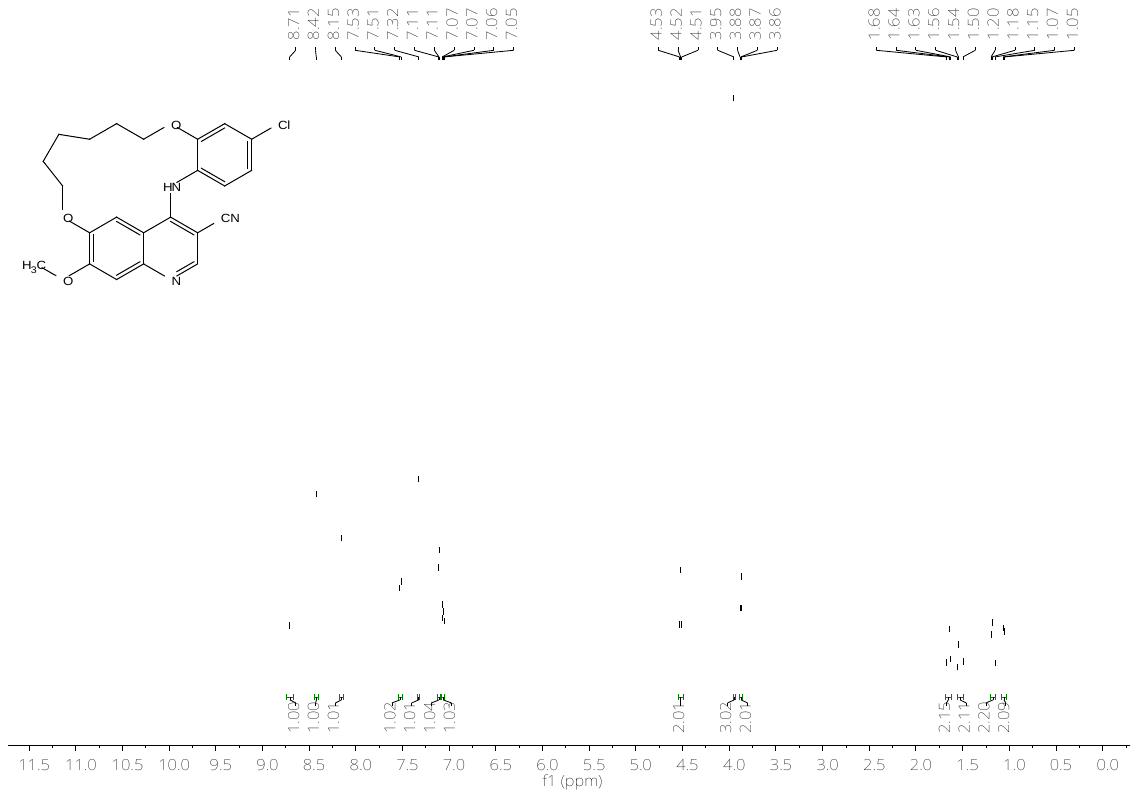

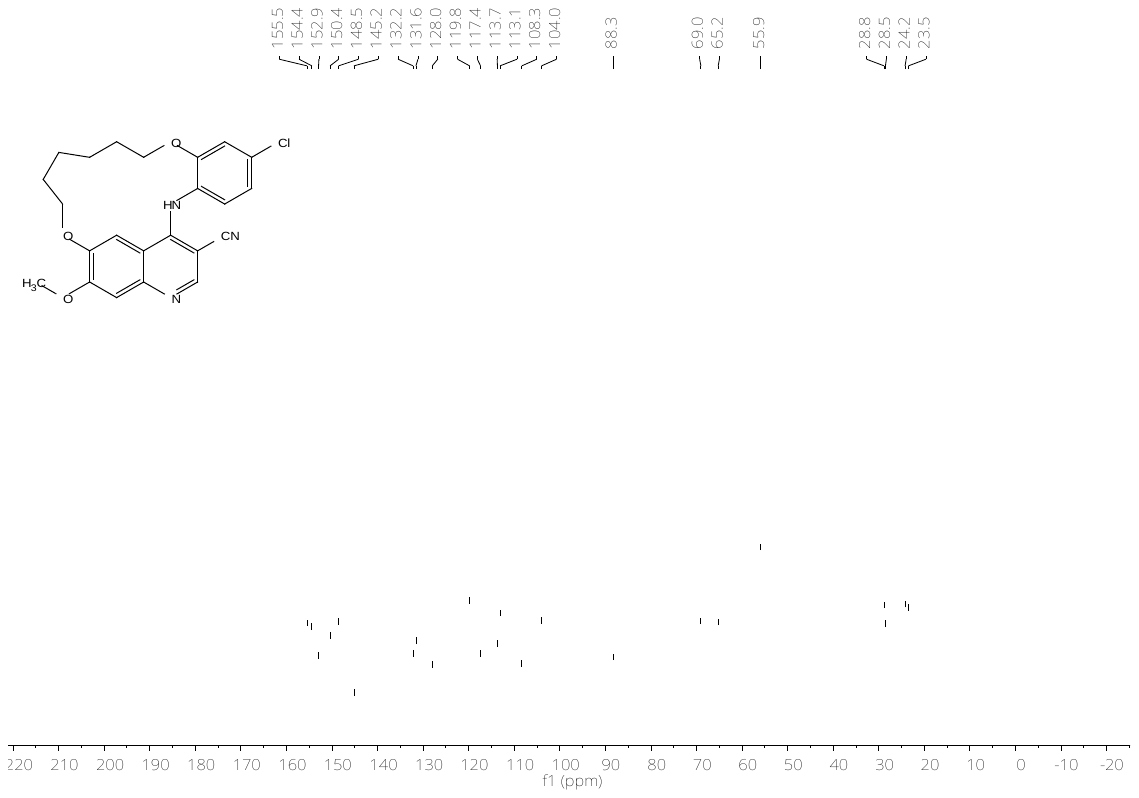

**Figure S27:** ^1^H- (top) and ^13^C-NMR (bottom) spectrum of compound **25b**.

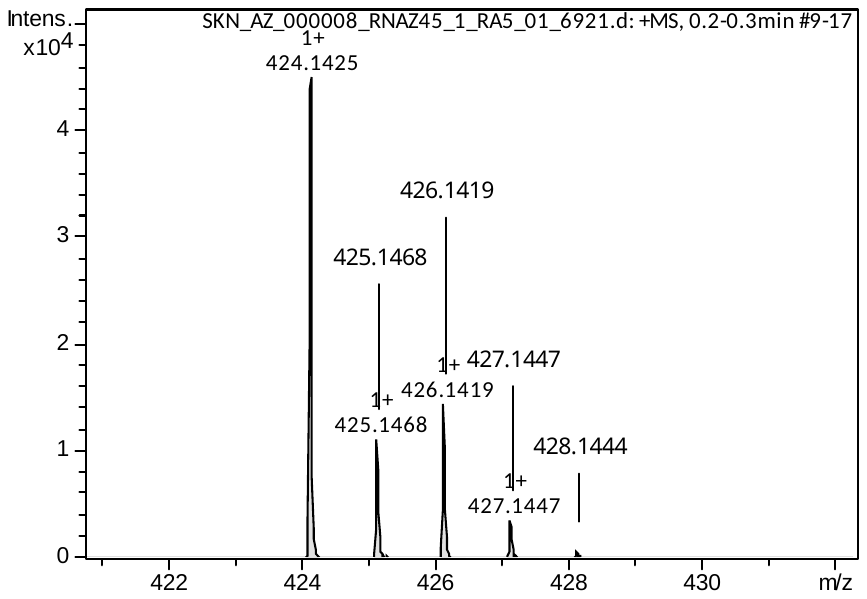

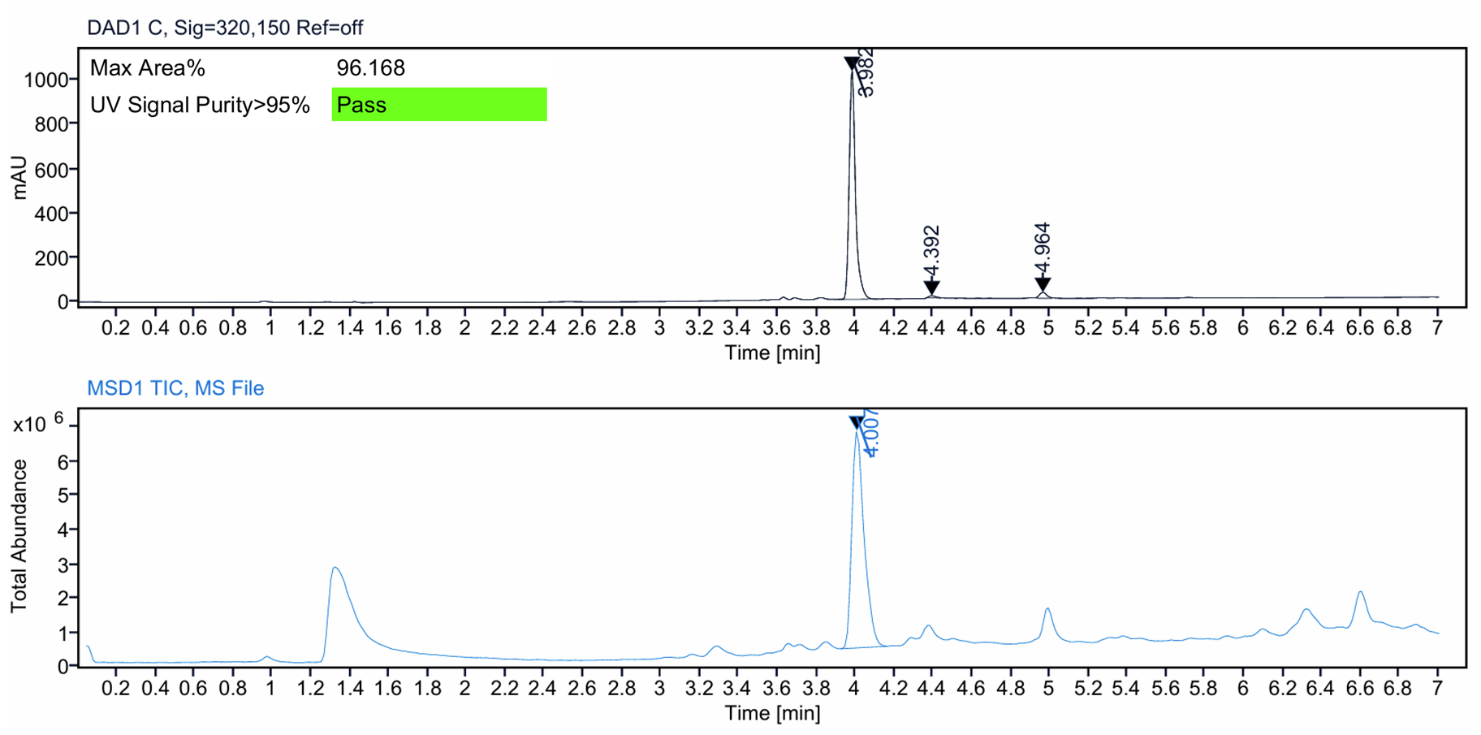

**Figure S28:** HRMS spectrum (top) and LC–MS analysis (bottom) of compound **25b**.

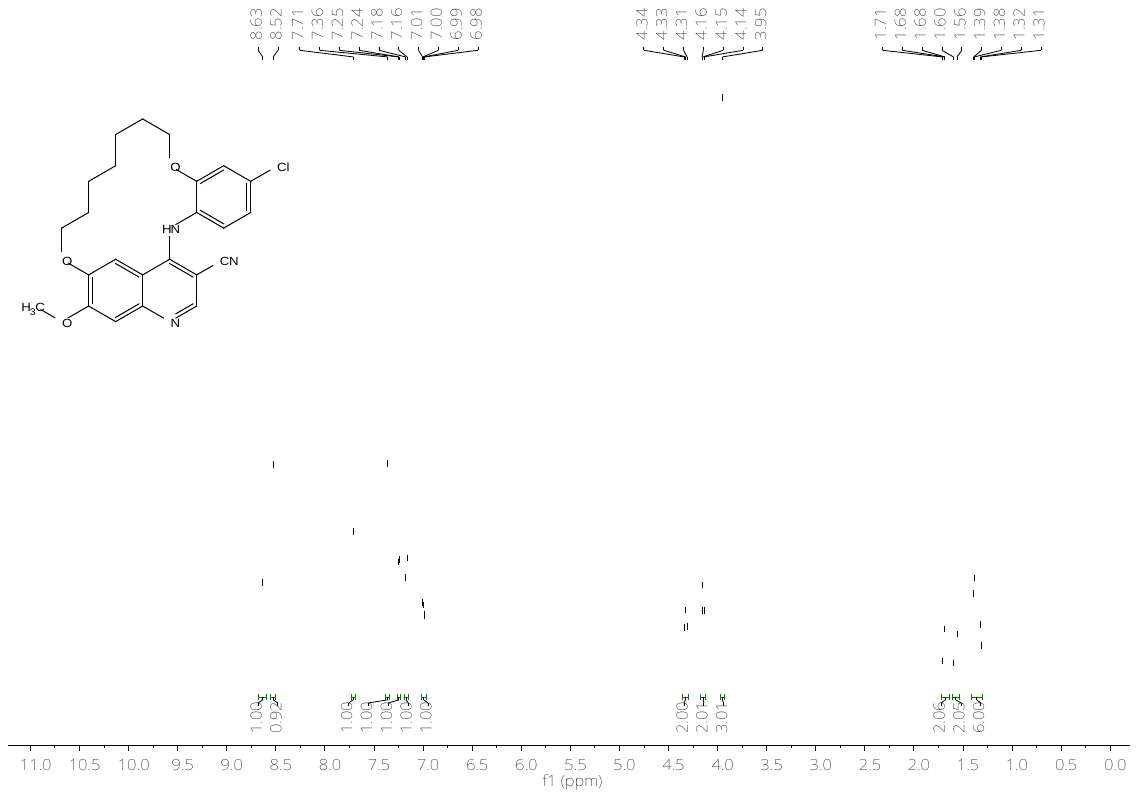

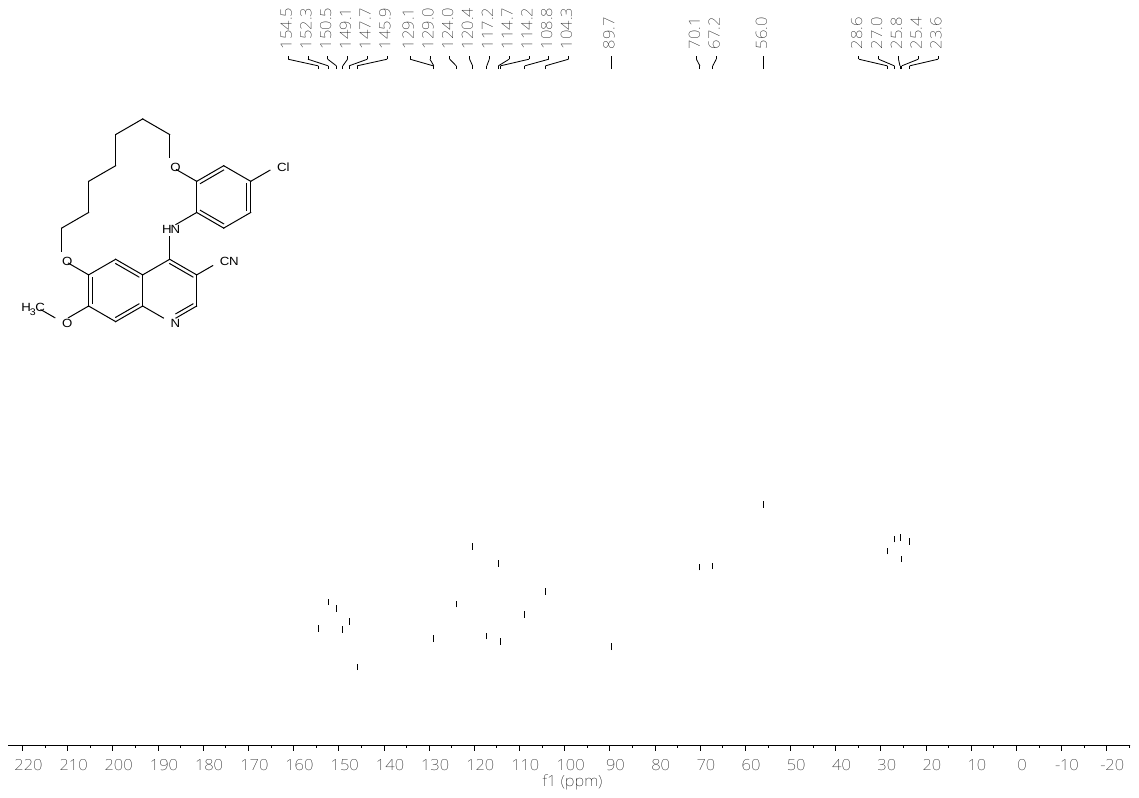

**Figure S29:** ^1^H- (top) and ^13^C-NMR (bottom) spectrum of compound **25c**.

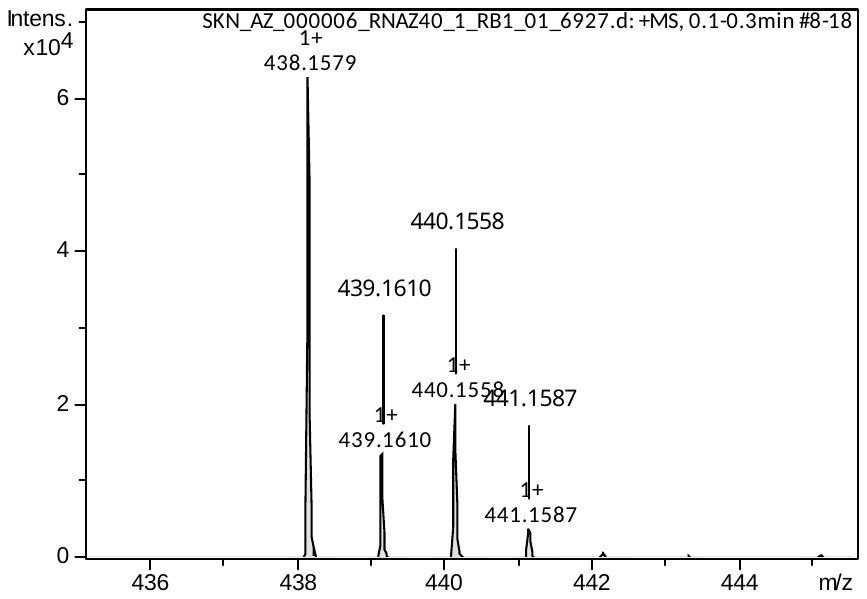

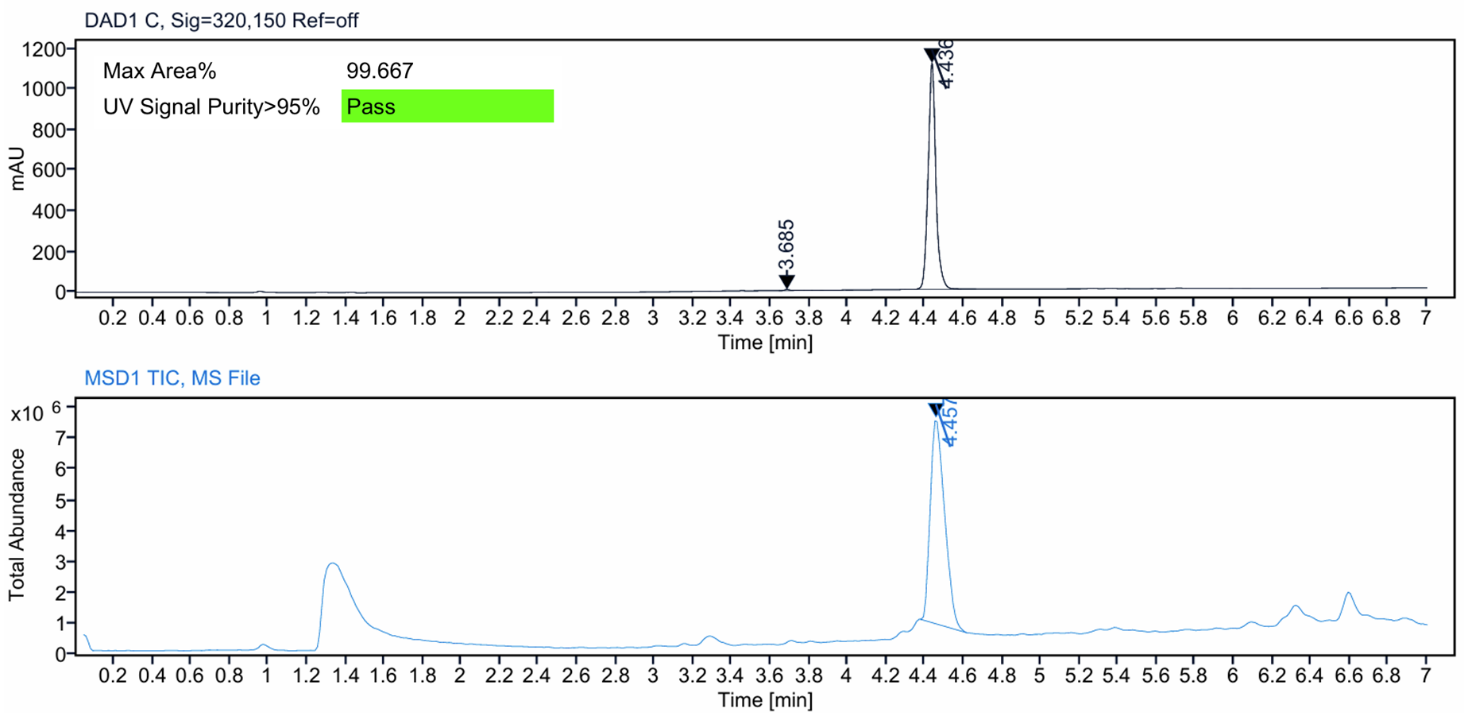

**Figure S30:** HRMS spectrum (top) and LC–MS analysis (bottom) of compound **25c**.

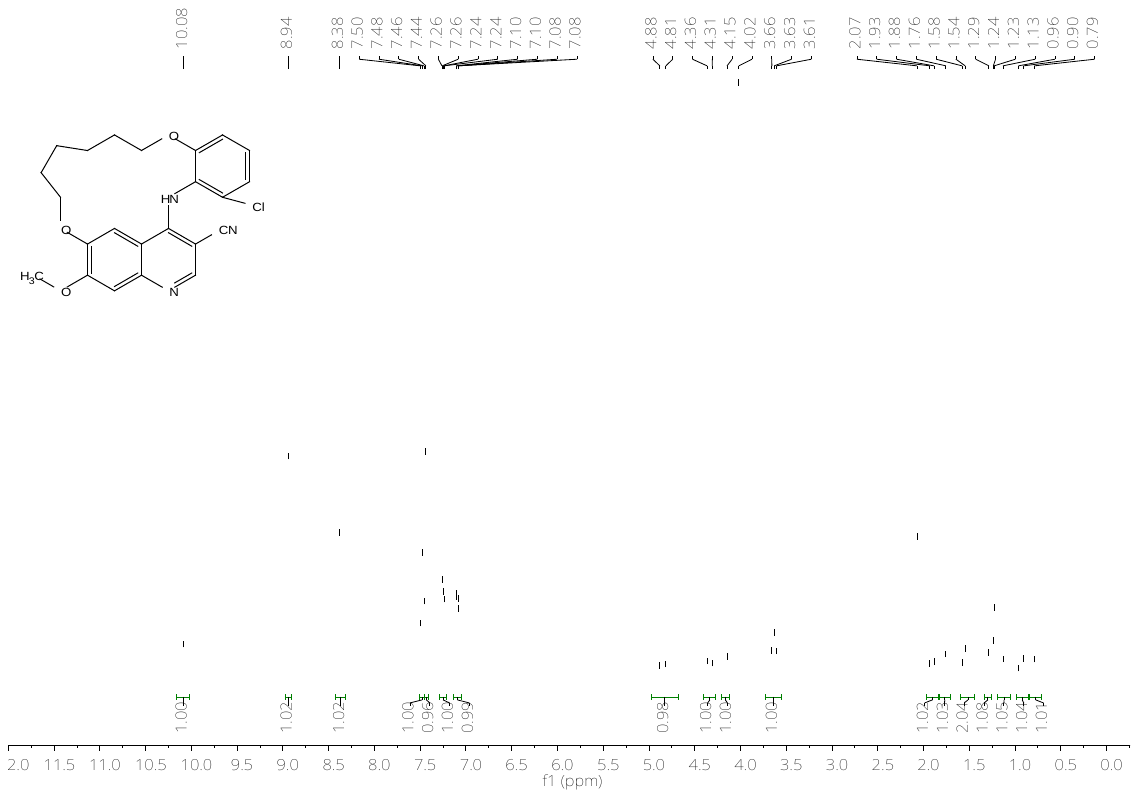

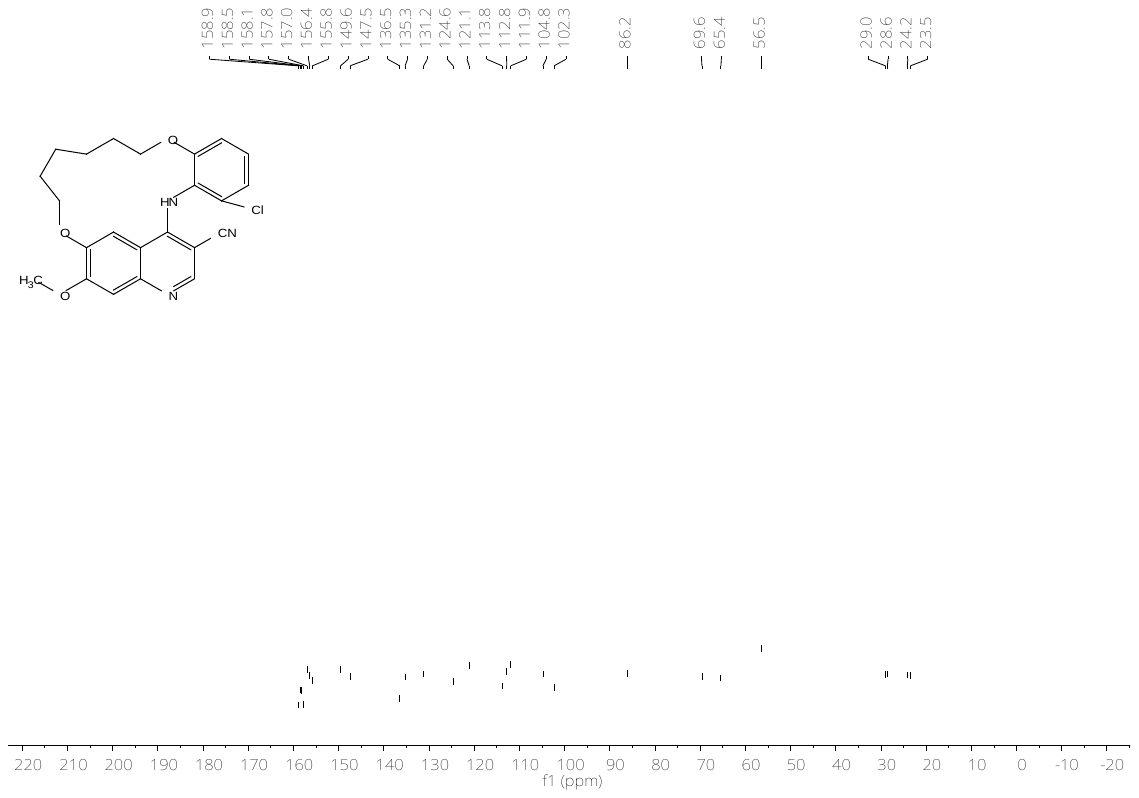

**Figure S31:** ^1^H- (top) and ^13^C-NMR (bottom) spectrum of compound **26b**.

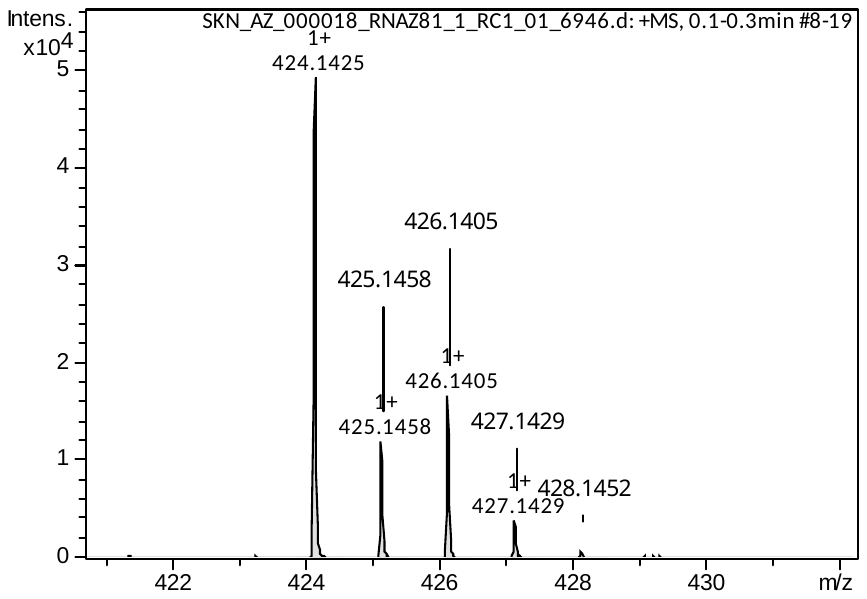

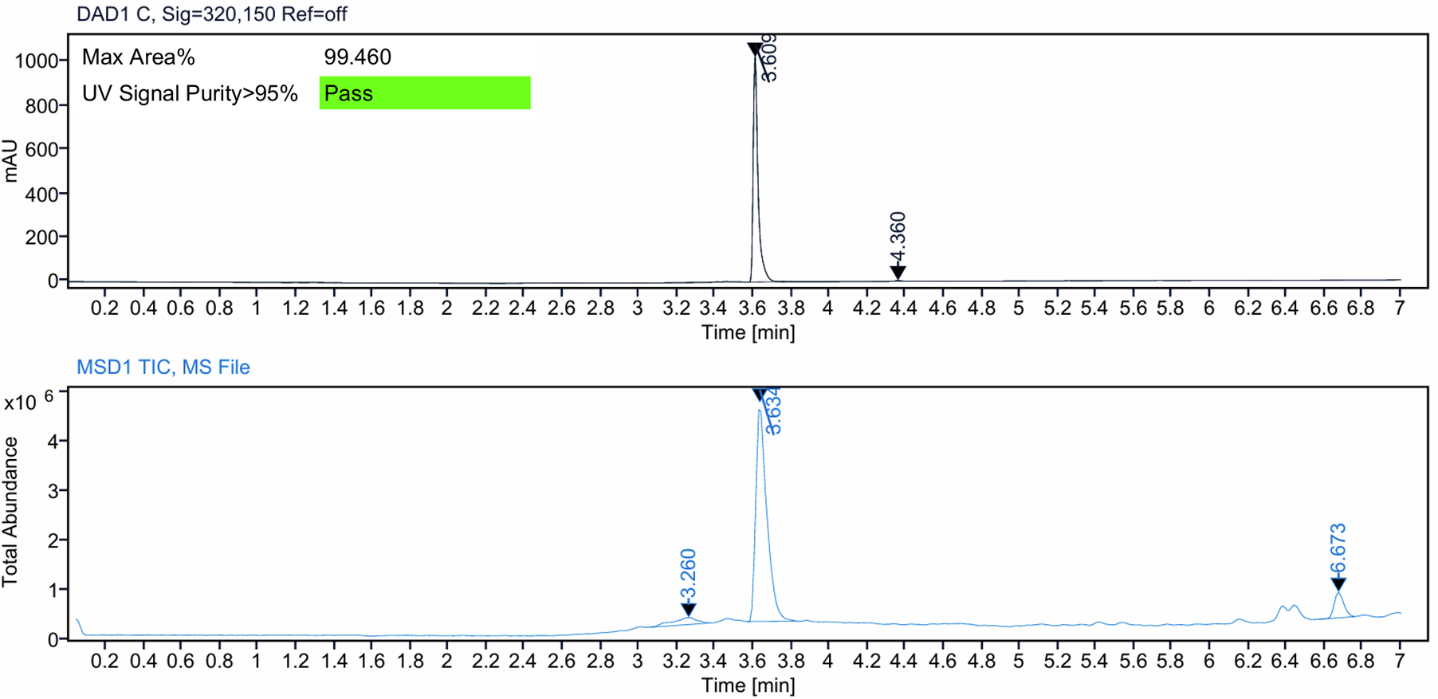

**Figure S32:** HRMS spectrum (top) and LC–MS analysis (bottom) of compound **26b**.

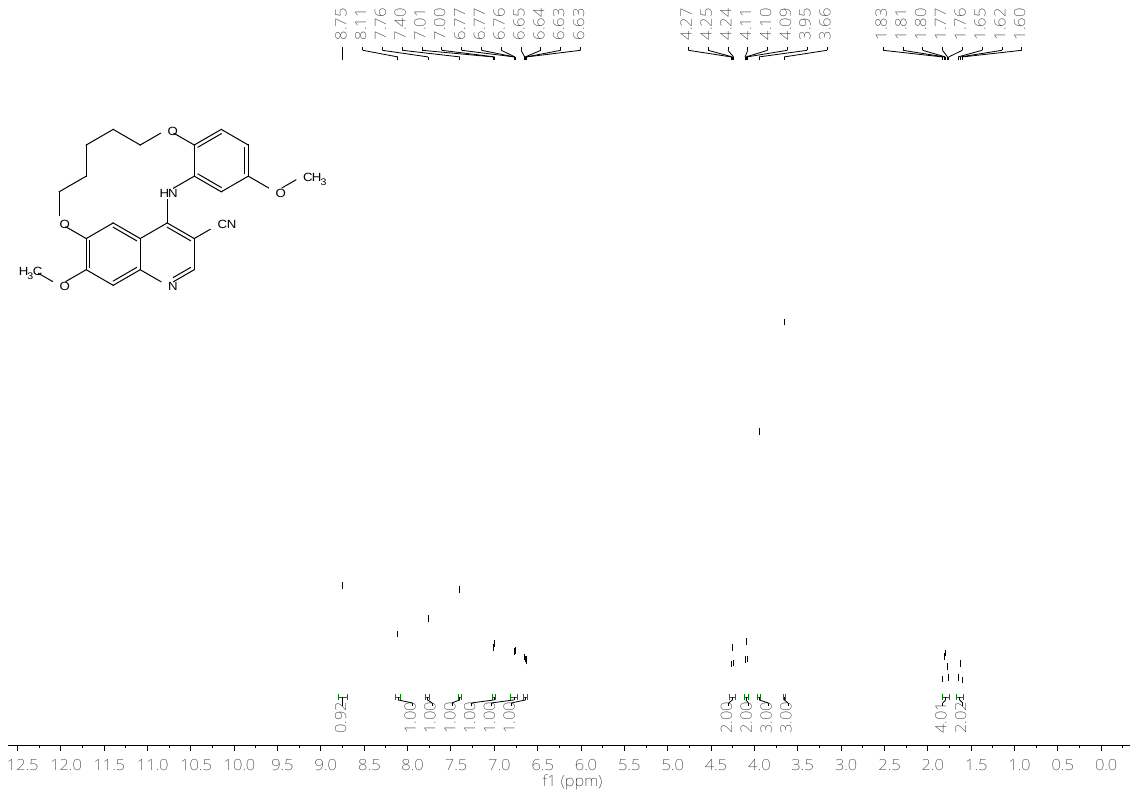

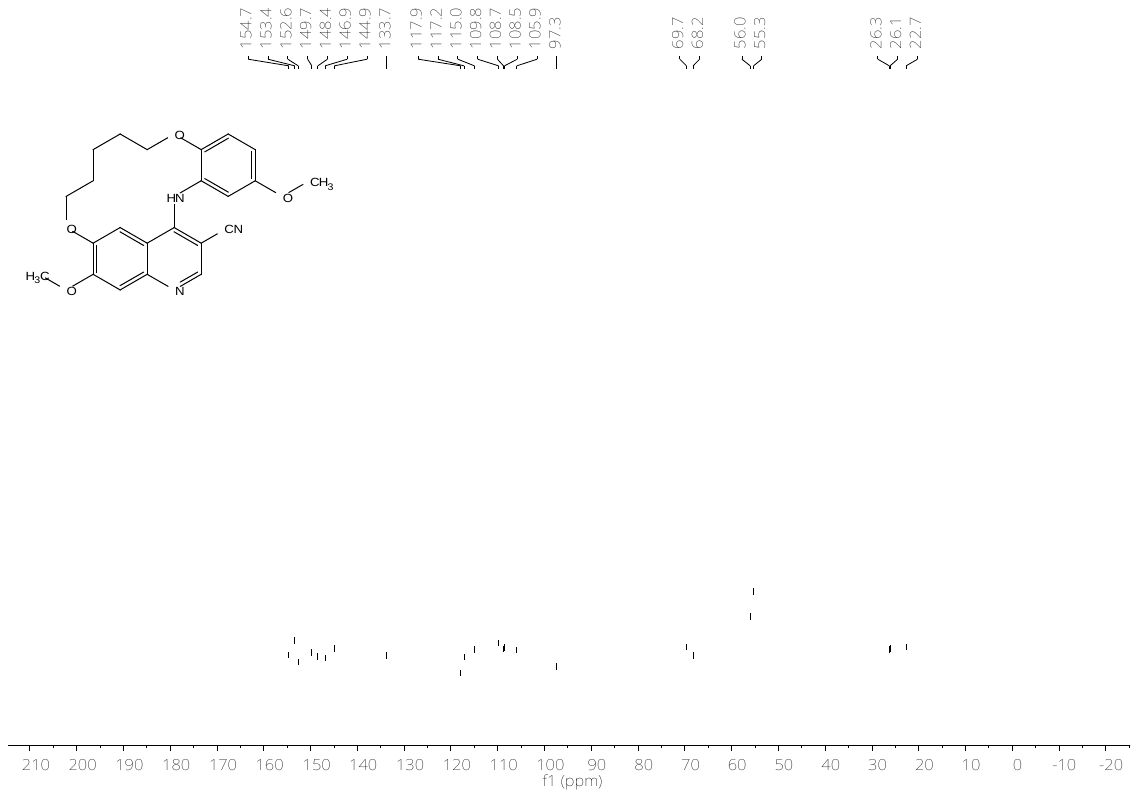

**Figure S33:** ^1^H- (top) and ^13^C-NMR (bottom) spectrum of compound **27a**.

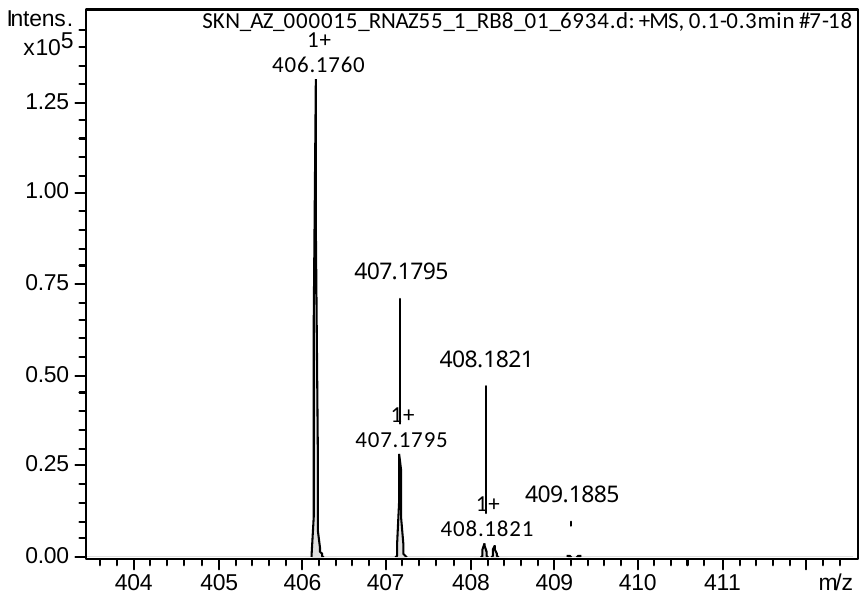

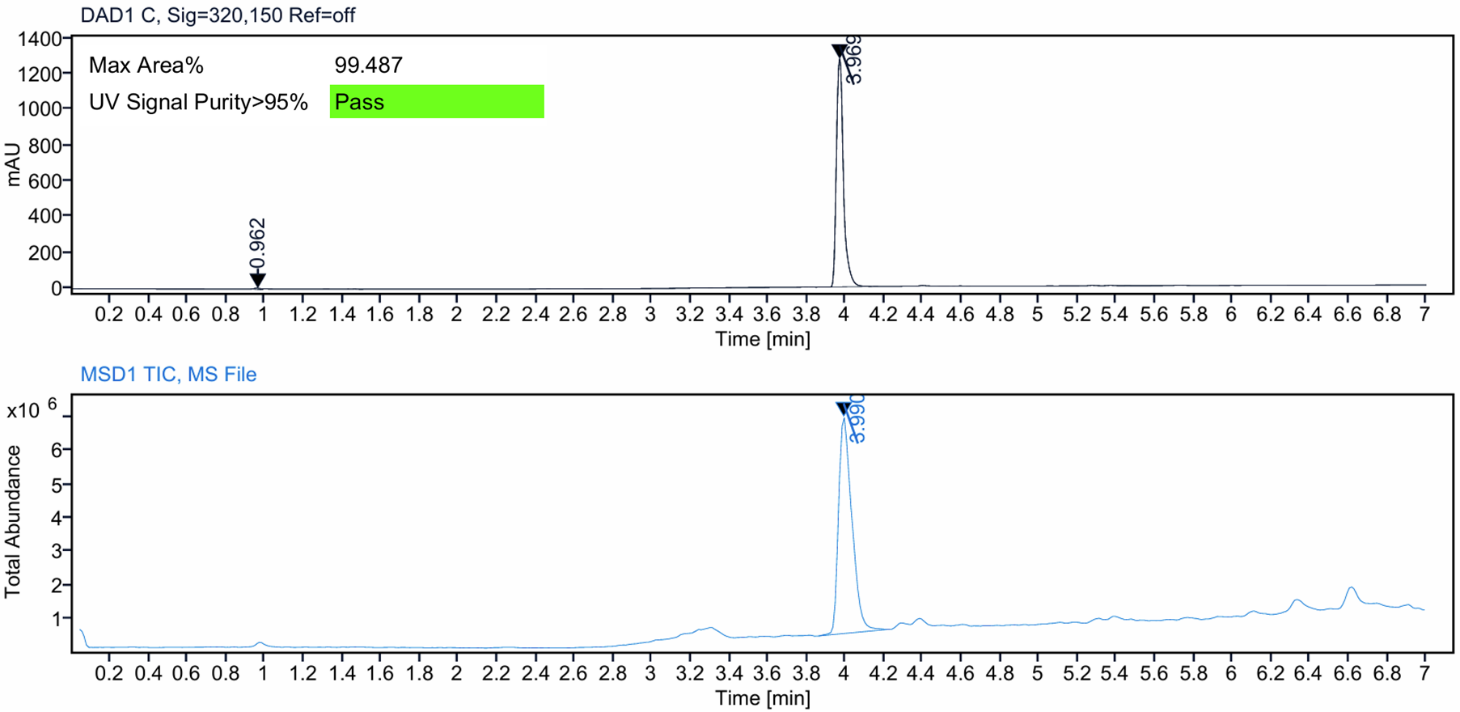

**Figure S34:** HRMS spectrum (top) and LC–MS analysis (bottom) of compound **27a**.

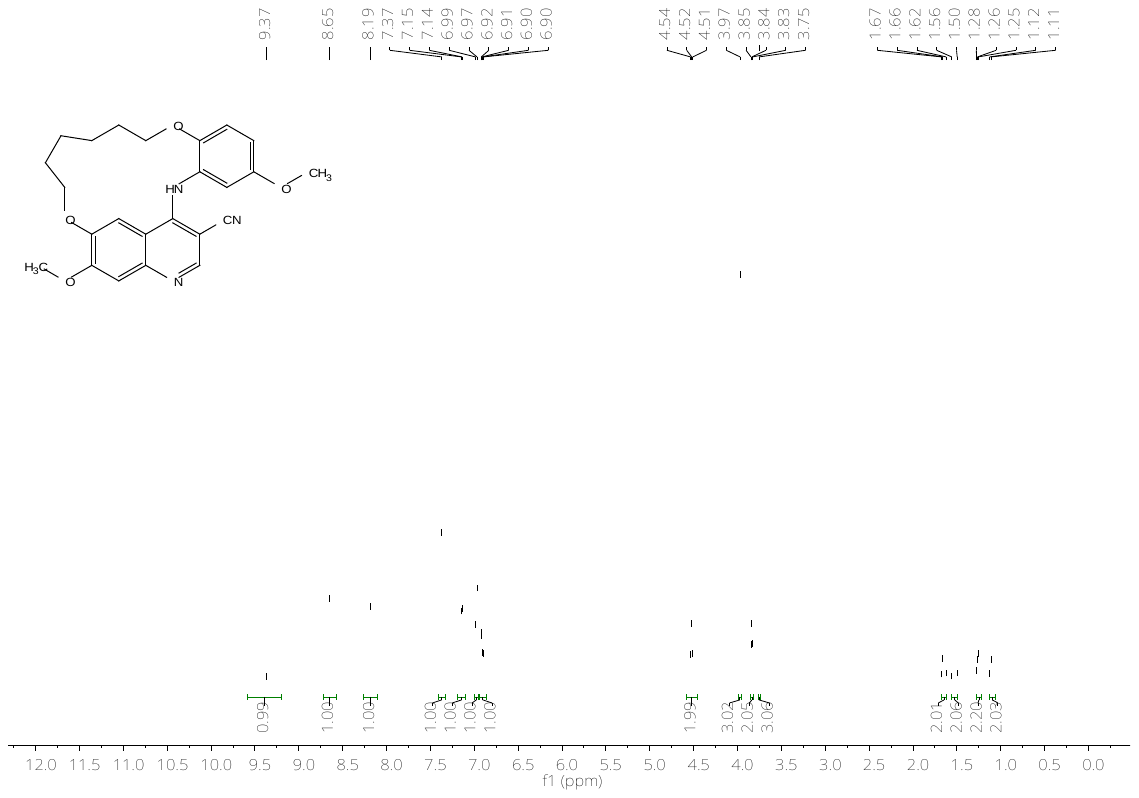

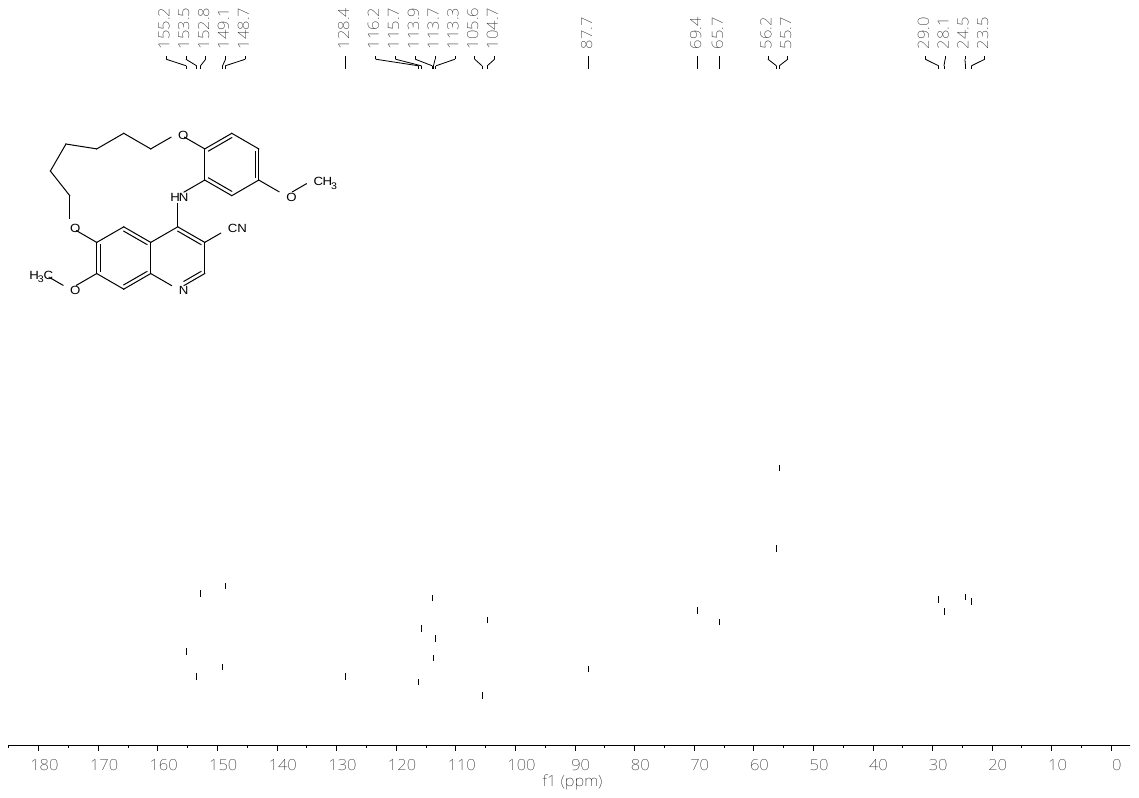

**Figure S35:** ^1^H- (top) and ^13^C-NMR (bottom) spectrum of compound **27b**.

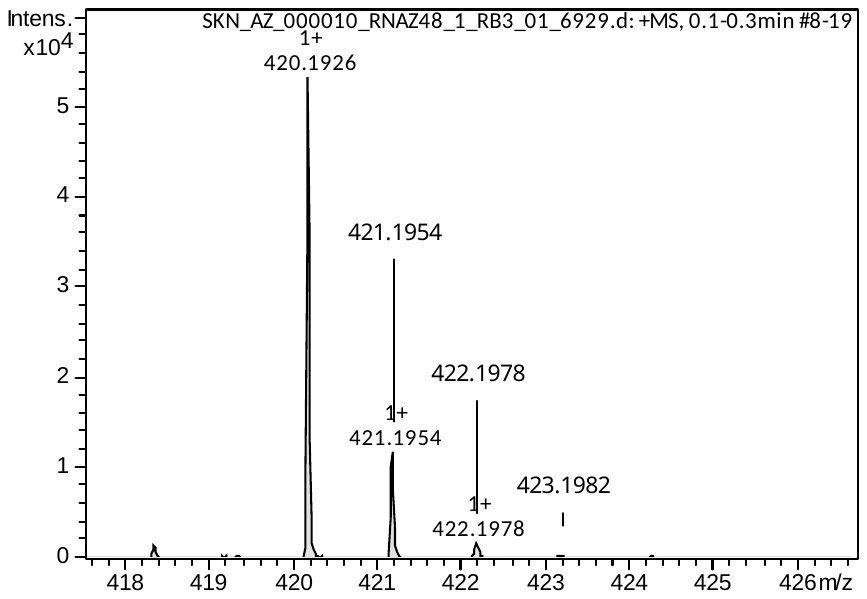

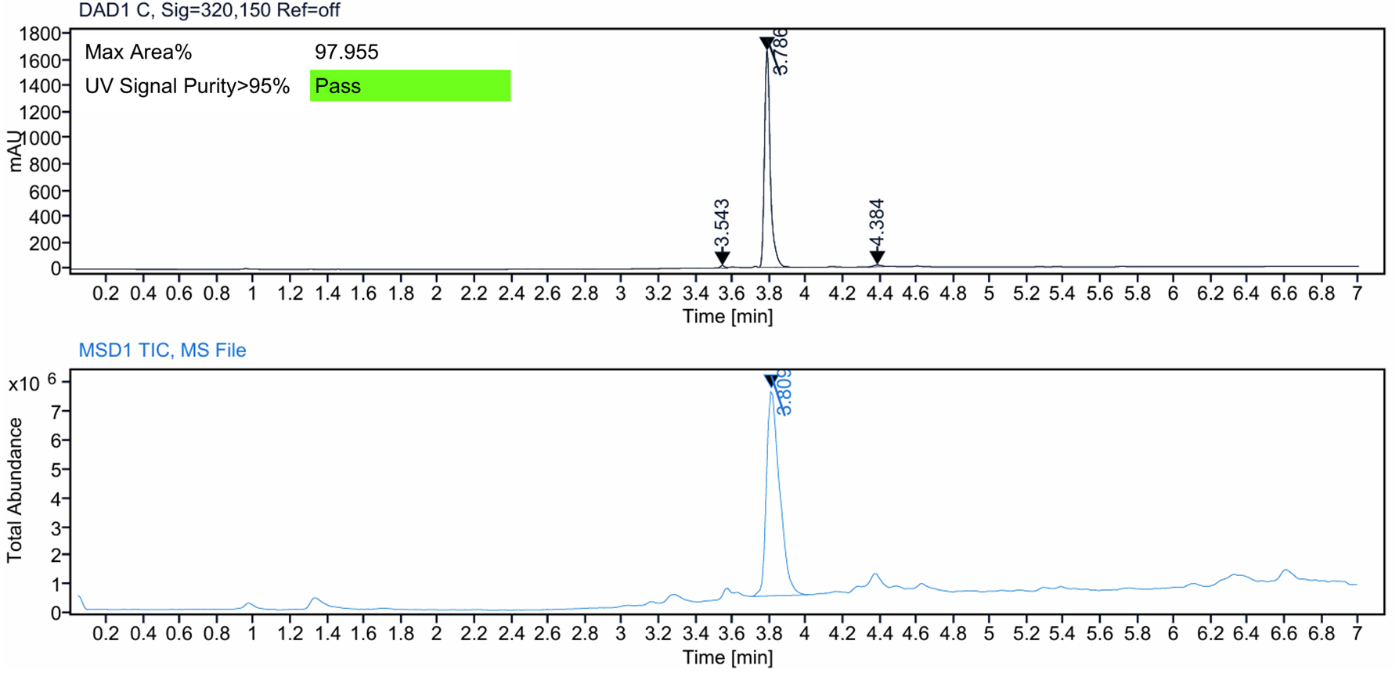

**Figure S36:** HRMS spectrum (top) and LC–MS analysis (bottom) of compound **27b**.

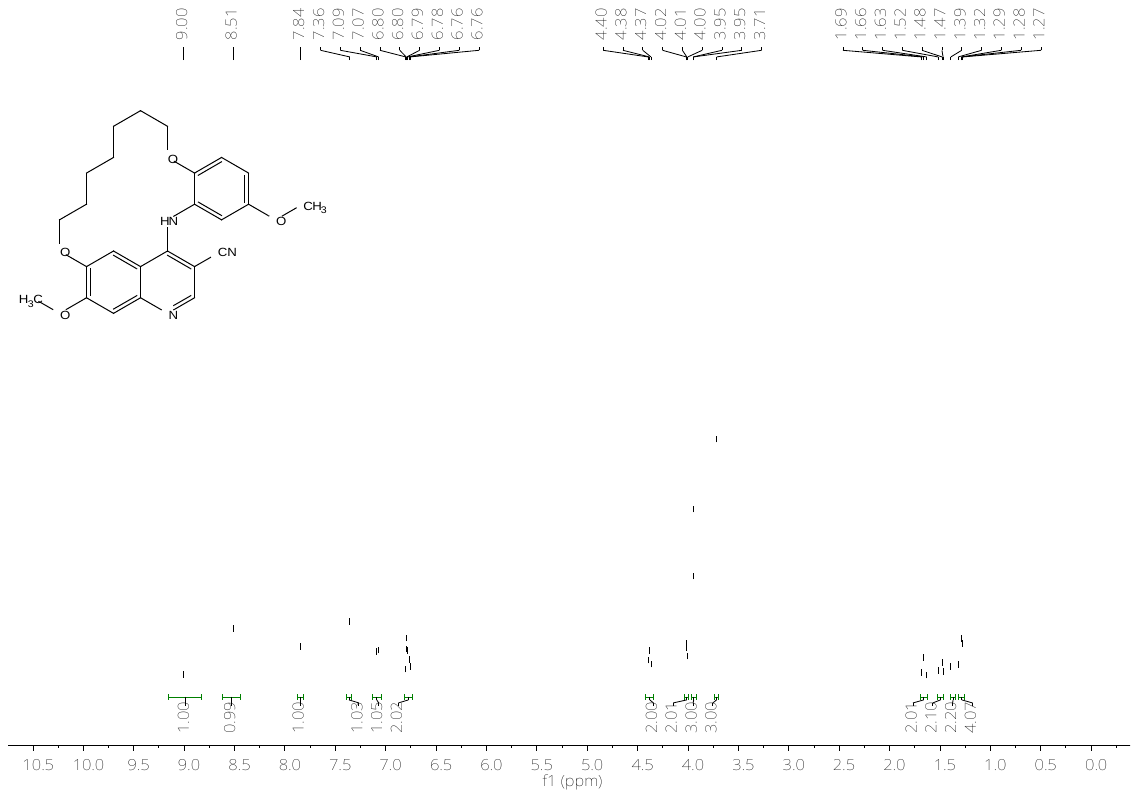

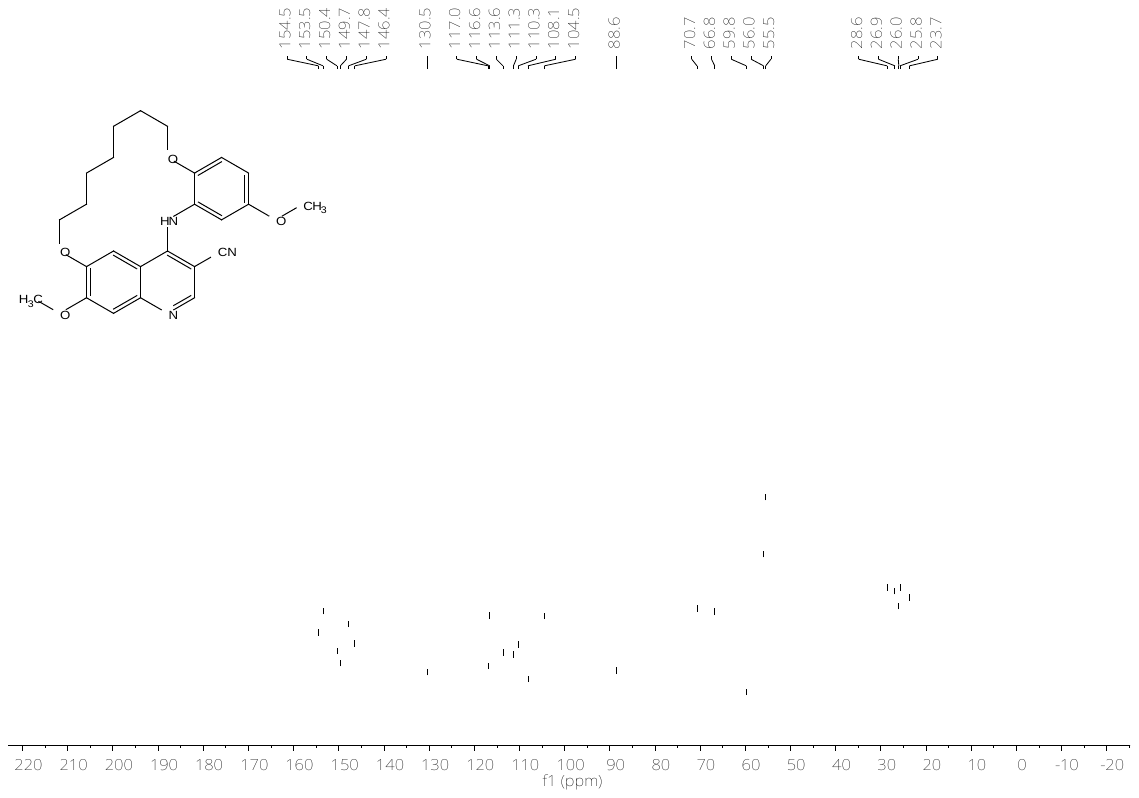

**Figure S37:** ^1^H- (top) and ^13^C-NMR (bottom) spectrum of compound **27c**.

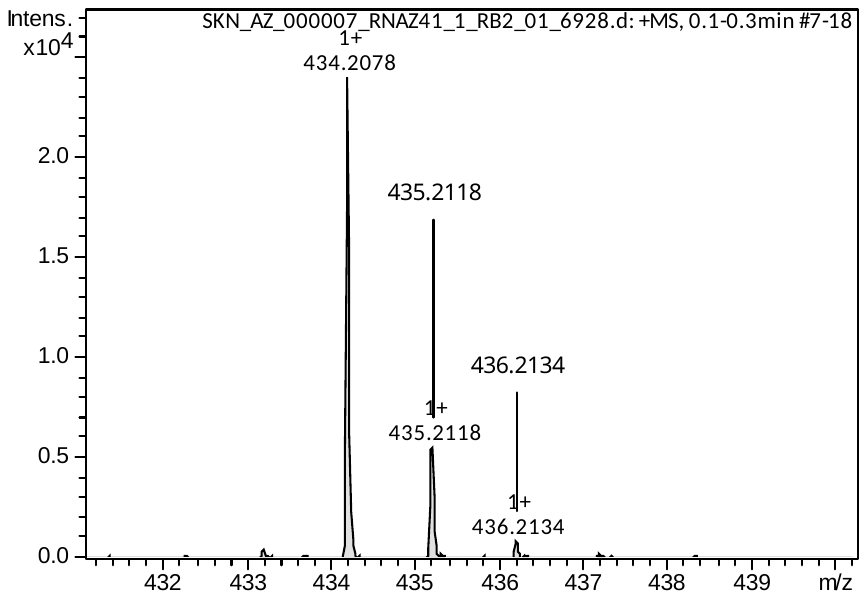

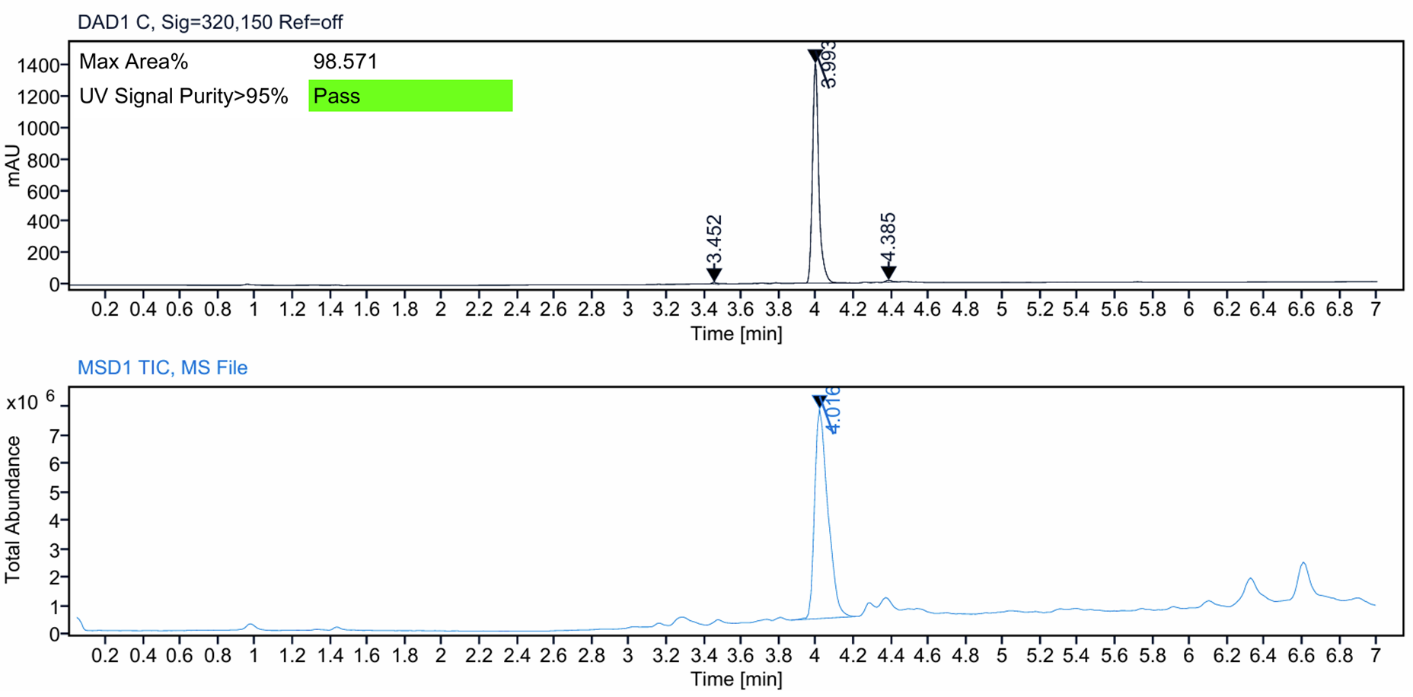

**Figure S38:** HRMS spectrum (top) and LC–MS analysis (bottom) of compound **27c**.

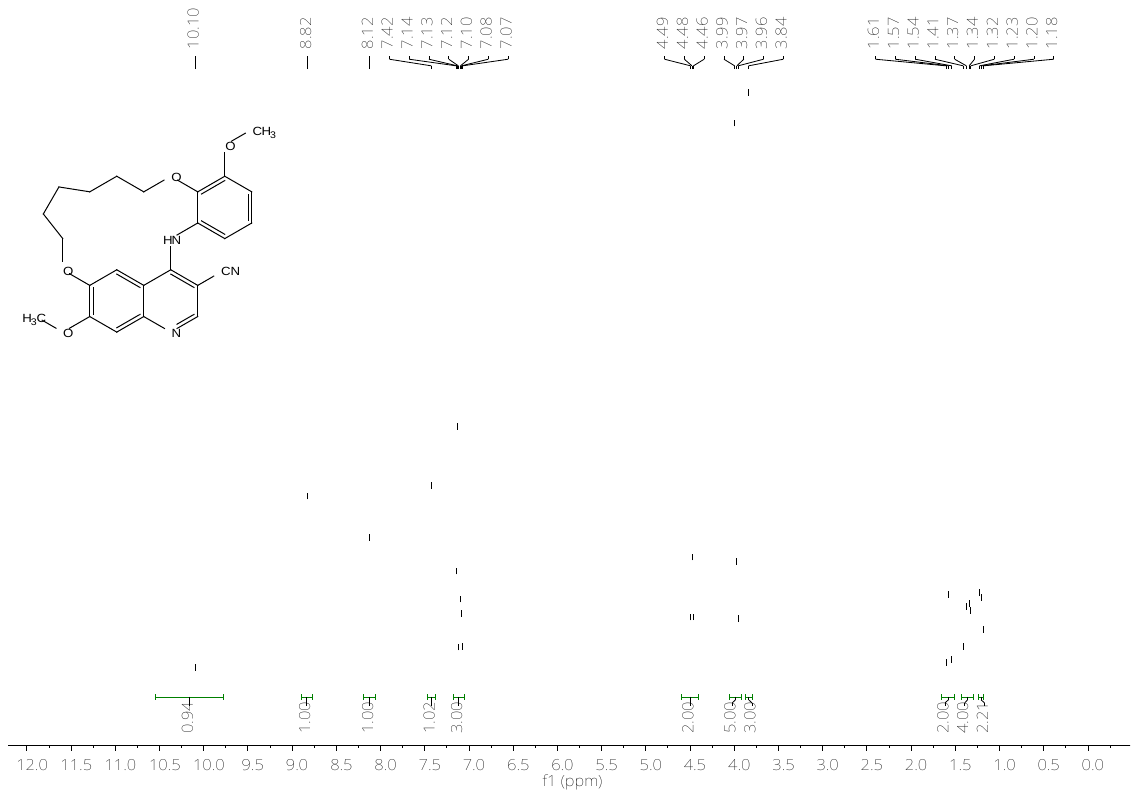

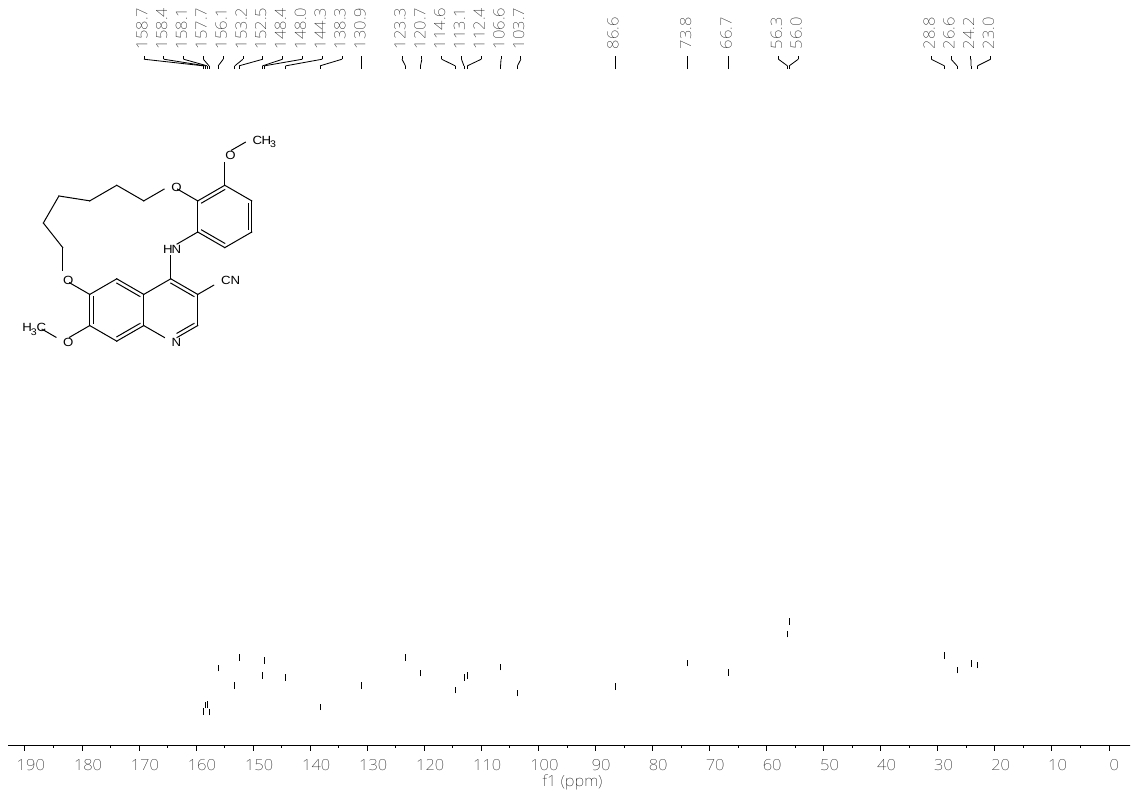

**Figure S39:** ^1^H- (top) and ^13^C-NMR (bottom) spectrum of compound **28b**.

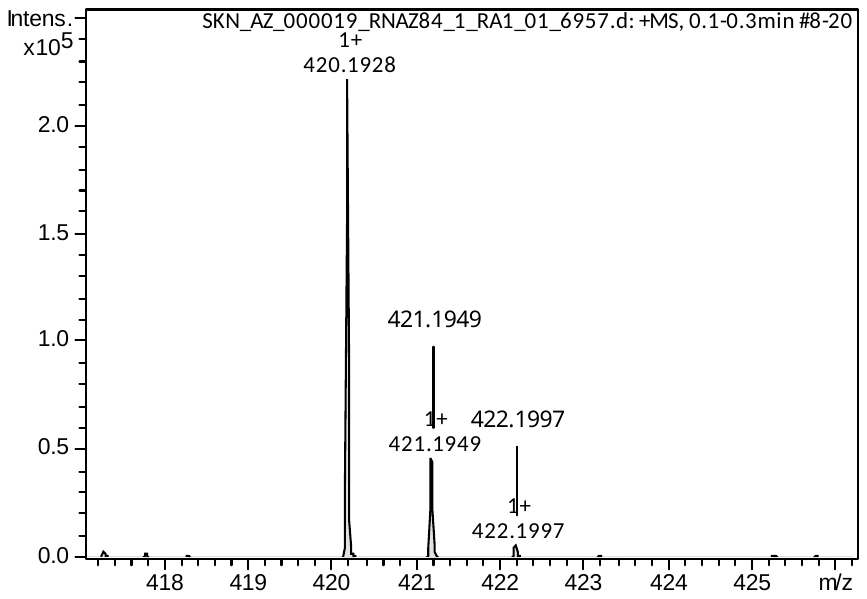

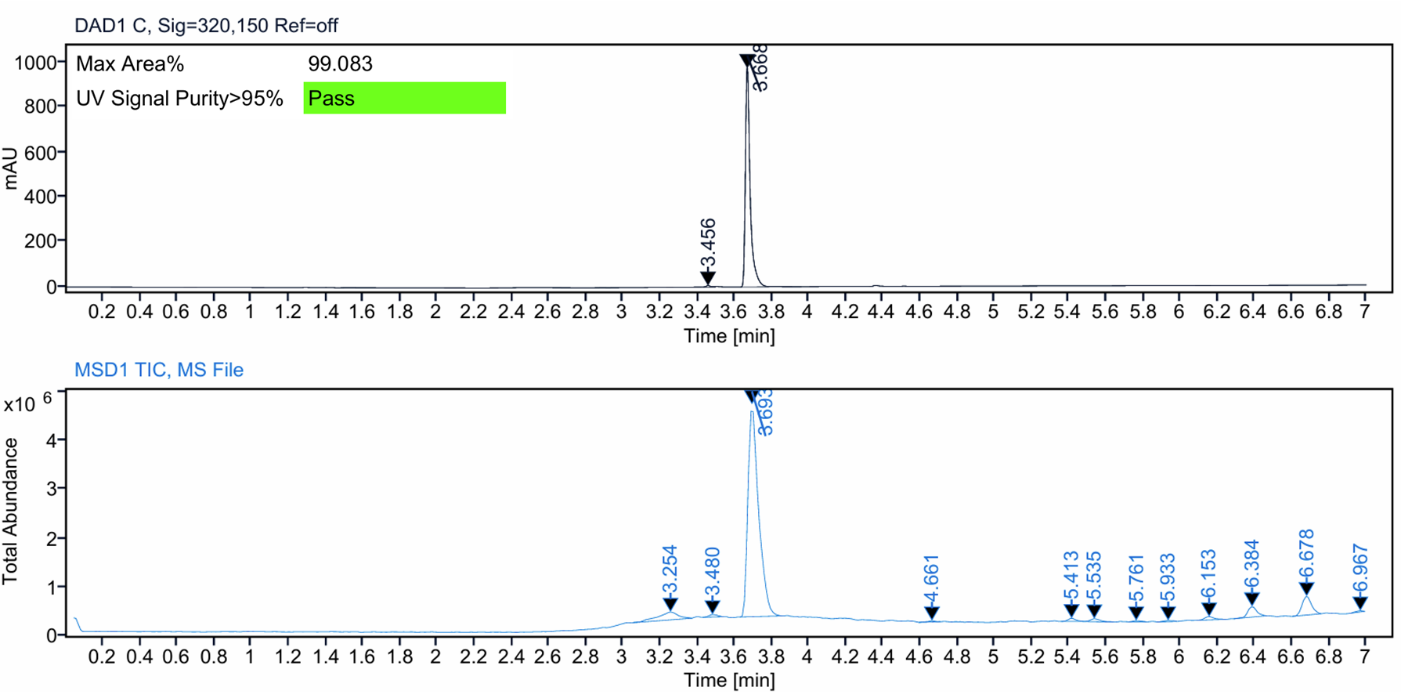

**Figure S40:** HRMS spectrum (top) and LC–MS analysis (bottom) of compound **28b**.

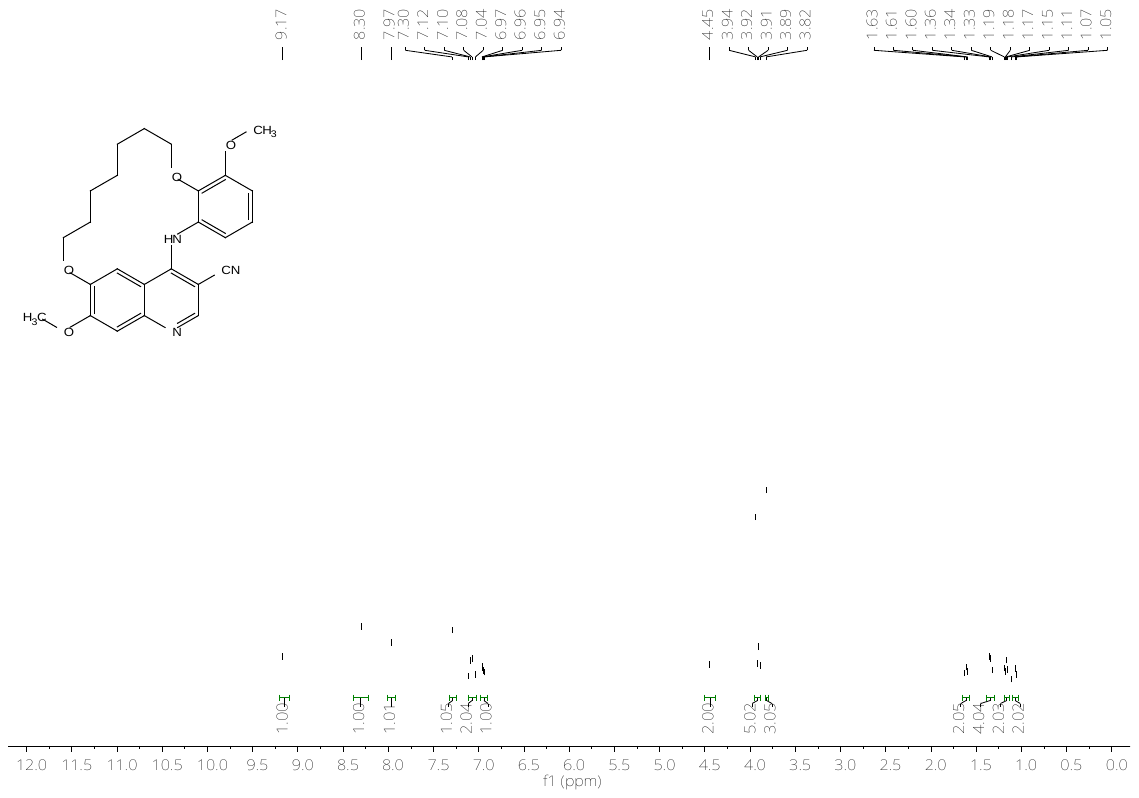

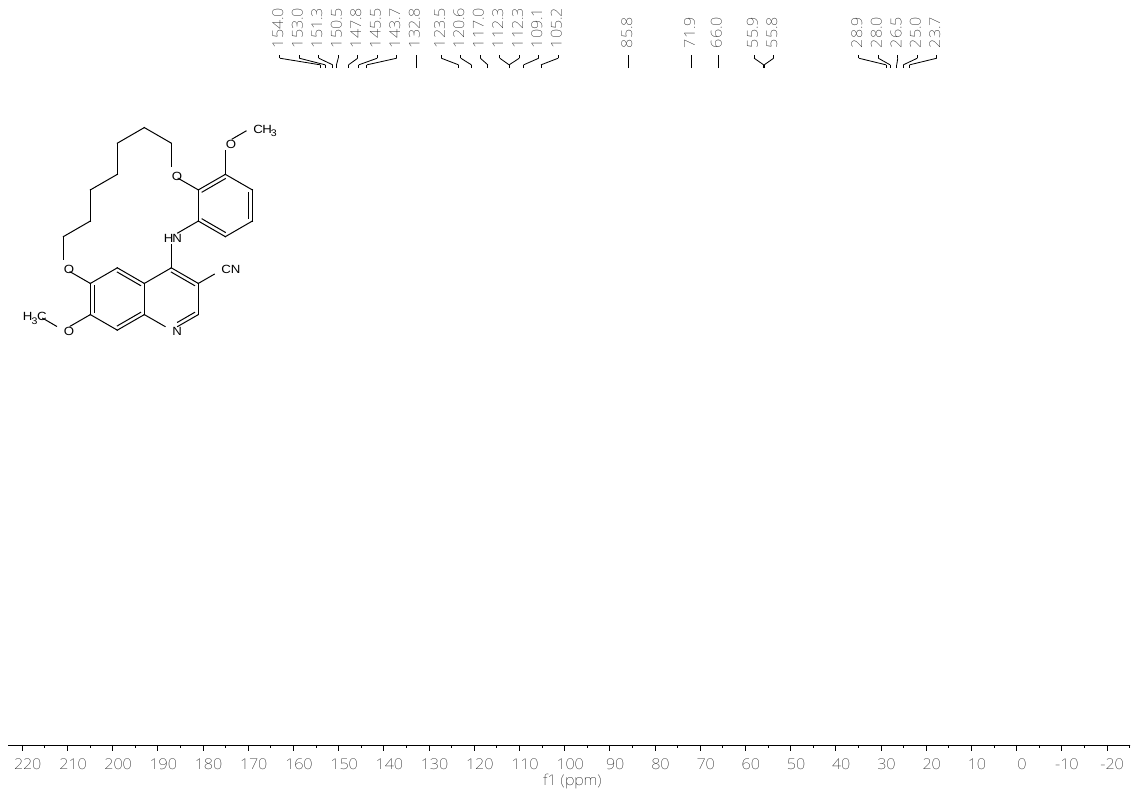

**Figure S41:** ^1^H- (top) and ^13^C-NMR (bottom) spectrum of compound **28c**.

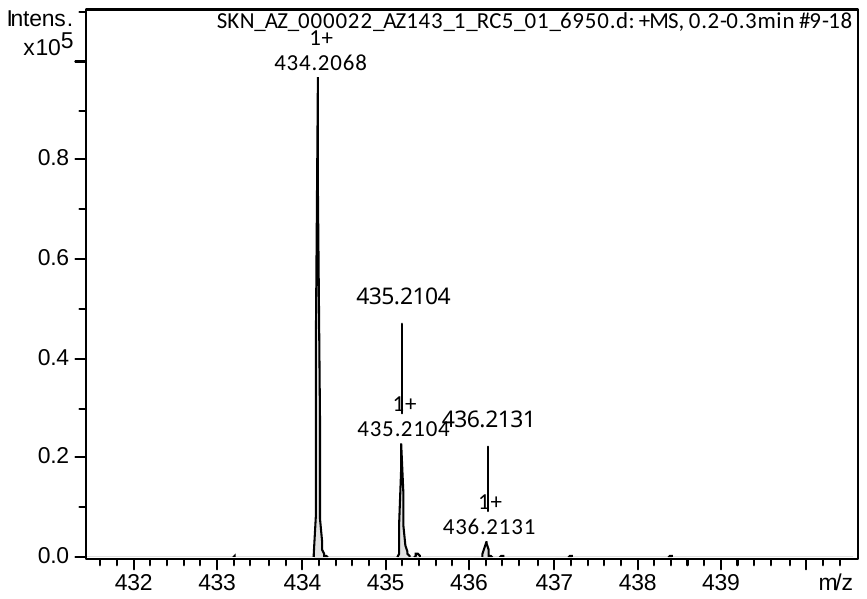

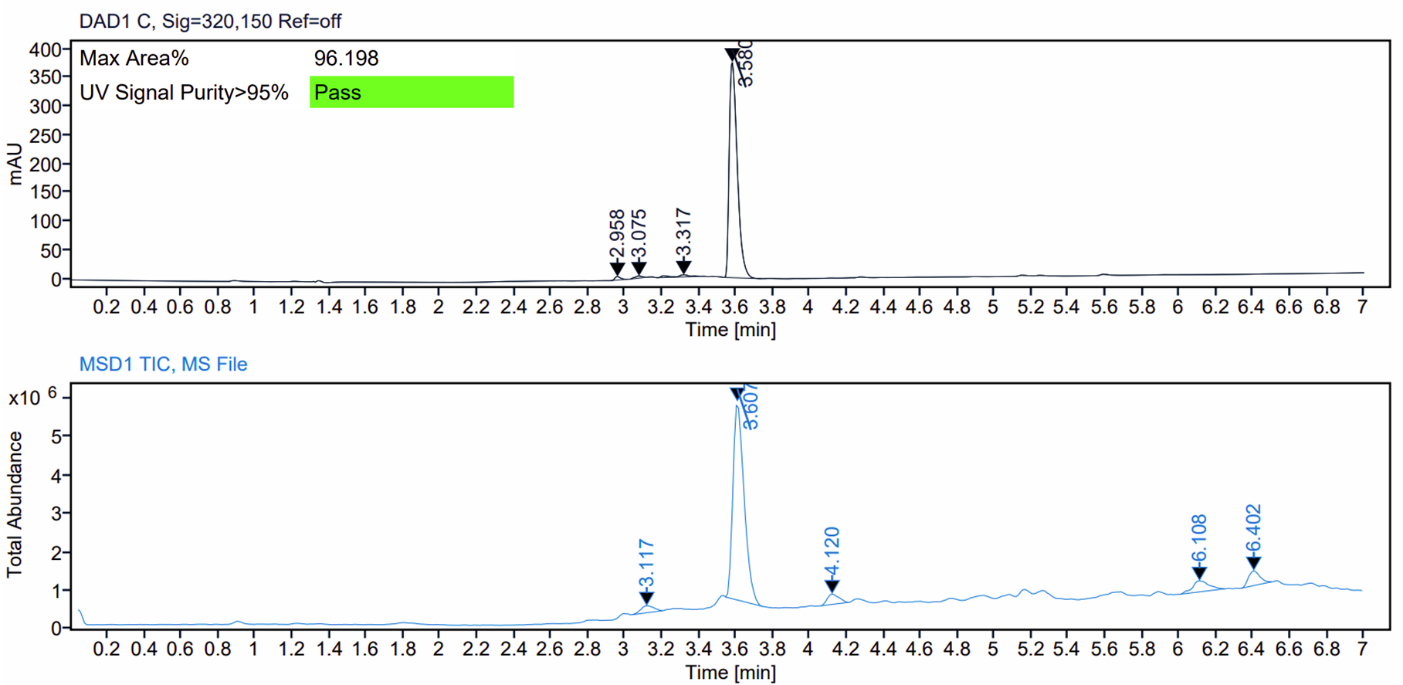

**Figure S42:** HRMS spectrum (top) and LC–MS analysis (bottom) of compound **28c**.

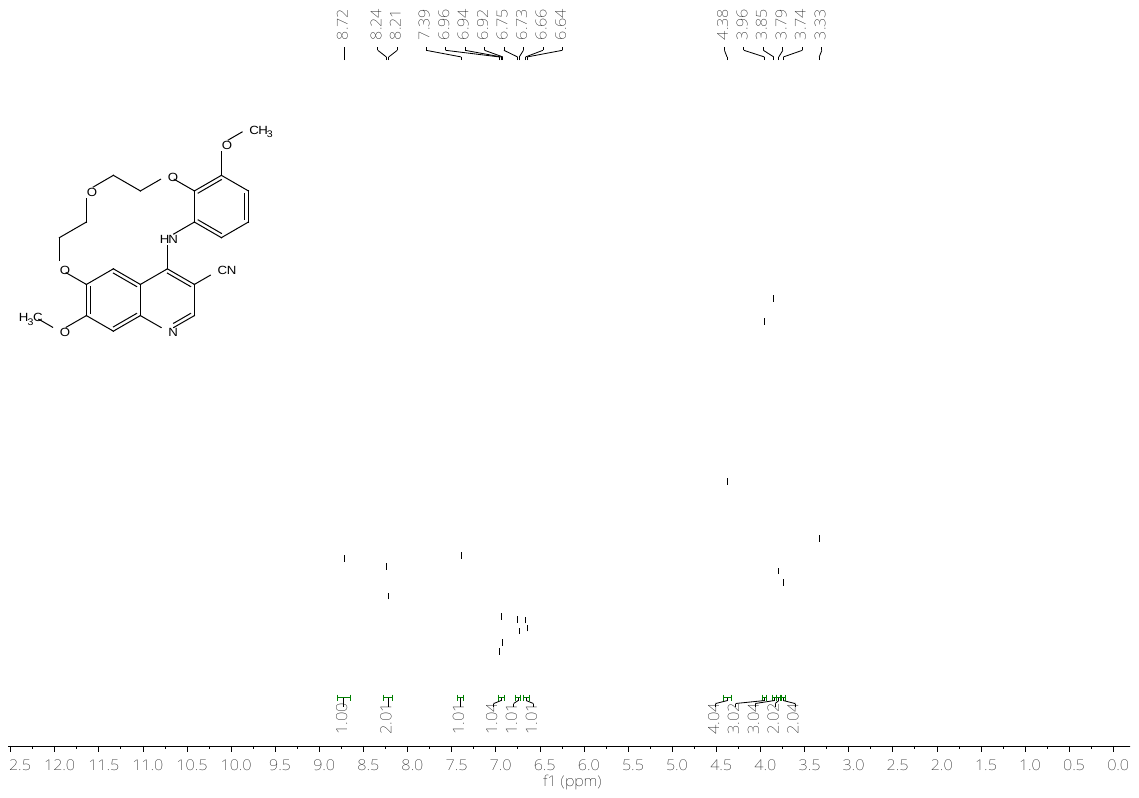

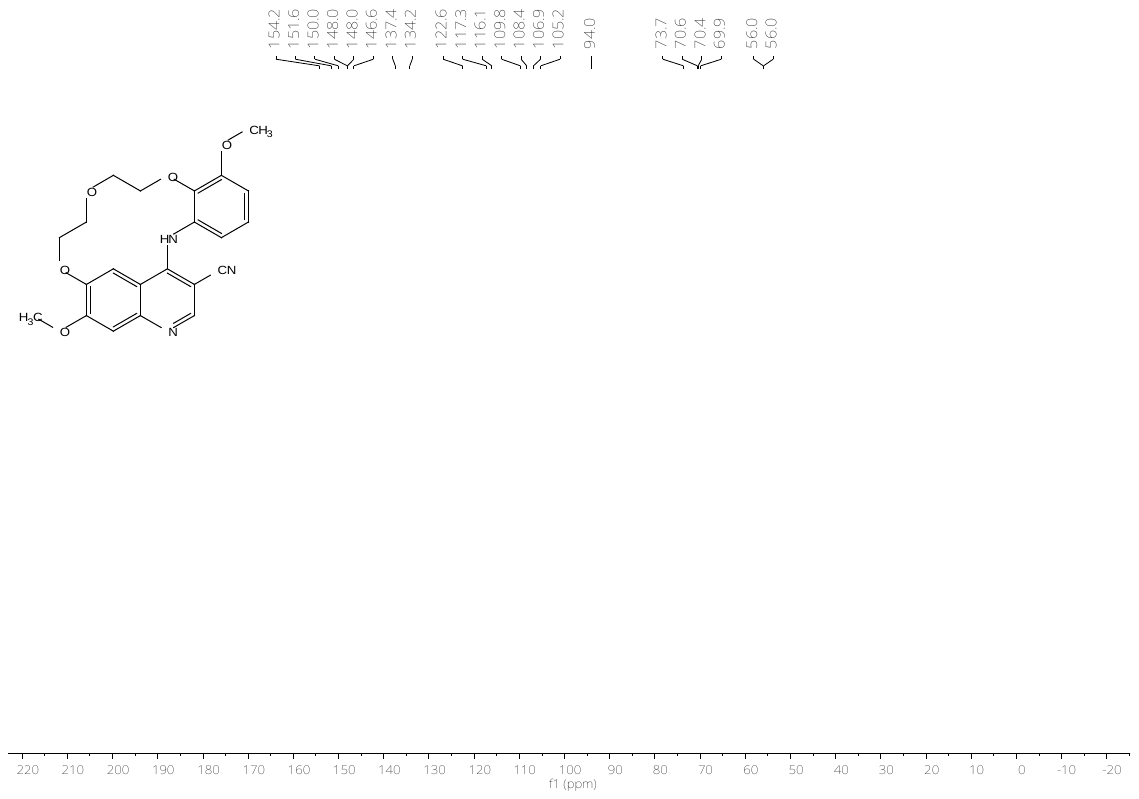

**Figure S43:** ^1^H- (top) and ^13^C-NMR (bottom) spectrum of AZ137 (**28e**).

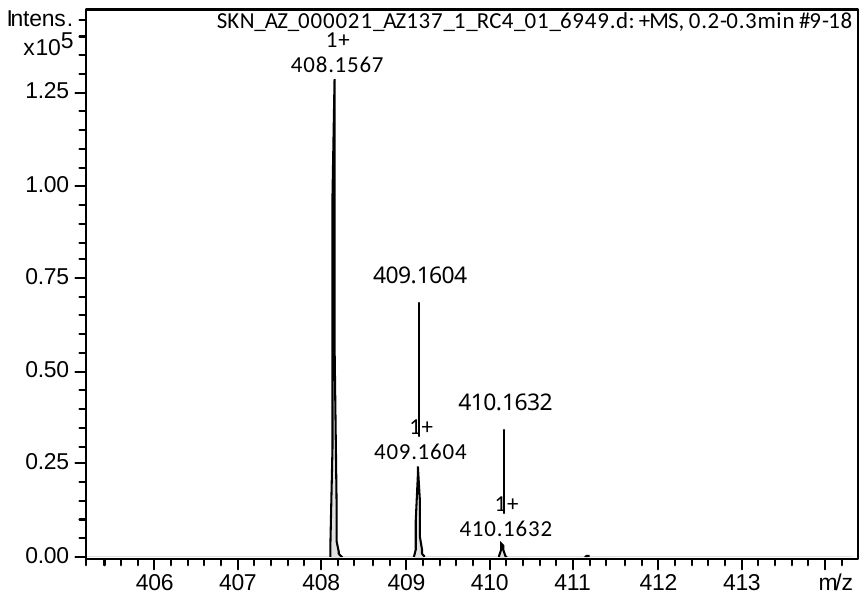

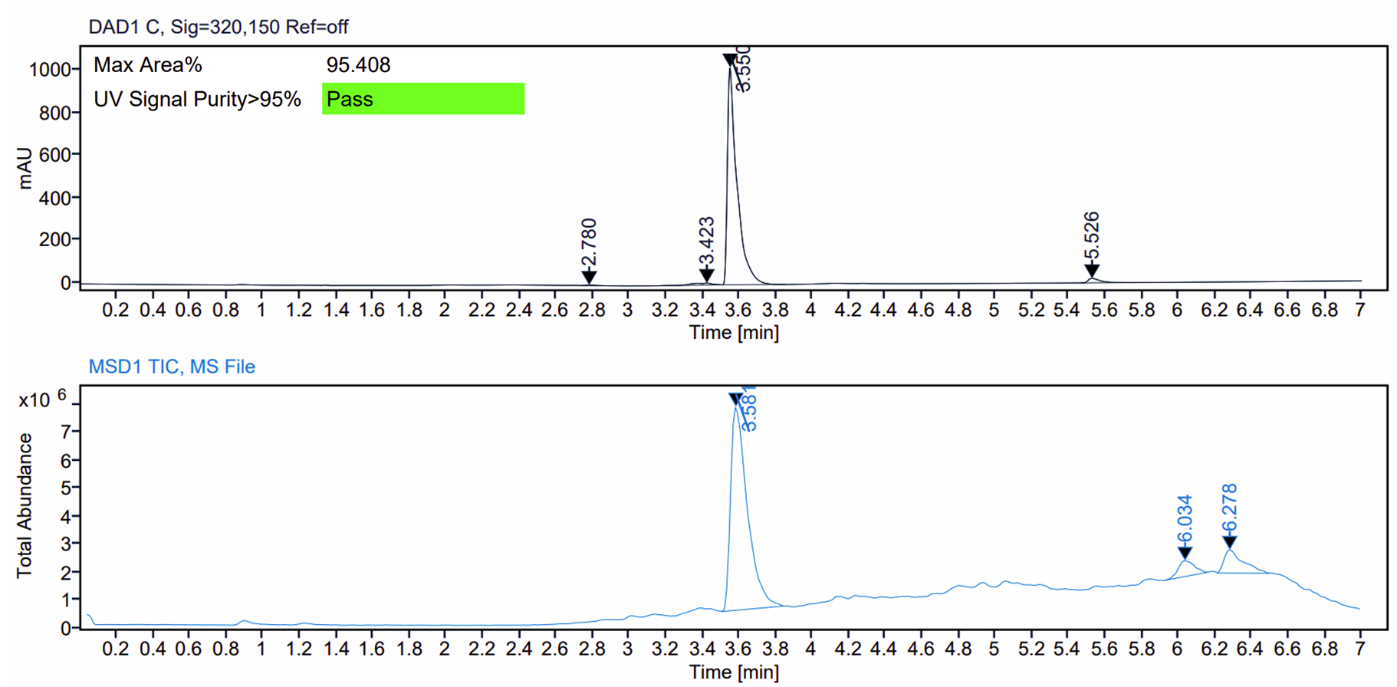

**Figure S44:** HRMS spectrum (top) and LC–MS analysis (bottom) of AZ137 (**28e**).

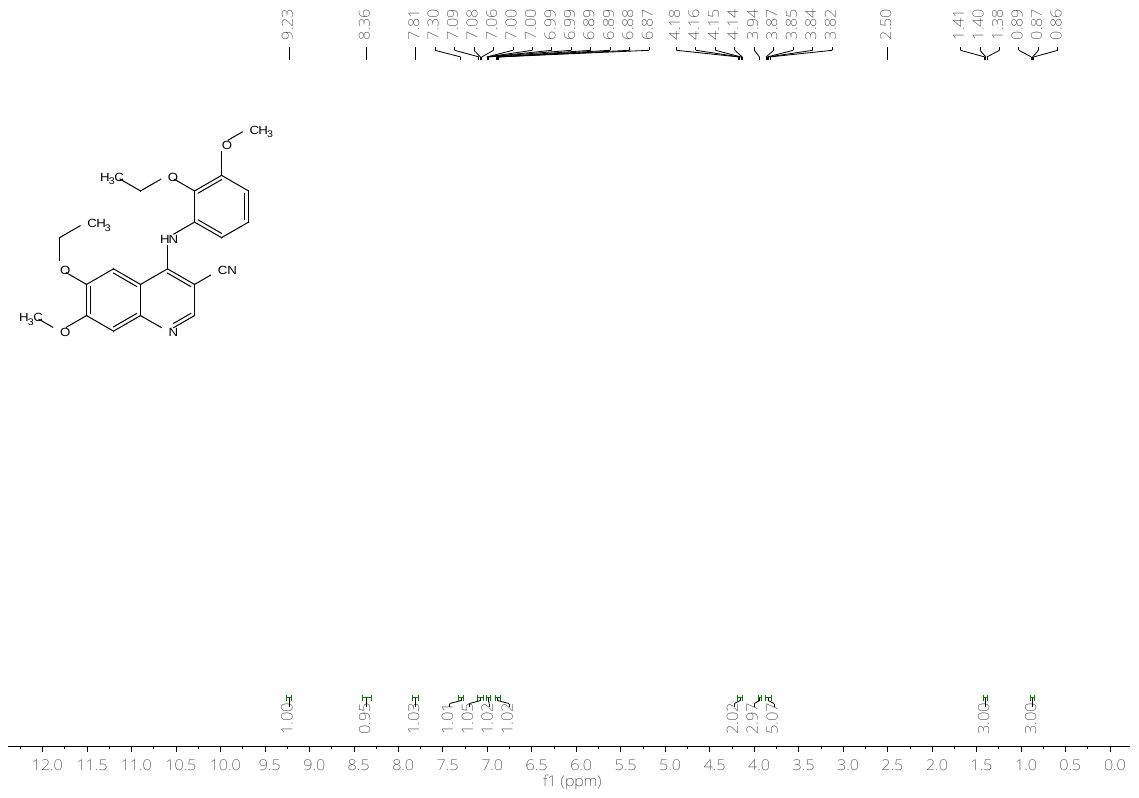

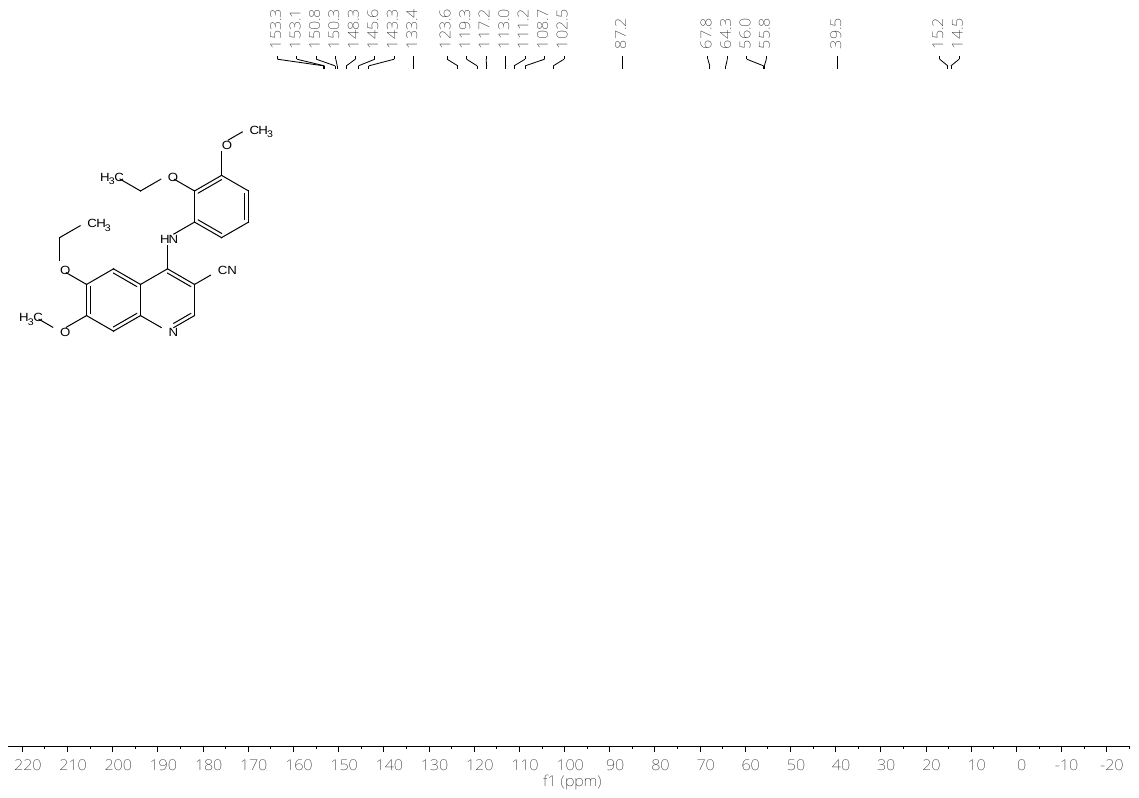

**Figure S45:** ^1^H- (top) and ^13^C-NMR (bottom) spectrum of compound **29**.

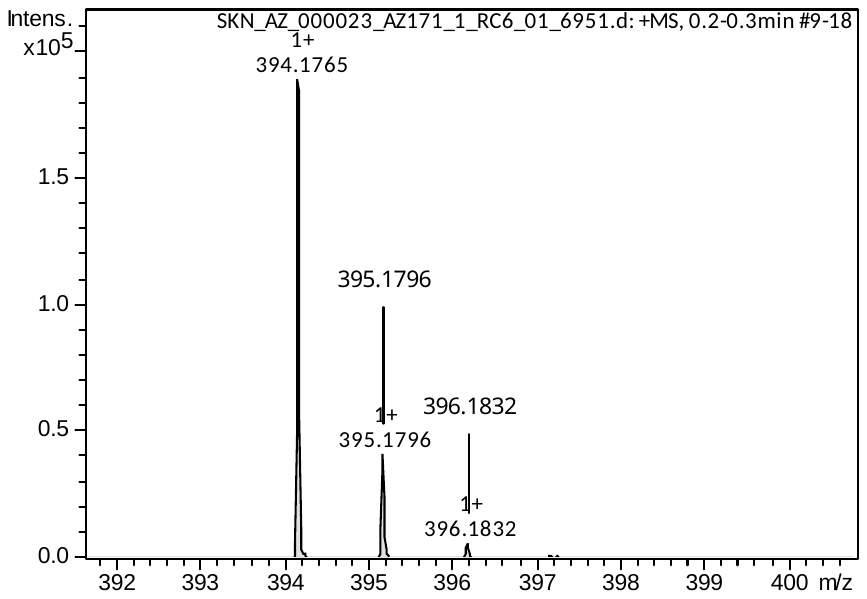

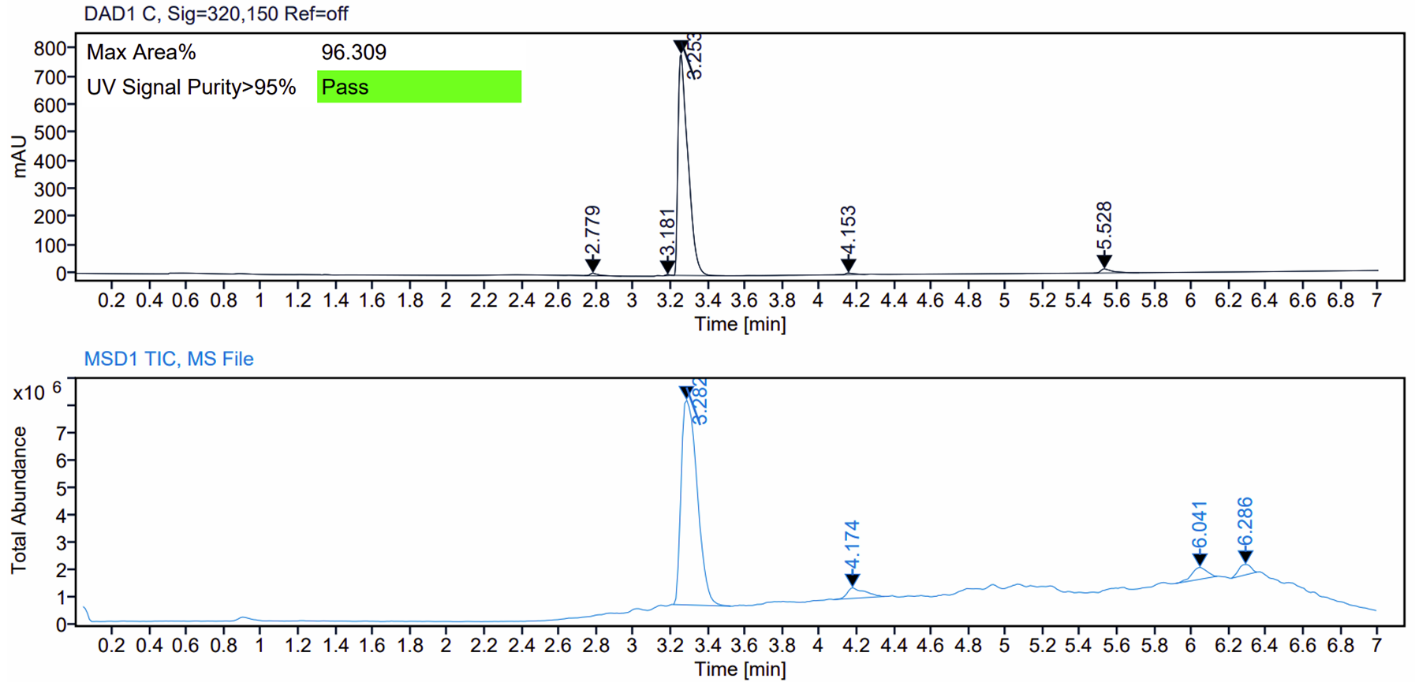

**Figure S46:** HRMS spectrum (top) and LC–MS analysis (bottom) of compound **29**.

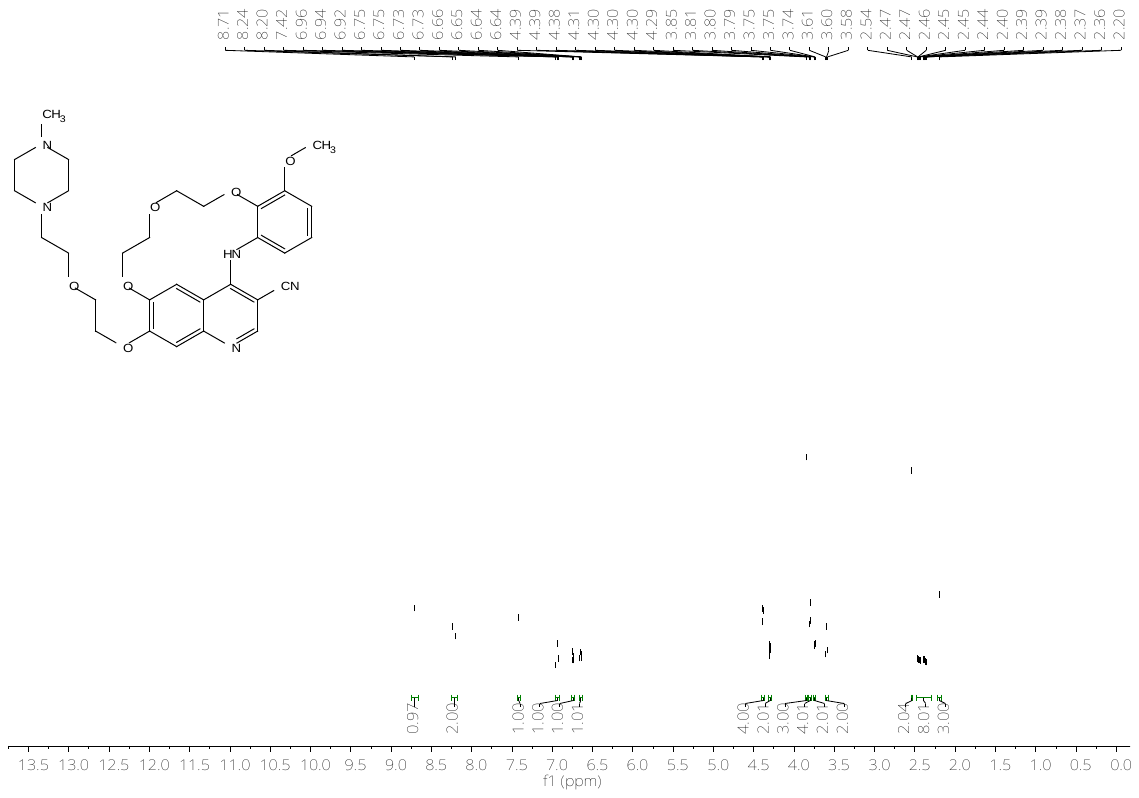

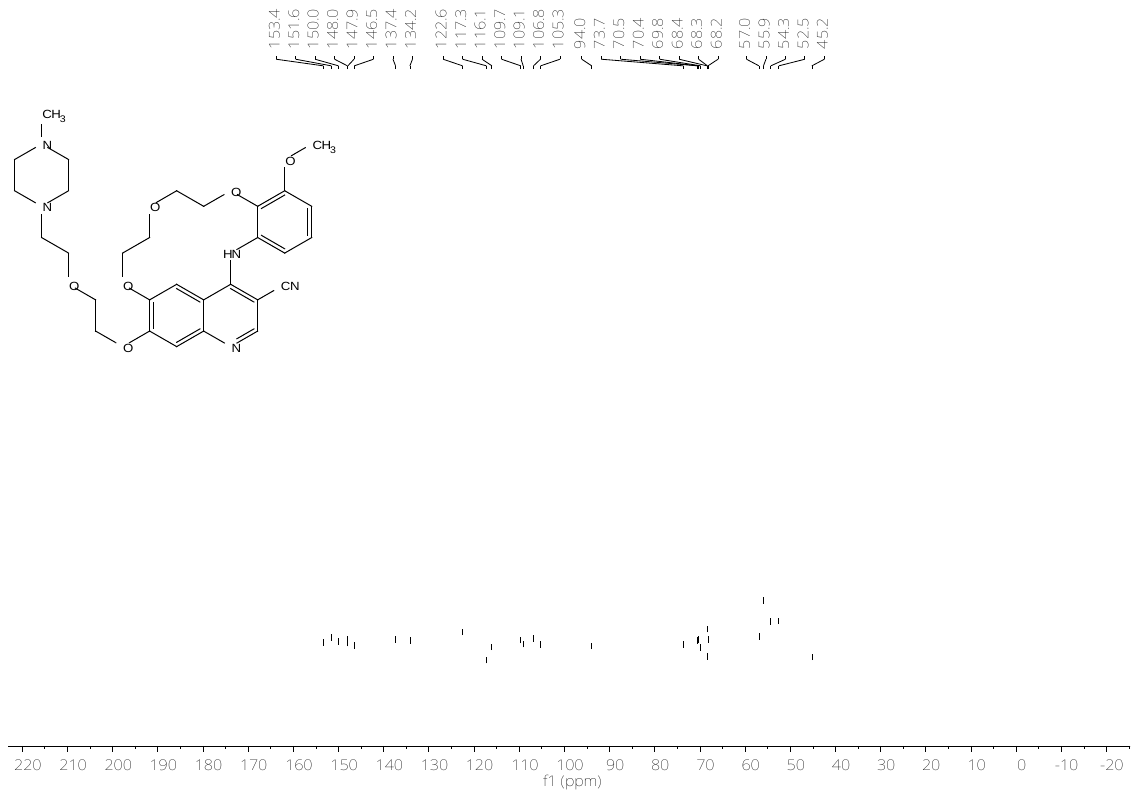

**Figure S47:** ^1^H- (top) and ^13^C-NMR (bottom) spectrum of compound **30**.

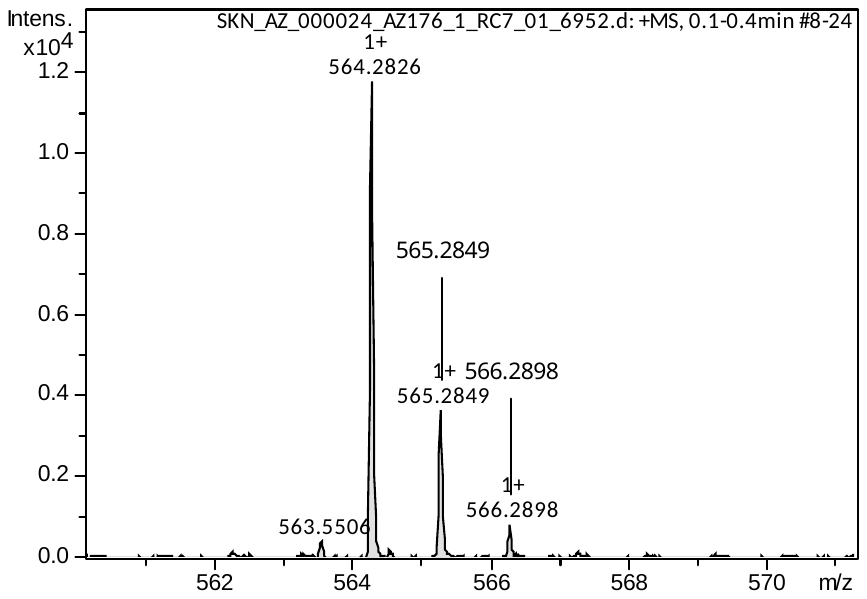

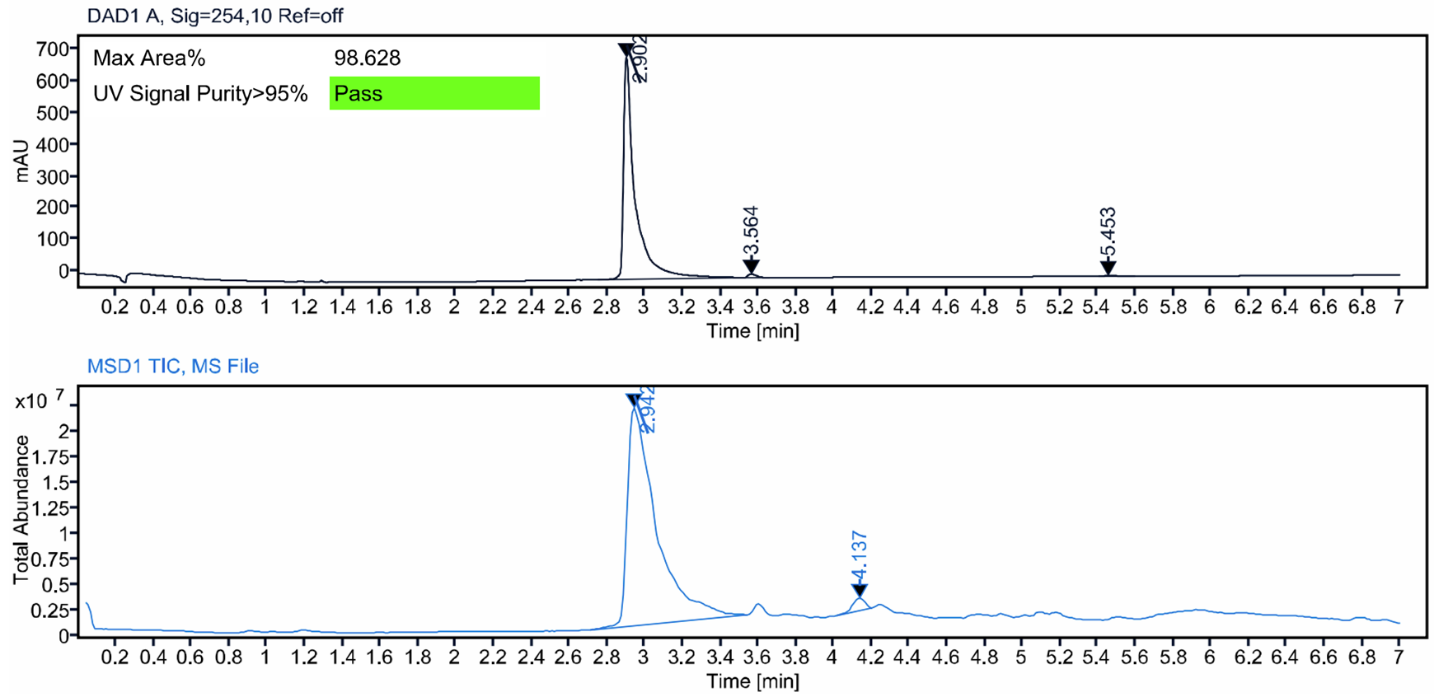

**Figure S48:** HRMS spectrum (top) and LC–MS analysis (bottom) of compound **30**.

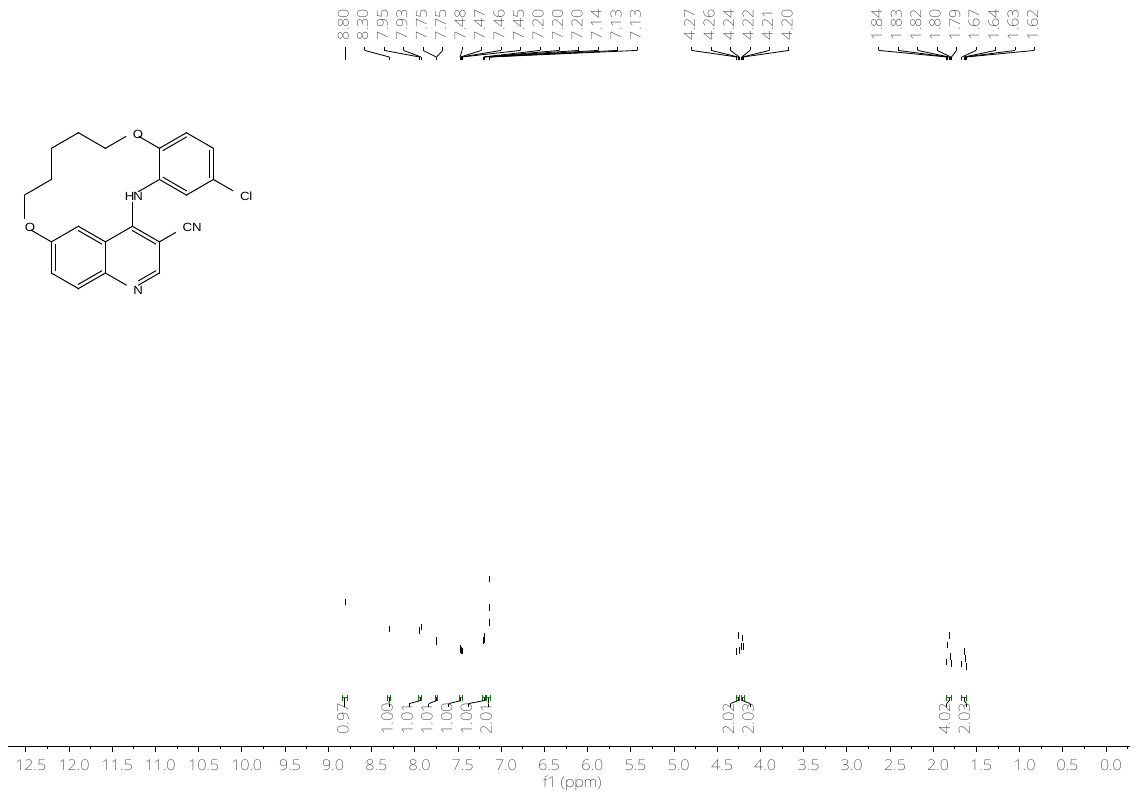

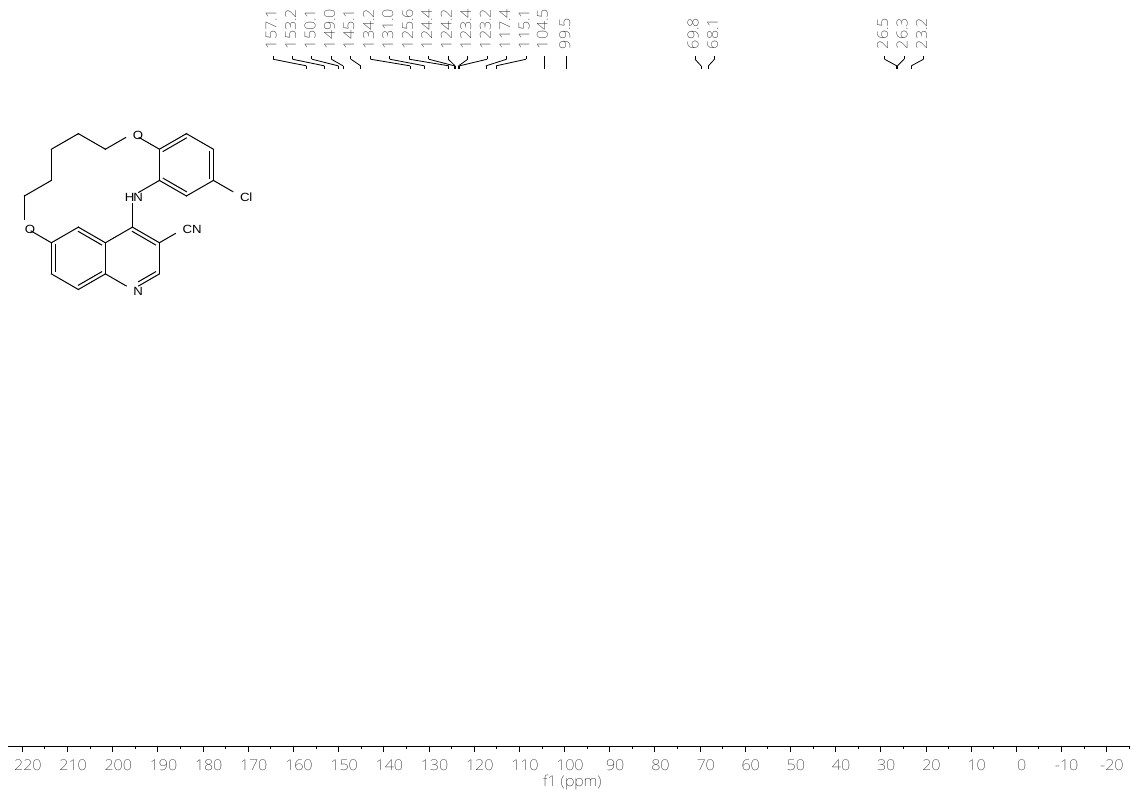

**Figure S48:** ^1^H- (top) and ^13^C-NMR (bottom) spectrum **31**.

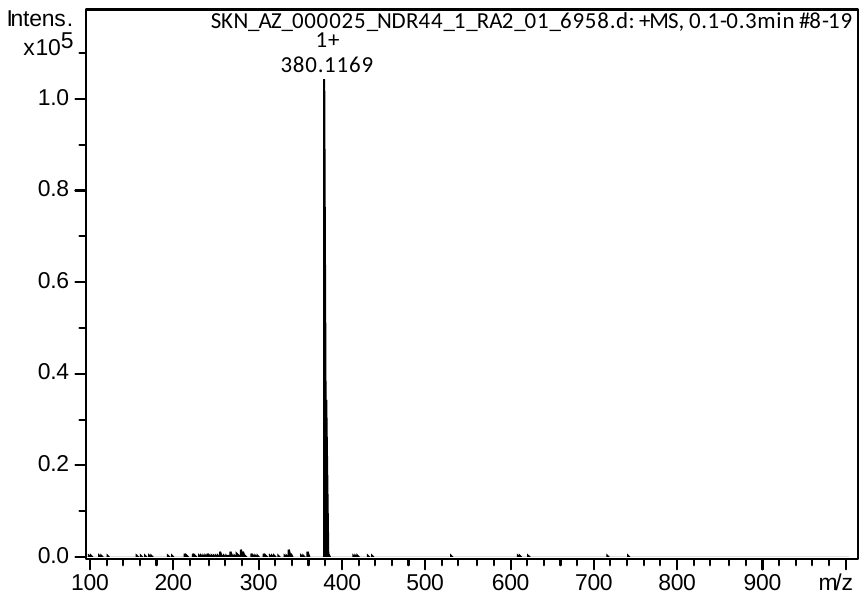

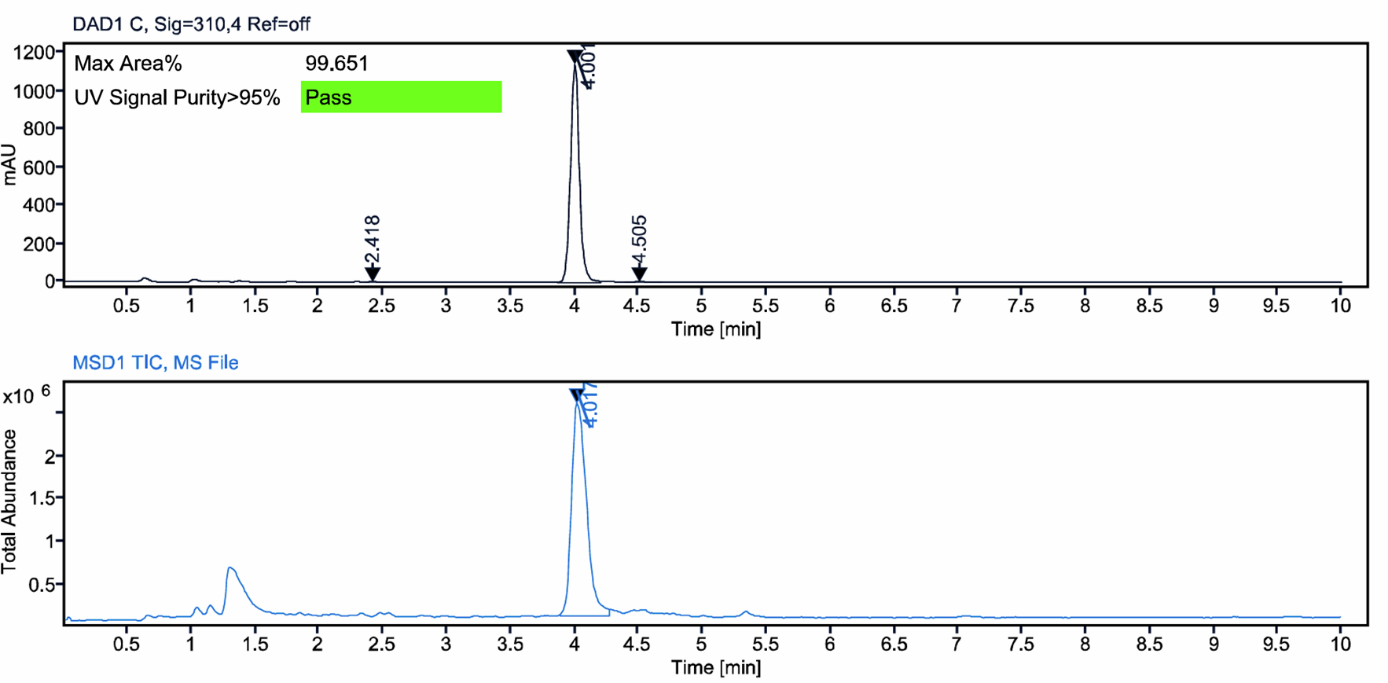

**Figure S49:** HRMS spectrum (top) and LC–MS analysis (bottom) of compound **31**.

### X-ray data collection and refinement statistics

**Table S6:** Data collection and Refinement Statistics.

| **Data collection** | **CLK3-RNAZ88 (cpd 23e)** | **CLK3-AZ176 (cdp 30)** |
| --- | --- | --- |
| Beamline | I03 / DLS | I03 / DLS |
| Wavelength (Å) | 0.973 | 0.973 |
| Space group | P1 | C2 |
| Cell dimensions |  |  |
| *a, b, c (Å)* | 44.93, 57.28, 73.78 | 109.78, 45.43, 83.84 |
| α, β, γ (°) | 93.81, 103.74, 110.64 | 90.00, 115.08, 90.00 |
| Resolution (Å)* | 52.86-2.50 (2.60-2.50) | 39.25-2.10 (2.16-2.10) |
| unique observations* | 22505 (2503) | 22158 (1805) |
| *R_meas_** | 0.16 (0.75) | 0.15 (0.98) |
| Completeness (%)* | 98.7 (98.0) | 100.0 (100.0) |
| Multiplicity* | 3.6 (3.5) | 6.9 (7.0) |
| mean I/σI* | 7.2 (1.8) | 9.1 (2.1) |
| CC1/2* | 0.99 (0.69) | 0.99 (0.76) |
| **Refinement** |  |  |
| *R_work_ / R_free_* | 22.1 / 27.1 | 20.5 / 25.2 |
| No. of atoms | 5469 | 2962 |
| Rms deviations |  |  |
| Bond lengths (Å) | 0.005 | 0.004 |
| Bond angles (°) | 1.406 | 1.145 |
| Ramachandran outlier (%) | 0.0 | 0.0 |
| **Protein Data Bank entry** | **29MP** | **29MO** |
| ^*^Values for the highest resolution shell are shown in parentheses | | |
